# ‘Phytopathological strolls’ in the dual context of COVID-19 lockdown and IYPH2020: transforming constraints into an opportunity for public education about plant pathogens

**DOI:** 10.1101/2021.02.03.429201

**Authors:** Frédéric Suffert, Muriel Suffert

## Abstract

The experience presented here relates to 2020, a particularly timely year for plant disease-related communication (‘International Year of Plant Health’ IYPH2020), but also a unique year because of the COVID-19 pandemic. Our goal was to illustrate the diversity and beauty of fungal plant pathogens through a naturalist approach that could be followed by any amateur. We achieved this end through ‘phytopathological strolls’, in which we observed and determined the origin of symptoms on diseased plants found in our garden, in the local streets, in nearby open spaces, and sharing this matter with a broad public. The lockdown imposed in France created an additional motivation to take up the challenge, and to involve our children, even under strong constraints, such as movement restrictions. We observed and described fungal pathogens through hundreds of photographs, shared our findings with a large audience on Twitter, and received feedback. The material used was deliberately simple and transportable: a digital reflex camera, an old microscope, a mobile phone, some books and an Internet connexion. Between March 17, 2020 and June 20, 2021 we found 196 plant pathogens, including 97 rusts, 27 powdery mildews and 28 septoria-like diseases. We discuss here the importance of promoting searches for plant pathogens, their description and conservation, through a combination of classical approaches and digital tools in tune with the times, such as Twitter, by treating pathogen identification like a detective game and, more surprisingly, by making use of the addictive nature of collection approaches, drawing a parallel with Pokémon Go.

## Introduction

Plant pathogens are harmful to crops and have, over the centuries, caused famines, ruined economies, and blighted landscapes. However, looking at these pathogens from a naturalist viewpoint, rather than in an agricultural context, we wondered whether it was possible to consider plant diseases, especially those caused by fungi, differently, given their rarity, diversity, and even beauty. This attitude is at odds with the perception of plant diseases among farmers and most plant pathologists, who are not used to approaching this subject from this angle. Like most plant pathologists, we typically start our scientific articles by describing the severity of crop losses and justifying our choice of study model as one of the principal diseases responsible for such losses. However, we also use to be struck by the “beauty” of some of the symptoms we observe, even if we generally prefer not to point this out to farmers (for whom these “beautiful” symptoms may spell financial disaster due to crop losses) or to friends and family unlikely to share our enthusiasm. As a result, the enjoyment of plant diseases is a somewhat solitary pleasure that can be shared only with colleagues, and potentially with partners or, more rarely, with children. We wonder whether, as plant health professionals, it might be possible to adjust our perspective and adopt a naturalistic approach not only for the overall surveillance of pests, but also to observe and share the beauty of fungal plant diseases. The second part of this question may appear frivolous in the context of a scientific journal such as *Plant Pathology*, but we are convinced that this aspect deserves greater attention, if only to improve the communication of our discipline to a broader public by a means other than cutting-edge- or technological science. Improving our observation of our environment and getting to know it better could be highly beneficial in a number of ways.

The ‘One Health’ paradigm emerged in the context of the increasing domination of our planet by human activities and is highly influential at the moment. This paradigm, the theme of the current *Plant Pathology* special issue, renders the concept of health more inclusive, by considering humans, domestic and wild animals, cultivated and wild plants to be indissociable (Cumming & Cumming, 2015). Fungal diseases of plants are a particularly important component of One Health, as they can contribute directly to the degradation of agrosystems (Desprez-Loustau *et al.,* 2007; Fisher *et al*., 2012), but also to the emergence of outbreaks with major but indirect consequences for animal and human health (Konopka et al., 2019). It is, thus, becoming increasingly important for pathologists to raise the awareness of policymakers concerning established and emerging fungal threats to food security and global health. However, causing fear is rarely a helpful strategy for communication. Admiring a human or animal disease would not be ethical, and, even if it were, it would be impossible to share this admiration with a wide audience from a purely naturalistic standpoint. Medical and veterinary doctors generally care for individual humans or animals. By contrast, ‘plant doctors’ rarely care for single plants (except for certain heritage trees or pot plants). Instead, they aim to improve the health of a whole plant population at the field or ecosystem scale. This makes it possible for everyone to observe single diseased plants in familiar places without apprehension or disgust, for the purposes of education or to inspire new vocations.

Plant health can be explored distinctively by a naturalist or normativist approach (Döring et al., 2012), if the observer is a farmer or an amateur botanist, for example. The naturalist concept of plant health proposed by Boorse (1977) was described by Döring et al. (2012) as a “circular argumentation” because it defines health as an absence of disease, whilst defining disease as being “inconsistent with health”. Plant diseases are often visible to the naked eye, even though they are caused by micro-organisms, and the symptoms may be ‘beautiful’ in the eyes of an amateur naturalist. Conversely, according to normativism, all measurements of a disease, including those obtained through biological sciences, have a cultural dimension and cannot, therefore, ever be completely objective. As a consequence, the search for diseases during strolls, as proposed here, is biased, because it results from an arbitrary choice: looking at a plant with flagrant symptoms rather than a plant that appears normal but has been physiologically weakened by disease, or being attracted to rare symptoms (not seen elsewhere) rather than common ones.

The United Nations General Assembly declared 2020 as the International Year of Plant Health (IYPH2020), with various objectives, including raising public awareness in a massive manner as concerns the protection of the environment and facilitation of economic and trade development, and the promotion and strengthening of global, regional and national plant health efforts (Routray, 2020). The objective of the experience reported here is consistent with the objectives of IYPH2020, namely informing, educating and engaging audiences through the sharing of knowledge about fungal pathogens on digital social media.

Our goal was to illustrate the diversity and beauty of fungal plant pathogens through a naturalist approach that could be followed by any amateur in any place colonised by plants, whether in a natural, rural or urban area. We used a strategy based on local ‘phytopathological strolls’, in which we observed and determined the origin of symptoms on diseased plants, sharing our exploits with the broad general public, from the scale of our family to a global social network, through the popular microblogging social media platform Twitter. The approach was simple: observe, describe through photographs, identify with simple means, share with a large audience, and receive feedback. This experience was initiated in 2020, a particularly timely year for communicating about plant diseases (IYPH2020) embracing the One Health paradigm, and very unusual because of the COVID-19 pandemic. The lockdown implemented to control this pandemic created a new motivation in our lives to take up this challenge under the imposed constraints, such as travel restrictions. We were convinced that illustrating the diversity and beauty of plant pathogens are would be all the more effective and striking under these uniquely constrained conditions. The problem of ‘collection’ is that its potentially unlimited nature can be daunting. The constraints of lockdown actually resolved this problem by forcing us to confine our efforts to short outings in the close to home.

## Materials and methods

### Context of strolls

There have been three lockdowns so far to counter the COVID-19 epidemic in France. The first extended from March 17 to May 11, 2020 (55 days), the second from October 30 to December 15, 2020 (47 days) and the third from April 3 to May 3, 2021 (28 days). Our ‘phytopathological strolls’ began at about the same time as the first lockdown and were completed some weeks after the end of the third lockdown. During the first lockdown, all the members of our family stayed at home (traditional single-family house with a garden) in Les Clayes-sous-Bois, about 30 km west of Paris (48°49’14’’, 1°59’33’’). We worked from home, the children had online lessons and we went out only to shop for food and to get some fresh air, initially in the nearby streets (as all parks and forests were closed to the public during the first few weeks) and later in open spaces (walks in a small park and in a nearby forest, such outings being limited to a radius of 1 km and a duration of 1 hour), or, exceptionally, to go to the INRAE laboratory and greenhouse at Thiverval-Grignon (48°50’49 N, 1°56’34’’) to deal with scientific and technical emergencies. The second lockdown was less strict. We worked from home two to three days per week, and were allowed to walk for up to three hours at a time, within a radius of 20 km from our home. The third lockdown was quite similar to the second one but we decided to spend it partially in alternative homes in the South-West of France. We saw every stroll as an opportunity to search — in our garden, the street, the urban green spaces, or the INRAE arboretum — for all the plants with visible symptoms on leaves. Between the lockdown periods, constraints were eased to such an extent that travel was possible, so we extended our search to various holiday destinations, such as Millau (44°5’50’’, 3°3’22’’) and Sermizelles (47°32’20, 3°47’33), and to weekend getaways (Figure 1).

**Figure 1.**
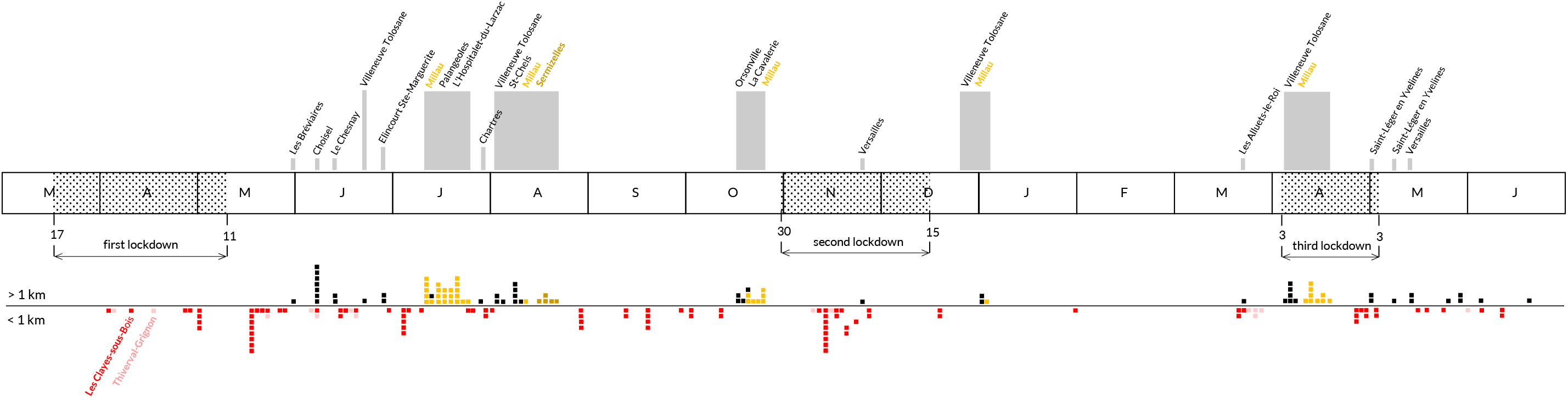
Distribution in time and space of our 2020 ‘phytopathological strolls’ in the context of the COVID-19 pandemic (see also the map in Figure 4). Each square indicates that a plant disease was photographed, collected and identified (first occurrence only). The colour of the square corresponds to the location. Grey rectangles indicate locations accessed after travel and their size are proportional to the distance from our home and to the duration of our stay. The prospecting perimeter around our home (Les Clayes-sous-Bois) and the INRAE workplace (Thiverval-Grignon) corresponded to the perimeter authorized during the first lockdown (< 1 km).

## Materials

We deliberately chose to use simple and transportable material within the reach of any amateur naturalist. During our communications, we made it clear that the various observations and identifications could be made by anyone looking at a symptomatic plant, without the need for specialist analytical techniques. The ‘old-school’ material used (Figure 2A) probably gave a cliché image of what it is like to be a plant pathologist in 2020. In *natura* photographs were taken with a Canon EO7D digital single-lens reflex camera with a 100 mm Macro lens (Canon Inc., 2012) and a Panasonic Lumix Dmc-Fx37 digital camera (Panasonic Corporation, 2008). Microscopy preparations were classically wet mounts, in which fine slices of symptomatic tissues were placed in a drop of water with some methylene blue held between a glass slide and a coverslip. We used a pin and a scalpel to prepare the slides, and a white ceramic tile was used as a transportable laboratory bench. Microscopy observations were performed with an old SM-M Leitz binocular microscope with a black enamel finish (Serial number 670103, circa 1965; Ernst Leitz GmbH, Wetzlar, Germany) operating with four lenses: x3.5, x10, x40, x100, and two eyepieces: x10, x1.25 (Overney & Overnay, 2008). Photographs of the microscope slides were taken with a Samsung Galaxy A40 (Samsung Electronics, 2019) mobile phone held by hand directly above the eyepiece of the microscope. The microscopy slides were sealed with nail polish and kept in a dedicated box. Diseased plant samples were pressed and protected with paper to build up a small herbarium.

**Figure 2.**
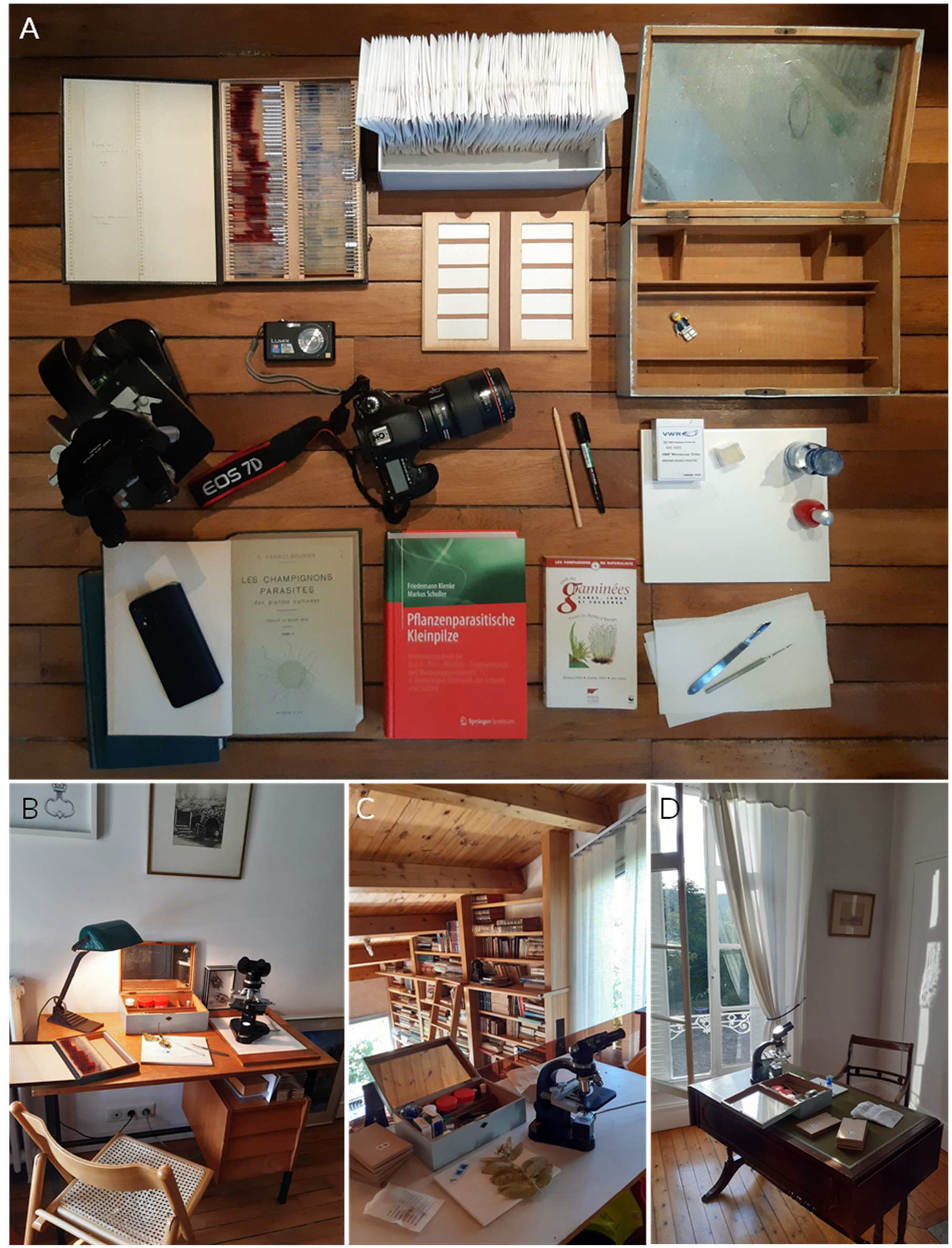
(A) Material used at home and microscopic observation areas for our 2020 ‘phytopathological strolls’: (B) Les Clayes-sous-Bois (Yvelines), (C) Millau (Aveyron) and (D) Sermizelles (Yonne).

We deliberately limited the bibliographic references used, focusing on two books on plant pathogens (Viennot-Bourgin, 1949; Gäumann, 1959; Klenke & Scholler, 2015) together with articles from a wide range of scientific journals. We used a few websites and blogs written by plant pathologists (e.g. Ellis, 2020; Kruse, 2020; Lechat, 2020; Sothmann, 2020). We also used basic botanical guide books to identify the host plants, particularly for difficult groups, such as Poacea or Carex (e.g. Fitter *et al*., 2016), and endemic species (e.g. Coste, 1887). We also used the Pl@ntNet mobile phone application (The PlantNet Consortium, 2020), which was very useful for confirming identifications during walks. All samples were prepared and identified at a desk, which was not always in the same location — at home in Les Clayes-sous-Bois (Figure 2B), during holidays in Millau (Figure 2C) or in Sermizelles (Figure 2D) — but always in the same pleasant ‘old school’ atmosphere.

### Public social media as a communication channel

The potential of social media is now largely recognized in science, especially in the environmental and ecological fields. The value of non-expert observations has been highlighted by the increasing number of individual initiatives from passionate amateurs or so-called “citizen science” projects accessible via websites or mobile applications (Silvertown, 2009; Daume & Galaz, 2016). Micro-blogging social media platforms for short messages (e.g. Twitter) are also used by many academic scientists as an informal arena for previewing work underway or for amplifying the impact of their publications (Darling et al., 2013). Furthermore, these platforms are appreciated in the day-to-day communication of scholars, as a means of actively engaging with the public and influencing perceptions about science (Ke et al., 2017). Our own experience on Twitter shows that the plant pathologist community is well developed and active. In this context, Twitter was the only tool used to communicate our findings. We used our personal accounts (mainly @wheatpath, but also @MurielSuffert for fuelling and sharing conversations), both created in 2012, with 2725 and 2345 followers, respectively, on June 20, 2021. We published hundreds of photographs of the plant pathogens that we found, presented their characteristics and discussed their identification in messages of limited size (<280 characters). In some cases, we developed ‘threads’ (groups of related tweets) for educational purposes providing more detailed explanations and bibliographic references, particularly during the first lockdown. This mode of communication was exclusive. We deliberately decided not discuss this experience with our INRAE and EPPO coworkers, with the exception of those with an active Twitter account.

## Results

### Photographs and descriptions of plant pathogens discovered during our strolls

We found and photographed pathogens causing 196 diseases: 97 rusts, 27 powdery mildews, 28 septoria-like diseases, 30 other fungal leaf diseases, 8 other fungal non-leaf diseases, 3 hyperparasite fungi and 3 non-fungal diseases (Table S1). These findings clearly demonstrate that it is possible to find a large diversity of plant pathogens in a relatively restricted area (Figure 3 4). An example of a sheet for a pathogen is presented in Figure 5, and all 195 sheets are available from Appendix 1, the core output of this experience. We were able to identify 158 of the 196 pathogens to species level without difficulty (sometimes after misidentification, corrected after posting on Twitter with the help of plant pathology experts and amateurs), and 27 with reasonable doubt. The identification of 12 other pathogens was uncertain, or limited to genus level, as for *Melampsora* sp. on willows, *Puccinia* sp. on sedges, or Erysiphe sp. These identifications are known to be problematic even for specialists (Pei & McCracken, 2005).

**Figure 3.** Illustration of our 2020 ‘phytopathological strolls’ in a familial context. (A) Cover of *Petasites hybridus*, host of *Coleosporium tussilaginis* f. sp. *petasitis* (Les Clayes-sous-Bois, 2020-06-07; Appendix 1, sheet n°3). (B) ‘Beastie the bug’ (@bug_beastie), mascot of EPPO travelling the world to raise awareness on plant health during the IYPH2020, here presented in *Pyrus communis*, host of *Gymnosporangium sabinae* (Millau, 2020-07-12; Appendix 1, sheet n°7). (C) Photograph of *Uromyces trifolii-repentis* on *Trifolium pratense* in the garden of our home (Clayes-sous-Bois, 2020-04-26; Appendix 1, sheet n°96). Practical work session during home schooling with our two youngest children: (D) in situ observation of *Puccinia lagenophorae* on *Senecio vulgaris* (Les Clayes-sous-Bois, 2020-03-29; Appendix 1, sheet no. 58); (E, F) microscopy observation; (G) diagnosis of *Phragmidium rubi-idaei* infection on a leaf of *Rubus idaeus* (Clayes- sous-Bois, 2020-06-05; Appendix 1, sheet no. 27) based on Viennot-Bourgin (1949).

**Figure 4.**
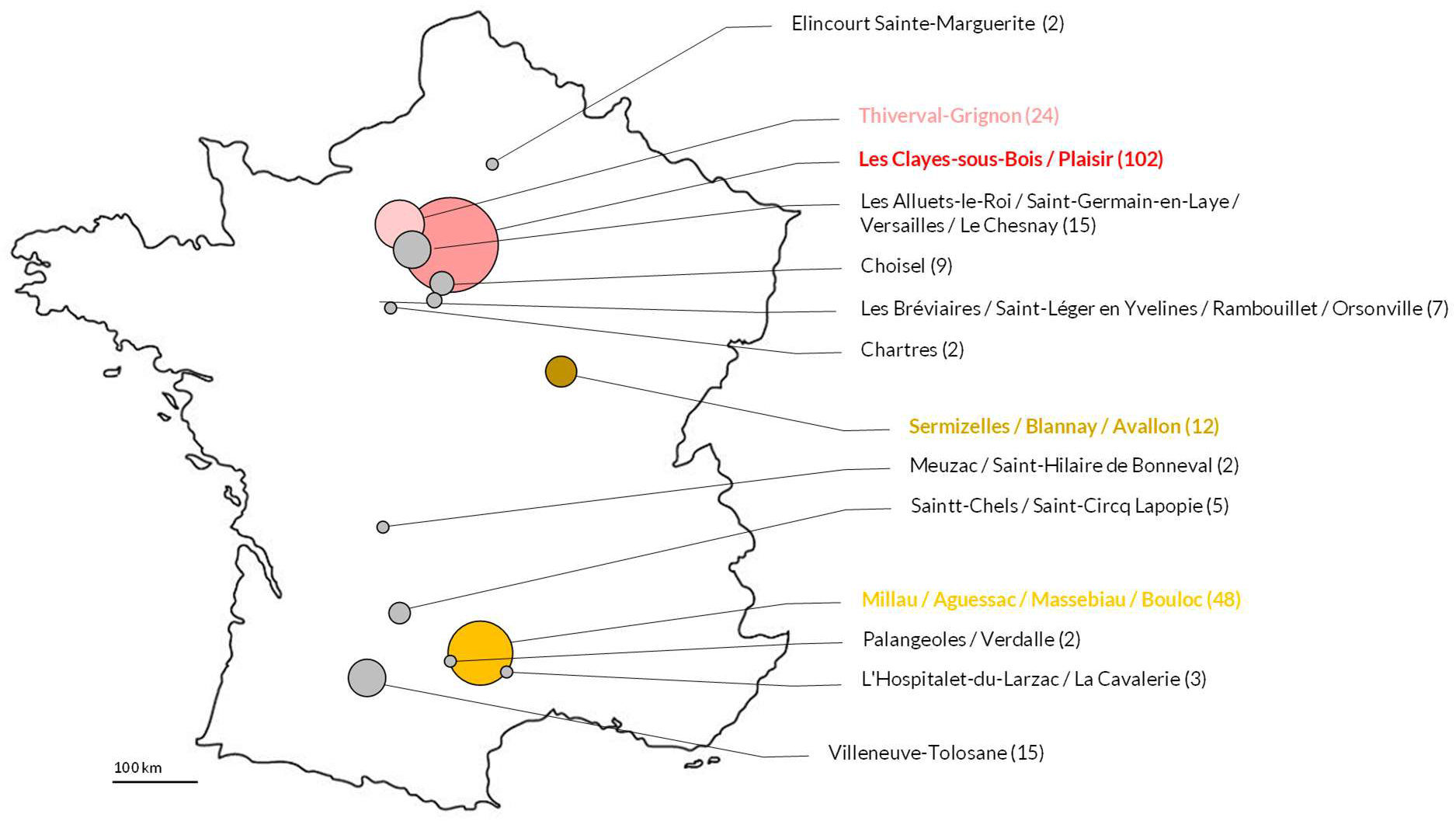
Location of the plant pathogens identified (first occurrence) during our 2020 ‘phytopathological strolls’ in France from March 17, 2020 to June 20, 2021. The size of the circles is proportional to the number of plant pathogens.

**Figure 5.**
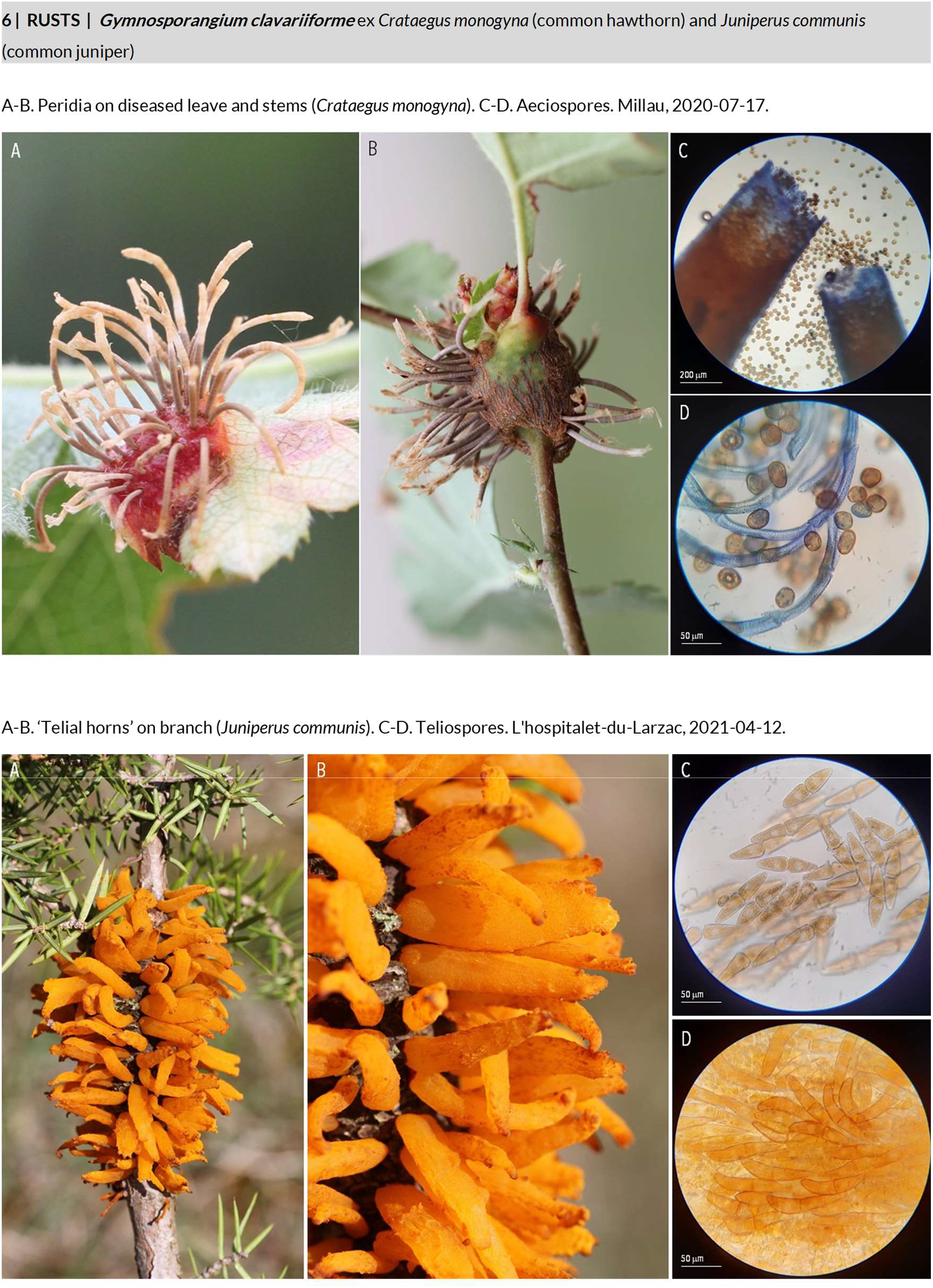
Example of a plant pathogen sheet (*Gymnosporangium clavariiforme* on *Crataegus monogyna* and *Juniperus communis*) from the 196 presented in Appendix 1.

### Some uncommon species

The vast majority of species we found are common in France, but some were quite rare or had never been reported in France or in Europe. For instance, we found *Uromyces plumbarius* on biennial gaura (*Oenothera gaura* syn. *Gaura biennis*) in a flowerbed just in front of our daughter’s school in Les Clayes-sous-Bois (Appendix 1, sheet no. 93). To our knowledge, this is the first time that this species has been reported in Europe (Bisby, 1916; Kern & Whetzel, 1926; Hennen & Cummins, 1967). For confirmation of this observation as a first detection in Europe, this identification would need to be completed by a molecular method (e.g. ITS sequencing), but the likelihood of this identification being confirmed is high, as no rust has ever before been described on biennial gaura in Europe (Klenke & Scholler, 2015). This ornamental species was imported from North America and its identification was certain in this case. We found *Septoria caricis* on specimens of grey sedge (*Carex divulsa* subsp. *divulsa*) growing in the gutter along a street in Villeneuve-Tolosane (Appendix 1, sheet no. 142). The leaves presented irregular spots, elongated parallel to the veins, with pycnidia releasing fusiform pycnidiospores, and morphological characteristics sufficiently distinctive to allow the identification of this fungal species with a high degree of certainty. To our knowledge, this is the first record of this fungal pathogen on this host plant in continental Europe: S. caricis has been reported on Carex sp. throughout the world (Poland, Iceland, Bulgaria, Romania, Uzbekistan, China, USA, Canada) but on *C. divulsa* only in the Canary Islands (Jørstad, 1958) and Turkey (Erdoğdu & Özbek, 2017). We found Blumeriella kerriae on a Japanese marigold bush (*Kerria japonica*) in both an urban horticultural garden in Chartres and a private garden in Villeneuve Tolosane (Appendix 1, sheet no. 157). This pathogen causes leaf spots and was described for the first time in the UK only recently (Stewart, 1917; Robinson et al., 2017). Its identification at two distant sites during our strolls suggests that this species may be widespread in France despite never having been officially reported. We found the widespread pathogen species *Microbotryum dianthorum* on *Dianthus longicaulis* (Appendix 1, sheet no. 185), a rare plant endemic to France, at L’Hospitalet-du-Larzac on the Larzac Causse (Coste, 1887). To our knowledge, this fungus, which causes anther smut disease, has never before been described on this carnation species. We found what appeared to bea secondary fungal pathogen on the leaves of a Japanese spindle tree (*Euonymus japonicus*) infected with the common powdery mildew pathogen *Erysiphe euonymi-japonici* (Appendix 1, sheet no. 104). A recent phyllosphere microbiome study (Zhang et al., 2019) showed that the genera most frequently found on the leaves of *Erisyphe* were *Pleosporales* and *Alternaria* (>10% of total sequences). Based on spore morphology, we therefore suspected this secondary pathogen to be Alternaria tenuissima (Appendix 1, sheet no. 153), which had already been identified on this host species, although attempts at host reinfection were not successful (Safari Motlagh & Bayegan, 2014). We also found Phaeobotryosphaeria visci on European mistletoe (*Viscum album*) in many locations (Appendix 1, sheet no. 133). This fungal pathogen received very little attention in France although it was reported to be a potential candidate for biological control (Varga et al., 2012), an interesting perspective as mistletoe is abundant in the trees of the French countryside. These six examples show that it is not rare to detect new occurrences of plant pathogens. Species can be identified morphologically with simple material, although more formal investigations would be required for confirmation of the identification. Organisms living in harmony with their host in a particular country may become invasive or pathogenic when introduced into new areas (e.g. Seebens et al., 2018). It is therefore useful to have records of native plant pathogens, to make it possible to trace their origins.

### Diversity of epidemiological stages and the search for alternate hosts: more fun than detective games?

Two of the 97 rust pathogen species we identified were found at all four epidemiological stages (pycnida-aecia-uredinia-telia), on the same host species for *Cumminsiella mirabilissima* (*Mahonia aquifolium syn. Berberis aquifolium*; Appendix 1, sheet no. 5) and on three different hosts for *Puccinia sessilis* (*Arum maculatum, Polygonatum multiflorum* and *Phalaris arundinacea*; Appendix 1, sheet no. 74). Six were suspected to be present at three stages on the same host (*Puccinia lapsanae* on *Lapasana communis*; Appendix 1, sheet no. 60) or on two different hosts (*Gymnosporangium clavariiforme* on *Crataegus monogyna* and *Juniperus communis*, *Puccinia caricina* var. *ribesii-pendula* on *Ribes rubrum* and *Carex pendula*, *Puccinia clematidis-secalis* or *Puccinia agropyri* on *Clematis vitalba* and *Elymus campestris*, *Puccinia urticae-hirtae* on *Urtica dioica* and *Carex hirta*, and *Uromyces dactylidis* on *Ficaria verna* and *Dactylis glomerata*; Appendix 1, sheet no. 6, no. 42, no. 44, no. 80, and no. 87, respectively). Forty were found at two developmental stages (32 uredinia-telia, 6 pycnia-aecia, 2 aecia-uredinia) and 49 at only one stage (30 uredinia, 13 telia, 6 aecia). It was not possible to establish unequivocally that the six aforementioned rust specimens on two different hosts belonged to the same species, but the level of suspicion was high for *Gymnosporangium clavariiforme* (Millau), *Puccinia caricina* var. *ribesii-pendula* (Orgeval), and *Puccinia clematidis-secalis* or *Puccinia agropyri* (Millau), as for Puccinia urticae-acutiformis found at the aecia and telia stages (Orgeval; Appendix 1, sheet no. 79; Appendix 2, conversation P), because we found the two infected hosts on the same day, separated by only a few meters. The identification of the rust found on Carex pendula at Les Clayes-sous-Bois as Puccinia caricina var. ribesii-pendula (Appendix 1, sheet no. 42), was uncertain. Indeed, we found uninfected specimens of the alternate host (Ribes rubrum) of this species in close proximity (<10 m away), and uninfected specimens of the alternate hosts (Tussilago farfara and Urtica dioica) of Puccinia petasites-pendulae and Puccinia urticata, respectively, which can also attack C. pendula (Klenke & Scholler, 2015), a bit further away (50 to 300 m away). For the 27 powdery mildews that we found, the perfect stage (sexual) was observed in 15 cases, rendering species identification much more reliable, based on the morphology of the cleistothecia (size, appendages) and asci (shape and number of ascospores).

### Fungal plant diseases are beautiful

Common pathogenic species with particularly aesthetic forms were observed, including *Rosa* sp. fruits contaminated with *Phragmidium mucronatum* resembling ‘olives stuffed with peppers’ (Appendix 1, sheet no. 26). We also observed a number of other attractive structures for which we posted photographs on Twitter. These structures included columns of *Cronartium flaccidum* teliospores on *Paeonia lactiflora* (Appendix 1, sheet no. 4), tubes containing *Gymnosporangium clavariiforme* aeciospores on leaves and stems of *Crataegus monogyna* (Appendix 1, sheet no. 6) and lantern-shaped growths containing *Gymnosporangium sabinae* aeciospores on *Pyrus communis* (Appendix 1, sheet no. 7). We also found spots on leaves of *Sambucus nigra* caused by *Cercospora depazeoides*, with such a beautiful graphical aspect that we presented this disease as ‘snake skin disease’ (Appendix 1, sheet no. 160).

### Pathogenic fungi are not the only outstanding feature of diseased plants

On some diseased plants, we found small animals associated with the fungal pathogen. For instance, we observed slugs feeding on *Coleosporium tussilaginis* f. sp. *petasitis telia* on *Petasites hybridus* (Appendix 1, sheet no. 2), with orange excrement containing the remains of teliospores. Interestingly, this observations was consistent with previous findings that some mollusc species prefer grazing on plants infected with rust fungi (Ramsell & Paul, 1990; Hajian-Forooshani et al., 2020). On various rust-infected plants, we found larvae of Mycodiplosis sp. feeding on: (i) aecia of Melampsora euphorbiae-dulcis on Euphorbia lathyris (Appendix 1, sheet no. 16); (ii) uredinia of Melampsora euphorbiae on Euphorbia amygdaloides (Appendix 1, sheet no. 15), Melampsora hypericorum on Hypericum perforatum (Appendix 1, sheet no. 17), Melampsora sp. on Salix atrocinerea (Appendix 1, sheet no. 22), Puccinia antirrhini on Antirrhinum majus (Appendix 1, sheet no. 35), Puccinia menthae on Mentha aquatica(Appendix 1, sheet no. 63), and Puccinia sessilis on Polygonatum multiflorum and Arum maculatum with a nymph of Symphypleones sp. (Appendix 1, sheet no. 74). We observed and described an emblematic example of mutualistic symbiosis between a grass (Dactylis glomerata), a fungal endophyte (Epichloë typhina) and an insect (Botanophila sp.; Appendix 1, sheet no. 183). In a dedicated thread on Twitter (see below), we thus presented several photographs of fungal stromata of E. typhina found on the sheath of the flag leaf of D. glomerata in the forest of Rambouillet (Viennot-Bourgin, 1949; Bultman & Leuchtmann, 2008). Most were still immature (white), but a few (turning orange) displayed the initiation of perithecia, the sexual stage of the fungus. We explained that the stromata are usually visited by Botanophila sp. flies, for feeding and egg laying, serving as the vector of spermatia and mediating cross-fertilisation in E. typhina, which is a heterothallic obligate outcrosser. We presented photographs with Botanophila sp. eggs and larval brood chambers, which we found on >85% of the stromata (on average, 4-5 per stroma), consistent with the findings of Górzyńska et al. (2010). A similar, but less common, interaction was found on *Brachypodium pinnatum* subsp. rupestre with Epichloë sylvatica (Appendix 1, sheet no. 184). We also observed aphids, such as *Macrosiphoniella artemisiae* on Artemisia vulgaris, but found no records of interactions established with the pathogen Puccinia artemisiella (Appendix 1, sheet no. 37). Finally, we identified three hyperparasite fungi: *Ampelomyces quisqualis* with *Golovinomyces cichoracearum*, *Tuberculina persicina* with *Puccinia sessilis*, and *Tuberculina sbrozzii* with *Puccinia vincae* (Appendix 1, sheet no. 191-193).

### Communication on Twitter

All 196 pathogens were presented in Twitter messages or threads, with the support of one or several photographic sheets, generally posted within two to three days of observation of the symptoms. Three examples of Twitter conversations from these messages are presented in Figure 6, and a dozen more are shown in Appendix 2. The various messages were intended for the general public, but also for specialists (farmers, scientists, students). We explained some of our practical activities, sometimes involving our children (Figure 3). Several comments from other plant pathology experts helped us to identify the pathogen species when our initial identification was erroneous or uncertain (Appendix 2, conversations G, H and I). Fruitful interactions occurred in the particular context of IYPH2020, promoted by the International Plant Protection Convention (@ippcnews) with the EPPO mascot “Beastie the bug” (@beastiebug), the British Society for Plant Pathology BSPP (@BS_PP) with the #WildPlantDisease and #WildFlowerHour challenge in August 2020 (Appendix 2, conversation B), the French Phytopathological Society SFP (@SFP_France) and the Plant Health and Environment SPE INRAE division (@INRAE_DPT_SPE) (Appendix 2, conversation A). Educational aspects were highlighted in several threads, mostly in English but sometimes in French, during the first lockdown, during which we had to organize home schooling (Appendix 2, conversations C, D, E and F). We adopted an ironic tone on several occasions, considering that visual or situational jokes might make the conversation more attractive (Appendix 2, conversations K, L, M, N, O). From March 17 to June 20, 2021, the messages relating to the 196 plant pathogens (single tweets or threads) on the @wheatpath account received a total of 5953 engagements (on average, 33 likes, retweets, citations, or responses per conversation). We analysed the number of engagements according to disease category (rusts, n = 97; powdery mildews, n = 27; septoria-like diseases, n = 28; other fungal leaf diseases, n = 30; other fungal non-leaf diseases, n = 8; hyperparasite fungi, n = 3; non-fungal diseases, n = 3; Table S1), and the results of this analysis are presented in Figure 7. If we consider the number of engagements to reflect interest in the pathogen presented (although there are, obviously, multiple biases), we can conclude that rusts are the most attractive plant diseases (on average 42 engagements per species), ahead of powdery mildew (28 engagements). Septoria-like diseases and other fungal leaf diseases were much less popular (19 engagements). We can explain these results by the nature of @wheatpath followers, some of whom are rust specialists, and by the attractive profile of rusts: they present considerable diversity, with many species, and a number of beautiful and diverse structures (stages 0 to IV). These characteristics are of a type likely to appeal to naturalists and collectors (see the discussion of Pokémon below).

**Figure 6.**
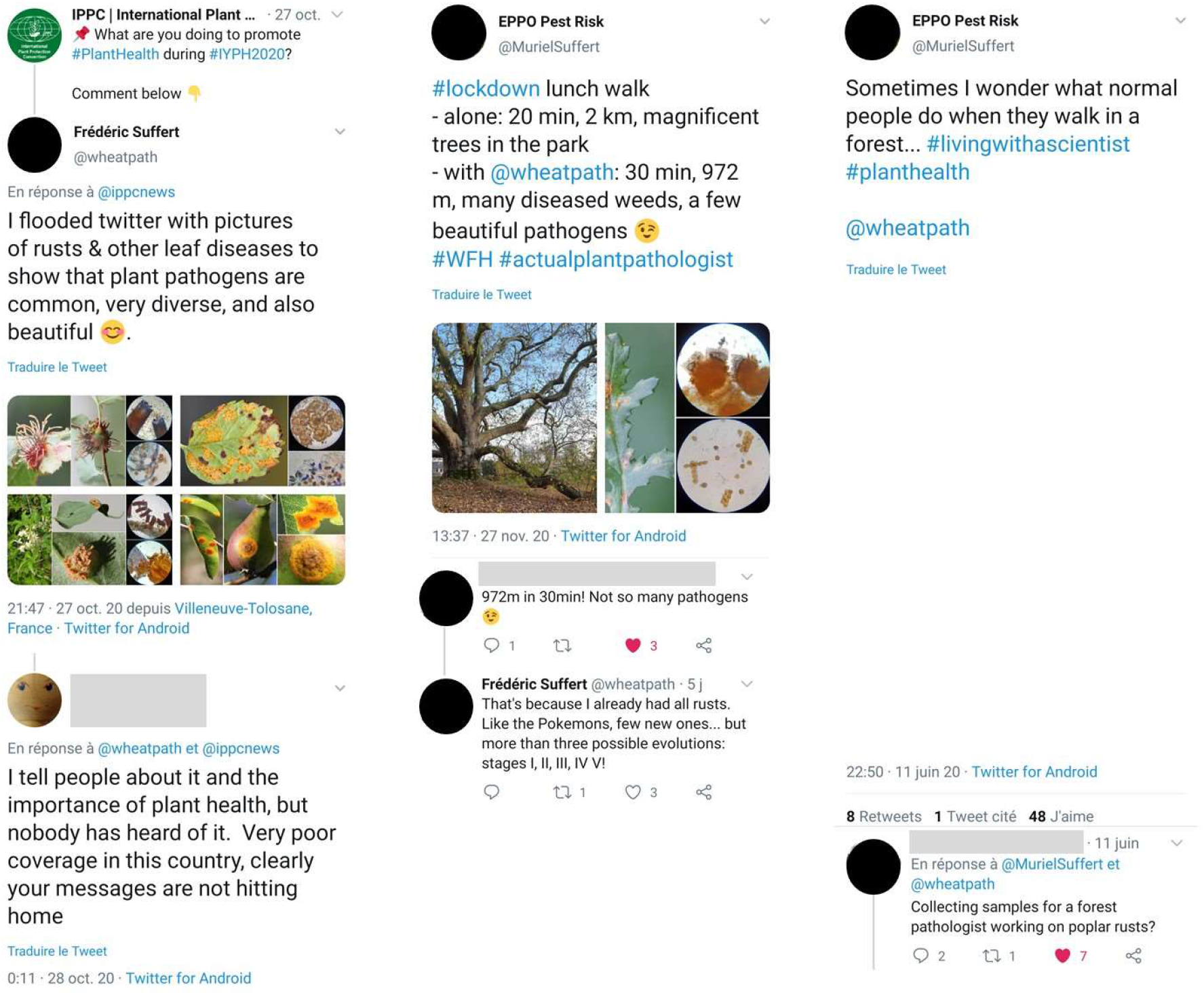
Three examples of conversations on Twitter from a dozen such conversations presented in Appendix 2.

**Figure 7.**
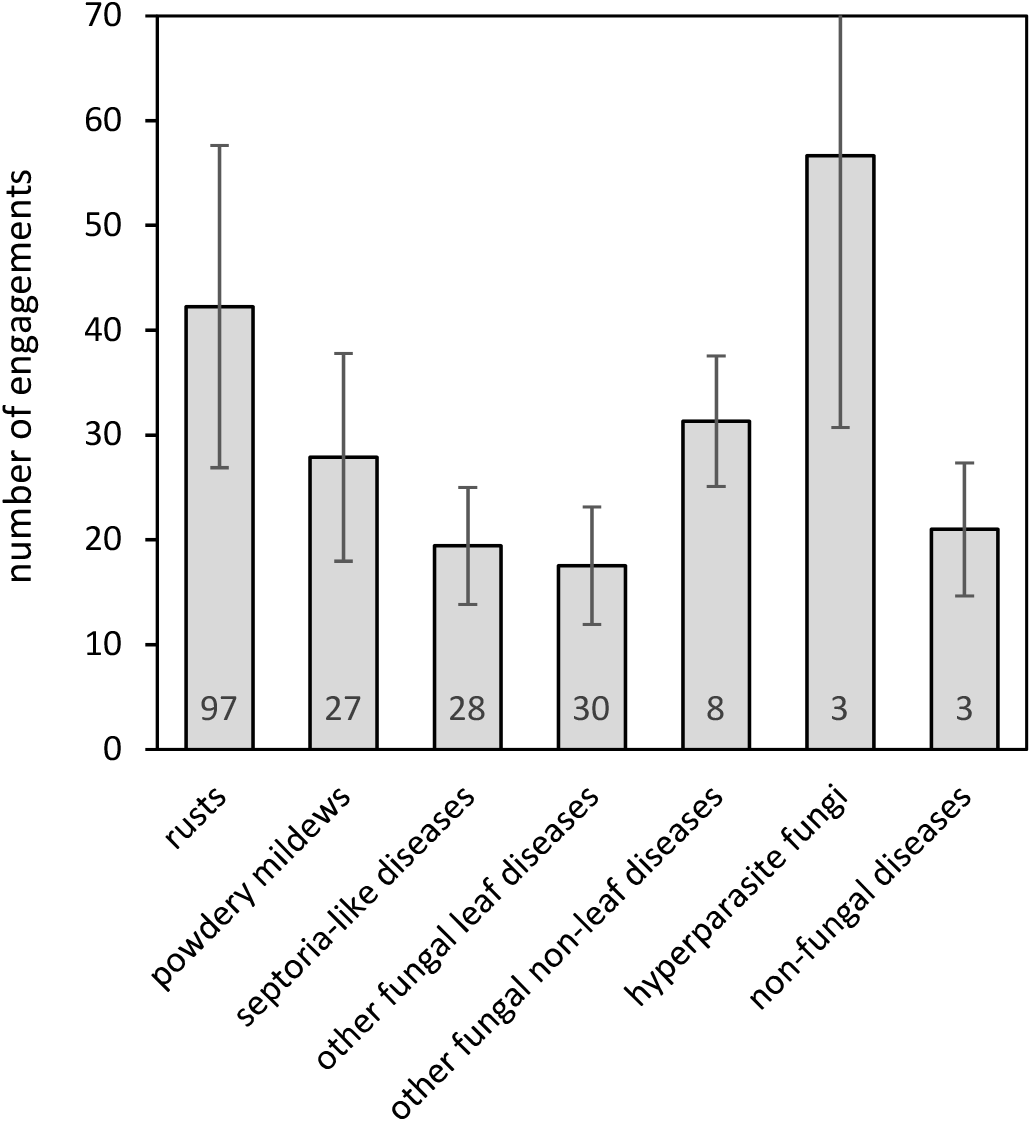
Preference for different categories of disease on Twitter based on the mean number of ‘engagements’ (like, retweet, citation or comment) for each tweet (unique message or thread). For each category, the number of diseases (total = 196) mentioned in at least one tweet is indicated. Bars indicate the standard deviation

Finally, our children were very interested in this experience, both in terms of the search for samples during strolls and the microscopy observations at home. Of course, these strolls also provided an opportunity to observe nature in general and to identify interesting other fungi, insects and plants, but we do not deal with this aspect here.

## Discussion

### Reviewing individual diseased plants: how can we highlight host-pathogen pairs?

The COVID-19 lockdowns have proved a real challenge for us, both professionally, as a scientific researcher focusing on plant health at national institute (INRAE) and a scientific officer in an intergovernmental organization (EPPO), and as parents responsible for educating our children (6, 12 and 13 years old) at home to compensate for school closures. This unusual experience showed us how difficult it can be to run experiments when access to laboratory facilities is limited, to co-ordinate meetings when colleagues cannot travel and to ensure that projects reach their milestones, in a broad sense, while keeping children motivated and curious about life in situation in which working from home has to be combined with home schooling. However, as shown by this article, these conditions also provided us with a tremendous opportunity to find new inspiration in plant pathology, in our surroundings, away from our usual work routines. Over the last eight months or so, we have seen many academic and non-academic colleagues taking the time to observe carefully and document many wild plants or pathogens on Twitter (see, for instance, the hashtags #WildPlantDisease, #WildFlowerHour, #MoreThanWeeds and #PavementPlants promoted by @BS_PP, @BSBIbotany, @wildflower_hour and @morethanweeds, or the @RPilze @SebbyRust and @pascal_frey personal account). Several of the plant pathogens found came from neglected urban areas (vacant green areas, waste ground, unmaintained portions of pavement, for example), highlighting a strong contrast, the naturalist curiosity of attractive and ephemeral biological structures found in ordinary, sometimes ugly, environments.

The COVID-19 lockdowns placed the spotlight on nature within urban areas, encouraging people to take a fresh look at the green areas around them. They have had a marked effect on citizen science projects, in Australia and South Africa for example, increasing the public response to initiatives for obtaining data on birds (Reside et al., 2020; Rose et al., 2020) or the flowering behaviour of grasses (Haeften et al., 2020). Moreover, bird watching was found to bring happiness to the observers in this difficult context (Nairn, 2020). In our opinion, observing plants and looking for their pathogens is as easy as bird-watching, and probably bring similar satisfaction to those practising it. As pointed out by Döring et al. (2012), there is no reason why healthy plants should be preferred over their pathogens, at least those that cause non-lethal infections. We are thus tempted to turn the adage “what doesn’t kill me makes me stronger” into “what doesn’t kill me makes me more attractive”: plant pathogens can be remarkable, in the most literal sense; they deserve to be ‘remarked’, i.e. to be seen, looked at, described, photographed, shared and used for emotional (artistic) or educational purposes. We show here that it is easy to develop such an approach within very limited spatial and logistic constraints, so long as there is a will and desire to do so. Our connections with some of our plant pathologist colleagues on Twitter helped us to get through lockdown by keeping us engaged with science while creating opportunities for engagement, whether with our children educated at home, or through communication with a broader public. Like Yannelli & Saul (2020), we found that despite (maybe even thanks to) the limitations and difficulties associated with lockdown, everybody can explore their immediate surroundings and discover the diversity and beauty of plant pathogens. We found this experience very encouraging, particularly given the likelihood of similar lockdown conditions being imposed again.

### Promoting the conservation of plant pathogens

Botanic gardens generally focus on plant conservation, but the conservation of native plant pathogens, as a component of local biodiversity, should also be considered. This point has already been made by Ingram, (1999, 2002), and came up again in the context of the IYPH2020 (Ingram, 2020), through three initiatives of the BSPP: (i) the #WildPlantDisease Twitter challenge, in which we participated in late August, by posting some of the pathogen photographs presented in Appendix 1; (ii) the Welsh Rust Group, producing red data lists and census catalogues for native plant pathogens in Wales; (iii) the Wyre Forest Study Group, a citizen’s science research group studying the natural history of the Wyre Forest in the West Midlands, which has recently decided to survey native plant pathogens. The conservation of plant pathogens essentially occurs ex situ, in microbiological collections, the activities of which are co-ordinated by the World Federation for Culture Collections (Hawksworth, 1997). Strict biotrophic pathogens, such as rust and powdery mildews, which account for more than half the pathogens described in this article (Appendix 1, Table S1), are obviously more problematic than fungi that can be cultured in axenic conditions. Consequently, the alternative principle of plant pathogen conservation in situ seems to be essential, even though it goes against the intuition of most plant health scientists, whose careers have been dedicated to preventing or eradicating plant diseases, and who have long adopted a normative view of plant health (Döring et al., 2012). This purely normative view no longer seems to be widely supported, at least in Europe. New conceptual frameworks for disease management, such as agroecology, are paving the way for the management of certain diseases by the stimulation of ecological regulation mechanisms, rather than aiming to achieve pathogen ‘eradication’ sensu lato, through the use of fungicides (field scale), through the implementation of phytosanitary measures (at the scale of national territories). The conservation of plant pathogens can benefit from the naturalist dimension, and can be appreciated for its educational and artistic value. In a context in which considerable means are legitimately devoted to limiting the emergence of certain pathogens, or eradicating them from vast territories (e.g. Puccinia graminis on its alternate host; Zhao et al., 2016), Ingram’s proposal (1998, 2002) merits further development: the in situ conservation of pathogens of the wild relatives of crop plants is desirable, not only in the centres of origin and diversity for these crops, to facilitate their management in the long term, but also in the gardens of amateurs, for purposes at the interface between science, art and education. For instance, it might be possible to imagine a ‘rust garden’, consisting of endemic host plants (main and alternate hosts to promote the completion of biological cycles), in which one waits each year for the plants to become covered with orange pustules, just as one might wait for a rose to bloom in a conventional garden. Such an idea was first proposed by Browning (1974) and Dinoor & Eshed (1990), who referred to “living gene parks” and “genetic reserves”, respectively. Of course, before such areas could be established, it would be necessary to characterise the risk for adjacent agroecosystems, applying the strict rule that the plant pathogens conserved should be restricted to those already present in an area. Academics and professionals, through national and international plant pathology societies (e.g. BSPP in UK and SFP in France) for example, could play a more significant role in coordinating and funding the efforts of amateur microbial systematists. The IYPH2020 acted as a catalyst for fostering collaboration, but this would probably not be sufficient. Innovative strategies should be developed, combining substantive issues placed firmly on the agenda of (supra)national institutions and individual initiatives based on agile but mature communication platforms. It should also be noted that certain specific national contexts, such as the ban on the use of synthetic phytosanitary products in French public urban areas since 2017, provide good opportunities for rediscovering urban plants and the pathogens that infect them, in addition to those present in parks and private gardens.

### Adapting the communication of descriptions of plant pathogens in tune with the times

During the first half of the 20th century, outputs equivalent to our ‘phytopathological strolls’ (in French, “excursions phytopathologiques”) were published in specialized journals (e.g. Massenot, 1953). They were generally quite arduous to read, as they targeted specialists, generally lacked illustrations and did not always include microscopic observations. In the second half of the 20th century, such publications were gradually replaced by systematic studies focusing on more or less restricted types of pathogen (e.g. Guyot, 1938, 1951, 1957; Gäumann, 1959), or geographical areas, such as regions (Kuhnholtz-Lordat & Blanchet, 1948) or countries ((Viennot-Bourgin, 1949; Savulescu, 1953), but these studies were still driven by a naturalist descriptive purpose. Such phytopathological descriptions have not completely disappeared (Scheuer, 2012), but they are now much less frequent and are more difficult to publish, as they do not correspond to modern scientific objectives and epistemological standards. Since the beginning of the 21st century, the few reference works that continue to be published (e.g. Klenke & Scholler, 2015) have been supplemented by citizen science, through individual websites (e.g. Ellis, 2020; Kruse, 2020; Lechat, 2020; Sothmann, 2020) or collective initiatives (nature platforms for the sharing of observational data about global biodiversity; e.g. Waarneming.nl, 2021). These initiatives make their content accessible to a broad public and, above all, involve that public as an active player in content development. National and international societies for plant pathology, such as BSPP and SFP, would benefit from linking academic and citizen science.

### Can we draw lessons from the ‘Pokémon paradox’ to develop an education strategy for plant pathology?

Generally speaking, the scientific curiosity of the public, and that of young children in particular, in a naturalist context, can and should be stimulated. The COVID-19 lockdowns, as shown by our experience described here, highlighted that simple actions promoting knowledge about plant pathogens can be attractive. Twitter has become an arena in which expert plant pathologists, professionals and the general public can exchange. More surprisingly, drawing a parallel with the social phenomenon ‘Pokémon mania’, can also provide inspiration for attracting an even wider audience, beyond the experts and amateurs already interested in plant pathogens. Pokémon is a turn-based role-playing game in which the user captures various species and uses them to fight other characters. This game was originally created in 1996 by Satoshi Tajiri, an amateur entomologist, and was partly inspired by his love of nature and his enjoyment of collecting insects. Several years later, zoologists established that British children were better at recognising imaginary creatures from Pokémon than they were at recognising local wild animals, and concluded, somewhat provocatively, that “conservationists are doing less well than the creators of Pokémon at inspiring interest in their subject” (Balmford et al., 2002). Acorn (2009) even referred to the ‘Pokémon paradox’, stating that this game is “both the antithesis of natural history education, and an inspiration to budding naturalists”. This paradox could be relevant for other organisms, such as plant pathogens, even though these organisms are much less popular and attractive to the general public than insects. One might think that this paradox increased in 2016 when ‘Pokémon GO’, the augmented reality mobile game developed by the company Niantic, brought the initial game into the urban world of video games, but this is would be without taking into account the fact that this game required walking outside, usually in urban areas, to capture the virtual creatures. Finally, the players of this game have many points in common with those who, like us, looked for plant pathogens in 2020 under lockdown constraints: more time on screens and social media (Qin et al., 2020), walks in spatially limited areas, including green open spaces and forests. Some organisations, such as the Royal Horticultural Society in the UK, have organised Pokémon-type tours, to help the public to discover gardens (Kondonis, 2016). Going beyond its frivolous nature, Pokémon-style approaches provide a good opportunity for science educators, and not just entomologists, to increase the interest of children in their natural environment (Dorward et al., 2017; Schmidt-Jeffris & Nelson, 2018). The creatures they find can “metamorphose” like insects, but fungal pathogens can do this too, with their anamorph and teleomorph stages, corresponding to changes in morphology in response to environmental stresses (e.g. pycnidiospores, blastospores and chlamydospores in Zymoseptoria tritici; Francisco et al., 2019), and they can even go so far as to have five stages, 0 (pycnial), I (aecial), II (uredinial), III (telial) and IV (basidial) — more than any Pokémon species — in the case of heteroecious macrocyclic rusts completing their life cycle on different host plants (Zhao et al., 2016). We are convinced that both biological bizarreness and biological complexity are attractive. The search for the teleomorph phase (sometimes unknown) or for the alternate host has obsessed several generations of plant pathologists and has occasionally led to the publication of emblematic treatises, book reviews, or Trojan horse methodological works (e.g. Savulescu, 1953; Anikster, 1986). Some alternate hosts took a very long time to be discovered (e.g. the “century-old mystery of Puccinia striiformis” elucidated in 2011 by Jin et al.) and the consensus decision to use only one name (“one fungus, one name”) was agree only a decade ago (Wingfield et al., 2012).

## Conclusion

It is crucial that professional plant pathologists, individually and collectively, play an active part in promoting a favourable perception of plant pathology among the public and in ensuring that relevant issues are taught in exciting and innovative ways in colleges and universities. The experience related in this article paves the way for potentially popular additional actions for communicating about plant diseases and making plant pathology more attractive to the general public, through more direct interactions. The goal is definitely not to focus on the acquisition of specialised knowledge or cutting-edge discoveries, but to share what already exists and to prove that anybody can discover these things in their immediate environment. We highlight the possibility of improving communication about plant pathogens by considering the identification of plant pathogens as a detective game, by adding jokes to descriptions (e.g. comparing aecidial cups with “churros” or with the head of Bart and Lisa Simpson; Appendix 2, conversation D), and finally by activating the addictiveness of a collection approach resembling that used in Pokémon Go. The role of Twitter as a tool for communicating and teaching is well established (Lopez-Franco & Hennen, 1990), but plant pathologists could make better use of this tool. For instance, opportunistic biodiversity observations published through Twitter represent a promising and, until now, unexplored example of data mining (Daume & Galaz, 2016). Furthermore, in 2020, Twitter played a fundamental role in enabling real-time global communication between scientists during the COVID-19 pandemic, at an unprecedented scale (e.g. Pollett & Rivers, 2020). It also made it possible to exchange knowledge and to disseminate scientific education in completely different fields, such as horticulture (Stafne, 2020) and sustainable soil management (Mills et al., 2019). There’s no reason why plant pathology could not play a part in this new trend.

## Appendix 1

Overview of the 196 (plant pathogens ordered by disease categories (rusts, *n* = 97; powdery mildews, *n* = 27; septoria-like diseases, *n* = 28; other fungal leaf diseases, *n* = 30; other fungal non-leaf diseases, *n* = 8; hyperparasite fungi, *n* = 3; non-fungal diseases, *n* = 3) and then by alphabetical order. The date of collection is indicated with the format YMD (year-month-day). Uncertainty in the species identification is indicated (‘?’ for a moderate uncertainty and ‘??’ for a high uncertainty). Unless otherwise stated, the determination was considered certain on the basis of Klenke & Scholler (2015) and Viennot-Bourgin (1949), confirmed by information found on different websites or blogs of plant pathologists (e.g. Ellis, 2020; Kruse, 2020; Lechat, 2020; Sothmann, 2020), or comments received in conversations on Twitter. Additional references were added for species which required further information for their identification.

**1.**
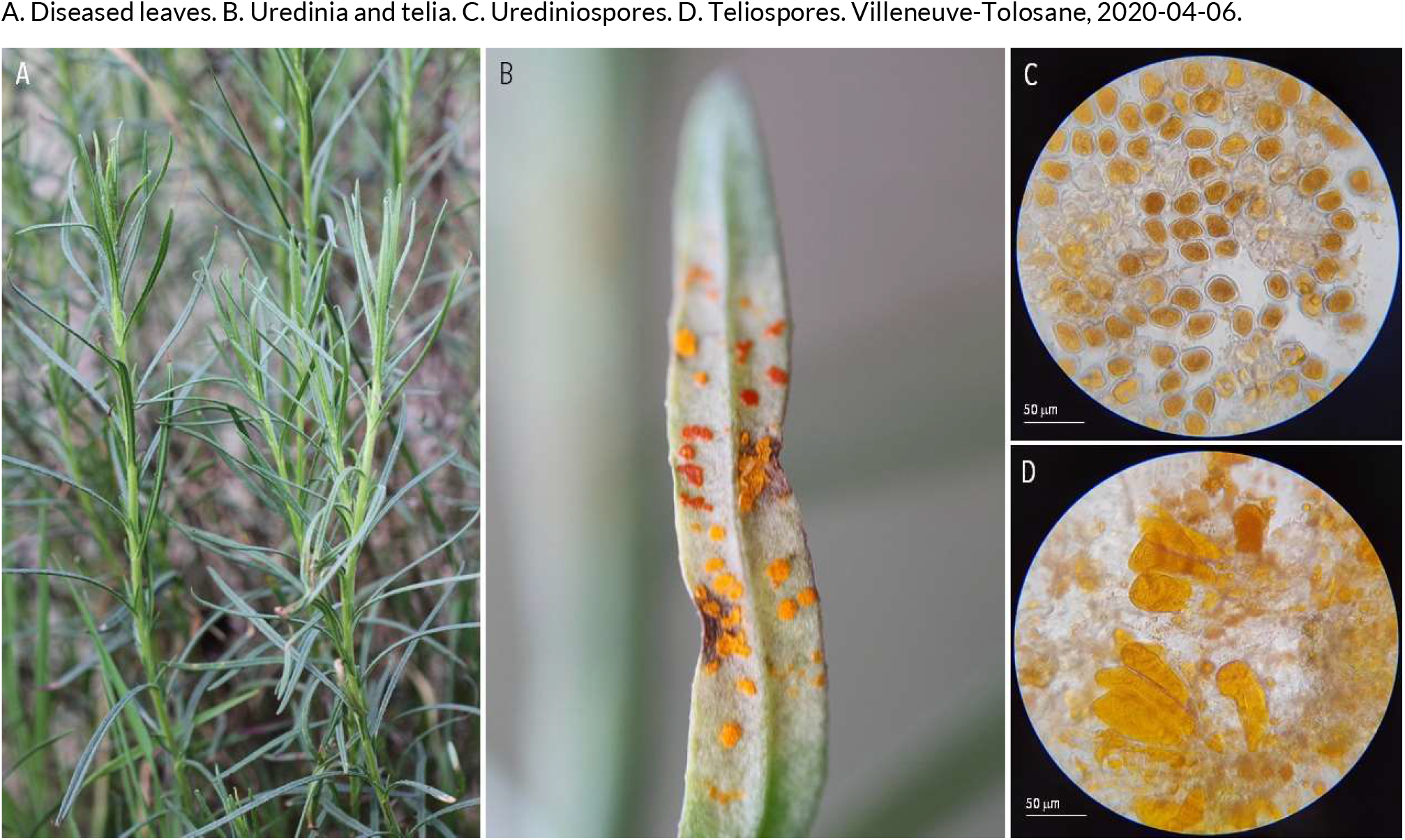
RUSTS | *Coleosporium senecionis* ex *Senecio inaequidens* (narrow-leaved ragwort)

**2.**
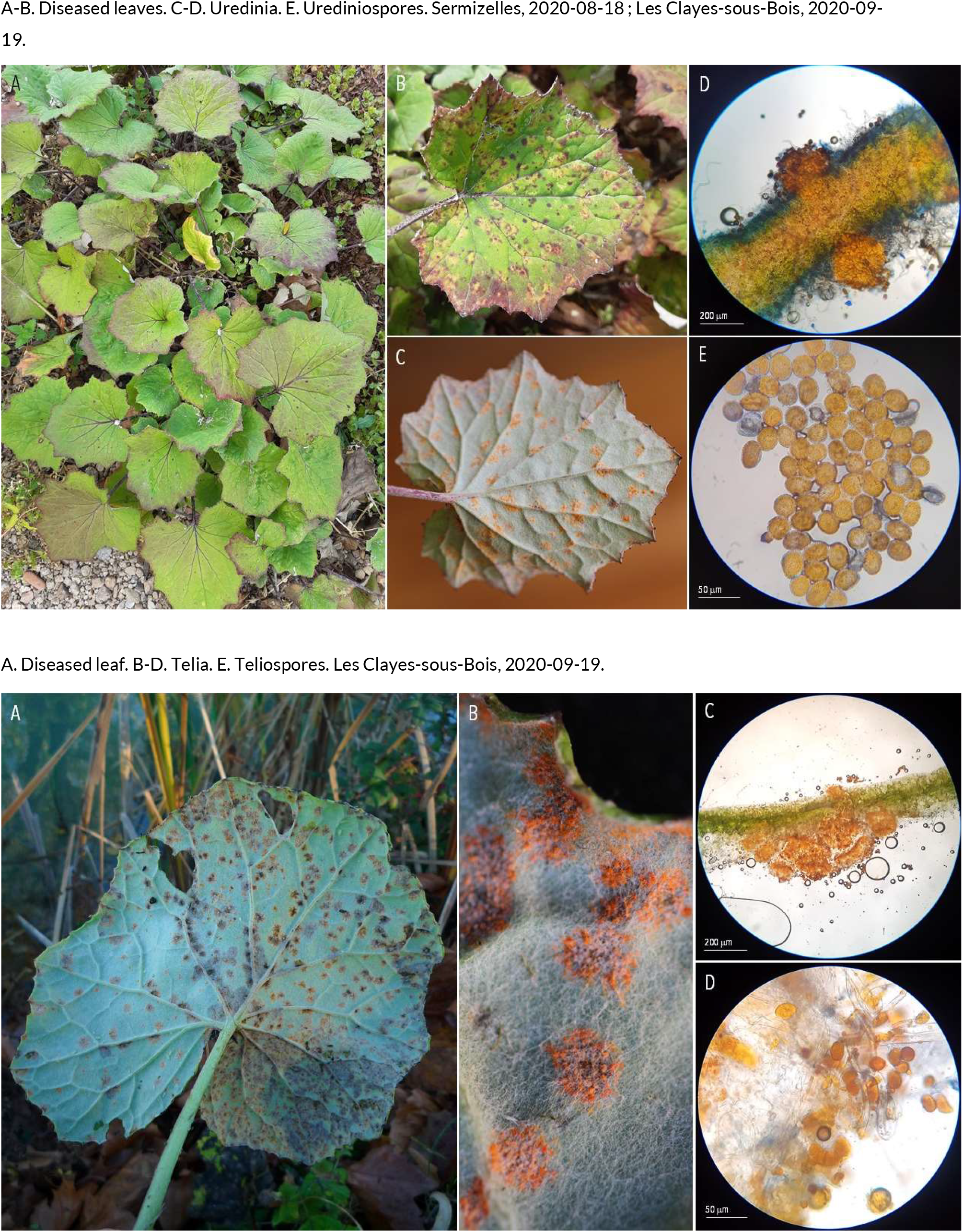
RUSTS | *Coleosporium tussilaginis* (?) ex *Tussilago farfara* (coltsfoot)

**3.**
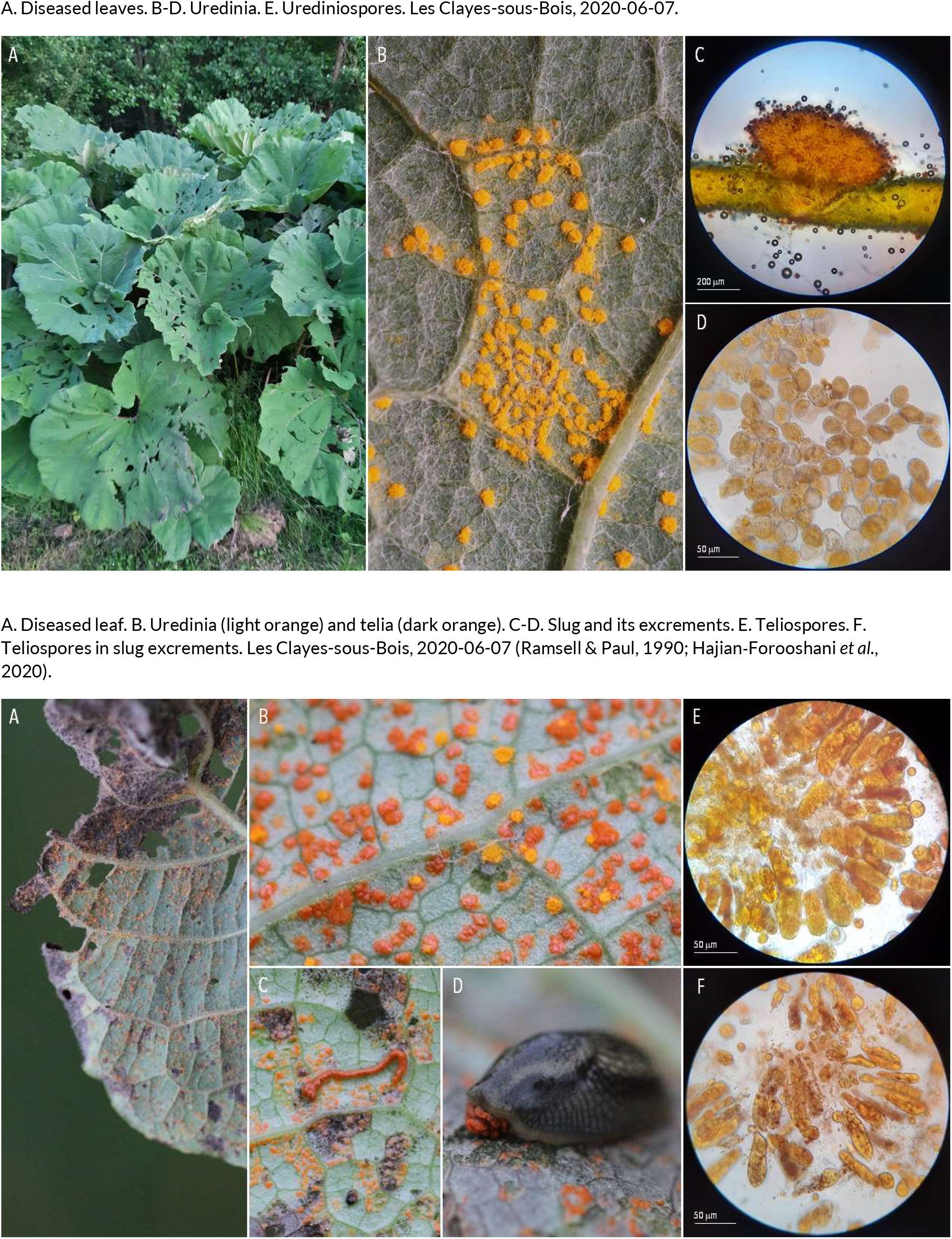
RUSTS | *Coleosporium tussilaginis* f. sp*. petasitis* (?) ex *Petasites hybridus* (butterbur)

**4.**
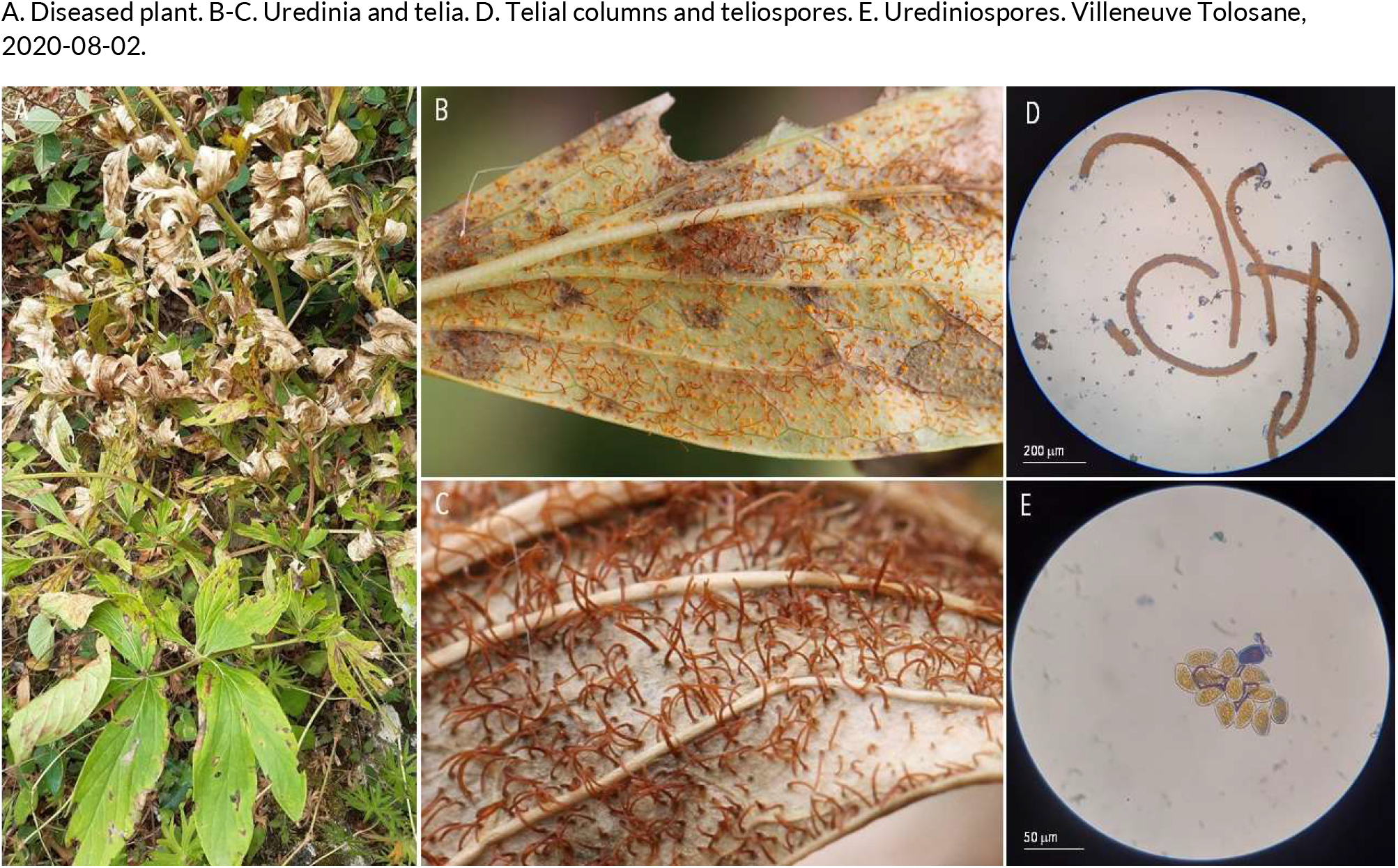
RUSTS | *Cronartium flaccidum ex Paeonia lactiflora* (common garden peony)

**5.**
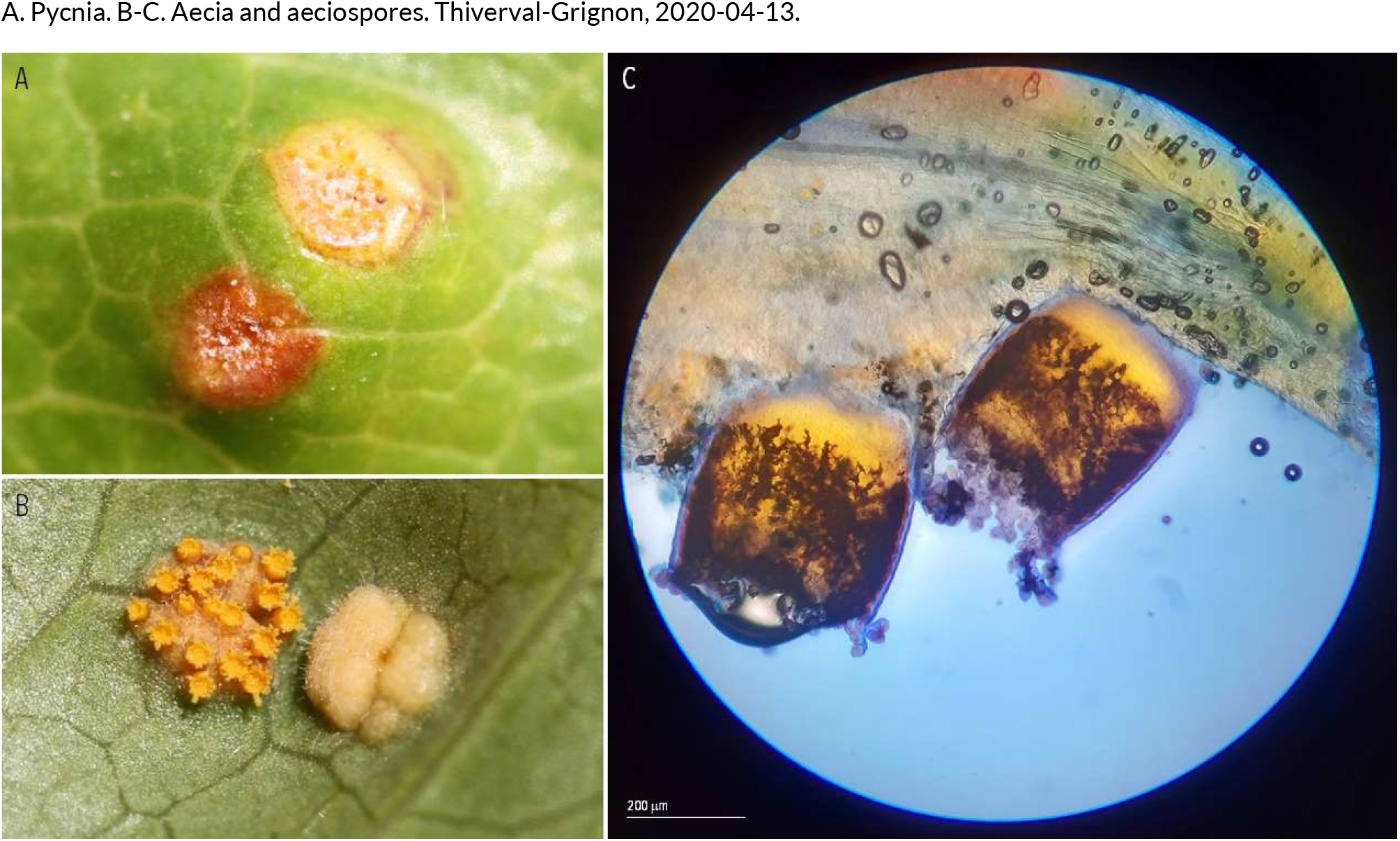
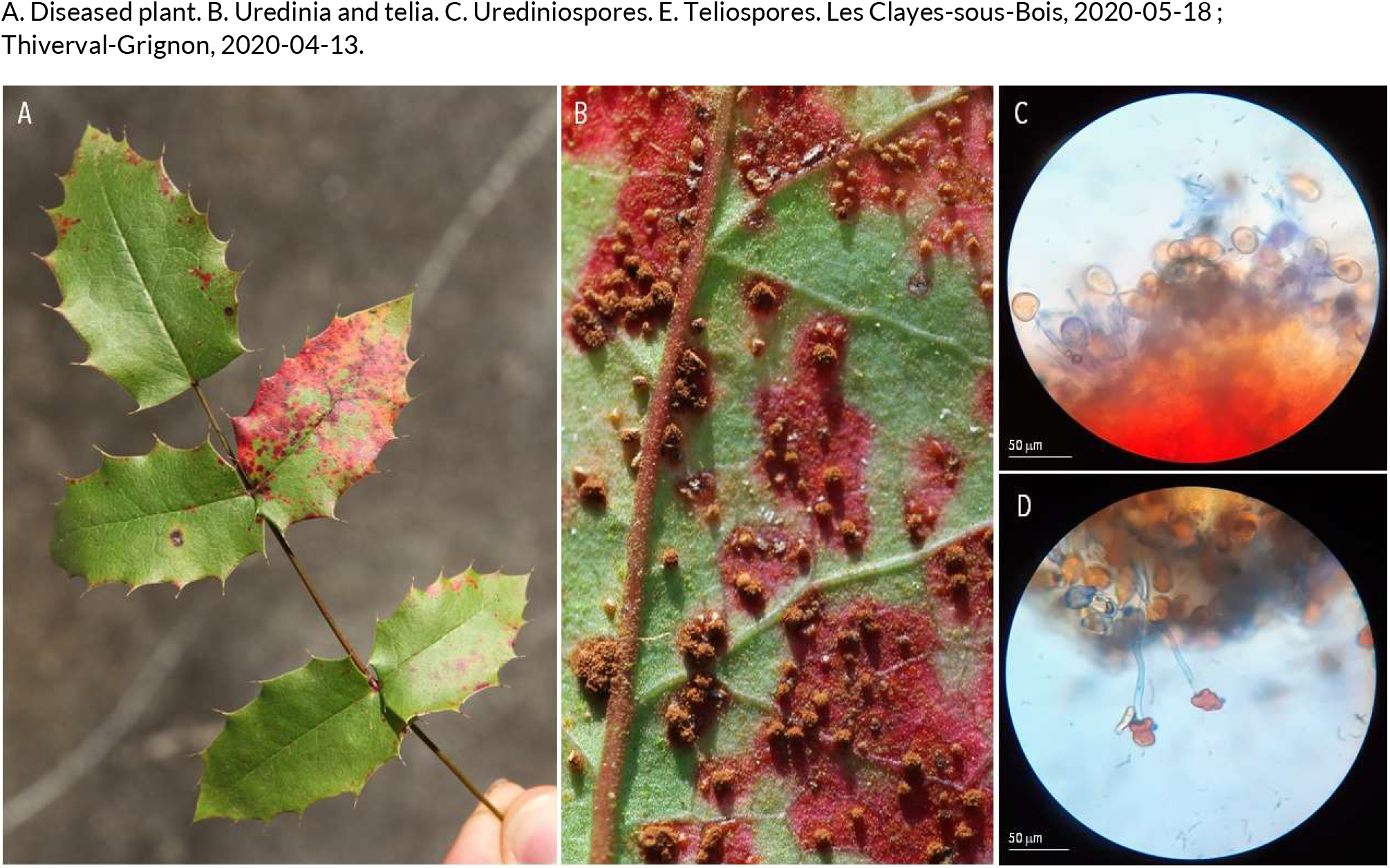
RUSTS | *Cumminsiella mirabilissima* ex Mahonia aquifolium syn. *Berberis aquifolium* (oregon grape)

**6.**
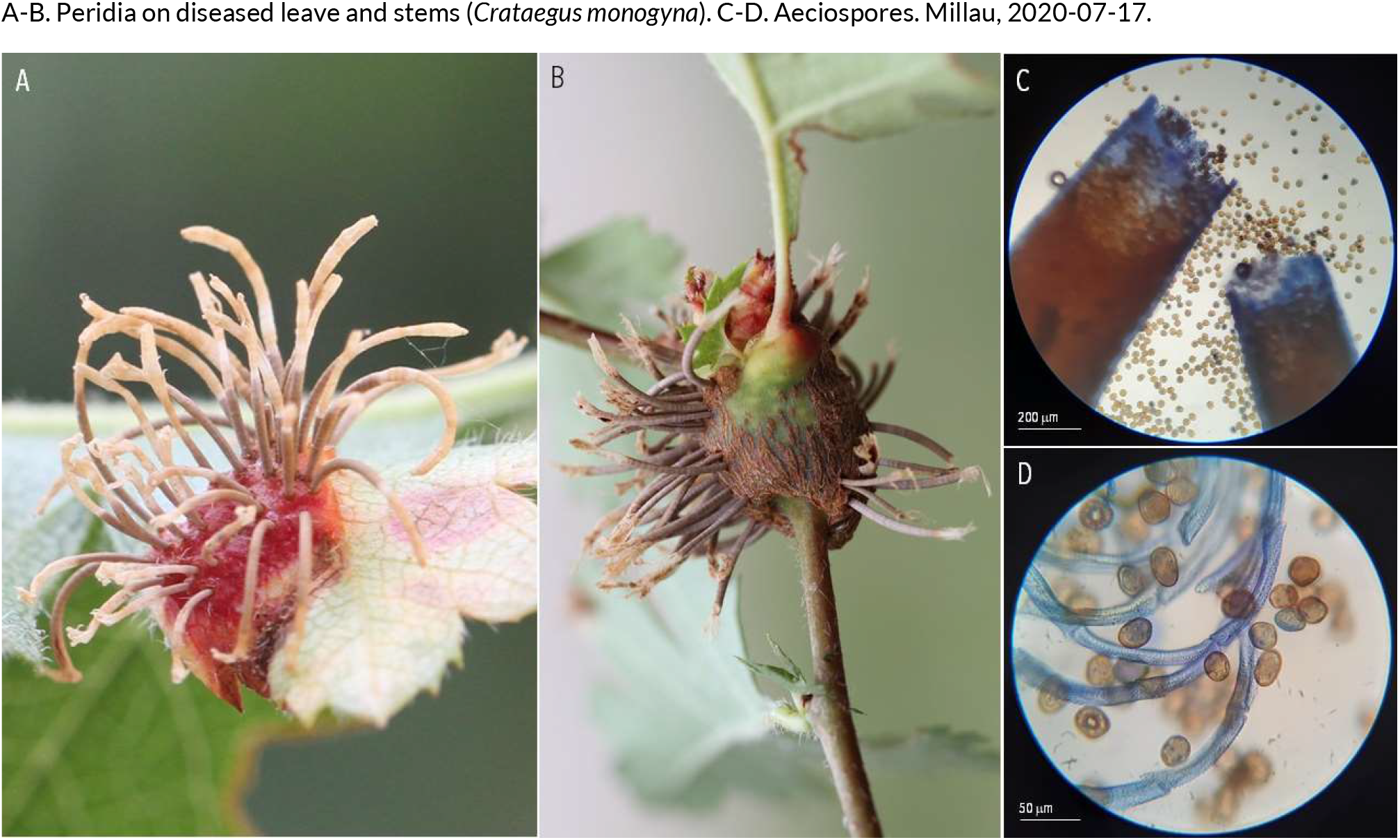
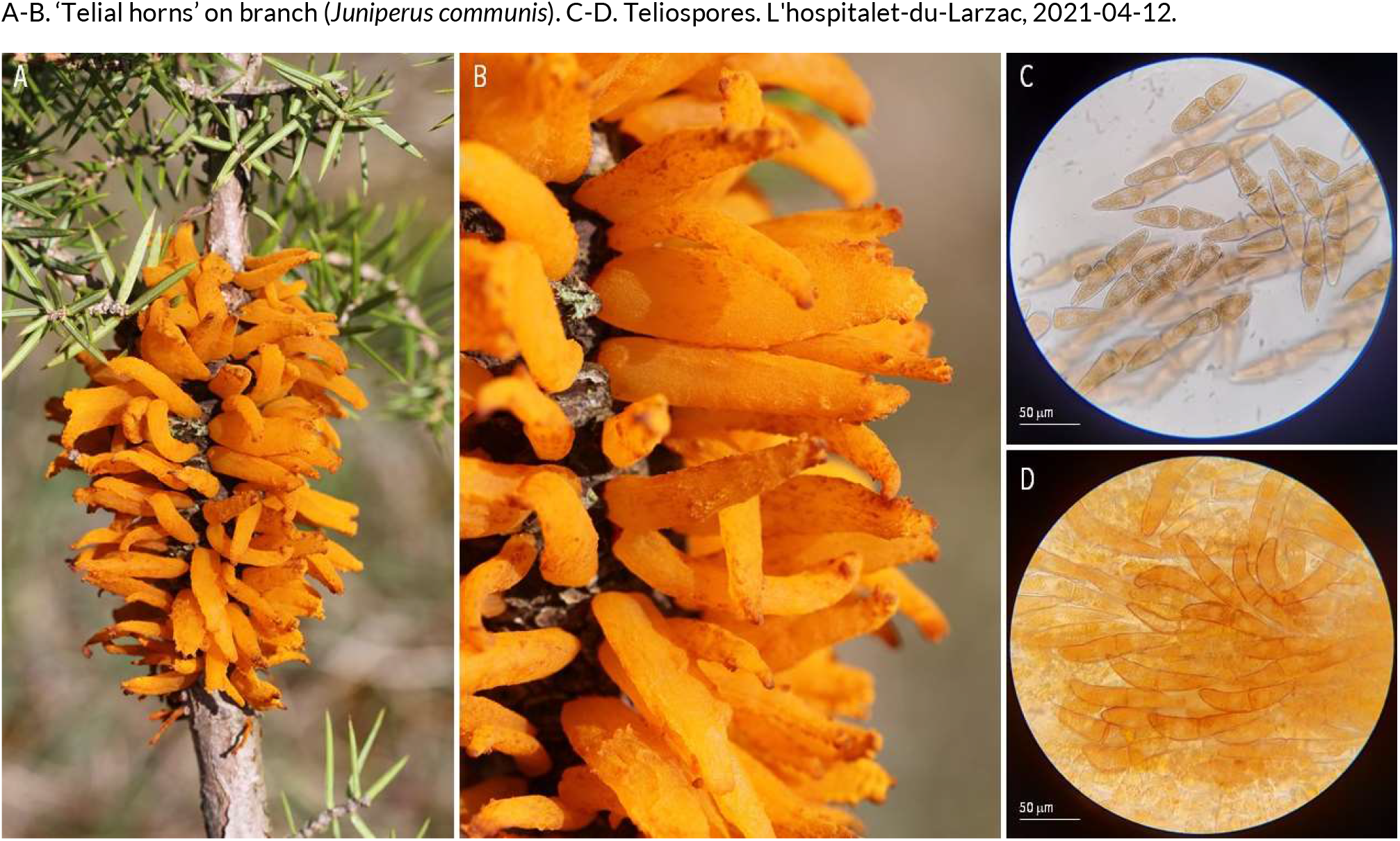
RUSTS | *Gymnosporangium clavariiforme* ex *Crataegus monogyna* (common hawthorn) and *Juniperus communis* (common juniper)

**7.**
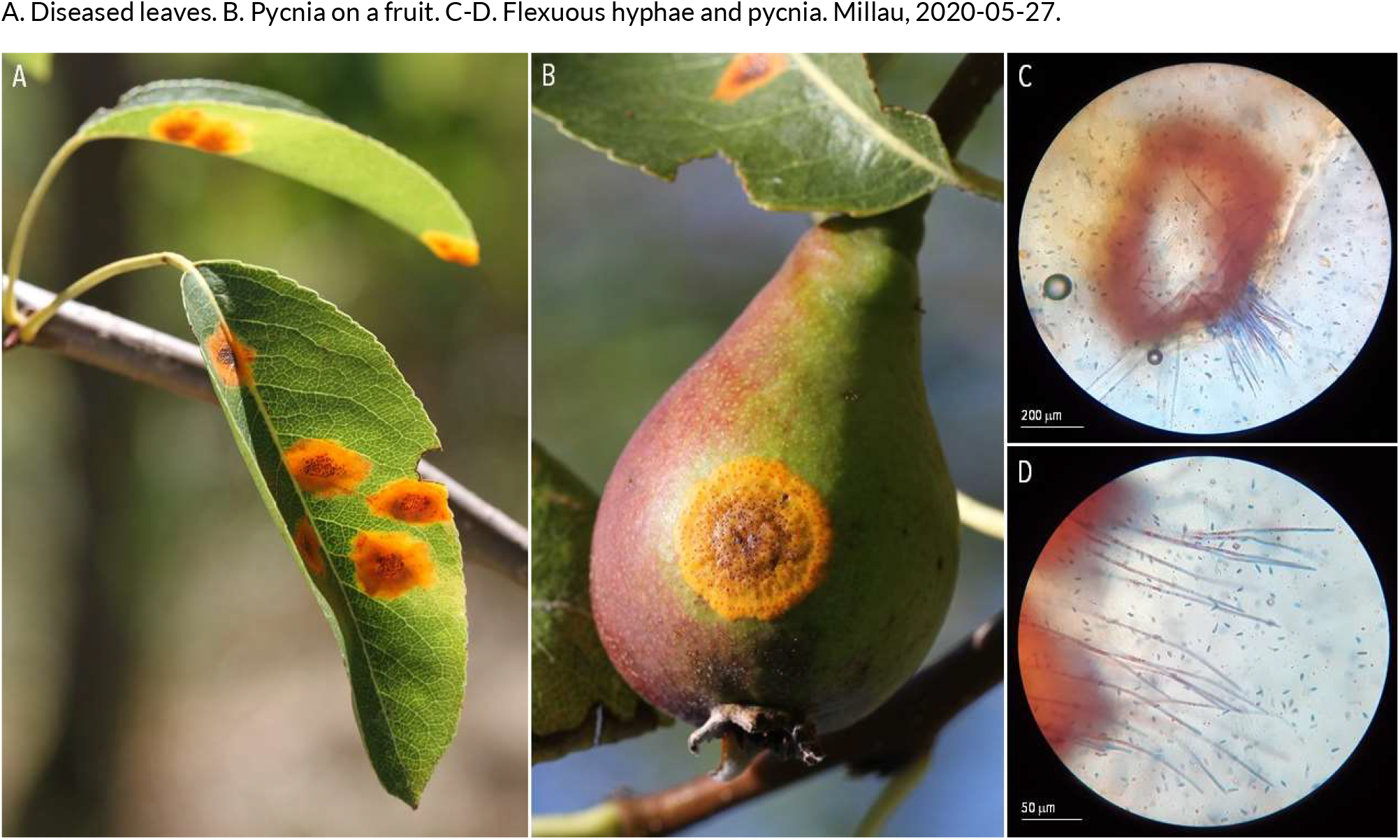
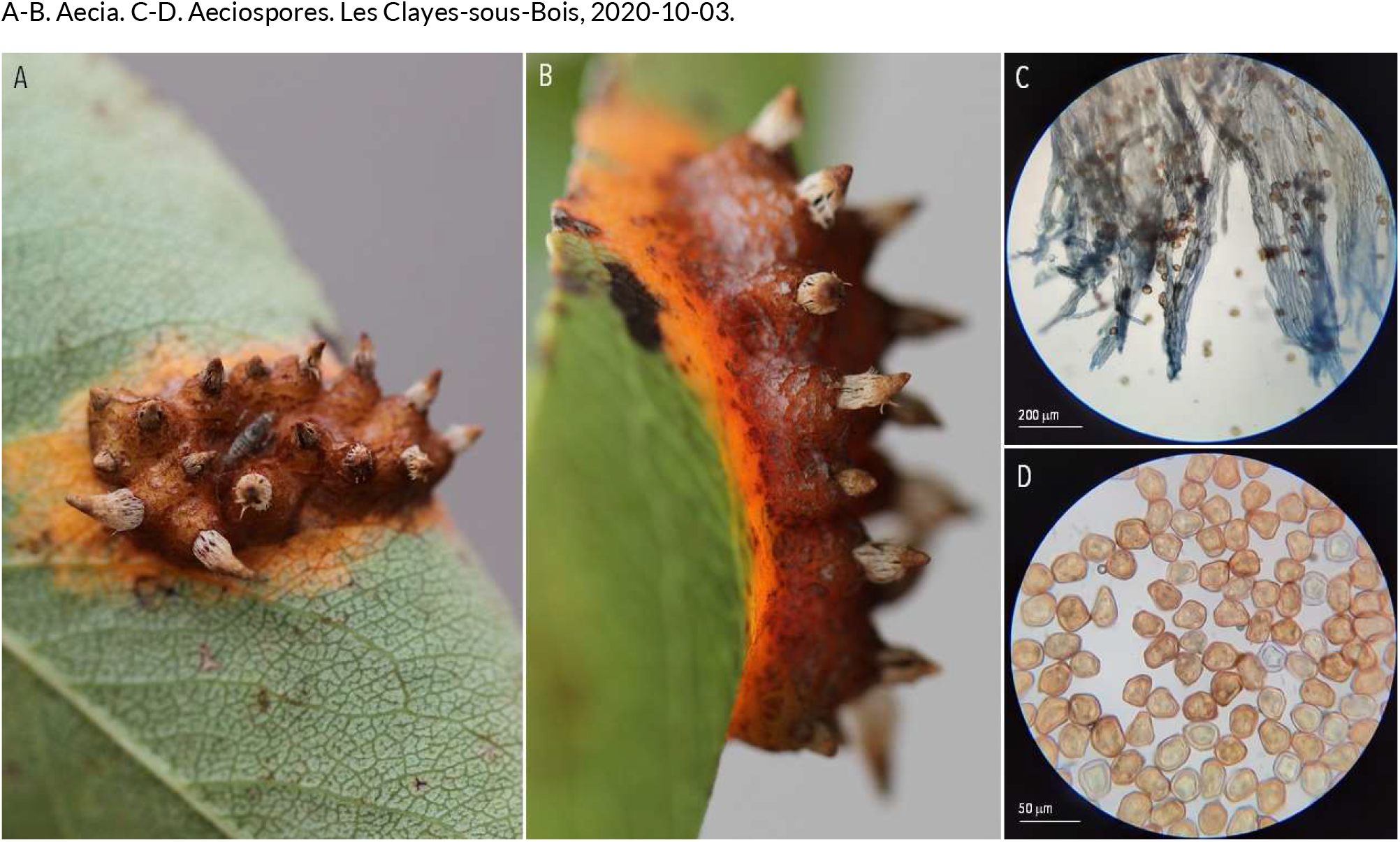
RUSTS | *Gymnosporangium sabinae* ex *Pyrus communis* (European pear)

**8.**
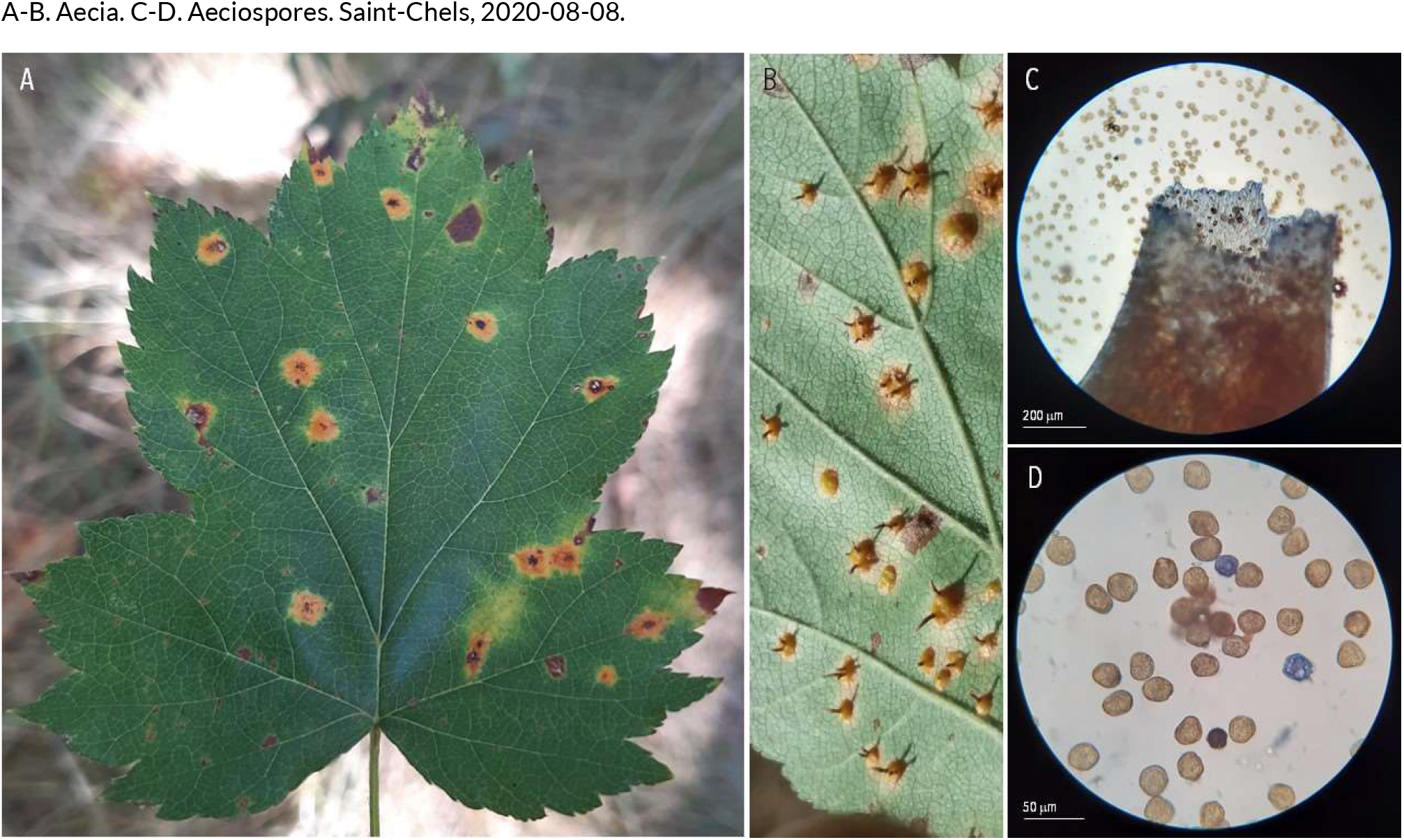
RUSTS | *Gymnosporangium torminali-juniperum* ex *Sorbus torminalis* (wild service tree. Saint-Chels, 2020-08-08.

**9.**
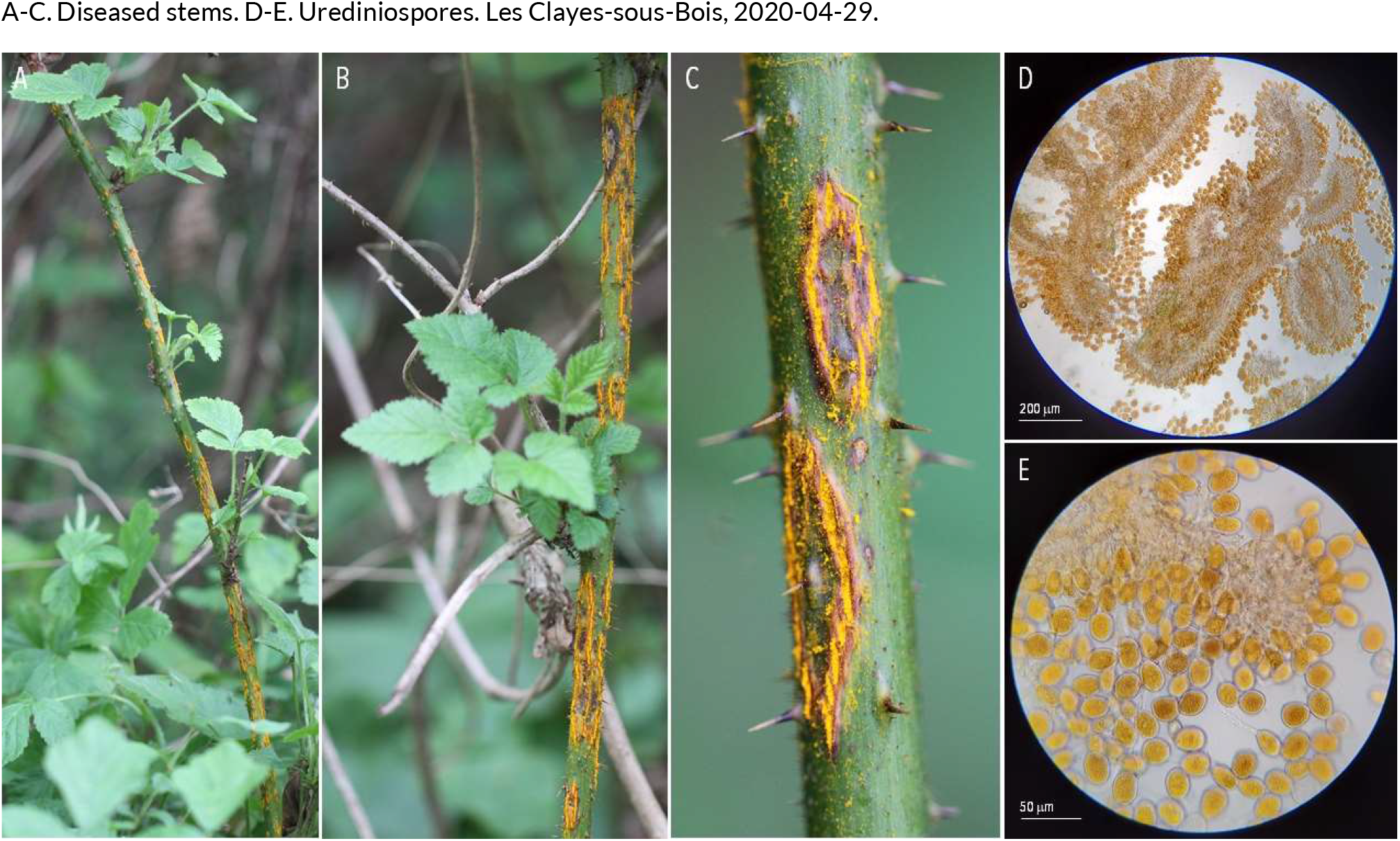
RUSTS | *Kuehneola uredinis* ex *Rubus fruticosus* (blackberry.

**10.**
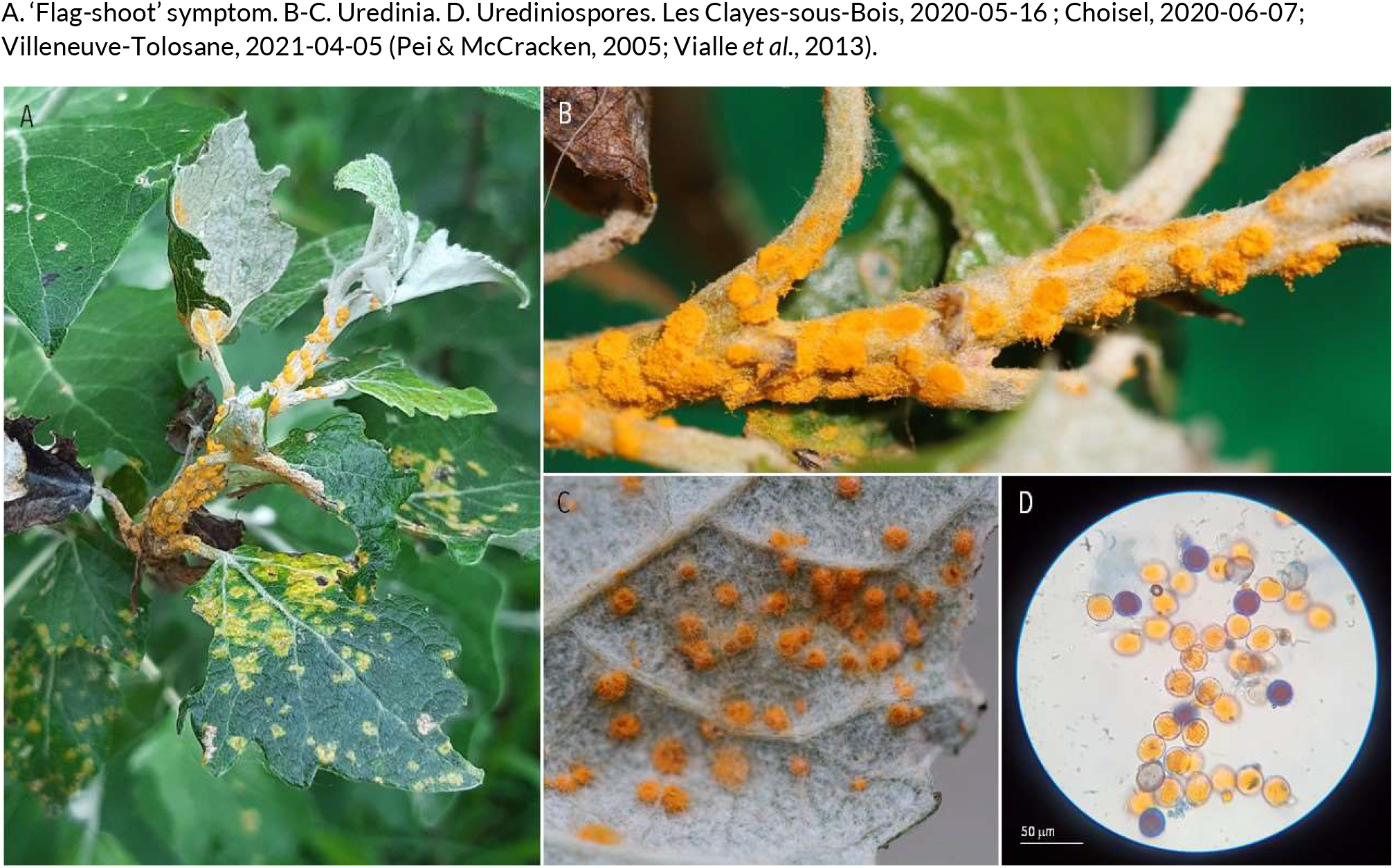
RUSTS | *Melampsora aecidioides* ex *Populus alba* (white poplar)

**11.**
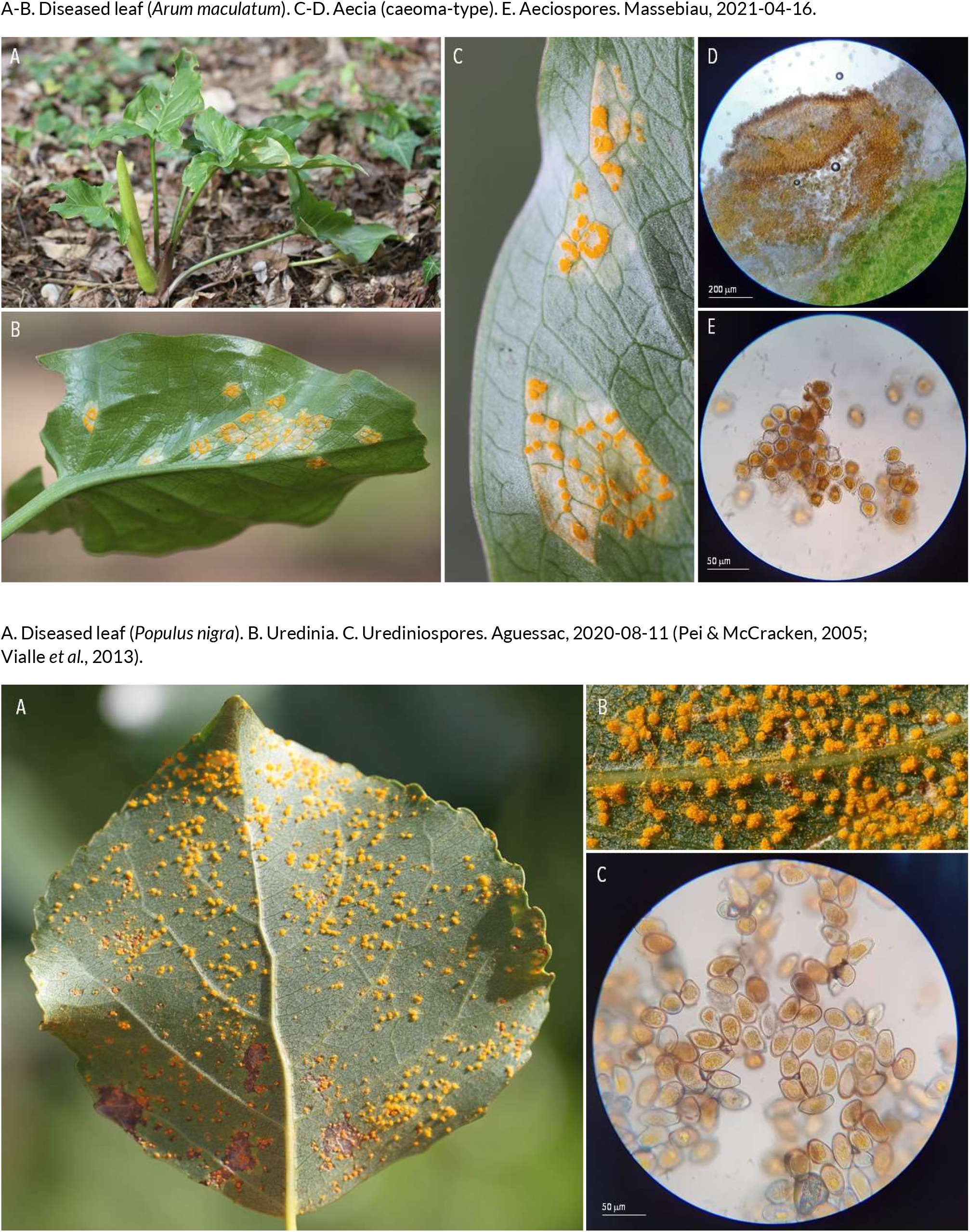
RUSTS | *Melampsora allii-populina* ex *Arum maculatum* (cuckoo pint) and *Populus nigra* (black poplar.

**12.**
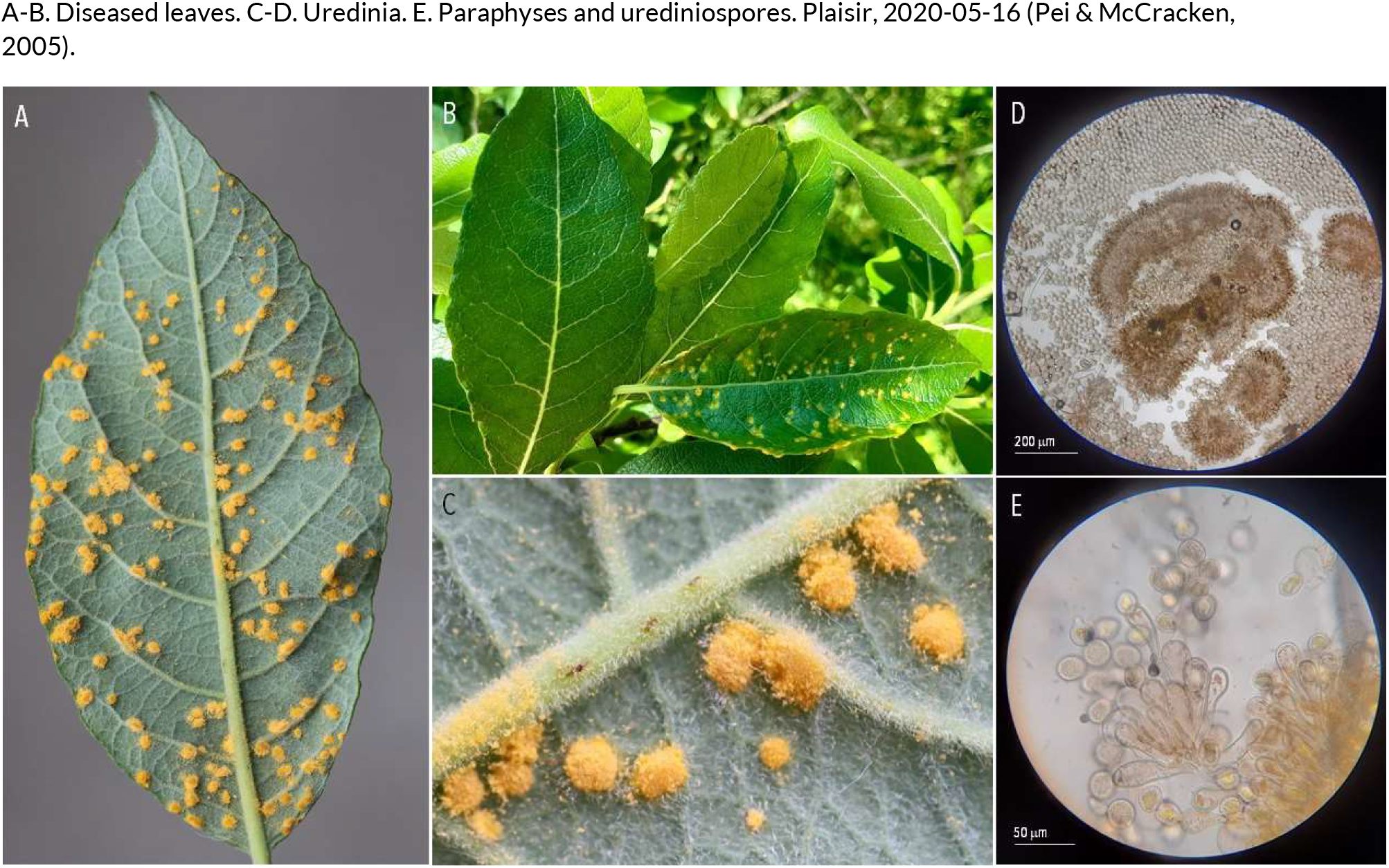
RUSTS | *Melampsora caprearum* (?) ex *Salix caprea* (goat willow)

**13.**
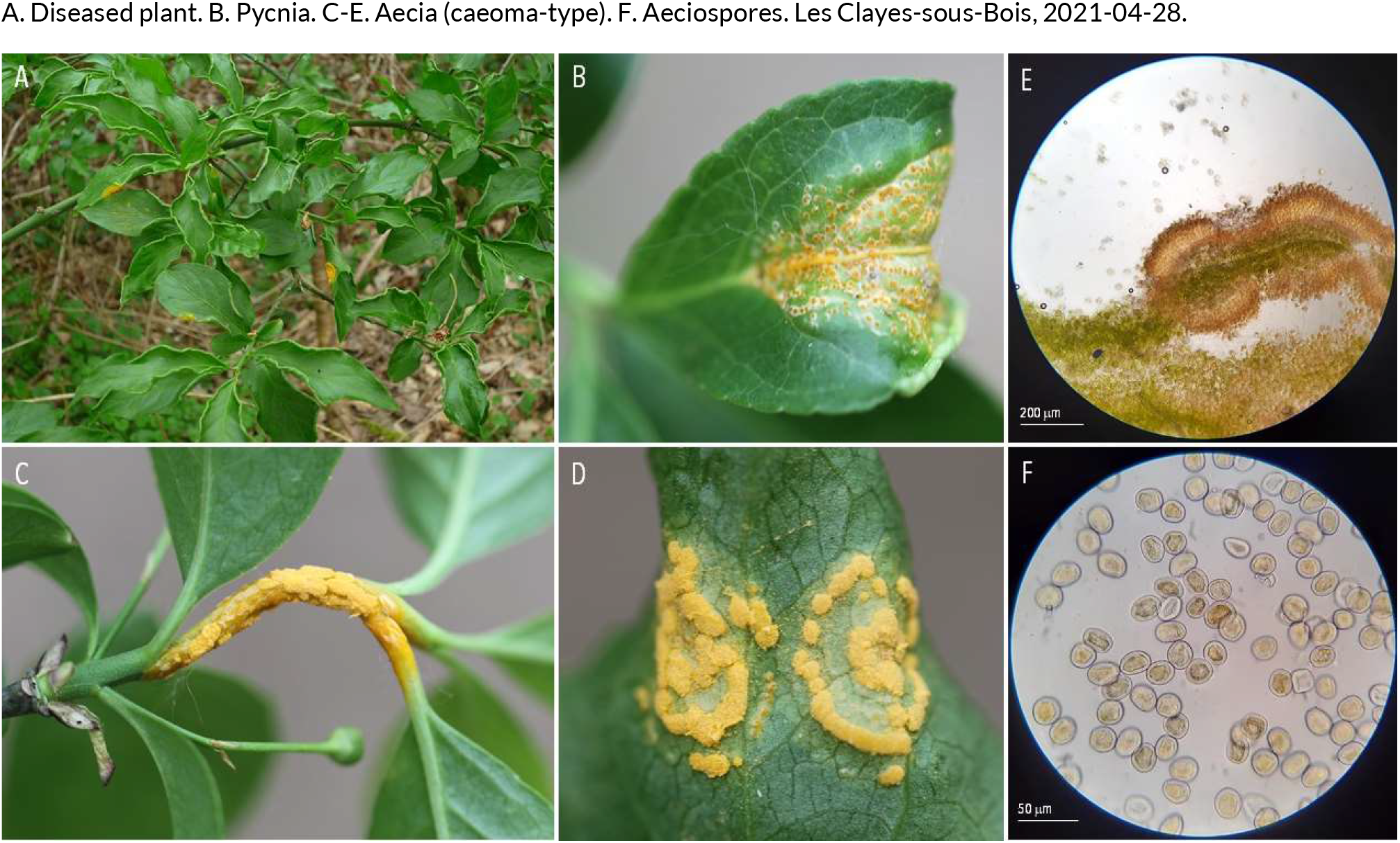
RUSTS | *Melampsora euonymi-caprearum* ex *Euonymus europaeus* (common spindle)

**14.**
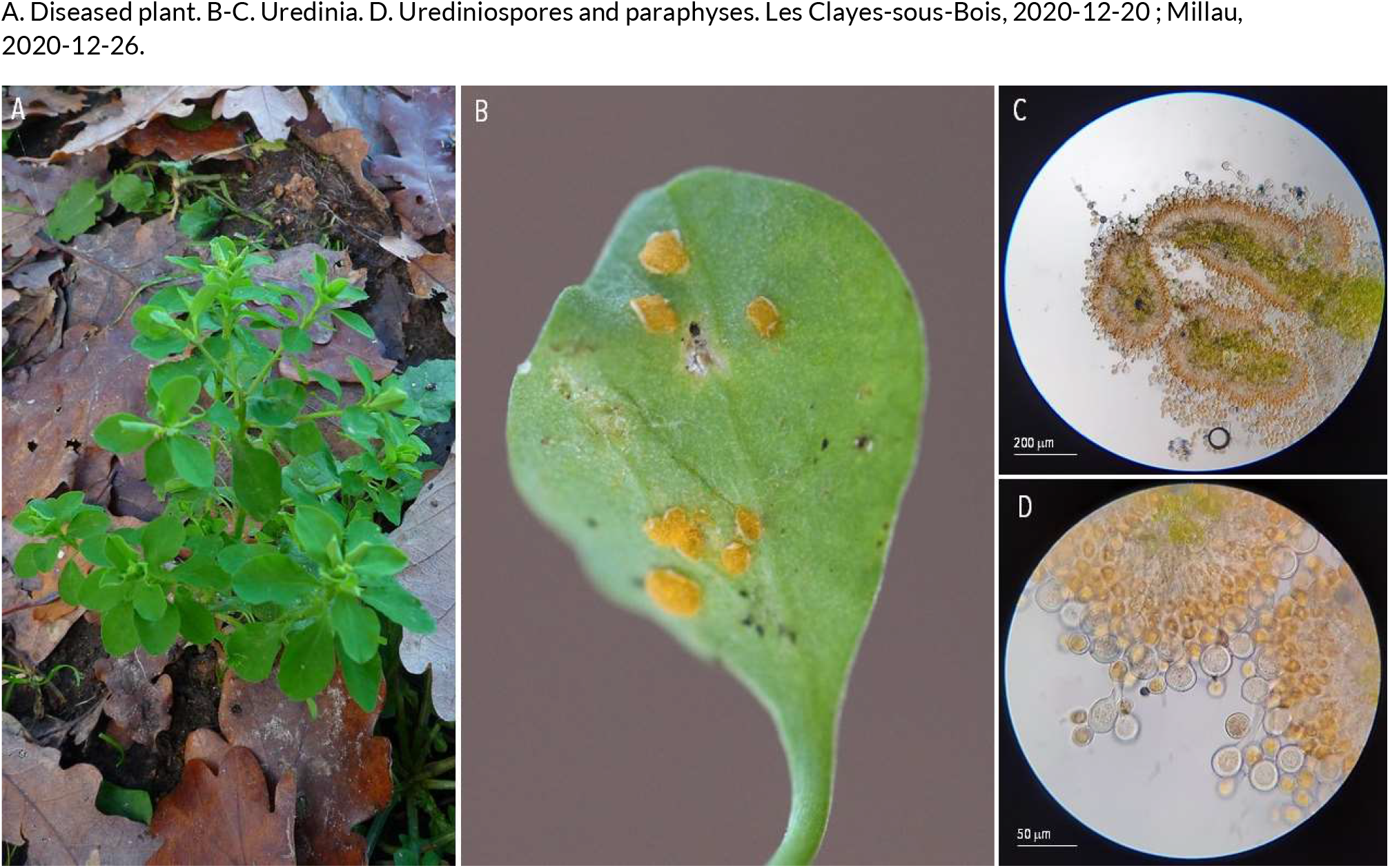
RUSTS | *Melampsora euphorbiae* ex *Euphorbia peplus* (petty spurge)

**15.**
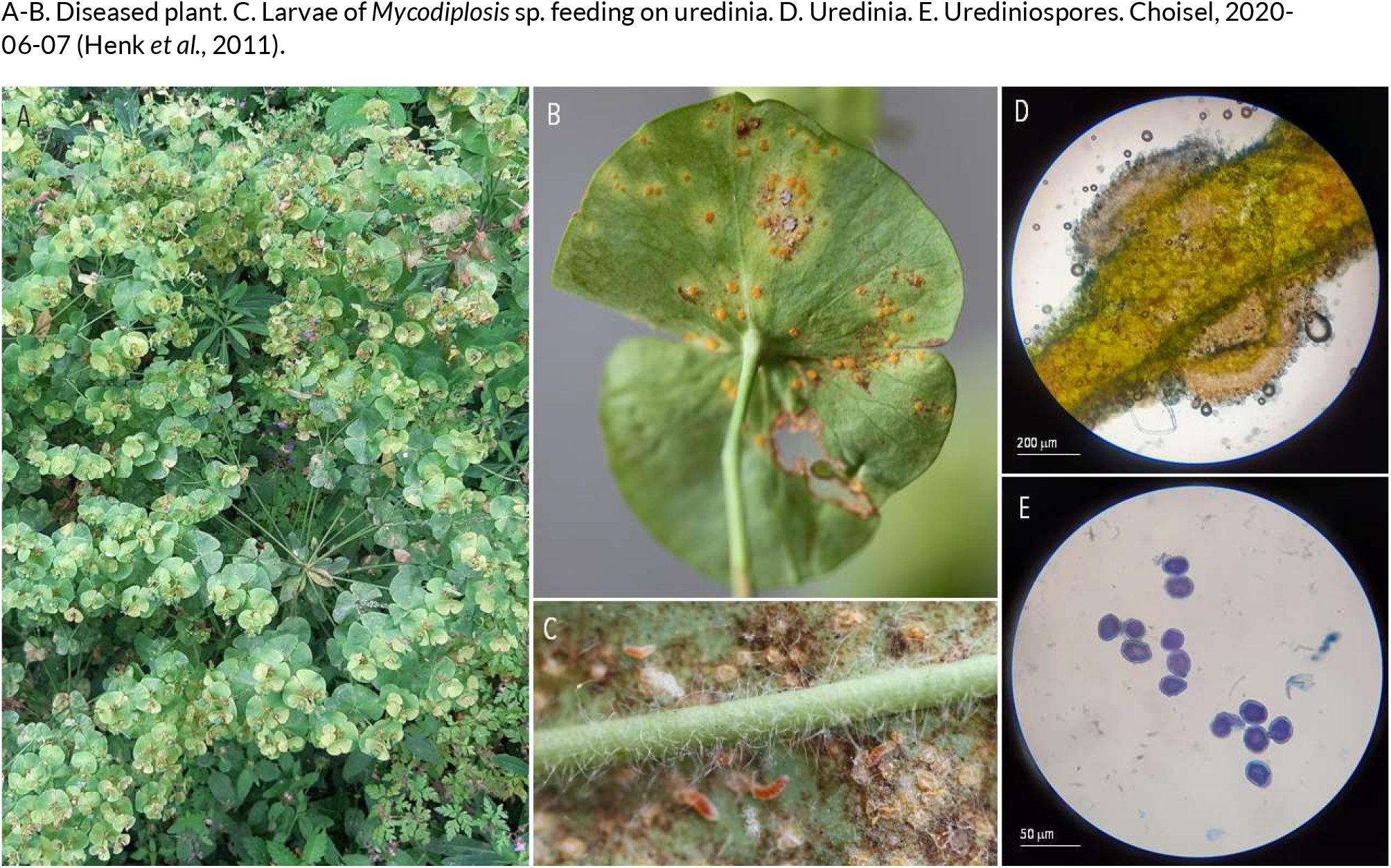
RUSTS | *Melampsora euphorbiae-amygdaloidis* ex *Euphorbia amygdaloides* (wood spurge)

**16.**
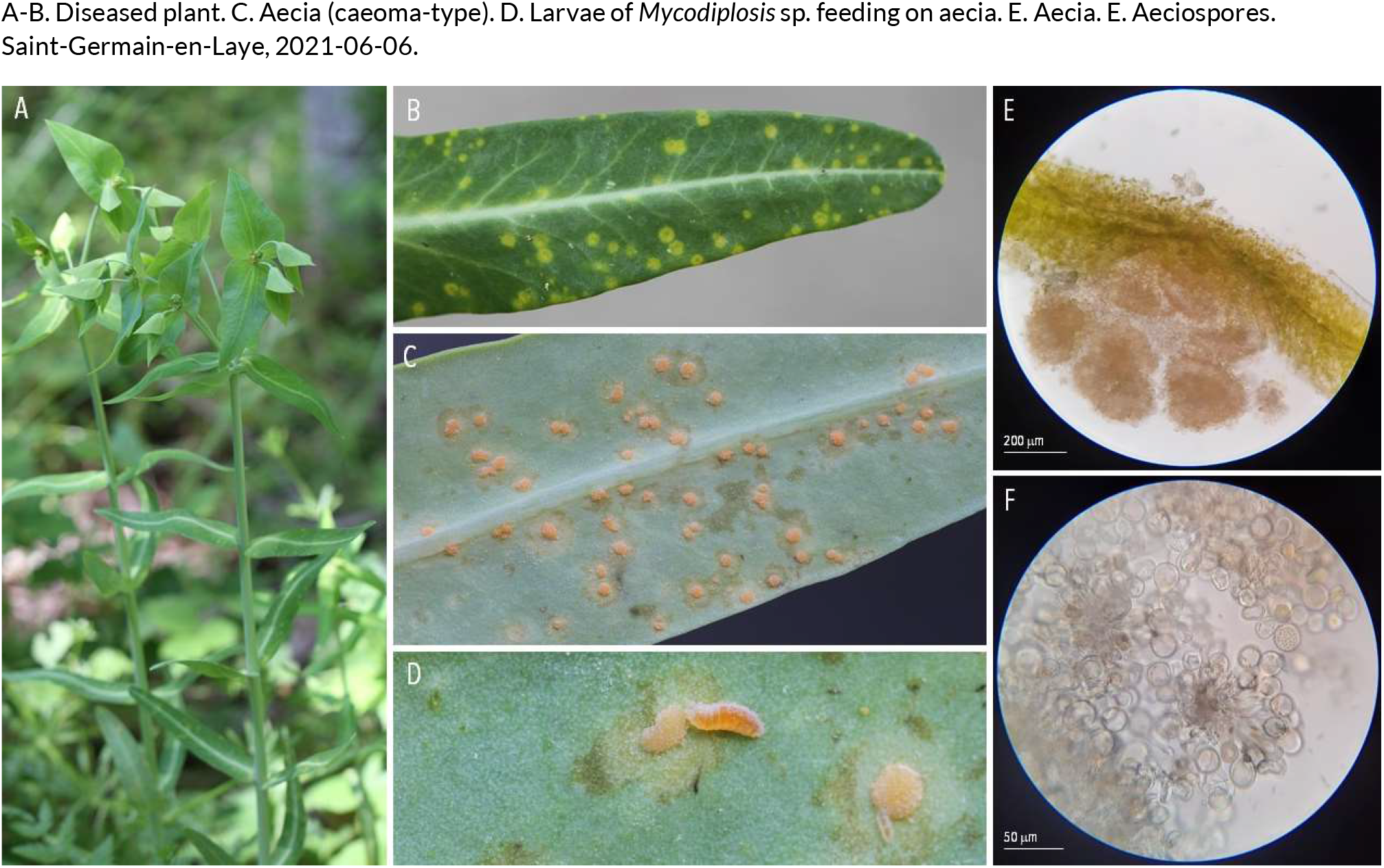
RUSTS | *Melampsora euphorbiae-dulcis* ex *Euphorbia lathyris* (caper spurge)

**17.**
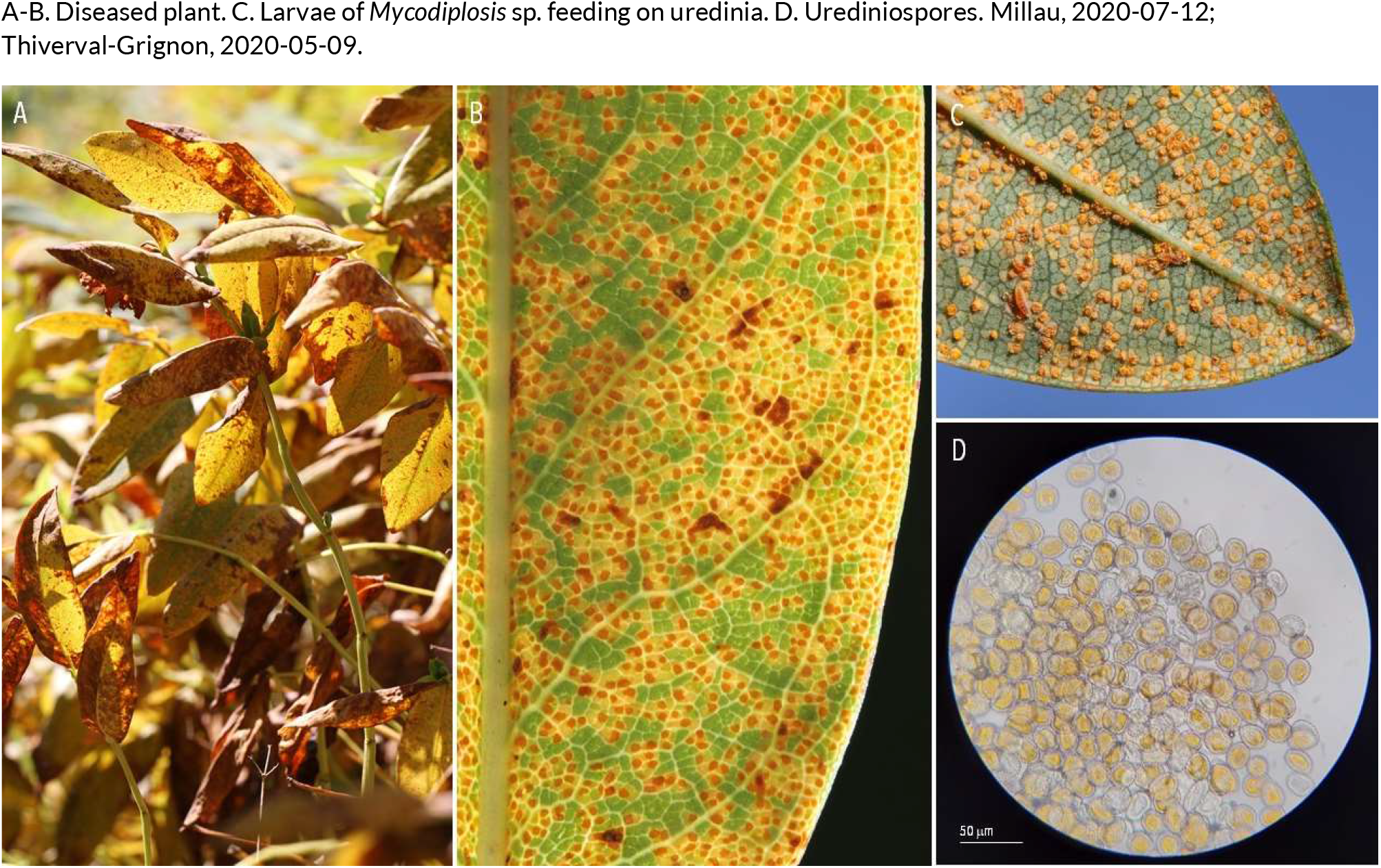
RUSTS | *Melampsora hypericorum* ex *Hypericum perforatum* (St. John’s wort)

**18.**
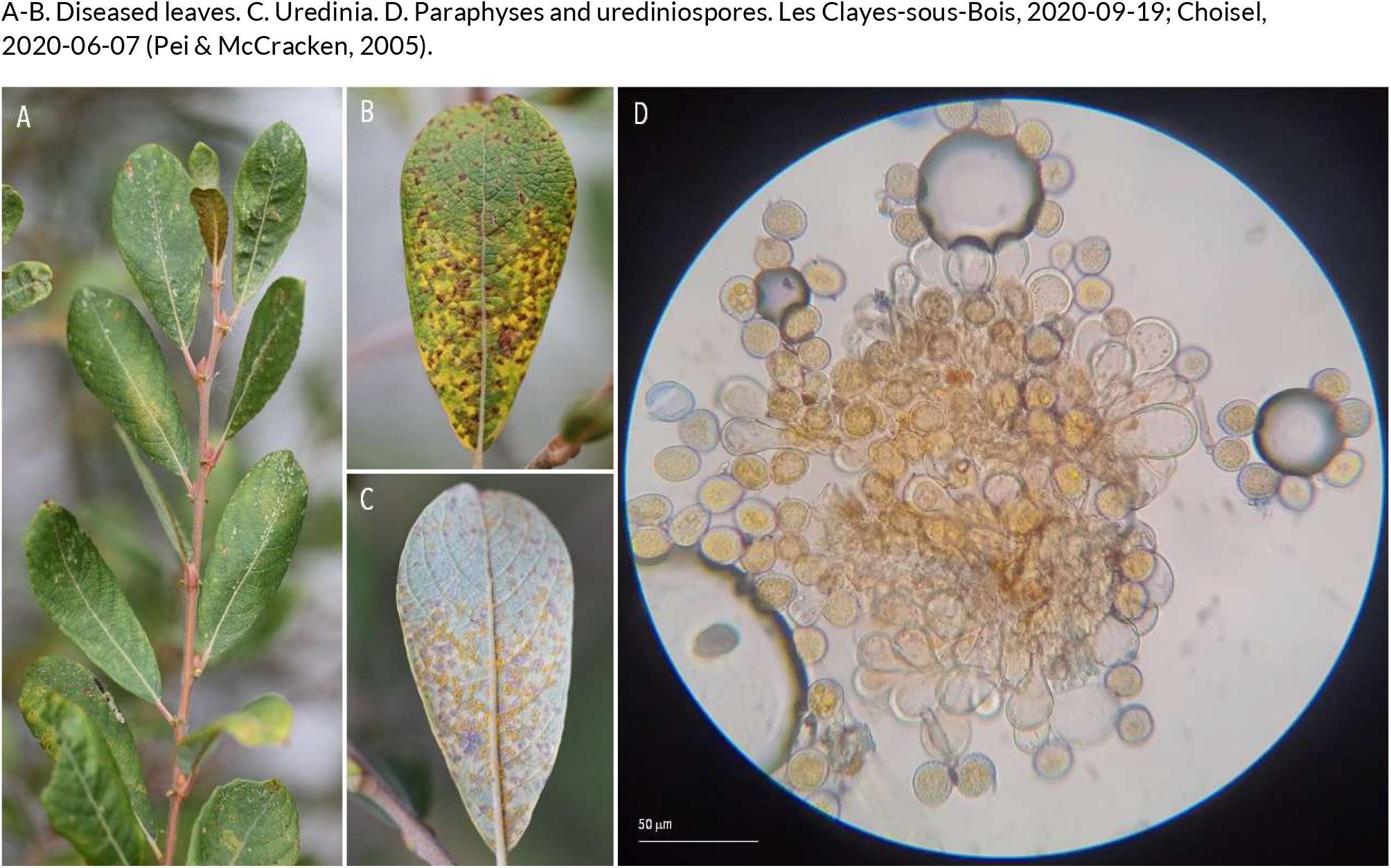
RUSTS | *Melampsora larici-epitea* (??) ex *Salix cinerea* (grey willow)

**19.**
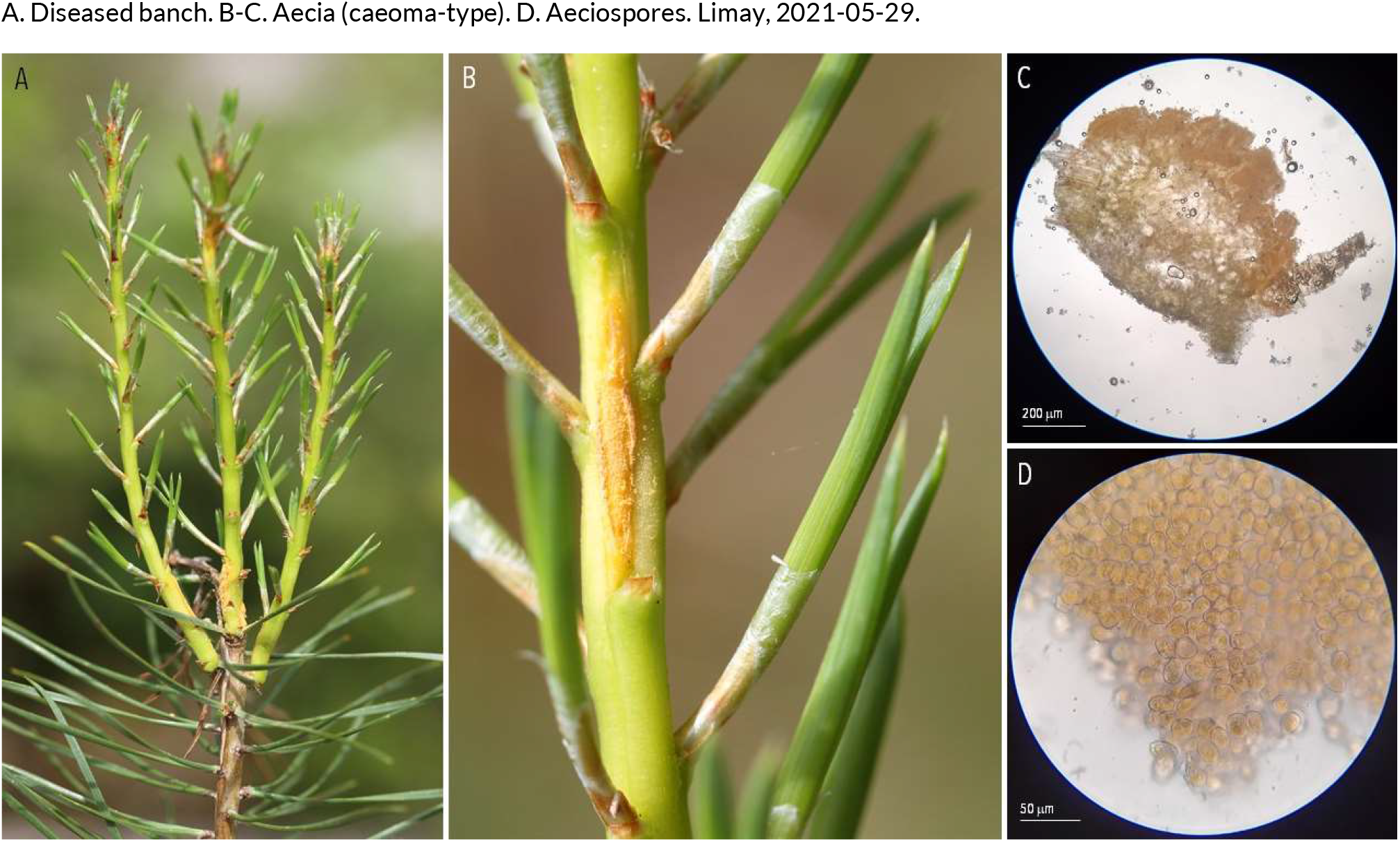
RUSTS | *Melampsora pinitorqua* ex *Pinus sylvestris* (scots pine)

**20.**
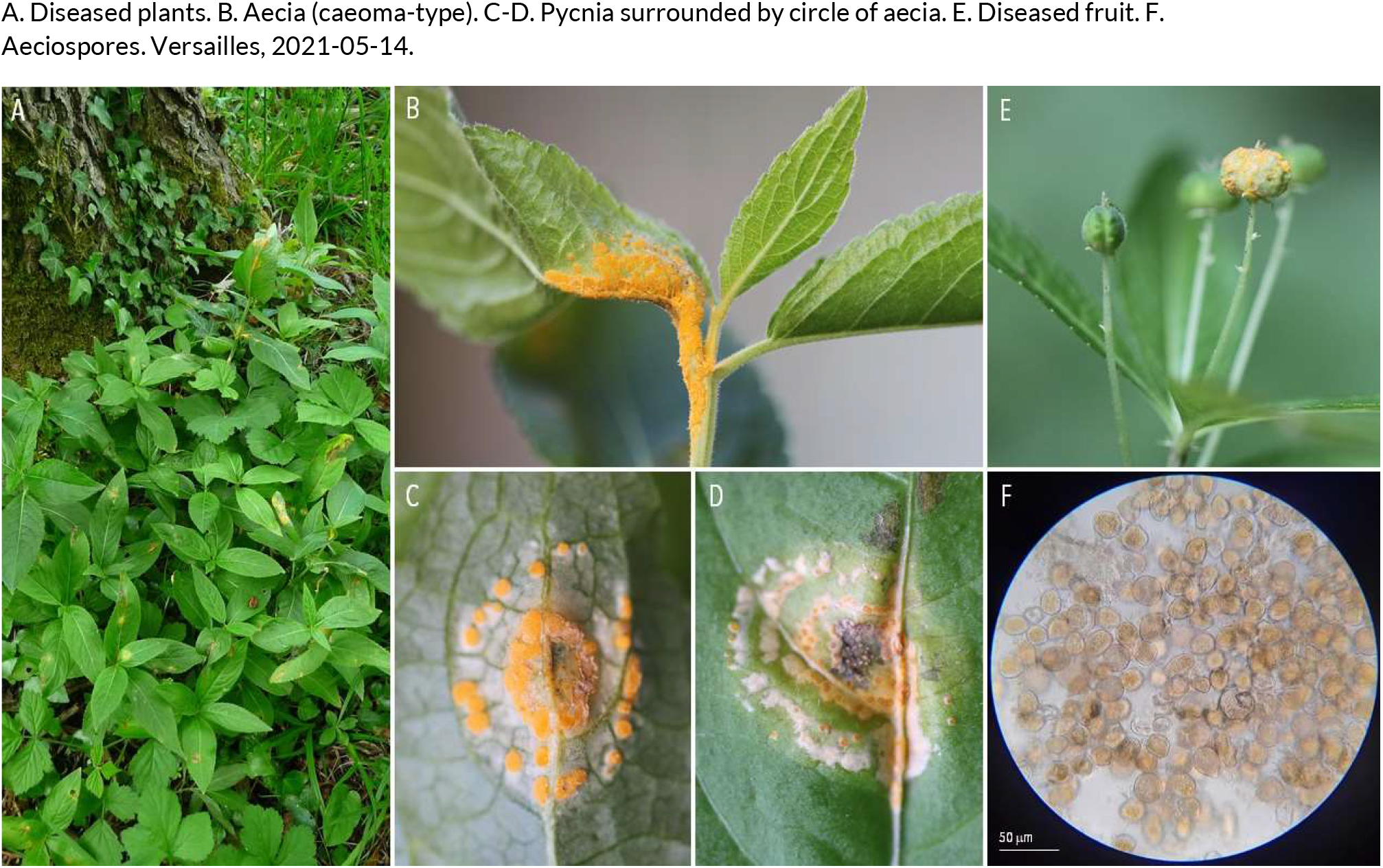
RUSTS | *Melampsora rostrupii* ex *Mercurialis perennis* (dog’s mercury)

**21.**
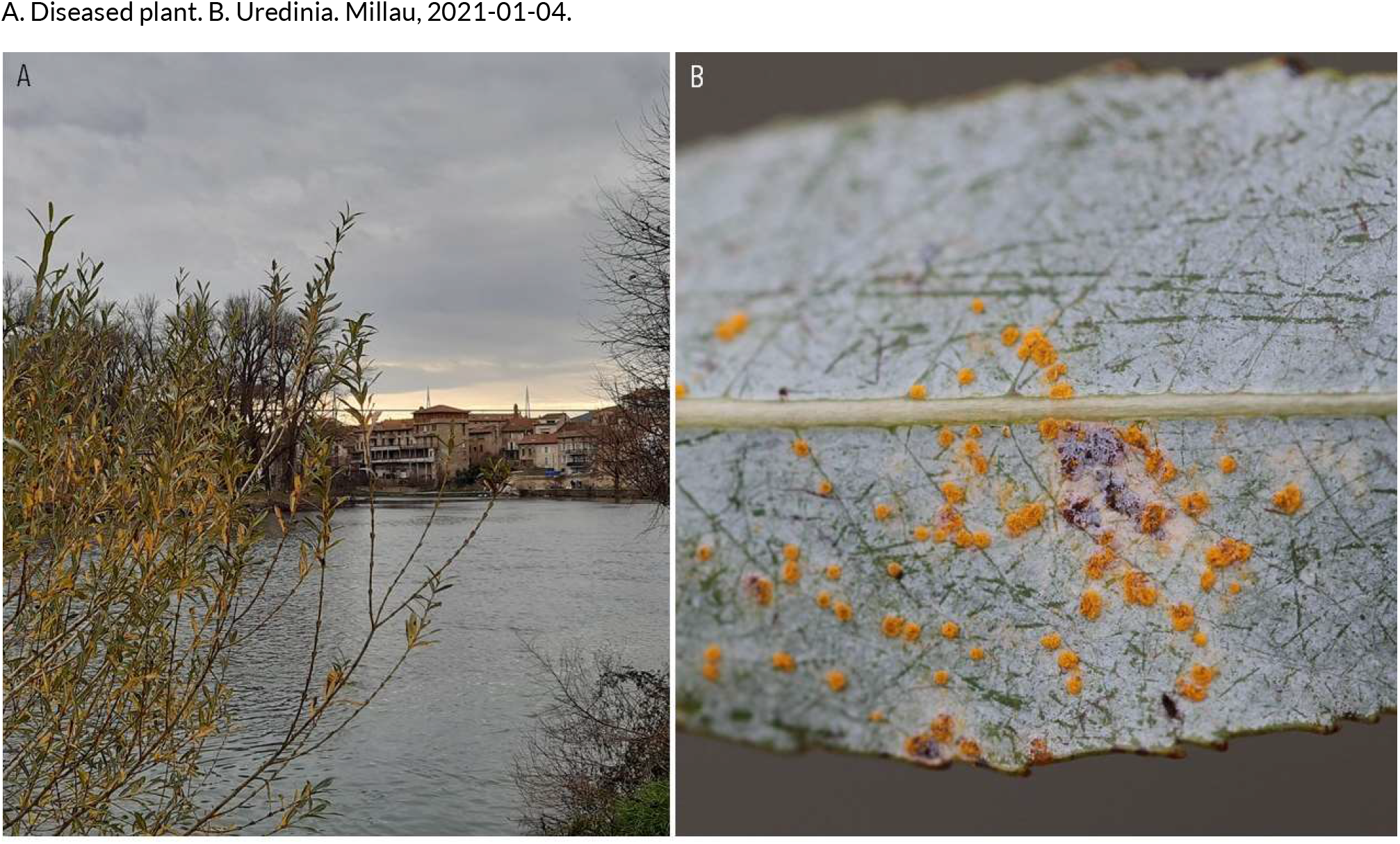
RUSTS | *Melampsora salicis-albae* (?) *ex Salix alba* (white willow)

**22.**
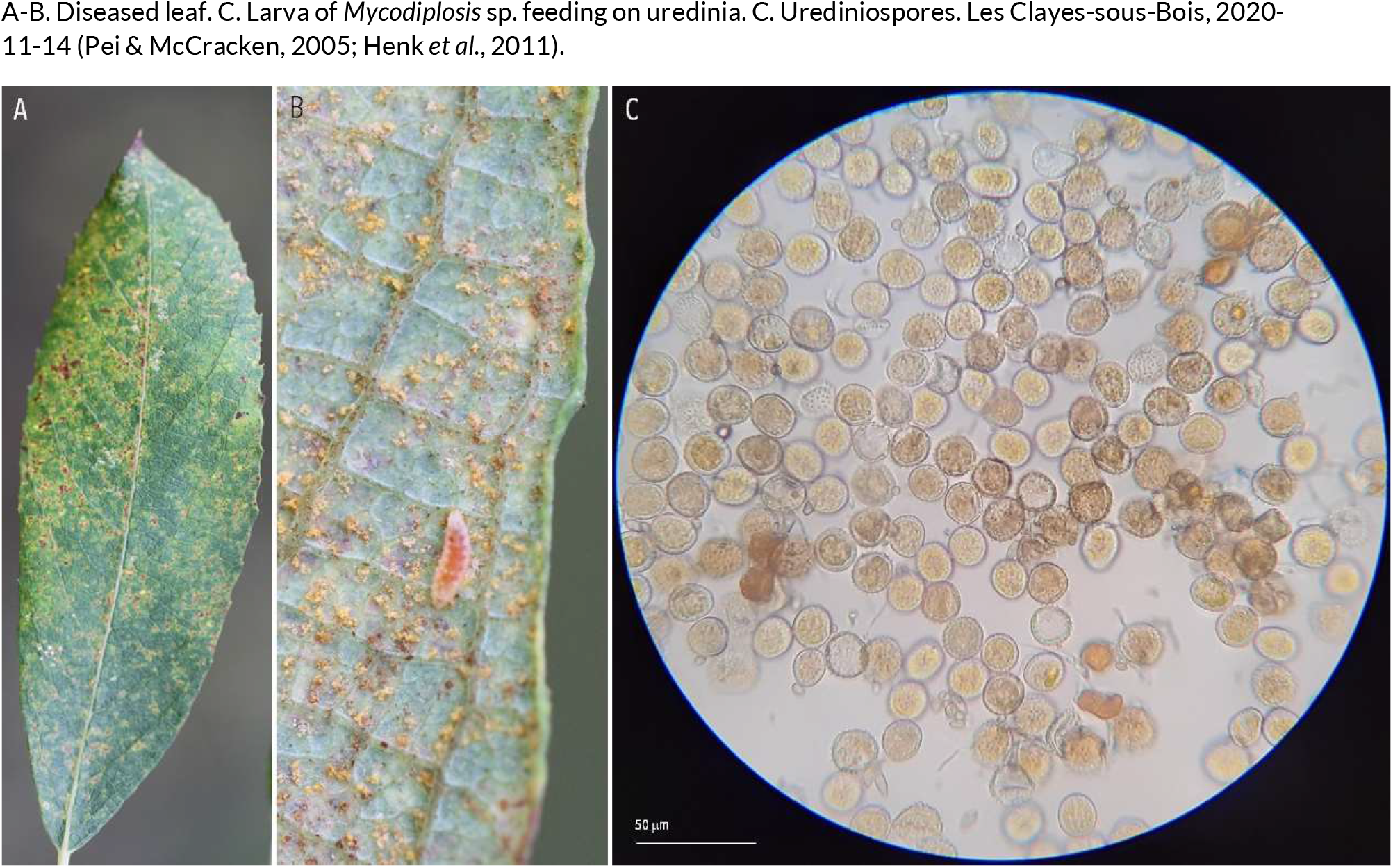
RUSTS | *Melampsora sp.* (??) ex *Salix atrocinerea* (large gray willow)

**23.**
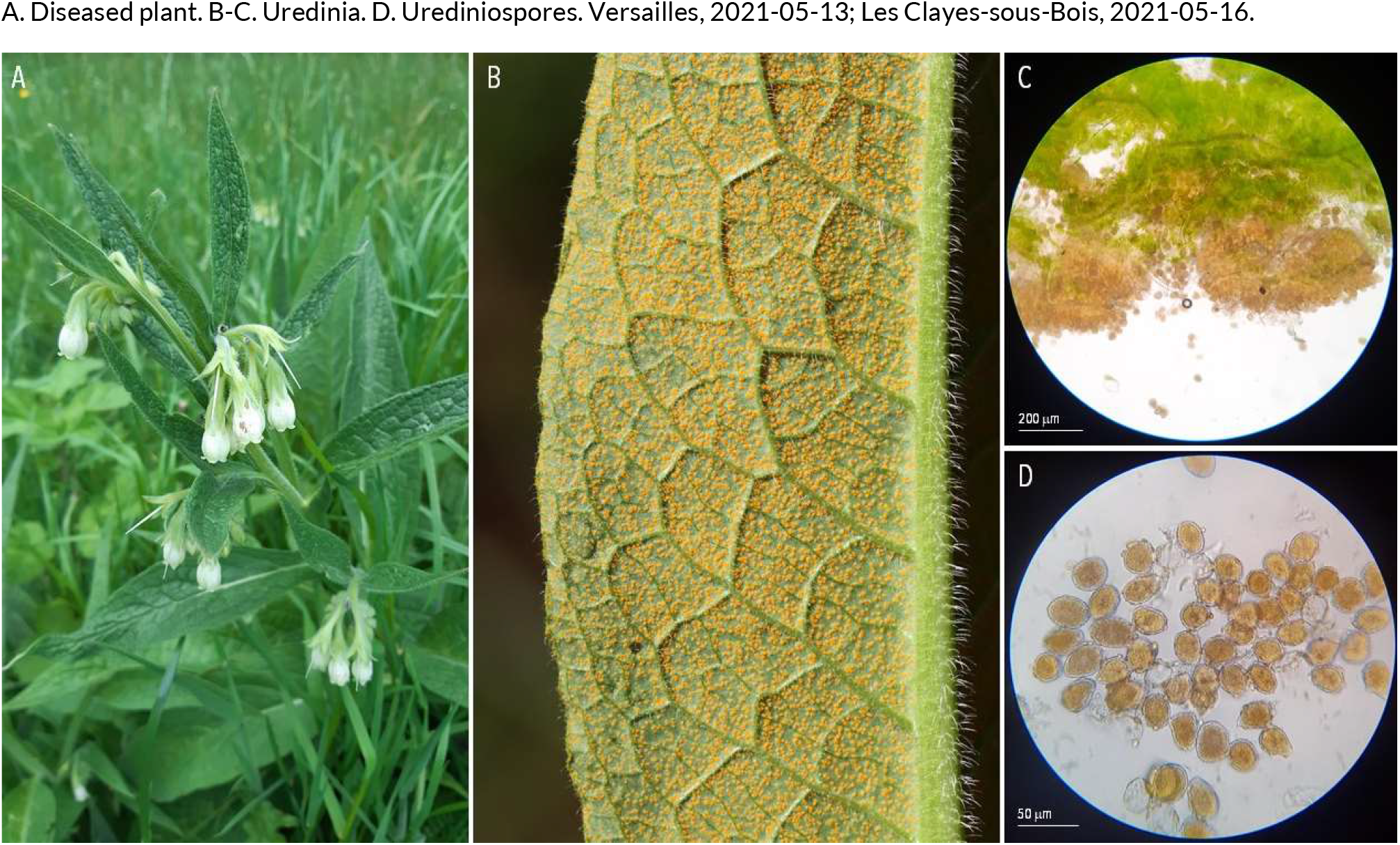
RUSTS | *Melampsorella symphyti* ex *Symphytum officinale* (common comfrey)

**24.**
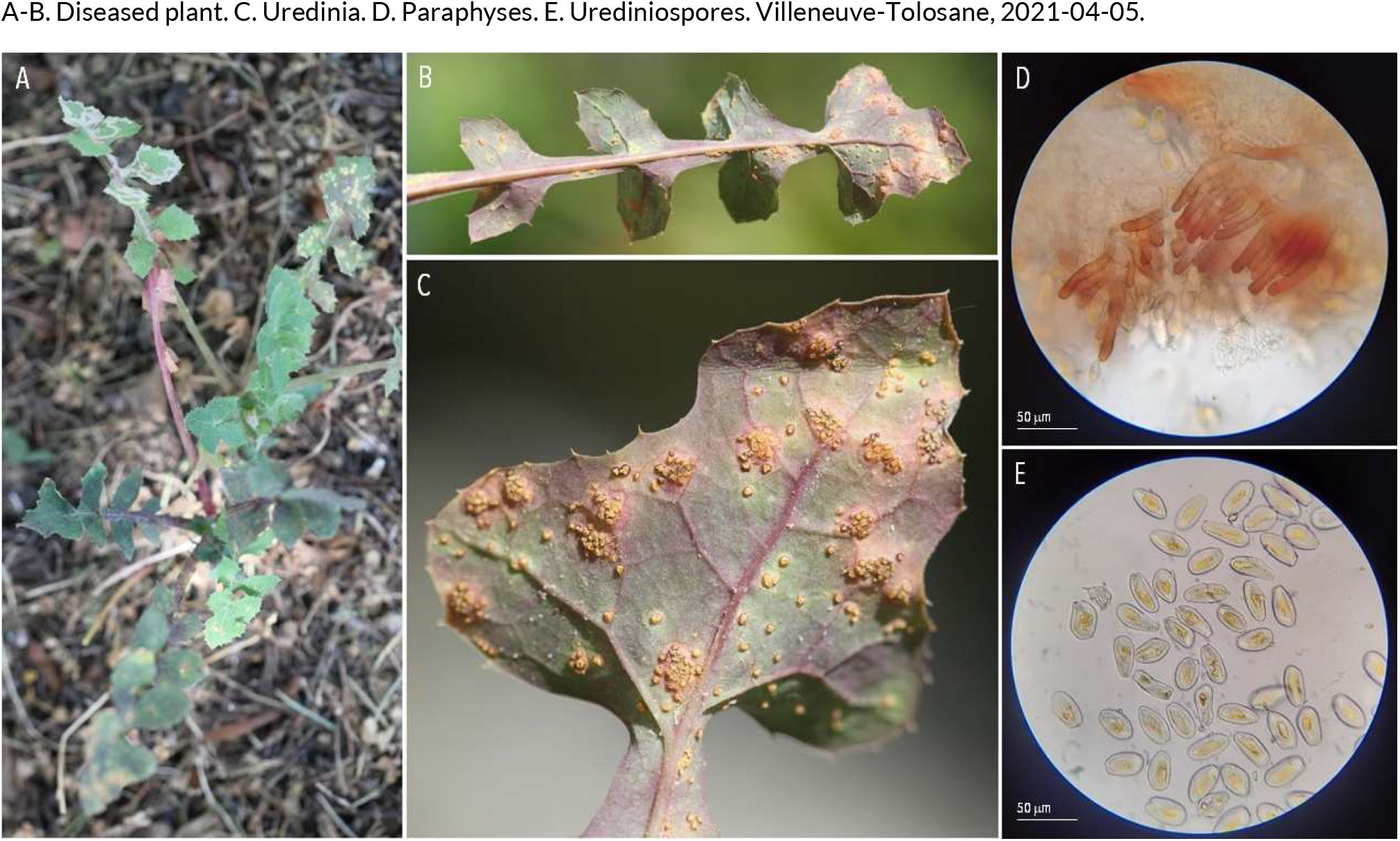
RUSTS | *Miyagia pseudosphaeria* (syn. Peristemma pseudosphaeria) ex *Sonchus oleraceus* (common sowthistle)

**25.**
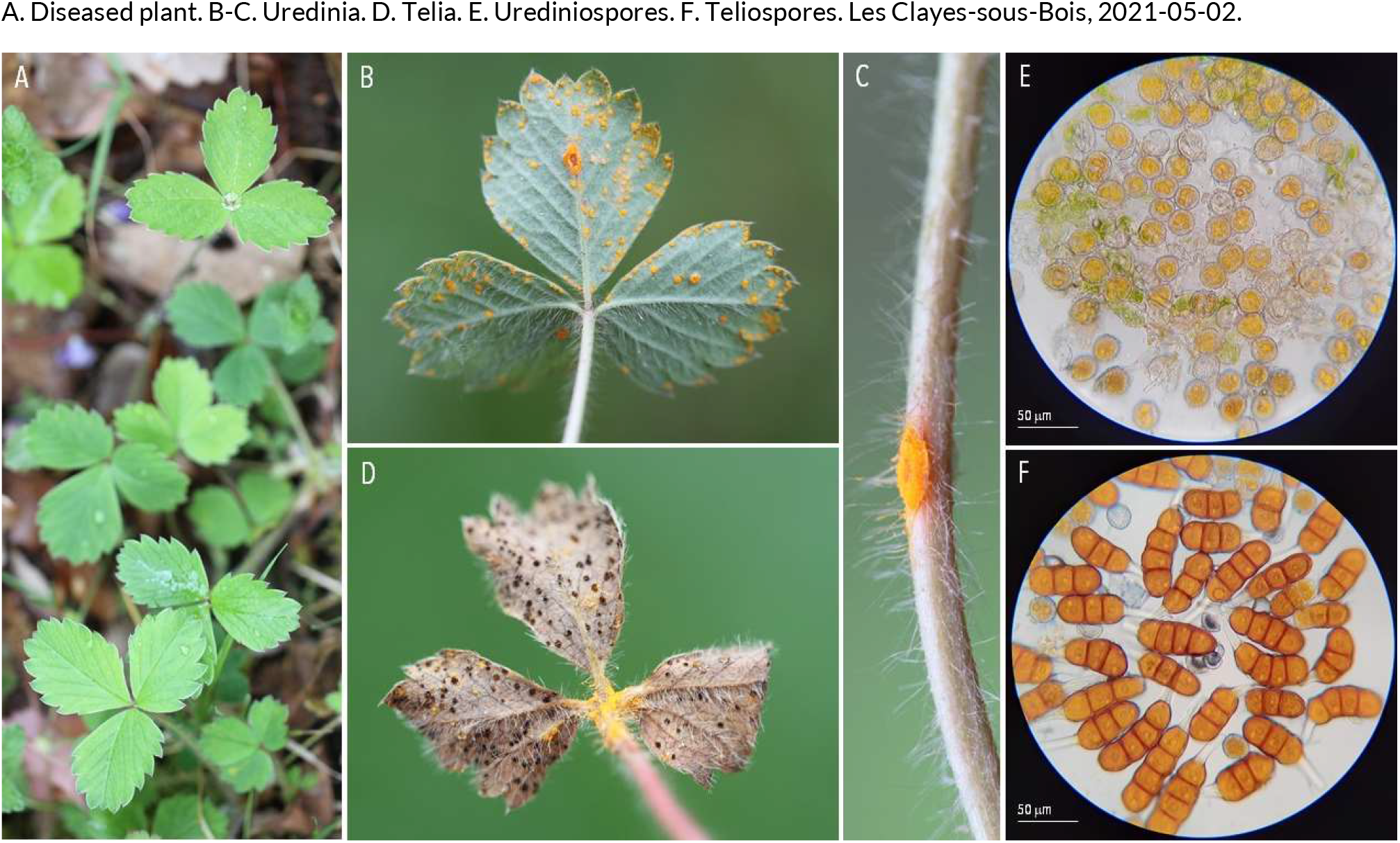
RUSTS | *Phragmidium fragariae* ex *Potentilla sterilis* (strawberryleaf cinquefoil)

**26.**
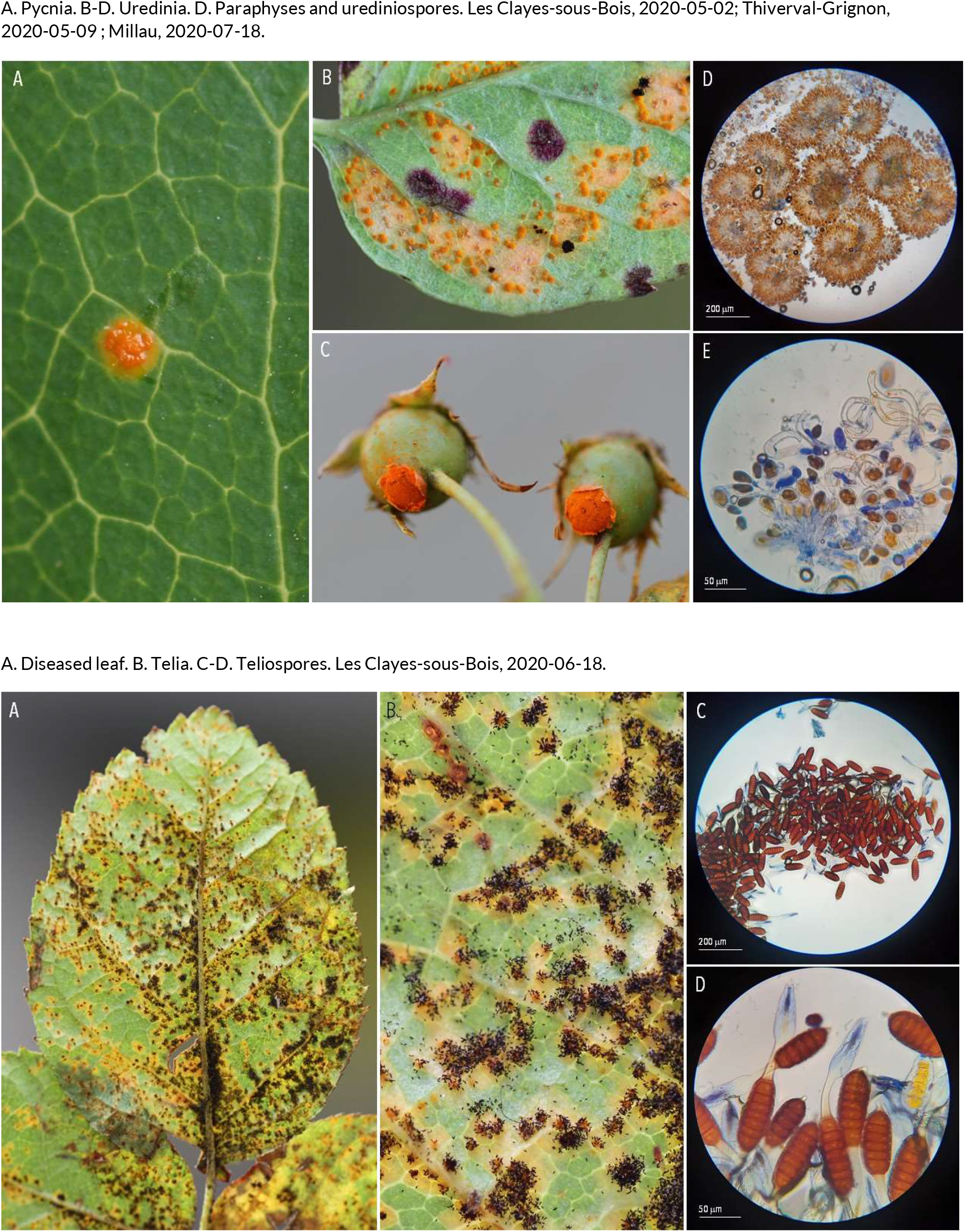
RUSTS | *Phragmidium mucronatum* ex *Rosa sp*. (rose)

**27.**
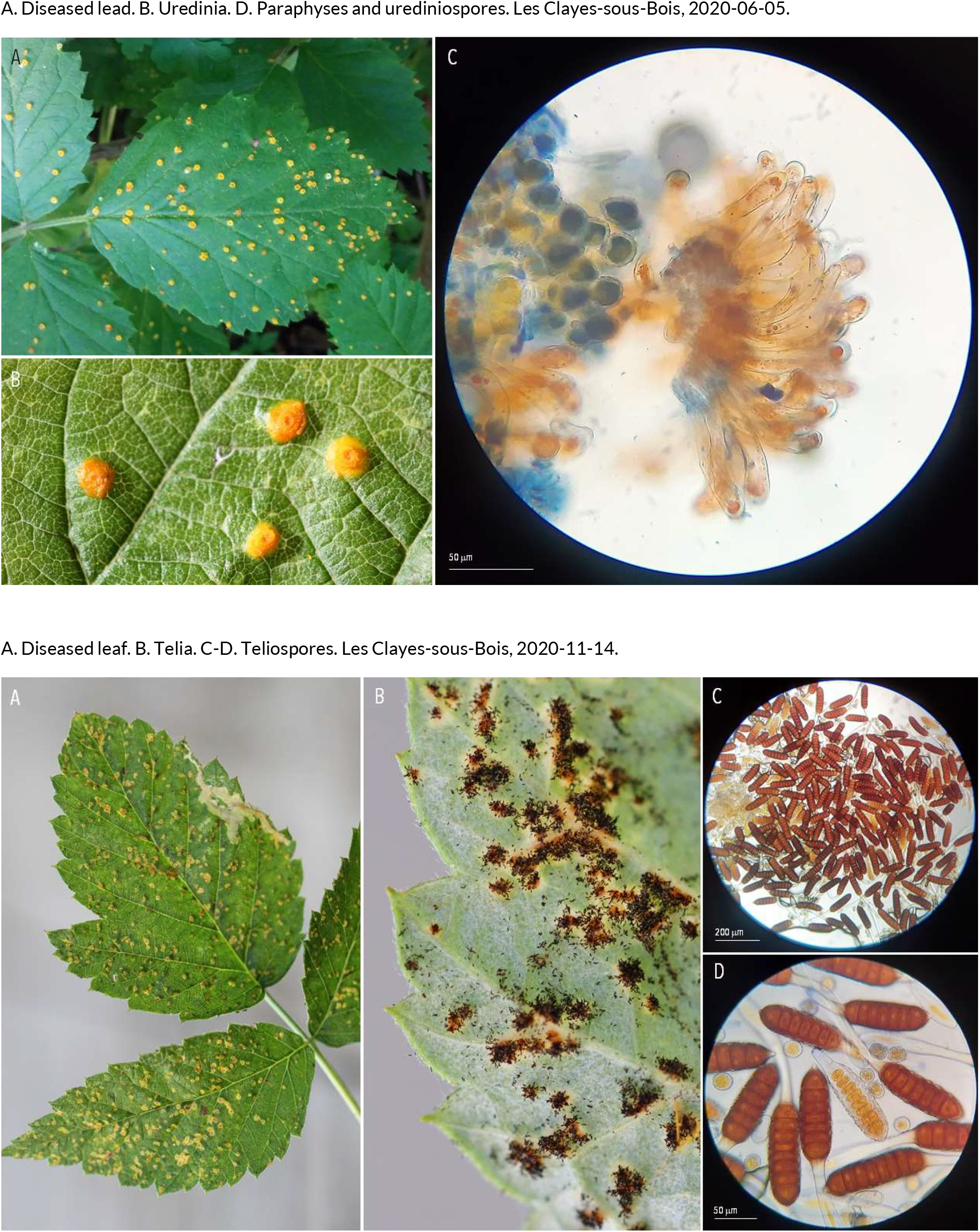
RUSTS | *Phragmidium rubi-idaei* ex *Rubus idaeus* (raspberry)

**28.**
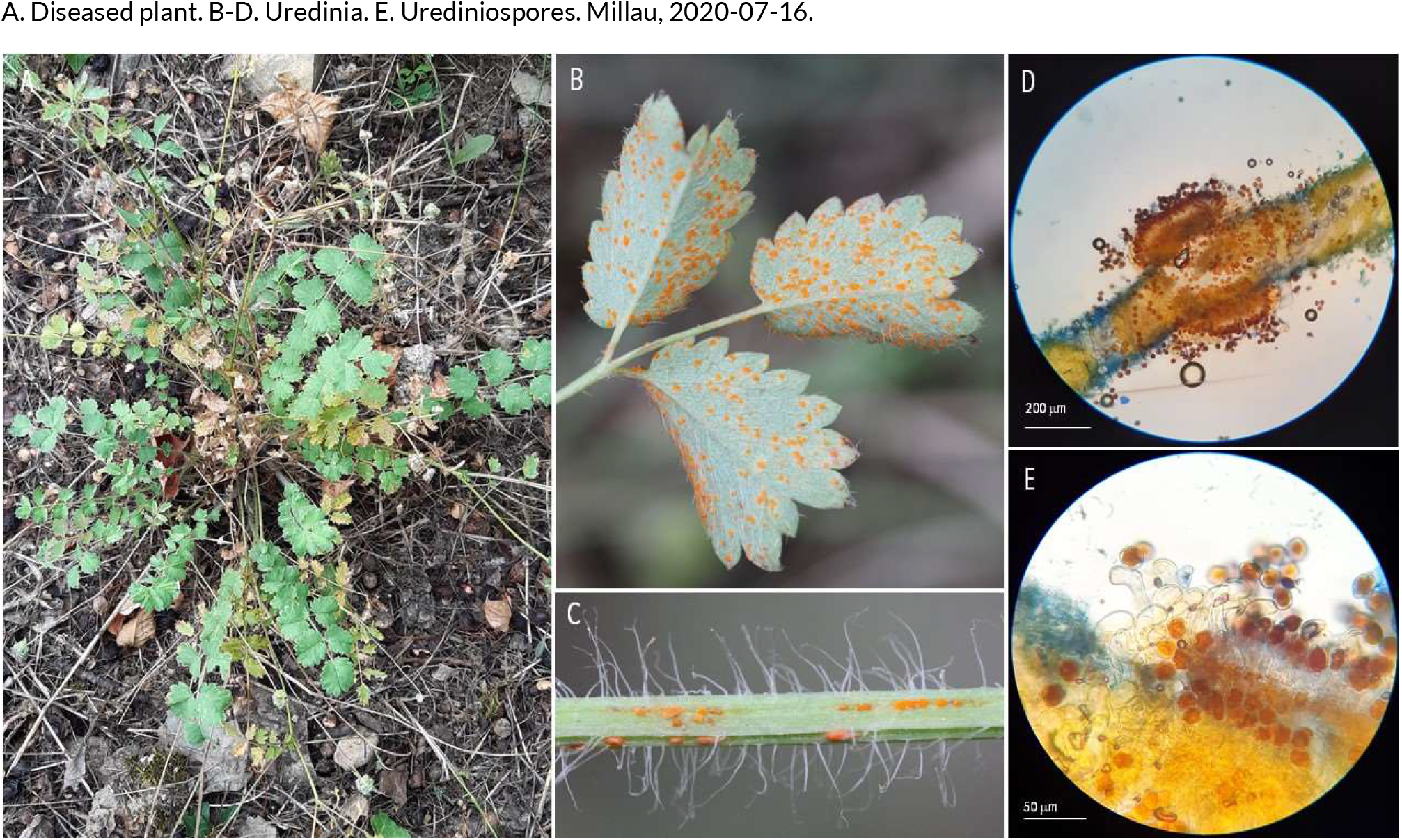
RUSTS | *Phragmidium sanguisorbae* ex *Sanguisorba minor* (salad burnet)

**29.**
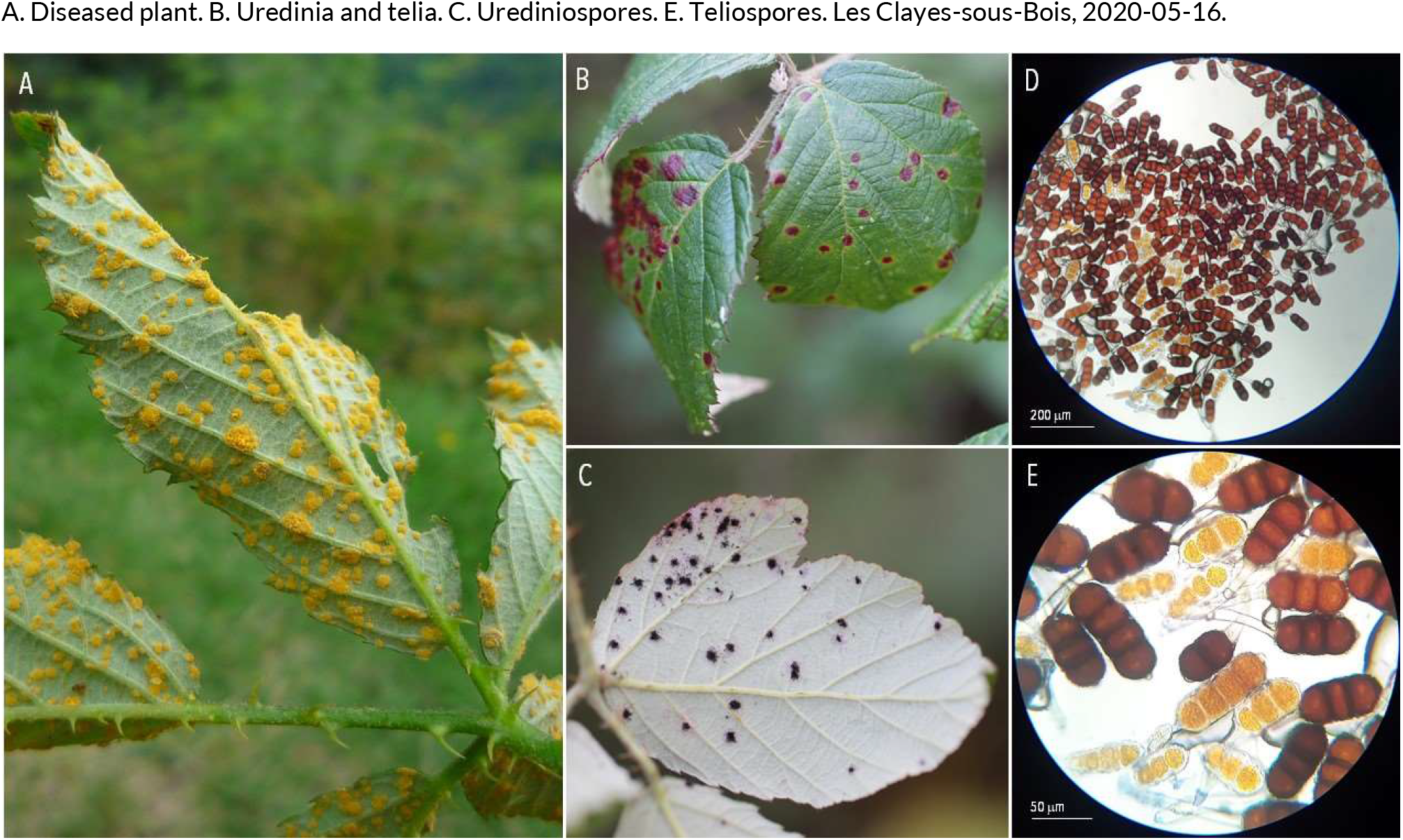
RUSTS | *Phragmidium violaceum* ex *Rubus fruticosus* (bramble)

**30.**
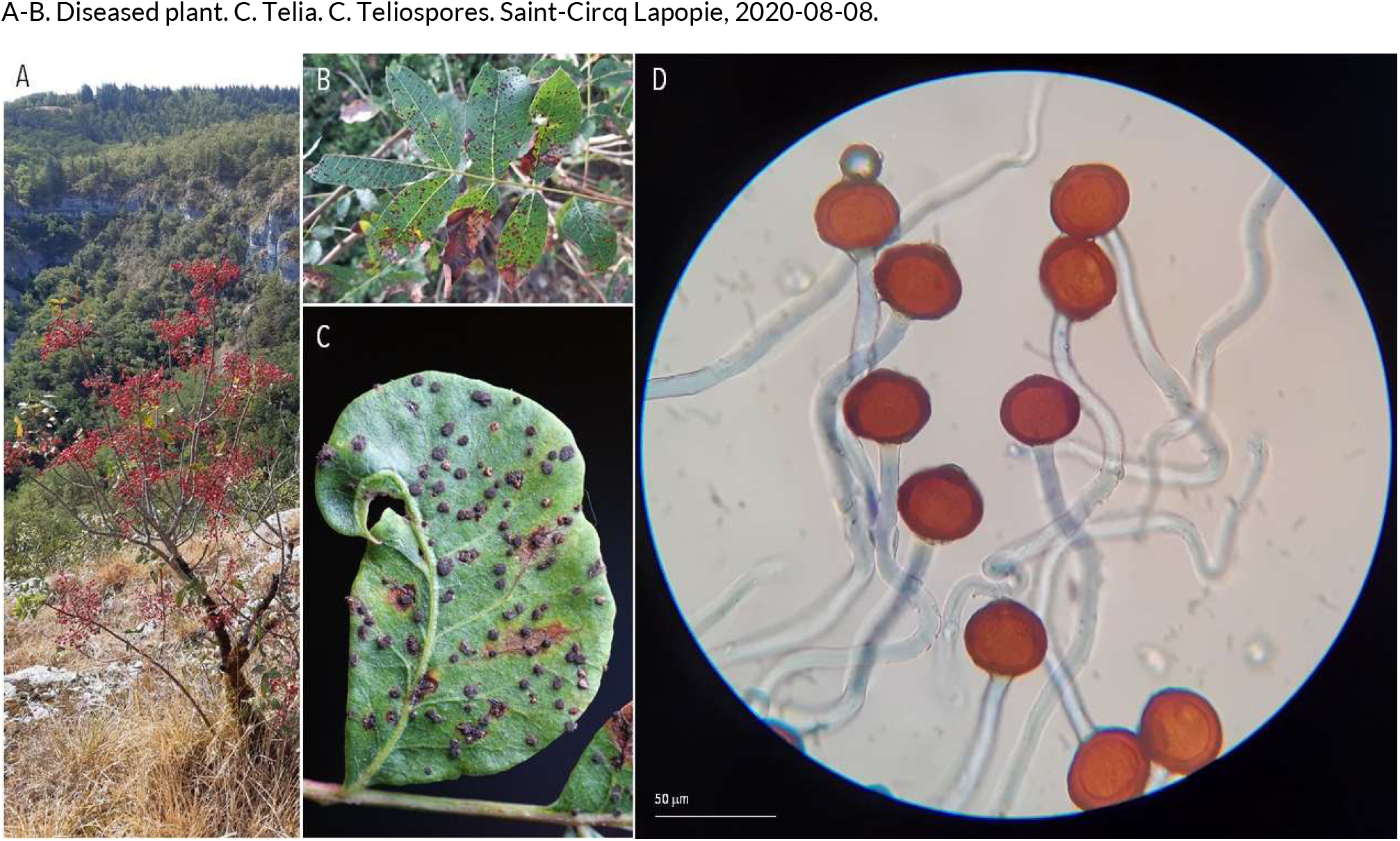
RUSTS | *Pileolaria terebenthi* ex *Pistachio lentiscus* (lentisk)

**31.**
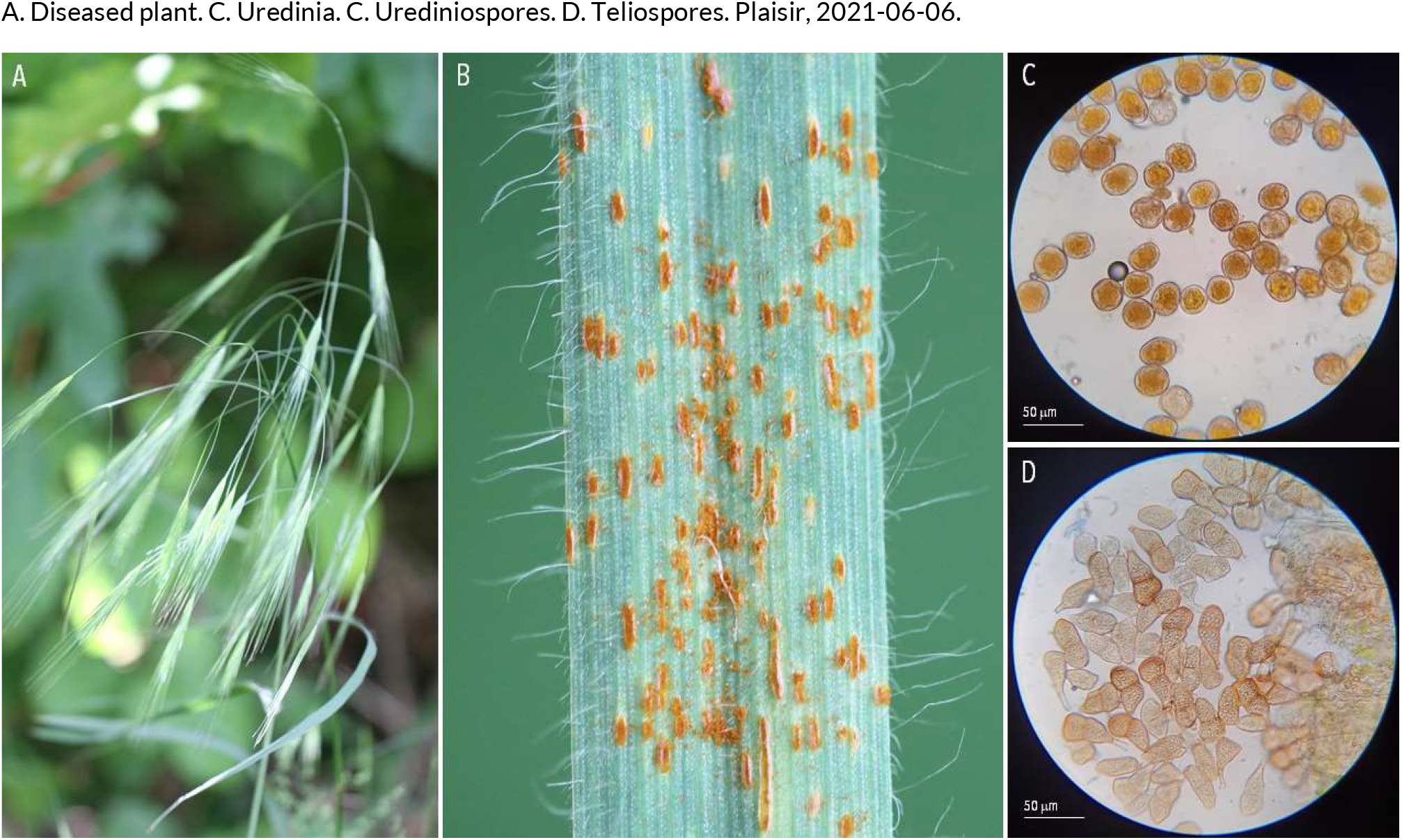
RUSTS | *Puccinia bromina* ex *Bromus sterilis* (barren brome)

**32.**
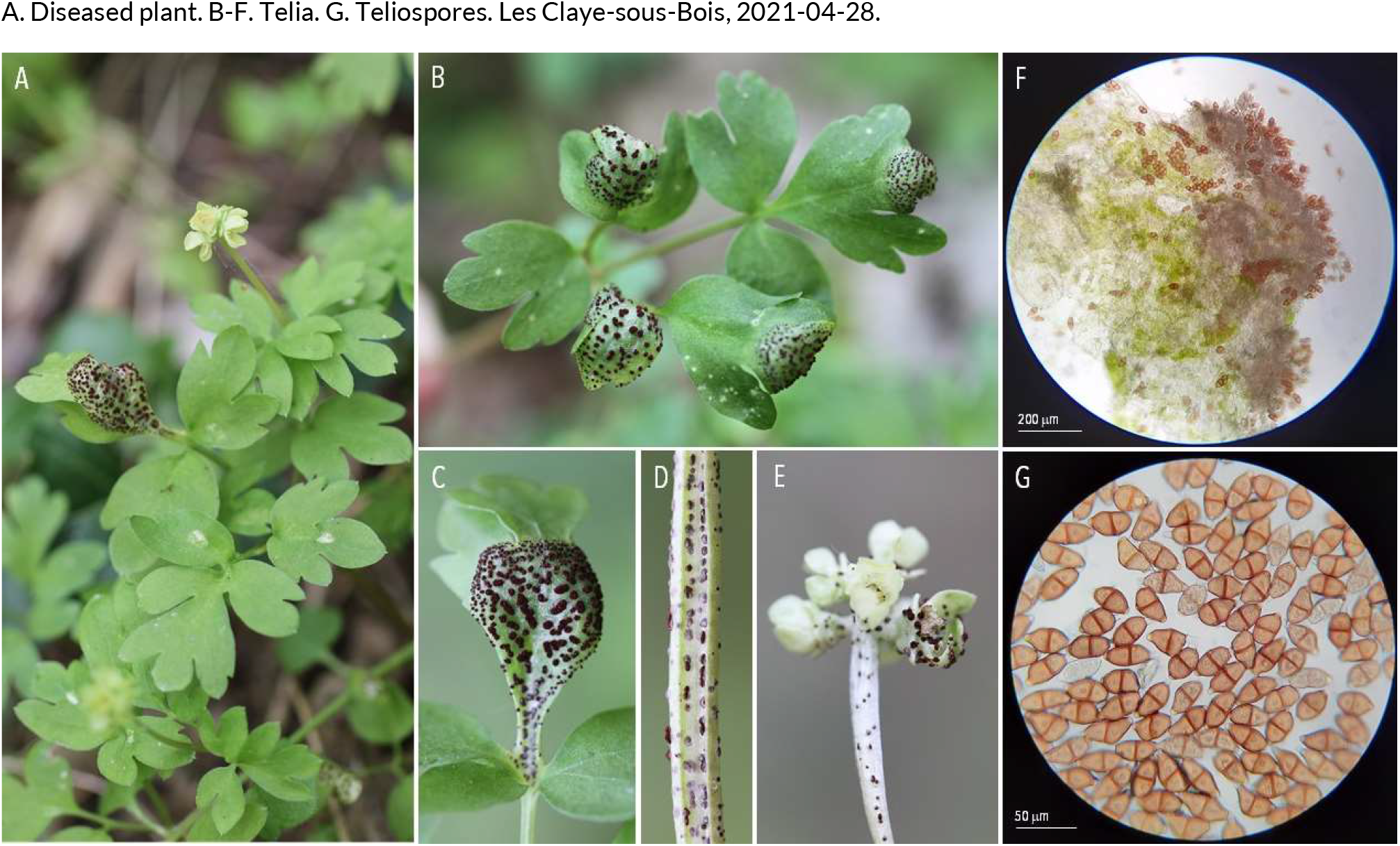
RUSTS | *Puccinia adoxae* ex *Adoxa moschatellina* (moschatel)

**33.**
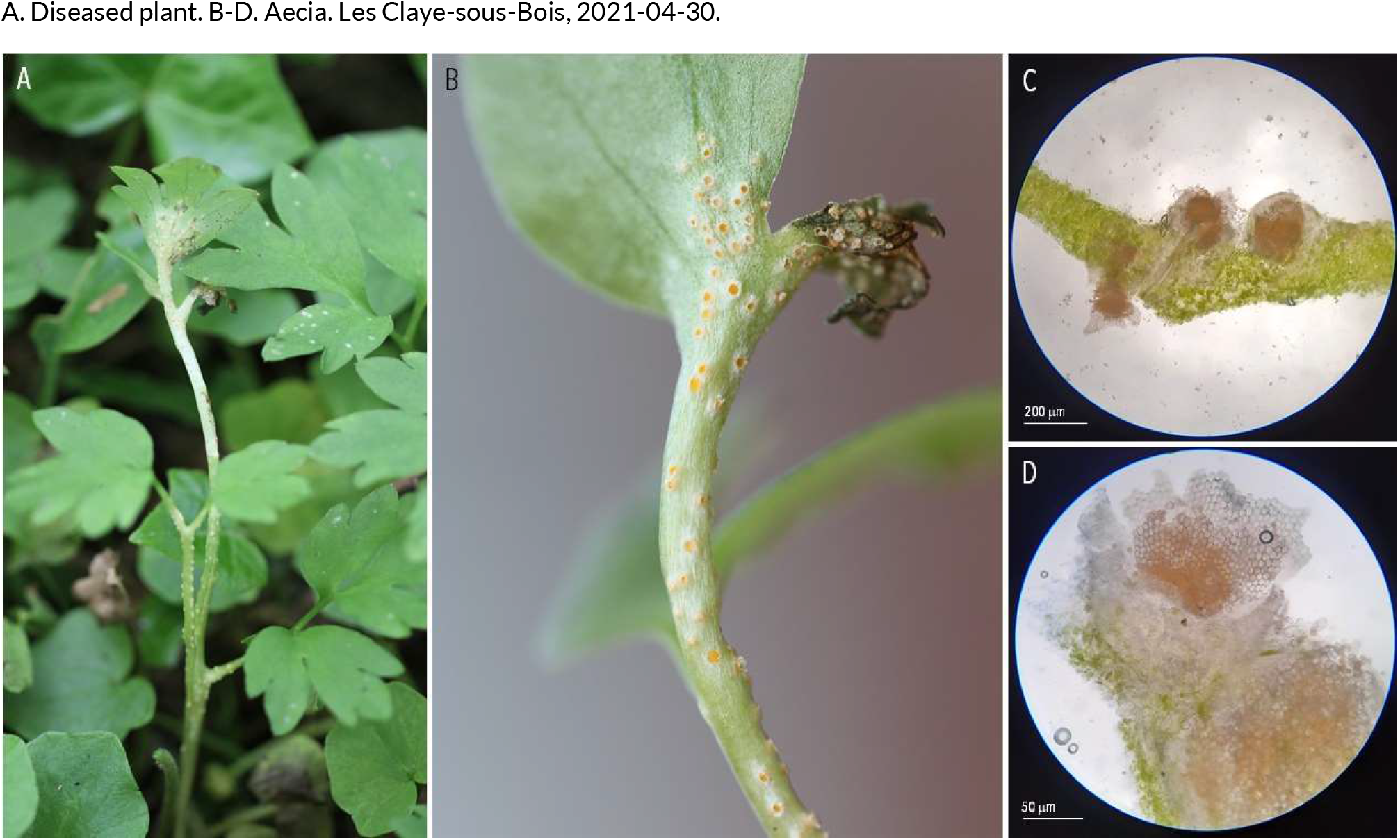
RUSTS | *Puccinia albescens* Adoxa moschatellina (moschatel)

**34.**
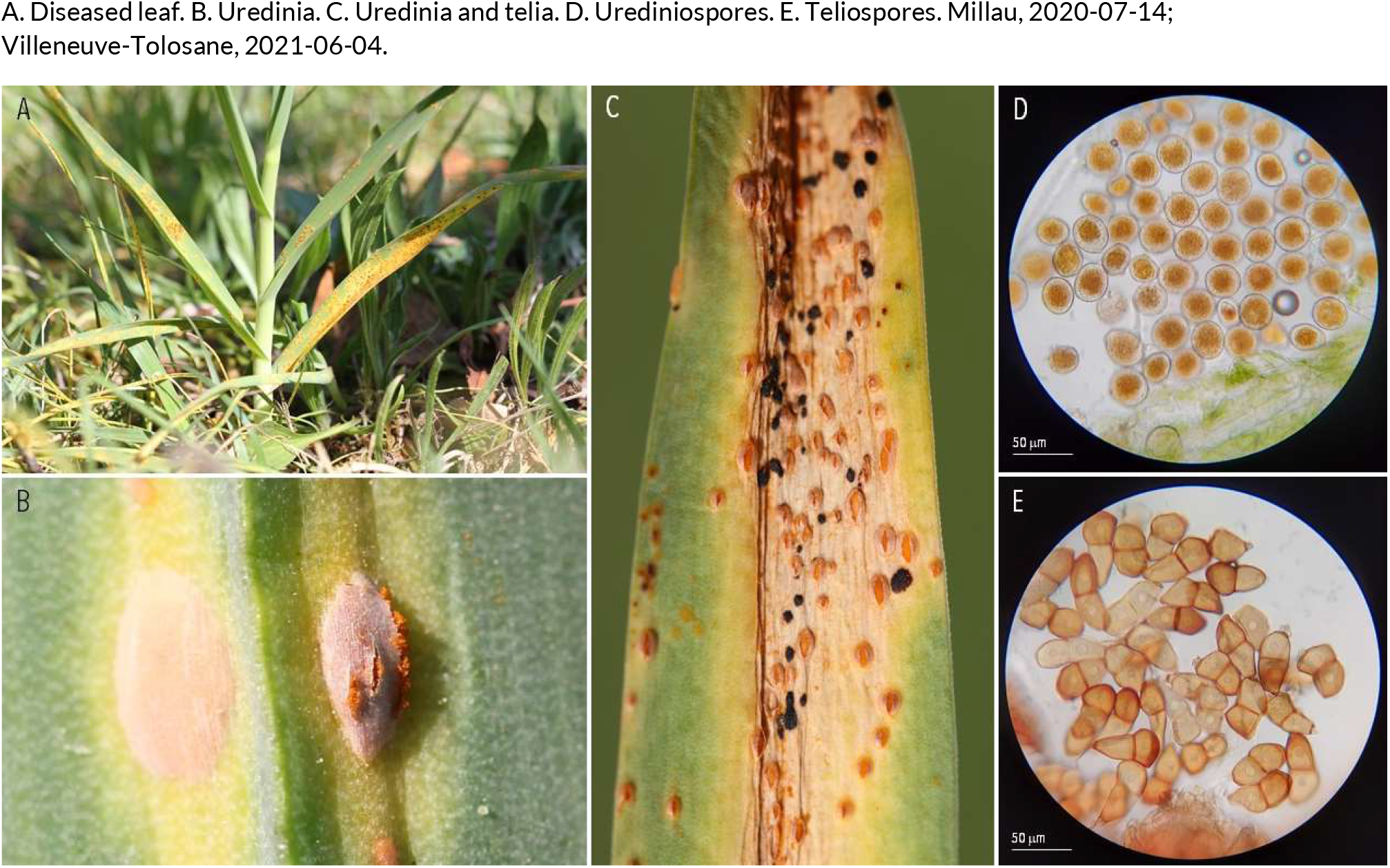
RUSTS | *Puccinia allii* ex *Allium polyanthum* (perennial sweet leek)

**35.**
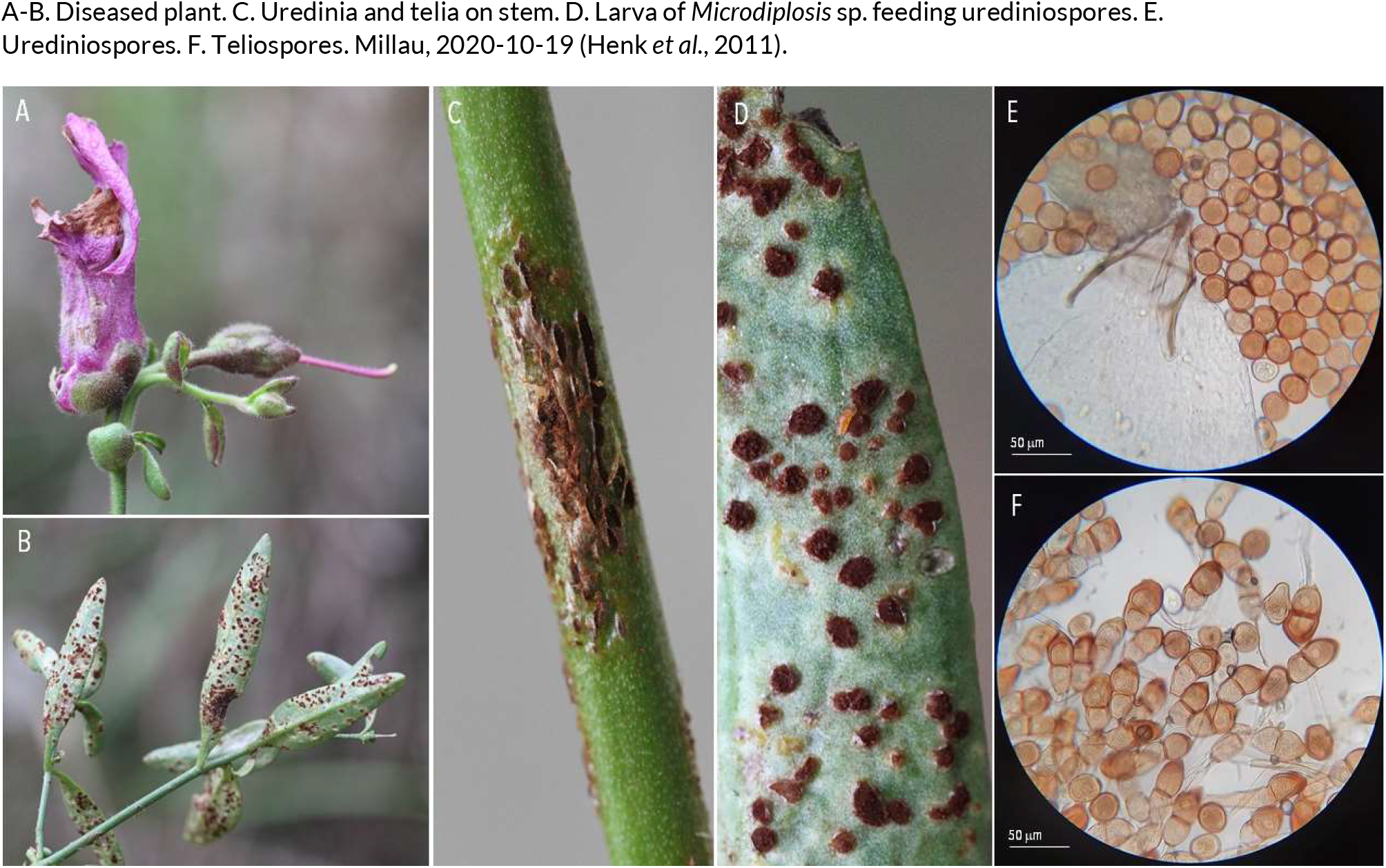
RUSTS | *Puccinia antirrhini* ex *Antirrhinum majus* (common snapdragon)

**36.**
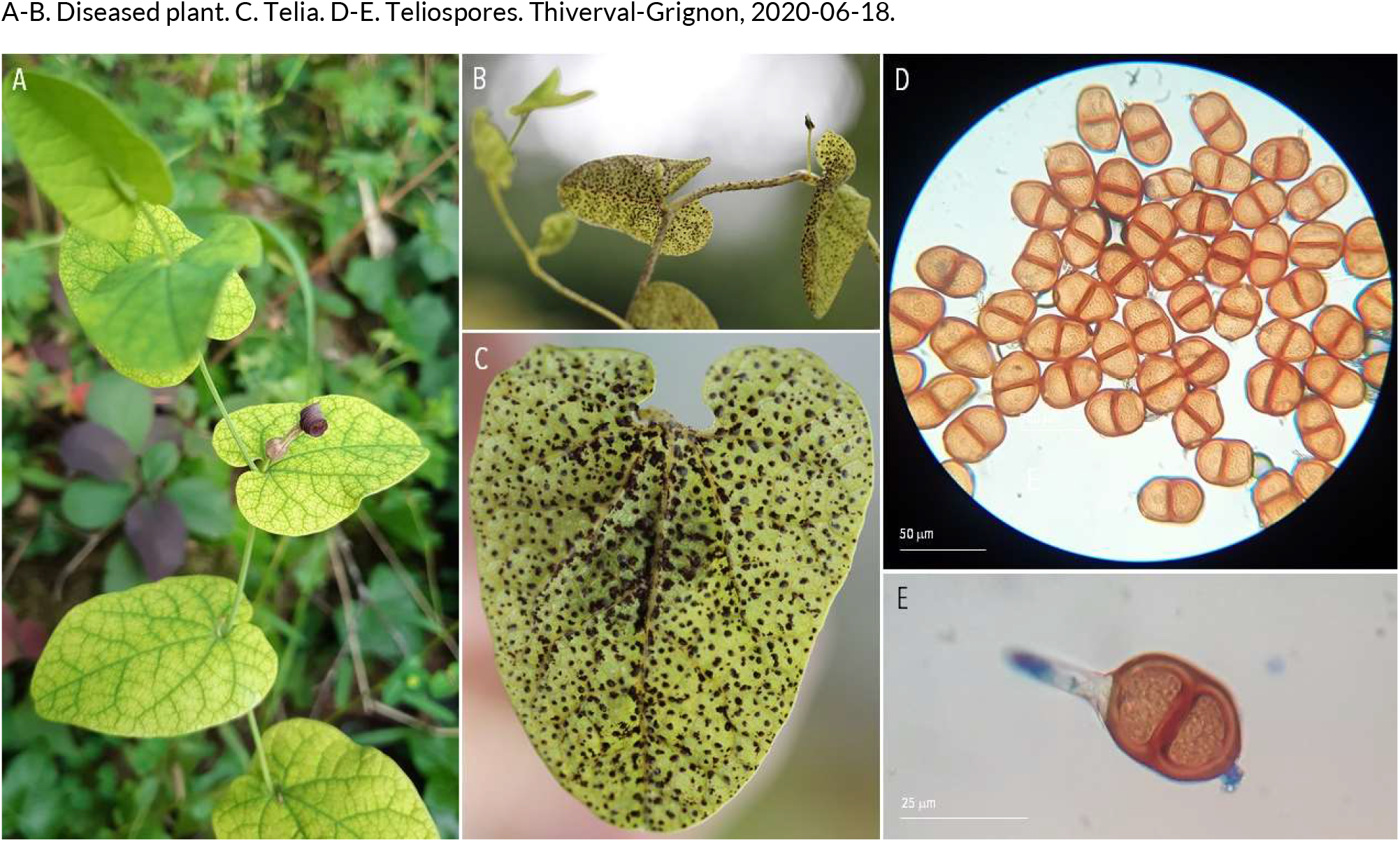
RUSTS | *Puccinia aristolochiae* ex *Aristolochia rotunda* (snakeroot)

**37.**
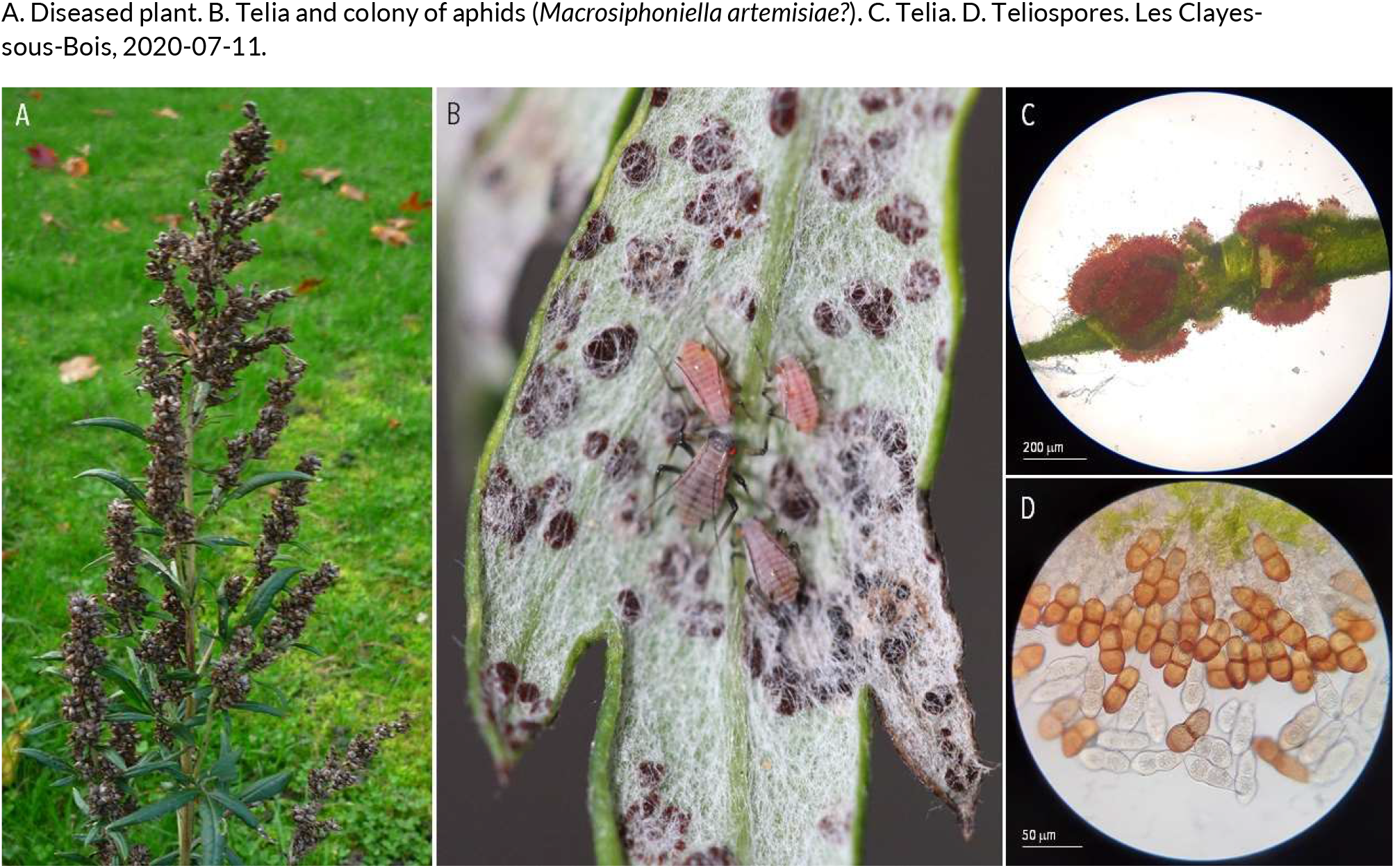
RUSTS | *Puccinia artemisiella* (?) ex Artemisia vulgaris (common mugwort)

**38.**
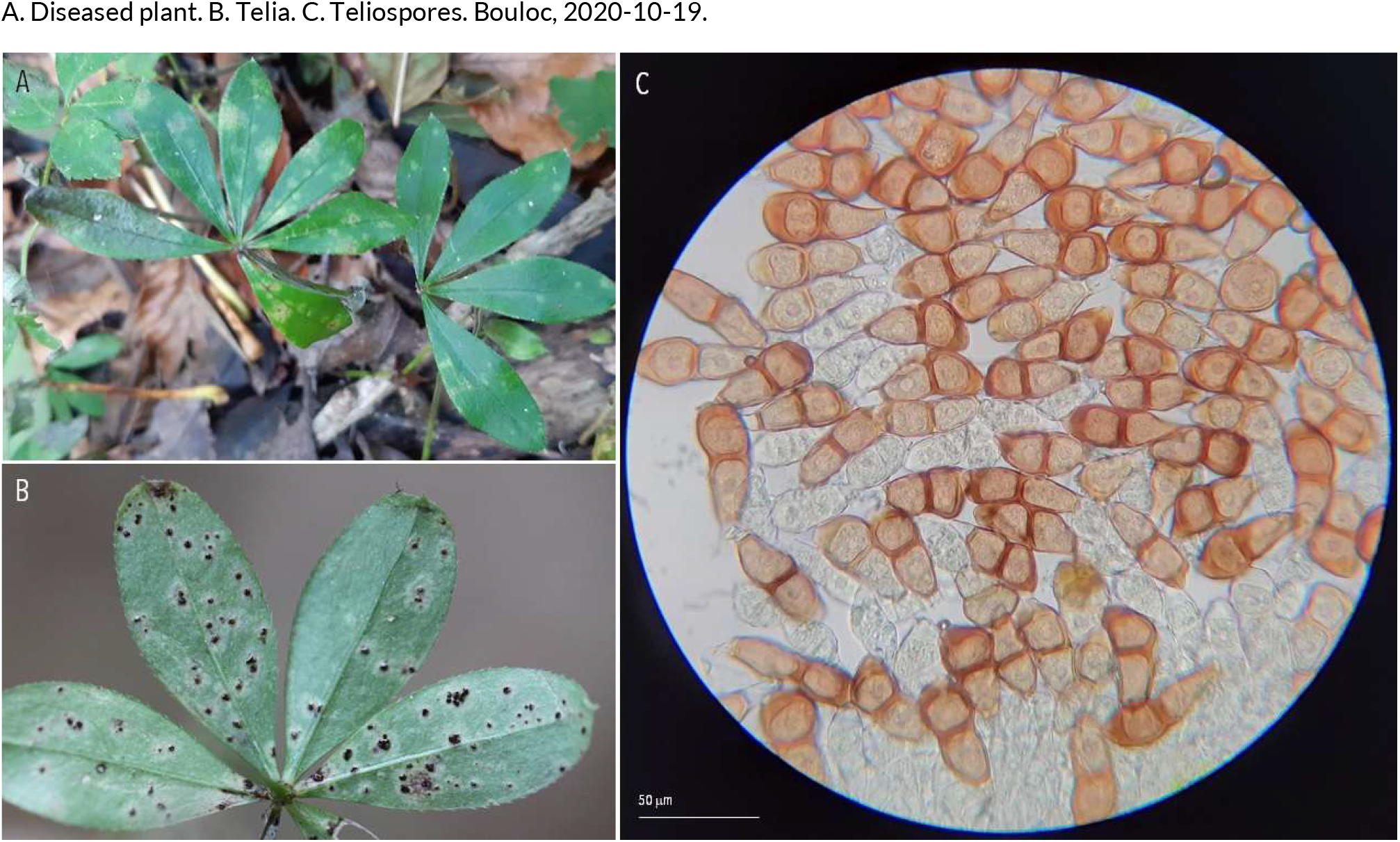
RUSTS | *Puccinia asperulae-odoratae* ex Galium odoratum (sweet woodruff)

**39.**
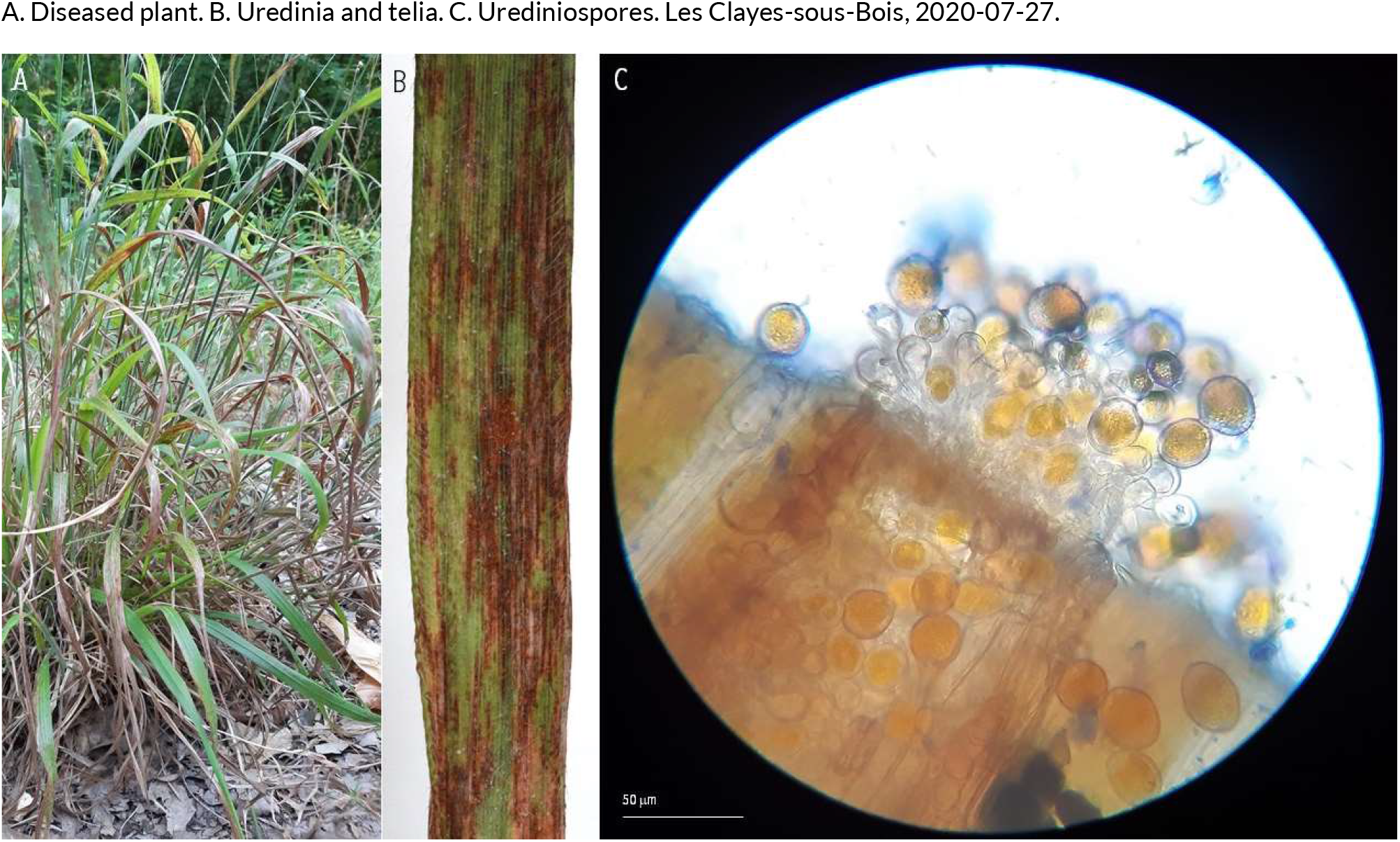
RUSTS | *Puccinia brachypodii* ex Brachypodium sylvaticum (slender false brome)

**40.**
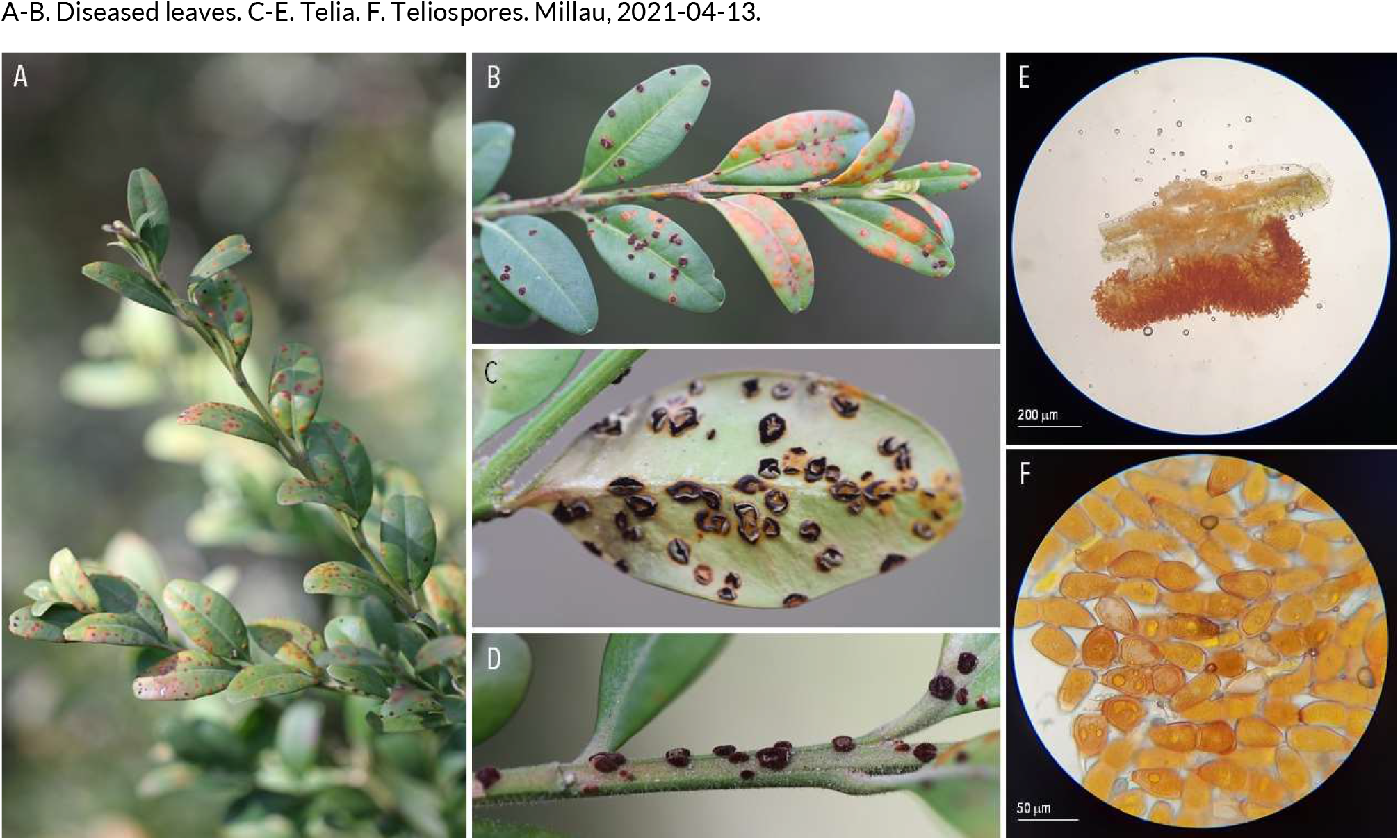
RUSTS | *Puccinia buxi* ex *Buxus sempervirens* (common boxwood)

**41.**
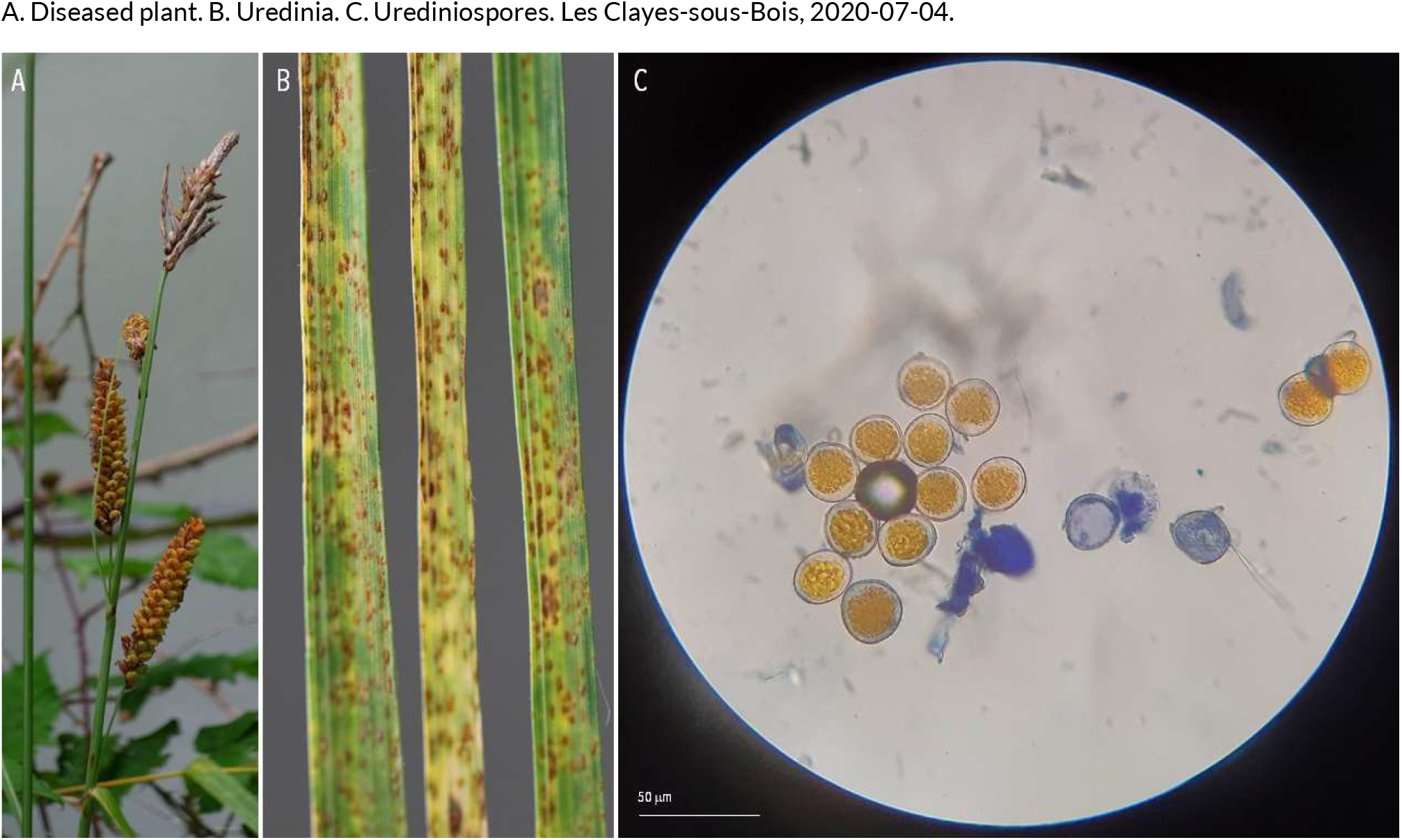
RUSTS | *Puccinia caricina* var*. ribesii-diversicoloris* (??) ex Carex flacca (blue sedge)

**42.**
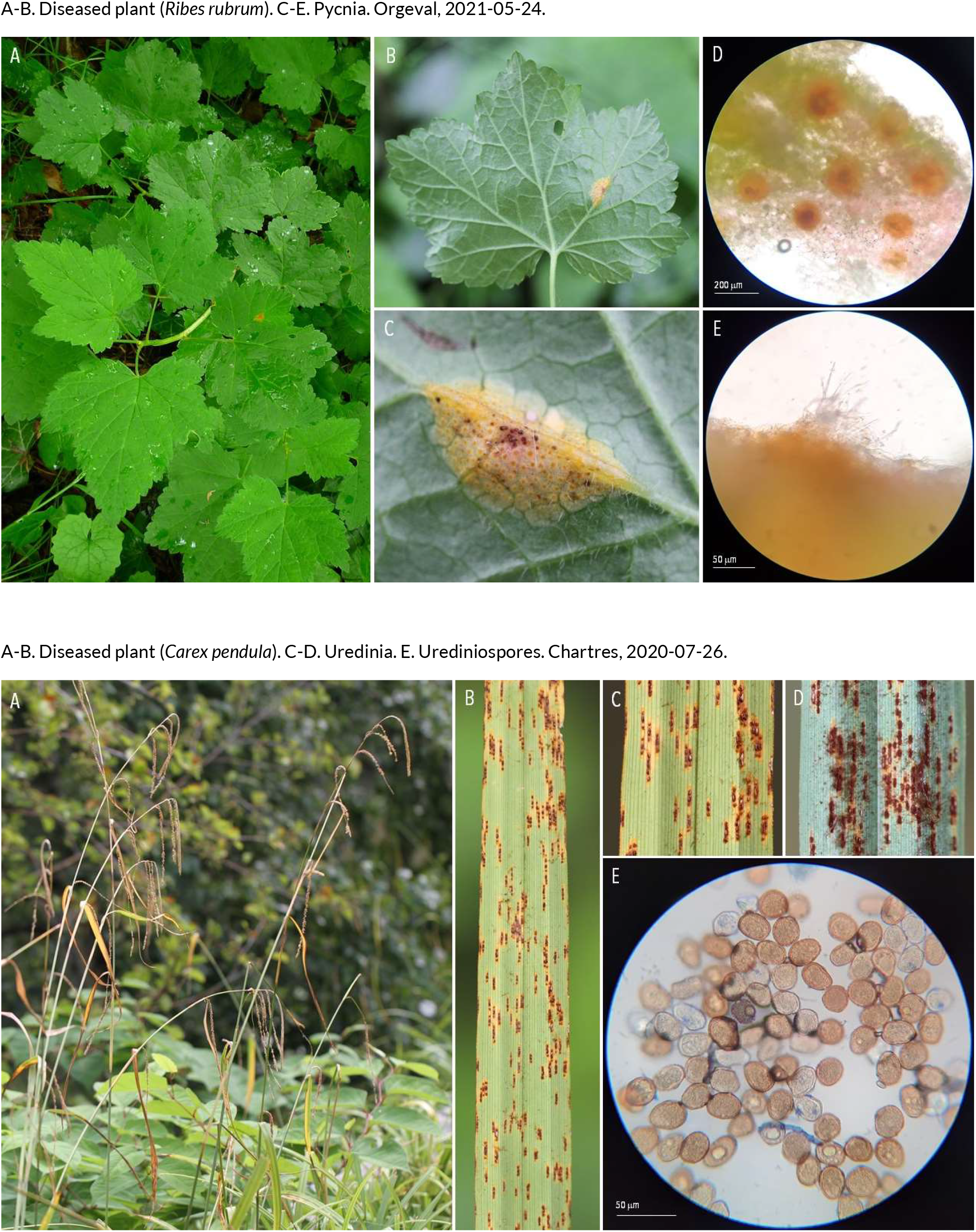
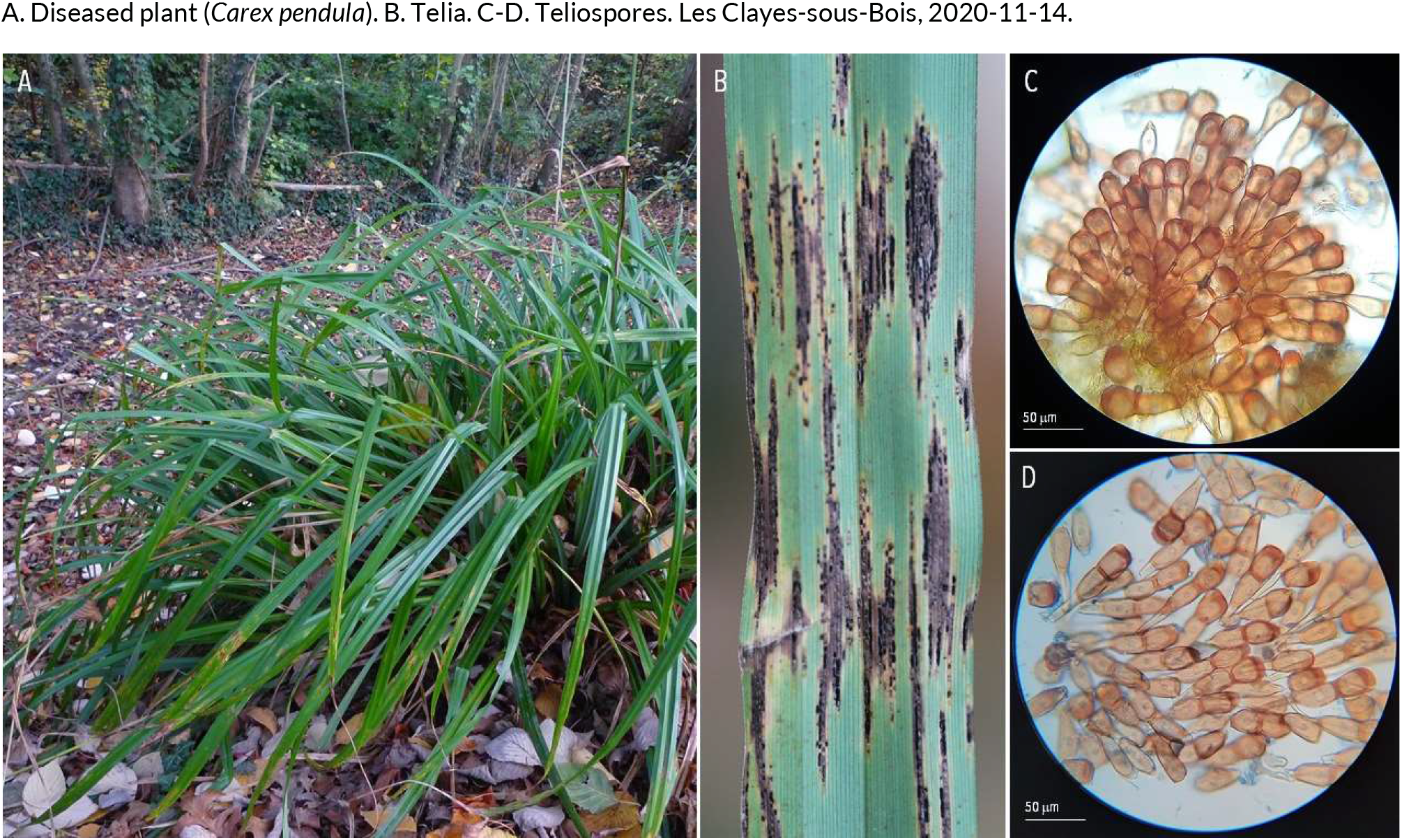
RUSTS | *Puccinia caricina* var. ribesii-pendula (?) *ex Ribes rubrum* (redcurrant) and *Carex pendula* (pendulous sedge)

**43.**
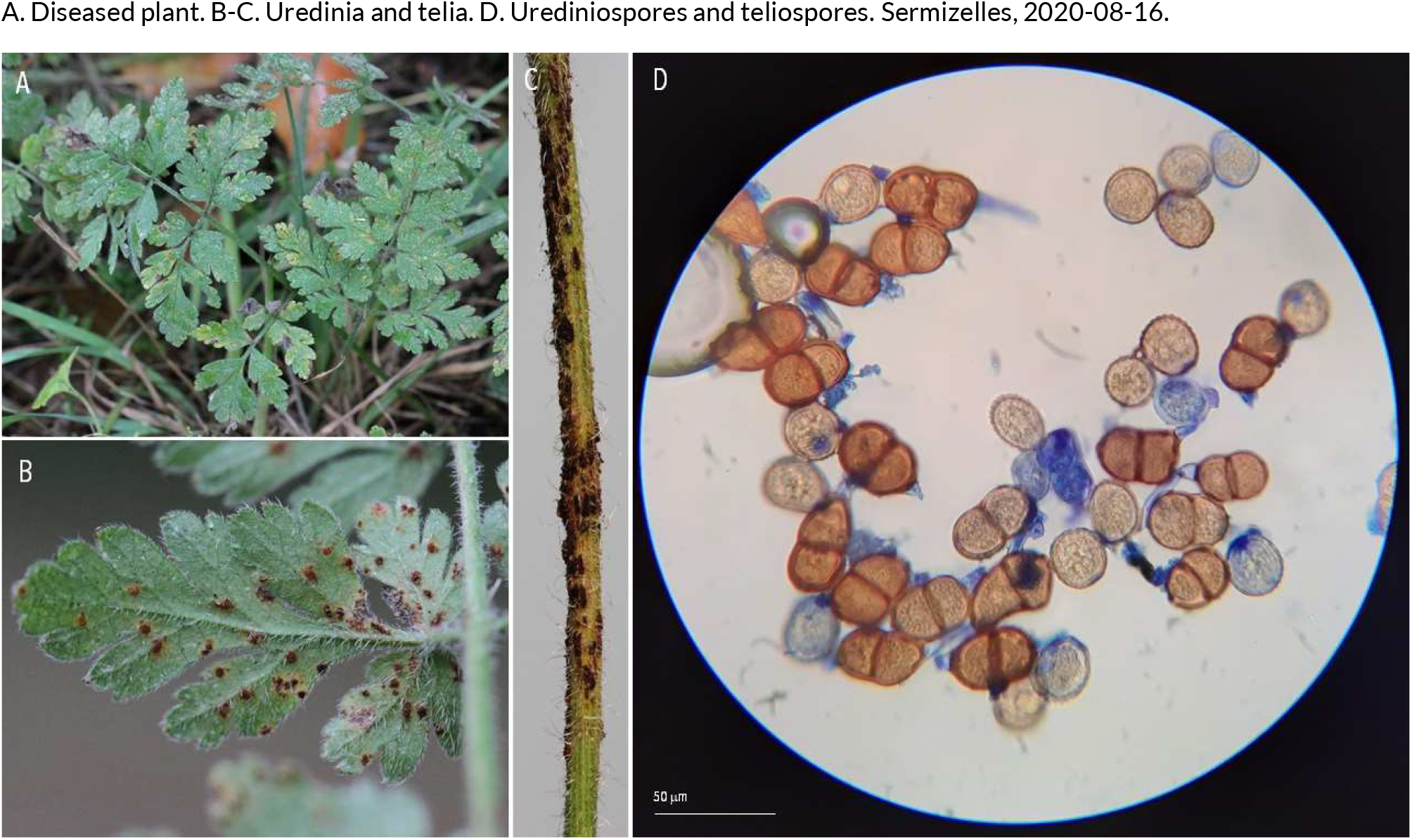
RUSTS | *Puccinia chaerophyllum* ex *Chaerophyllum temulum* (rough chervil)

**44.**
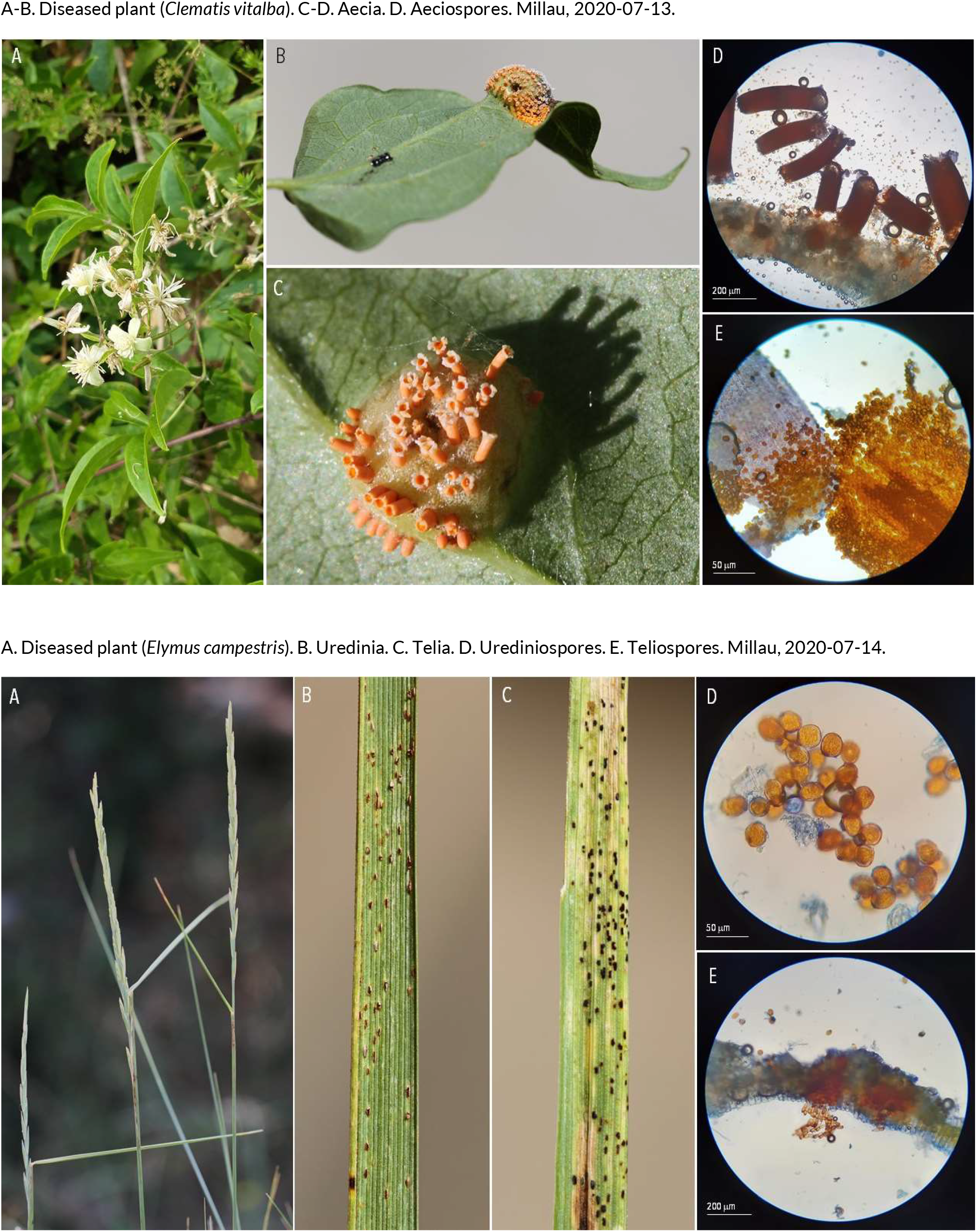
RUSTS | *Puccinia clematidis-secalis* or *Puccinia agropyri* (?) ex *Clematis vitalba* (old man’s beard) and *Elymus campestris* (couch grass)

**45.**
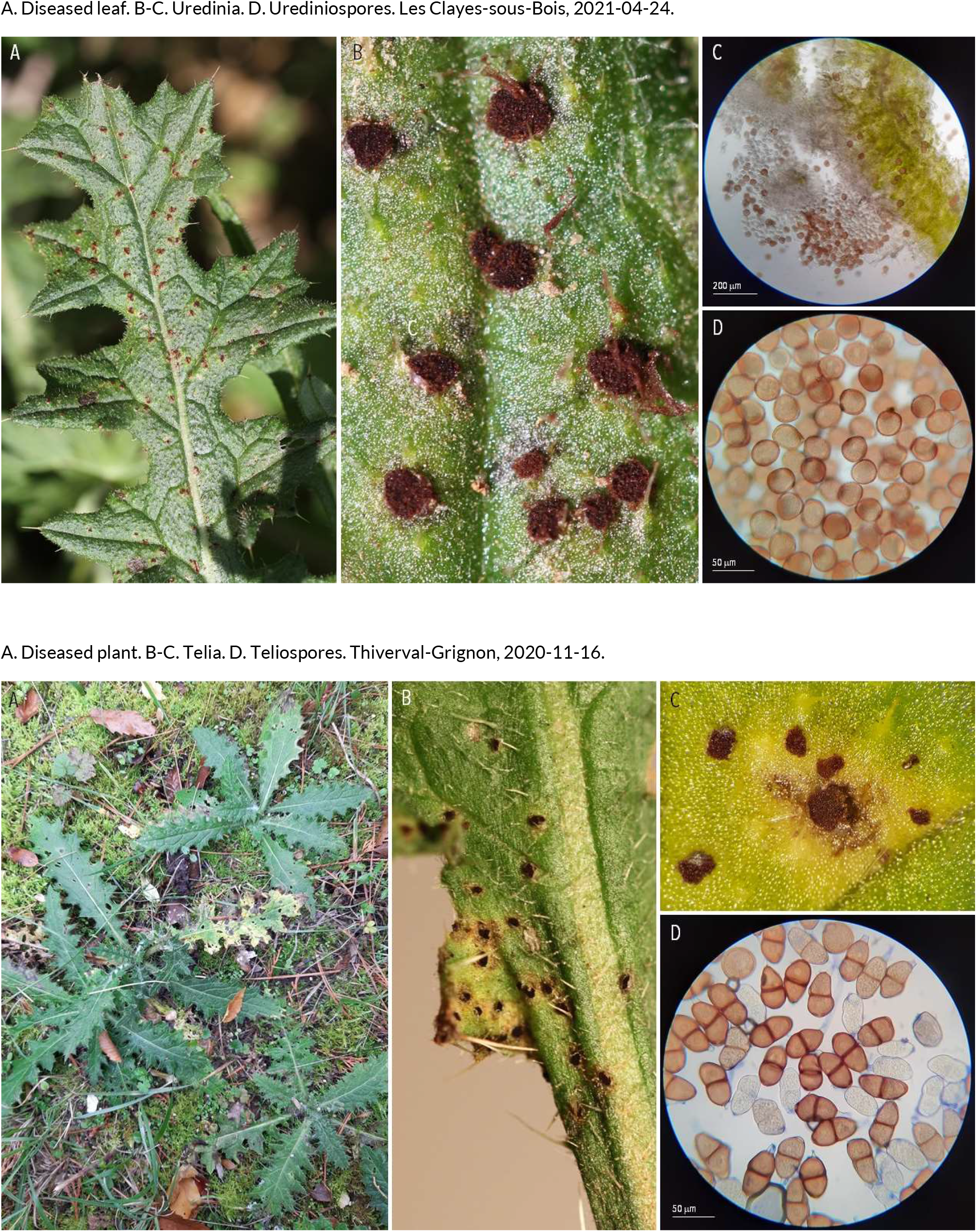
RUSTS | *Puccinia cnici* ex *Cirsium arvense* (spear thistle)

**46.**
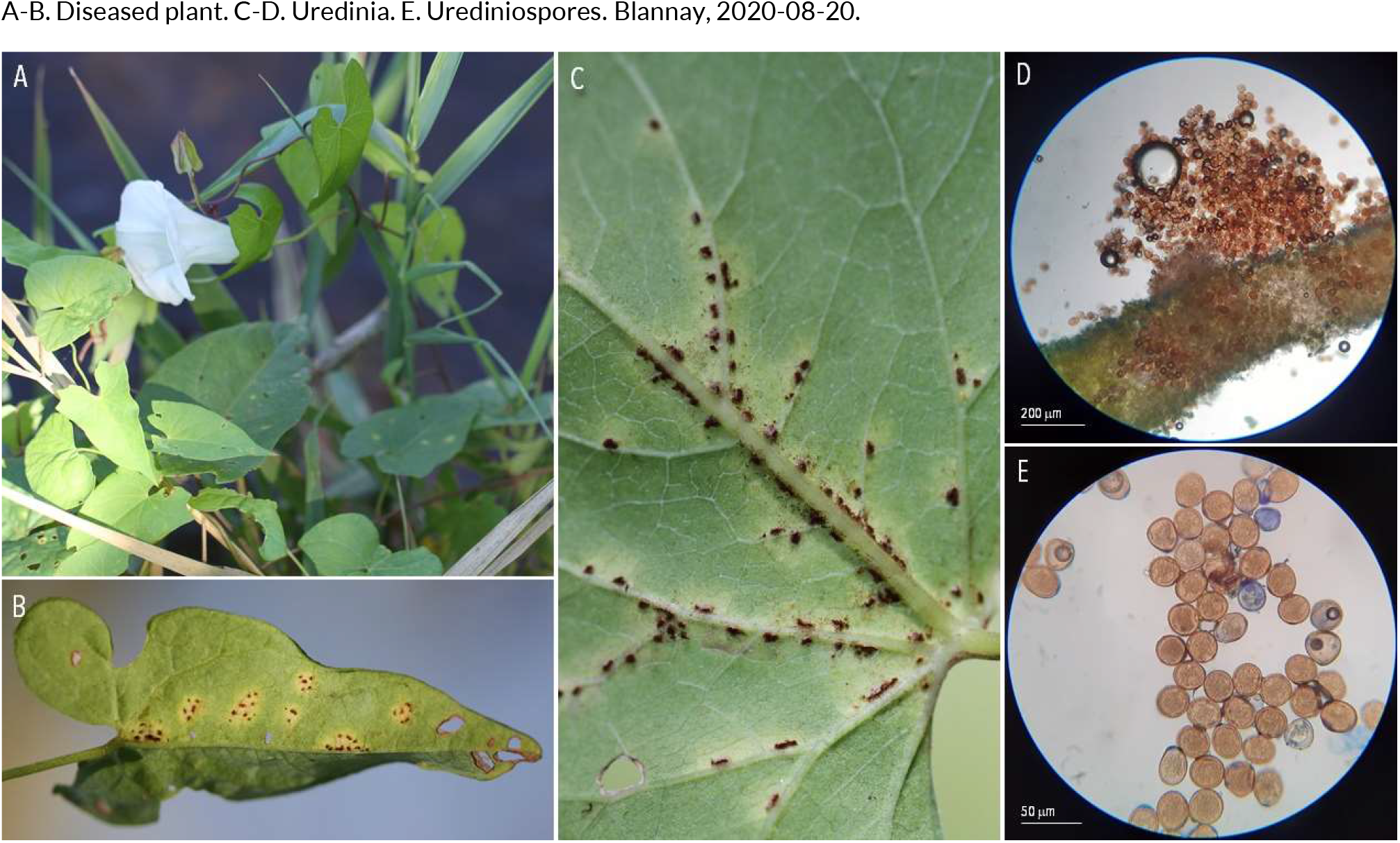
RUSTS | *Puccinia convolvuli* ex *Calystegia sepium* (hedge bindweed)

**47.**
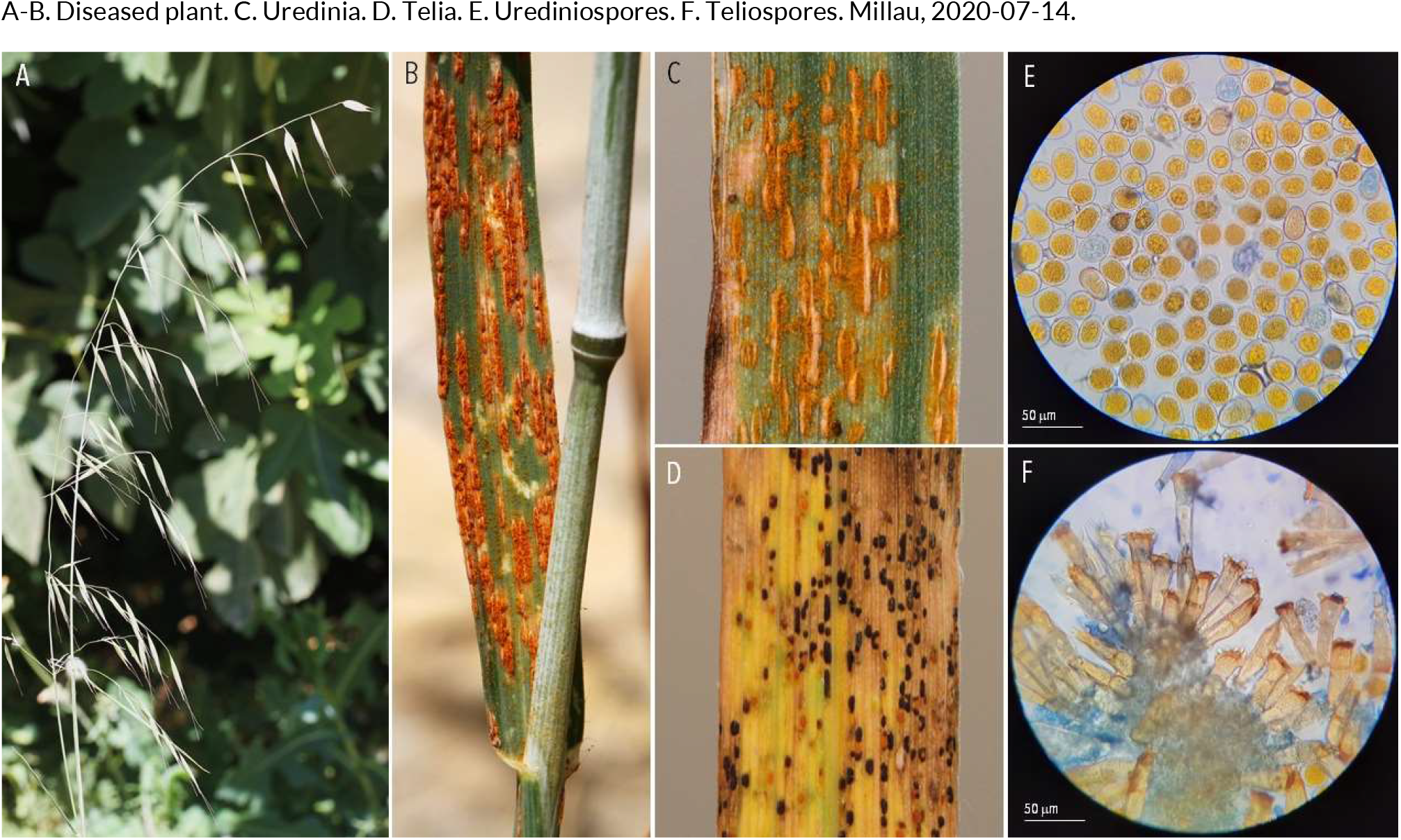
RUSTS | *Puccinia coronata* f. sp. *avenae* ex *Avenae fatua* (common wild oat)

**48.**
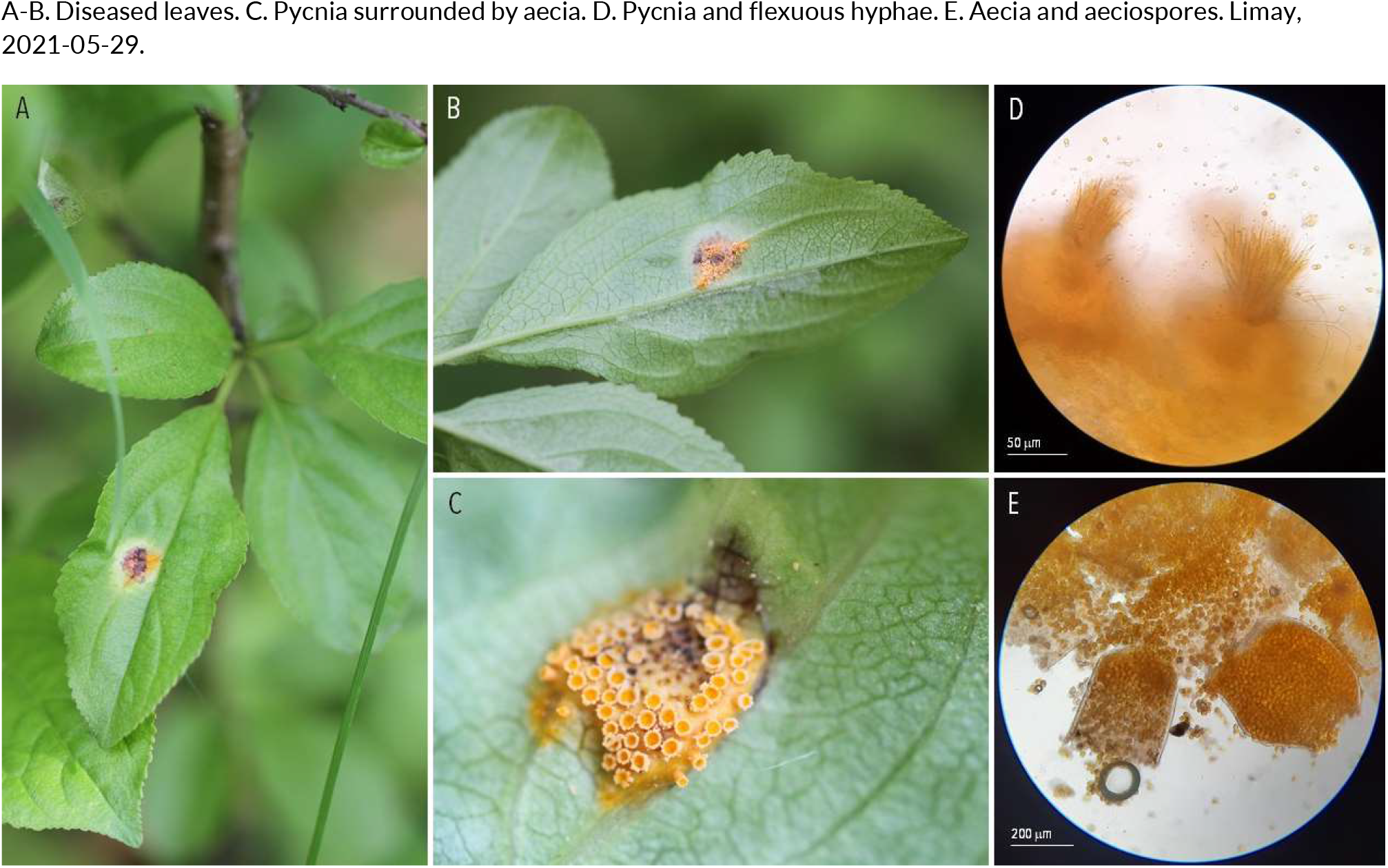
RUSTS | *Puccinia coronata* f. sp*. bromi* ex *Rhamnus cathartica* (common buckthorn)

**49.**
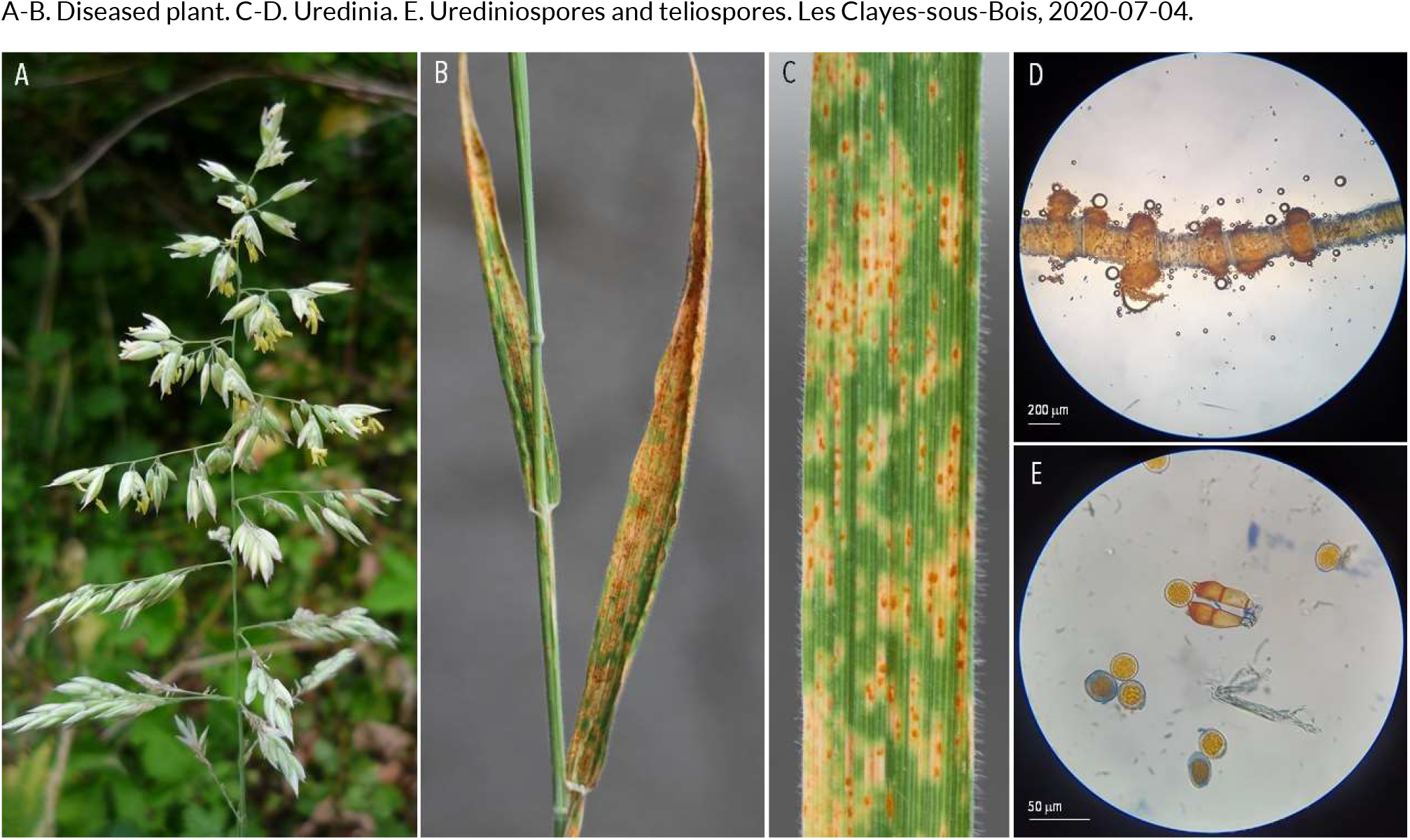
RUSTS | *Puccinia coronata* f. sp. *holci* ex *Holcus lanatus* (common velvet grass)

**50.**
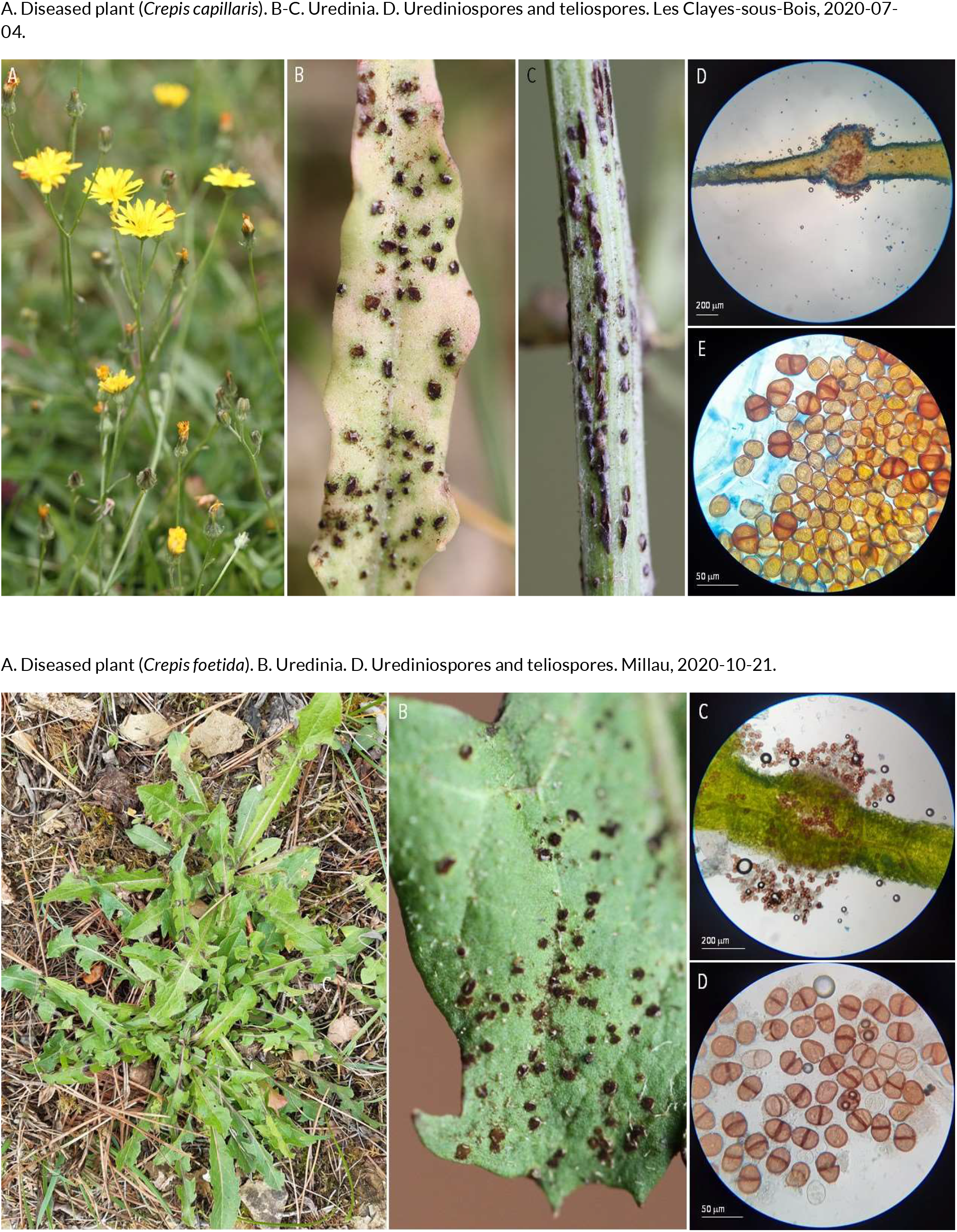
RUSTS | *Puccinia crepidicola* ex *Crepis capillaris* (smooth hawksbeard) and *Crepis foetida* (stinking hawksbeard)

**51.**
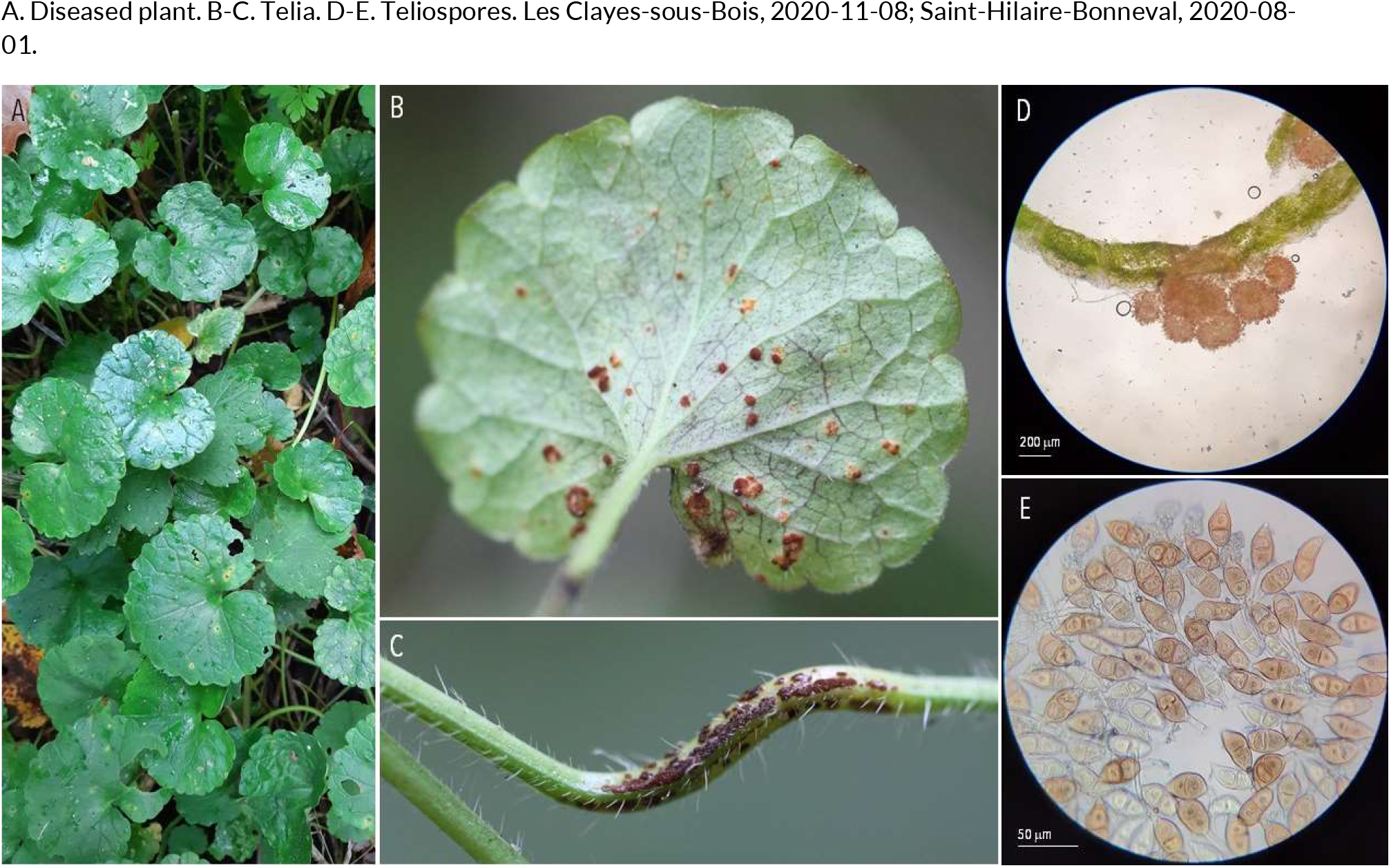
RUSTS | *Puccinia glechomatis* ex *Glechoma hederacea* (ground ivy)

**52.**
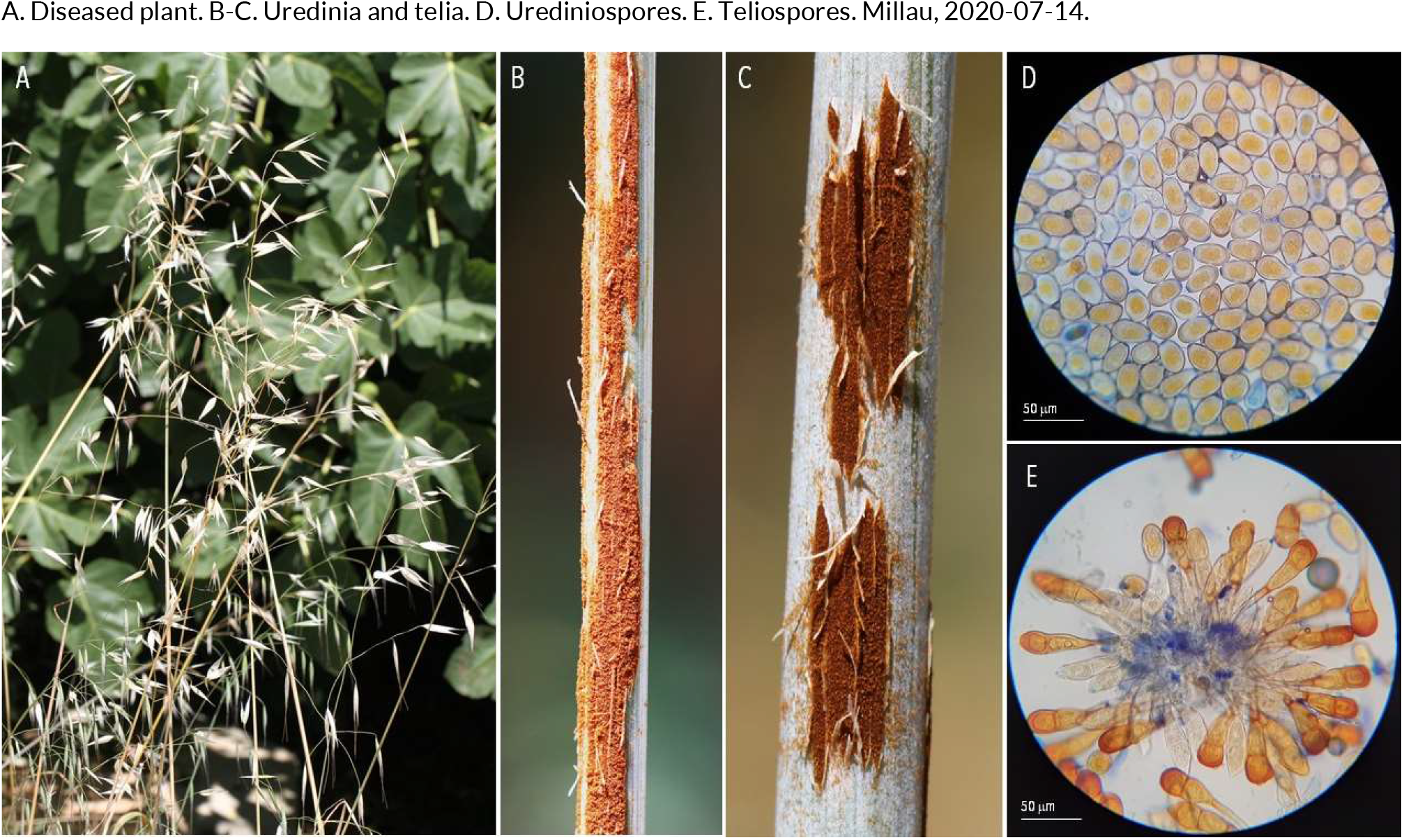
RUSTS | *Puccinia graminis* f. sp. *avenaceum* ex *Avenae fatua* (common wild oat)

**53.**
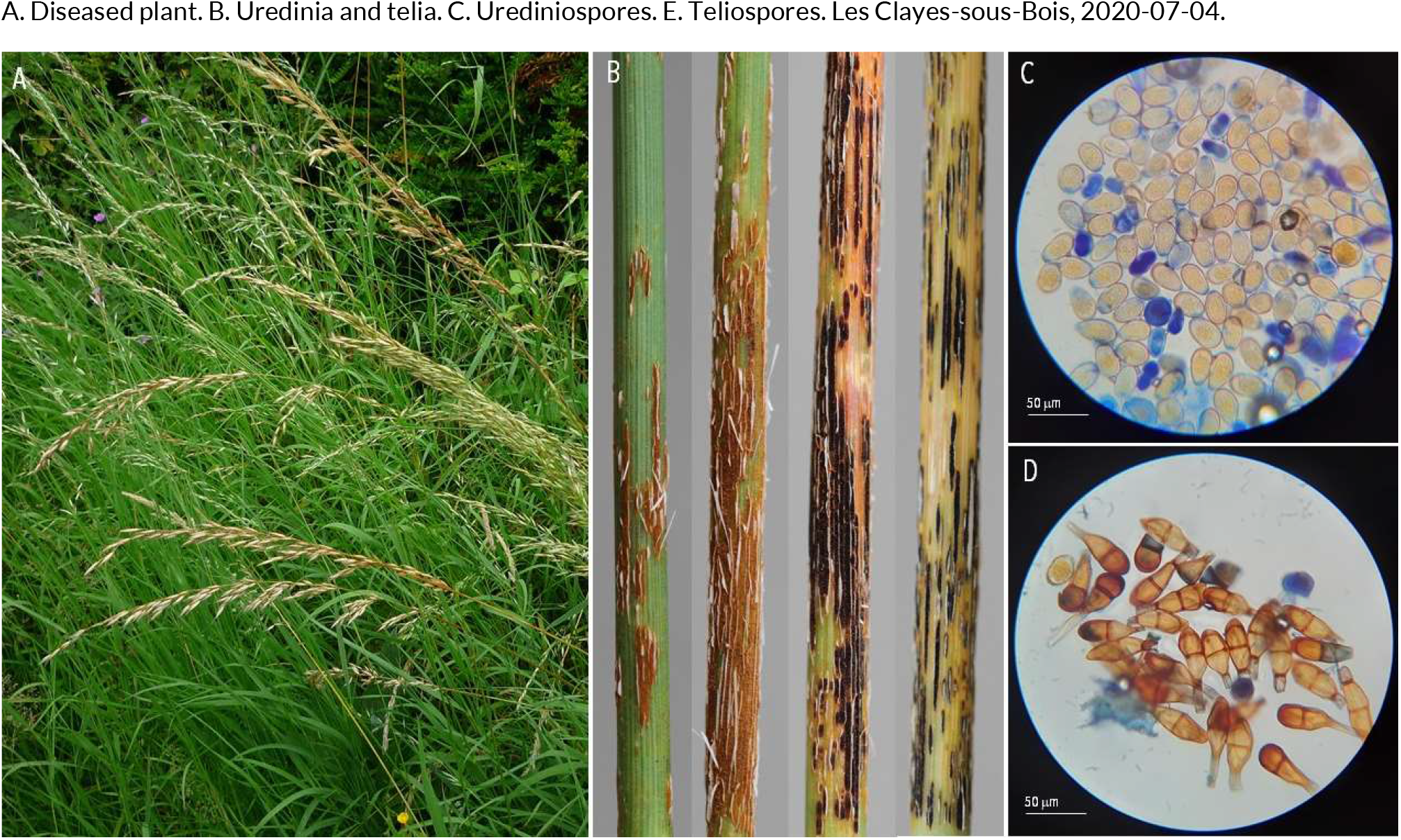
RUSTS | *Puccinia graminis* f. sp. *avenae* (?) ex *Arrhenatherum elatius* (tall oatgrass)

**54.**
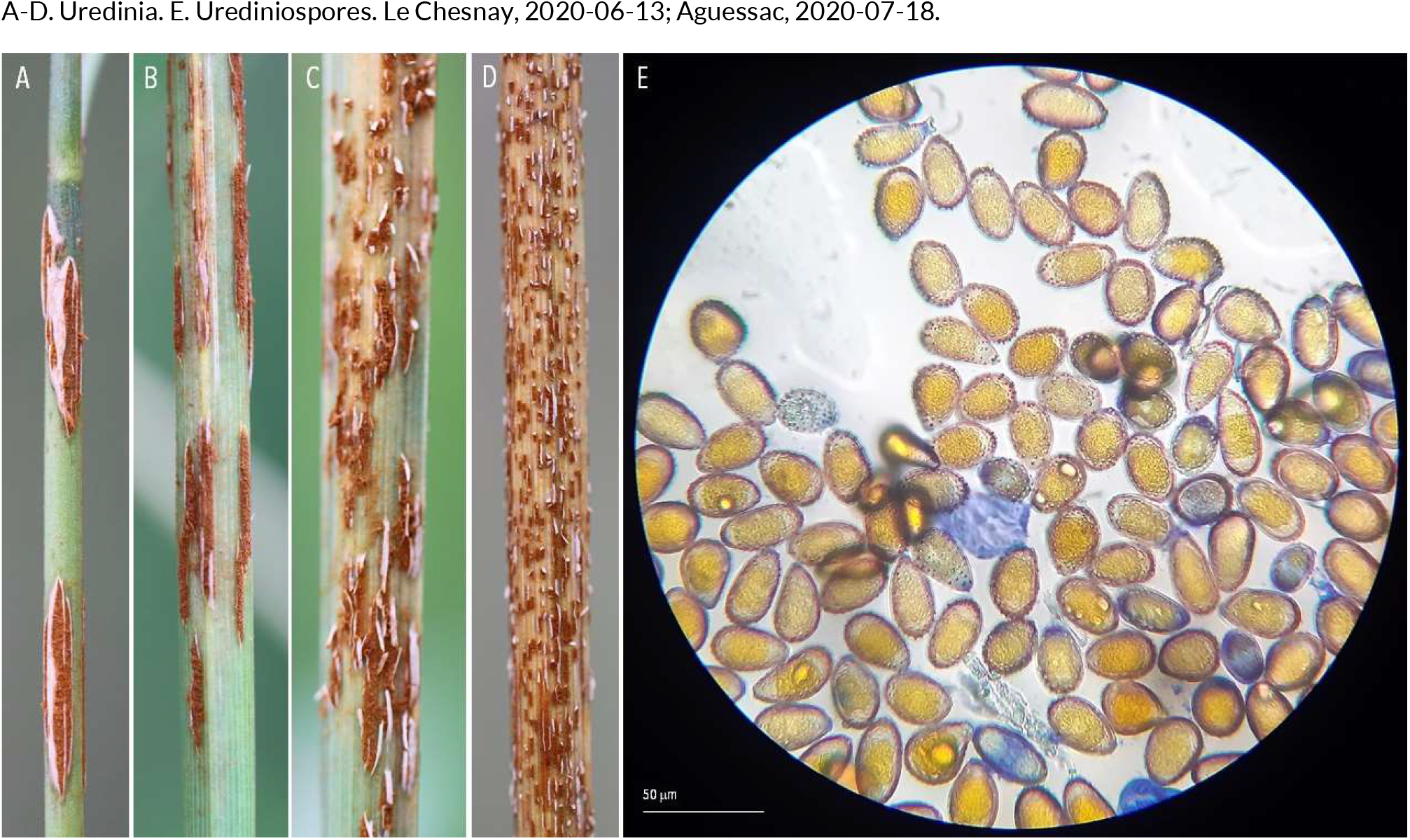
RUSTS | *Puccinia graminis* f. sp. *dactylidis* ex *Dactylis glomerata* (orchard grass)

**55.**
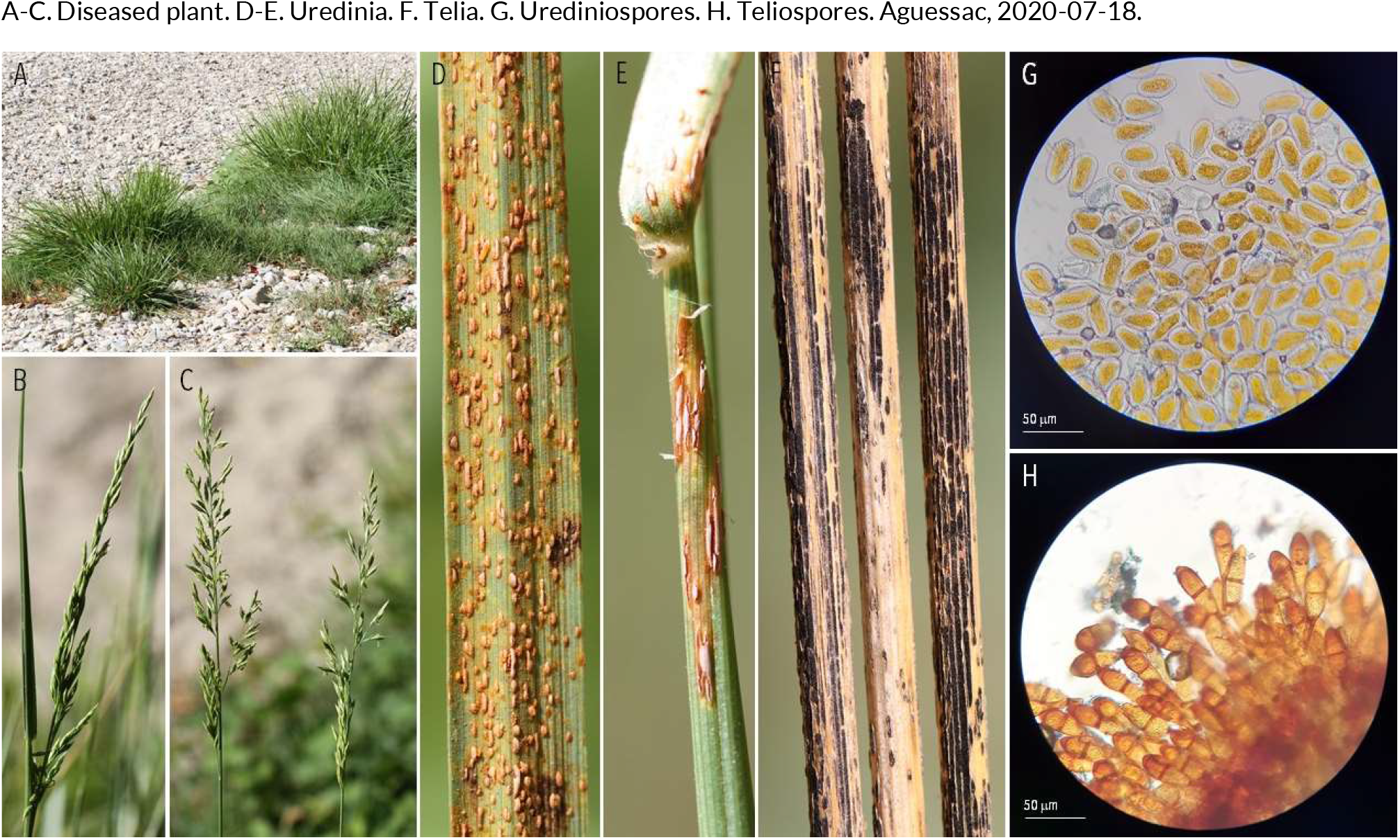
RUSTS | *Puccinia graminis* f. sp. *graminicola* (?) ex *Festuca arundinacea* (tall fescue)

**56.**
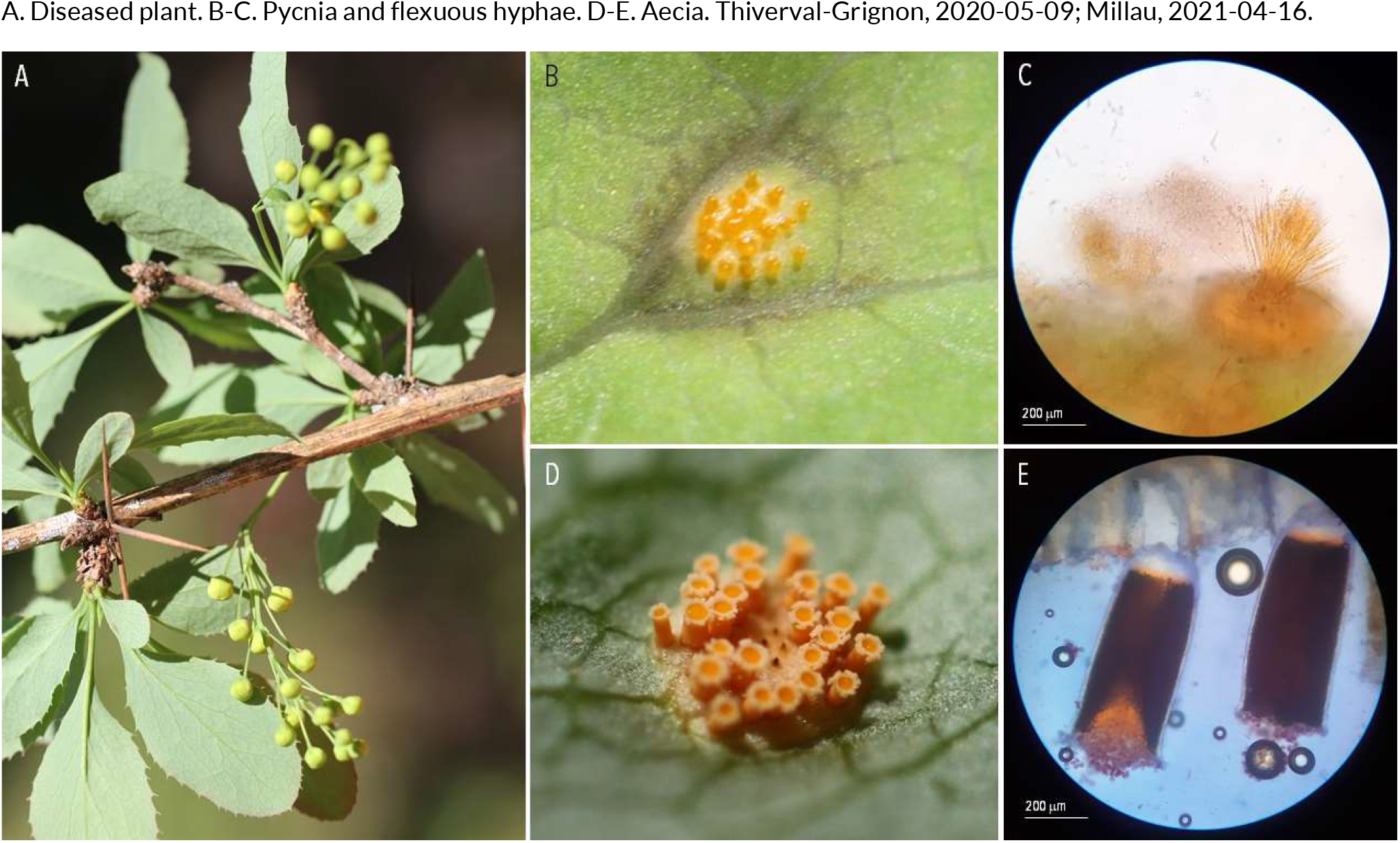
RUSTS | *Puccinia graminis* sp. *secalis* (?) ex *Berberis vulgaris* (common barberry)

**57.**
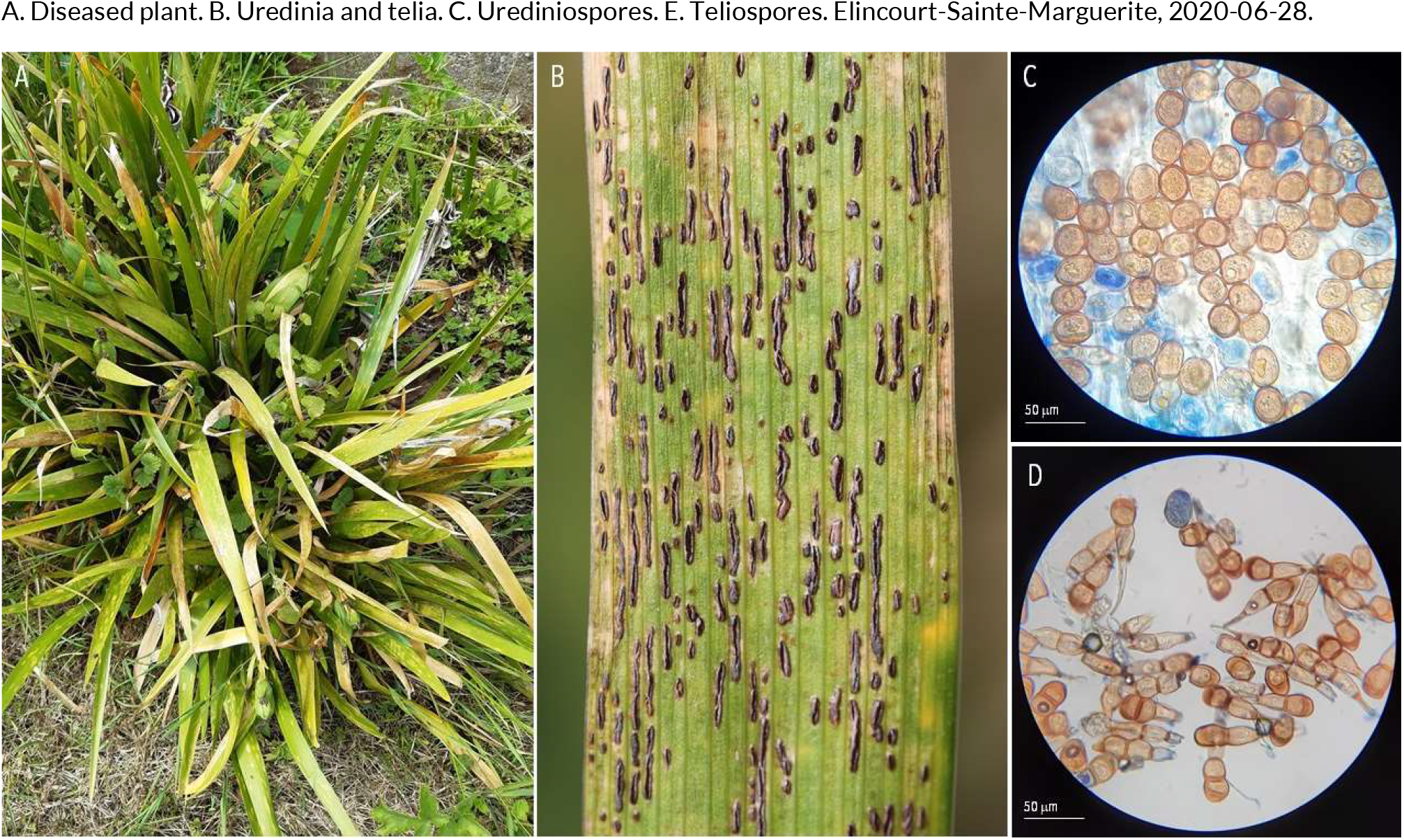
RUSTS | *Puccinia iridis* ex *Iris foetidissima* (stinking iris)

**58.**
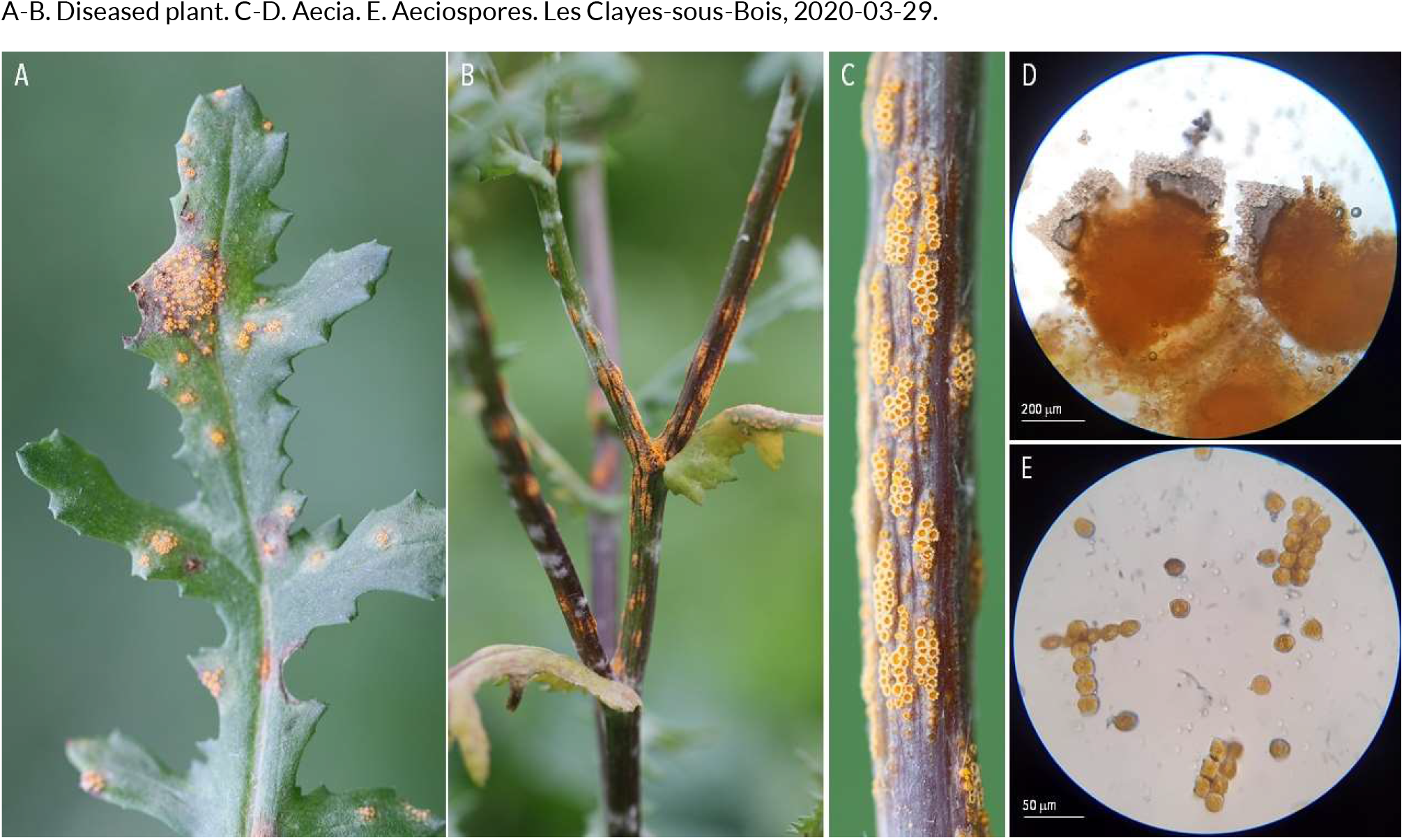
RUSTS | *Puccinia lagenophorae* ex *Senecio vulgaris* (common groundsel)

**59.**
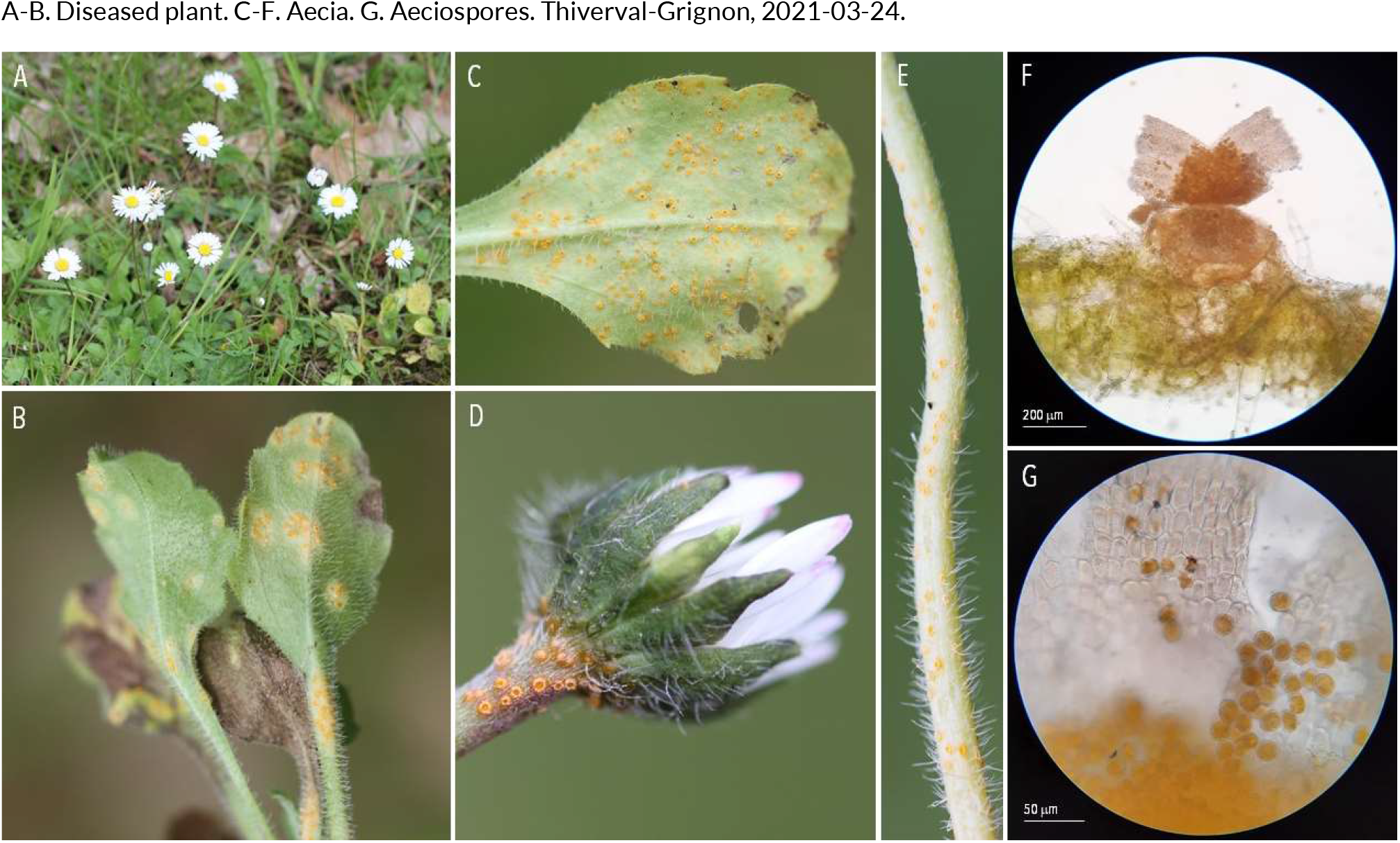
RUSTS | *Puccinia lagenophorae* or *Puccinia distincta* (?) ex *Bellis perennis* (English daisy)

**60.**
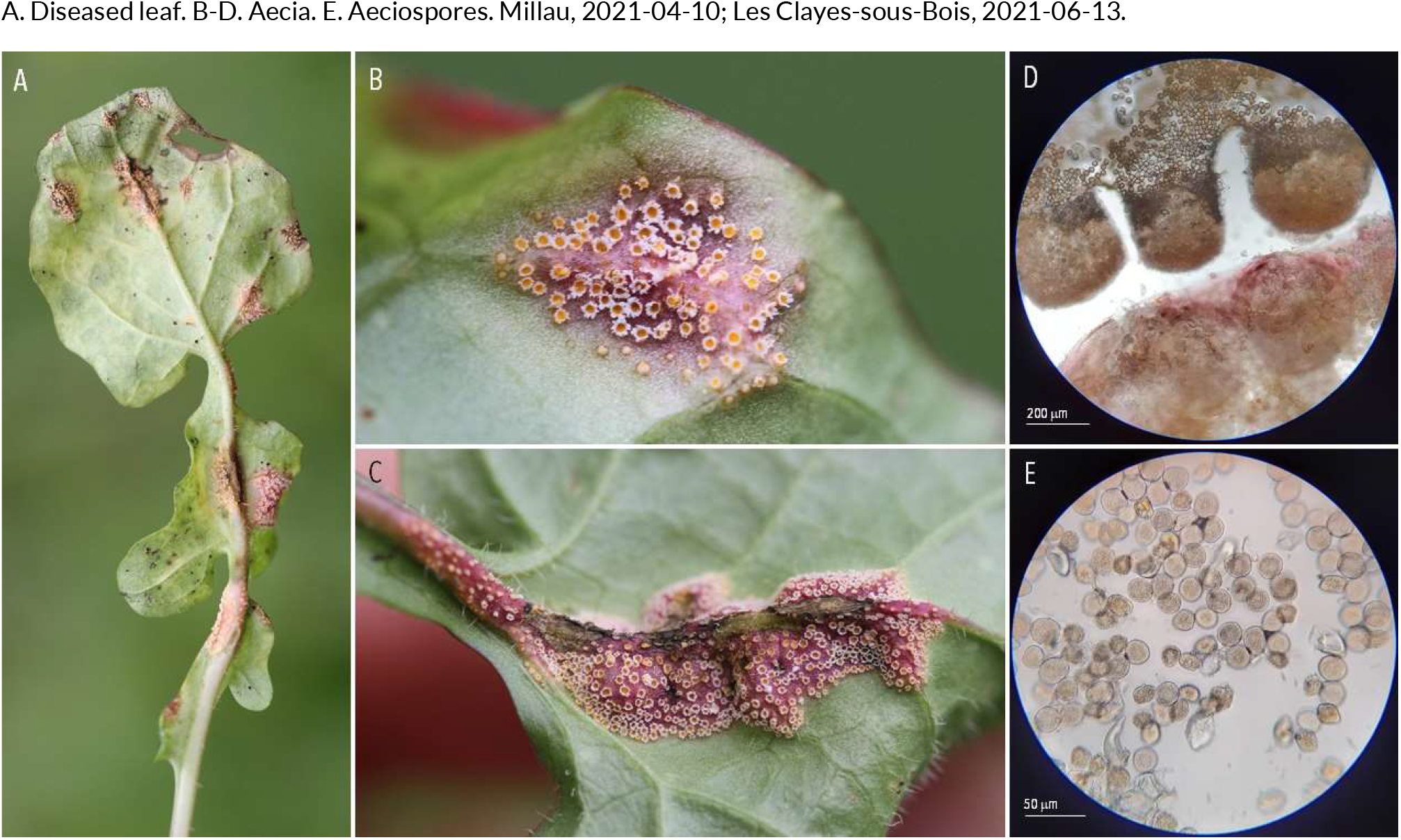
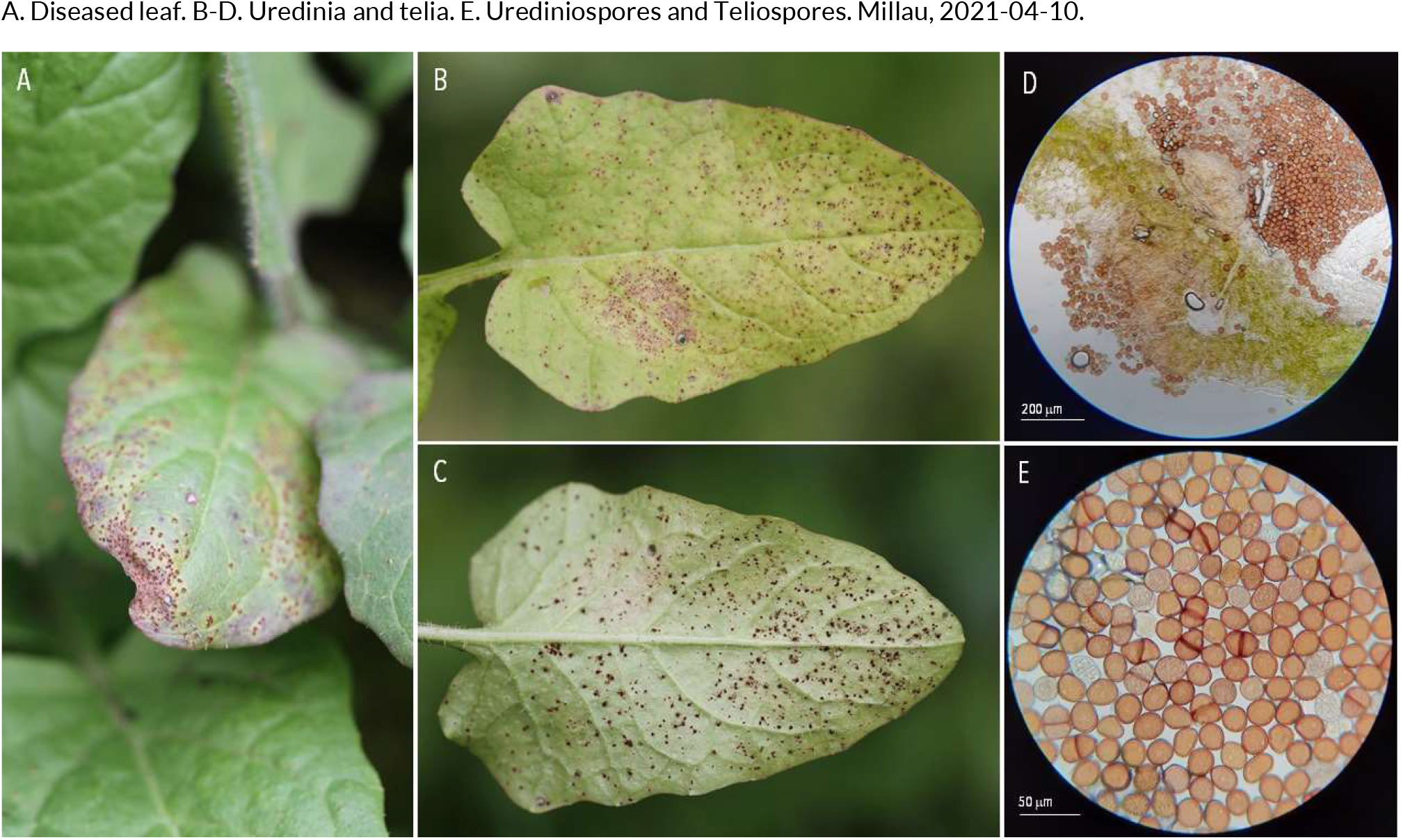
RUSTS | *Puccinia lapsanae* ex *Lapasana communis* (nipplewort)

**61.**
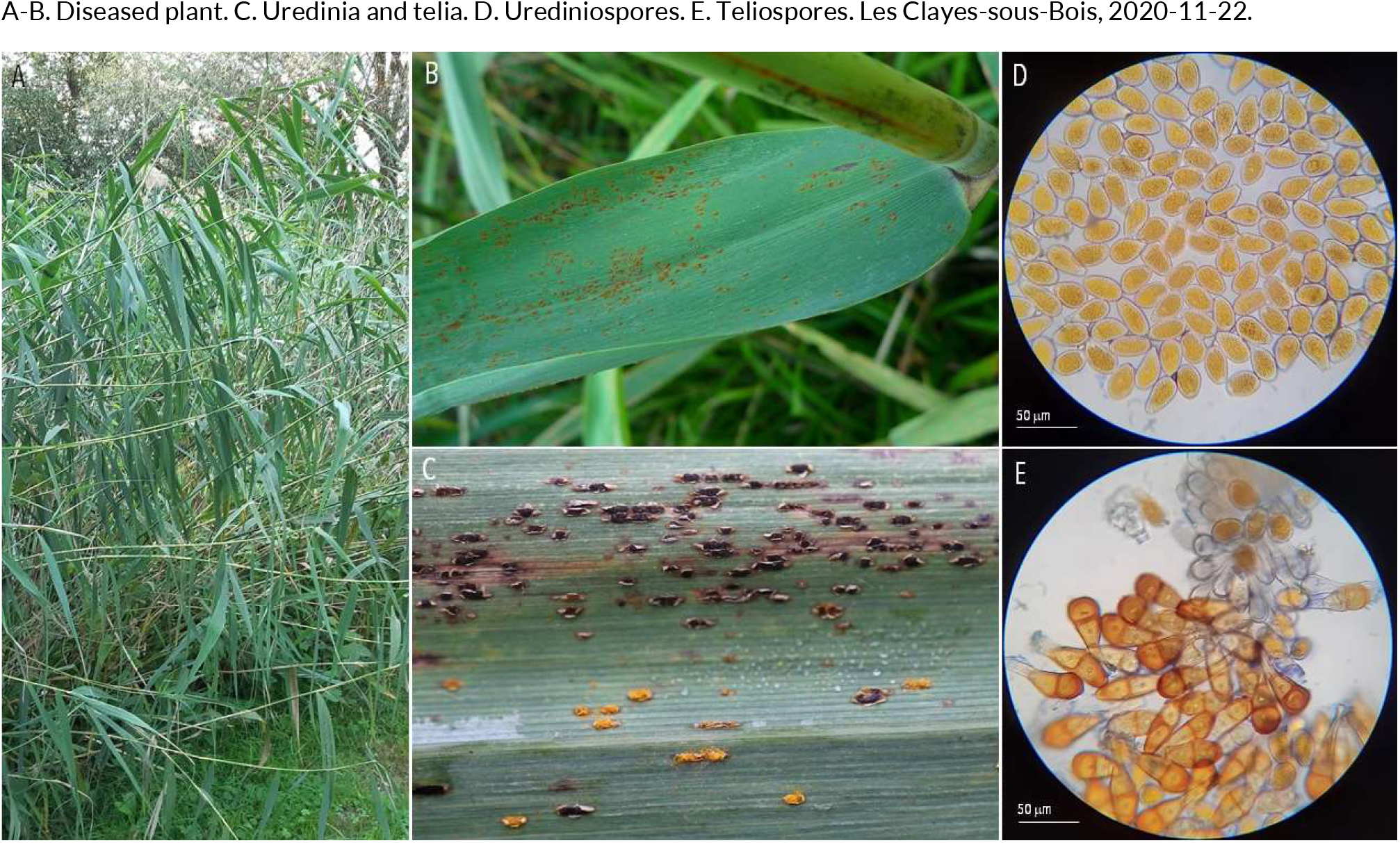
RUSTS | *Puccinia magnusiana* ex *Phragmitis australis* (common reed)

**62.**
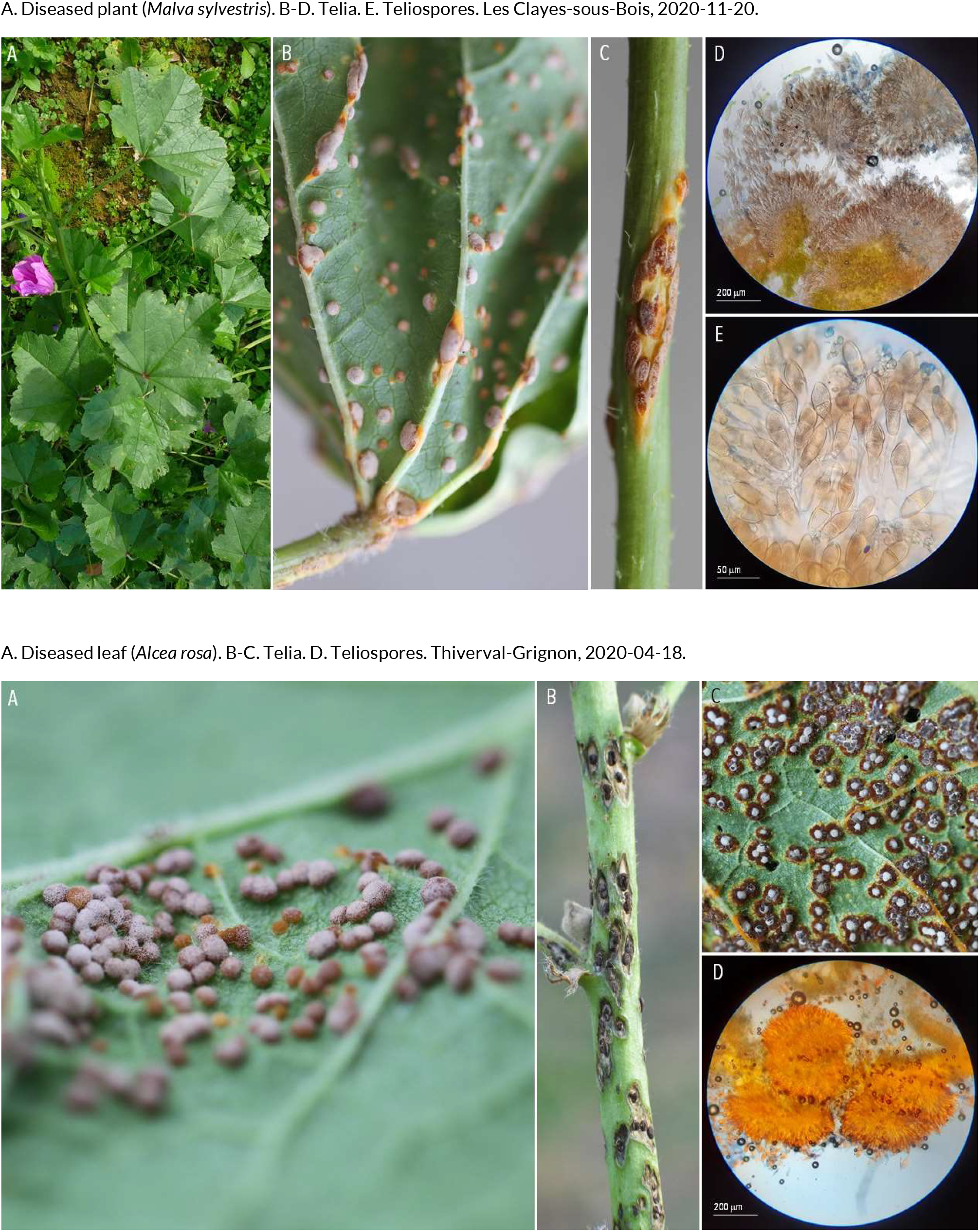
RUSTS | *Puccinia malvacearum* ex *Malva sylvestris* (common mallow) and *Alcea rosa* (common hollyhock)

**63.**
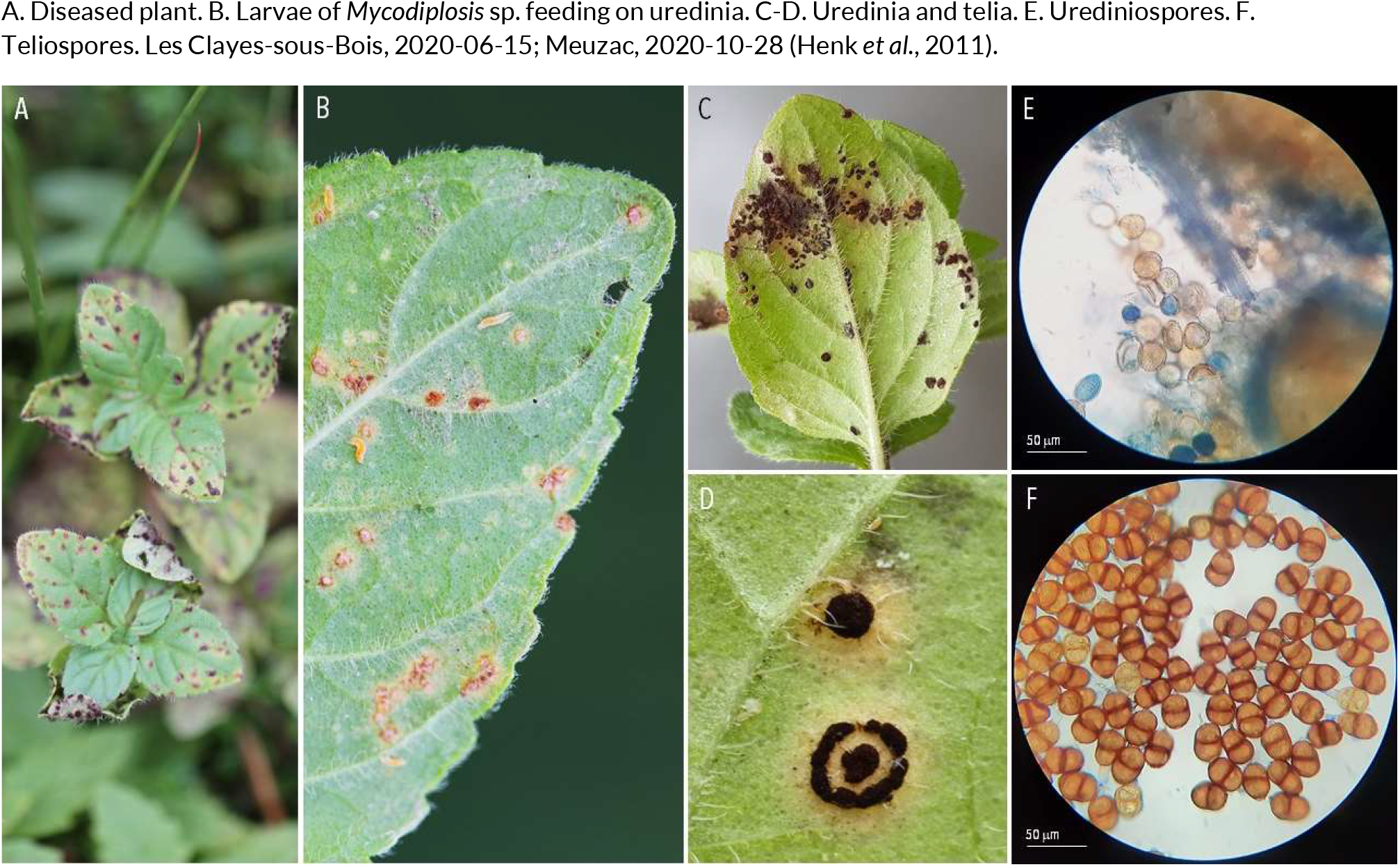
RUSTS | *Puccinia menthae* ex *Mentha aquatica* (water mind)

**64.**
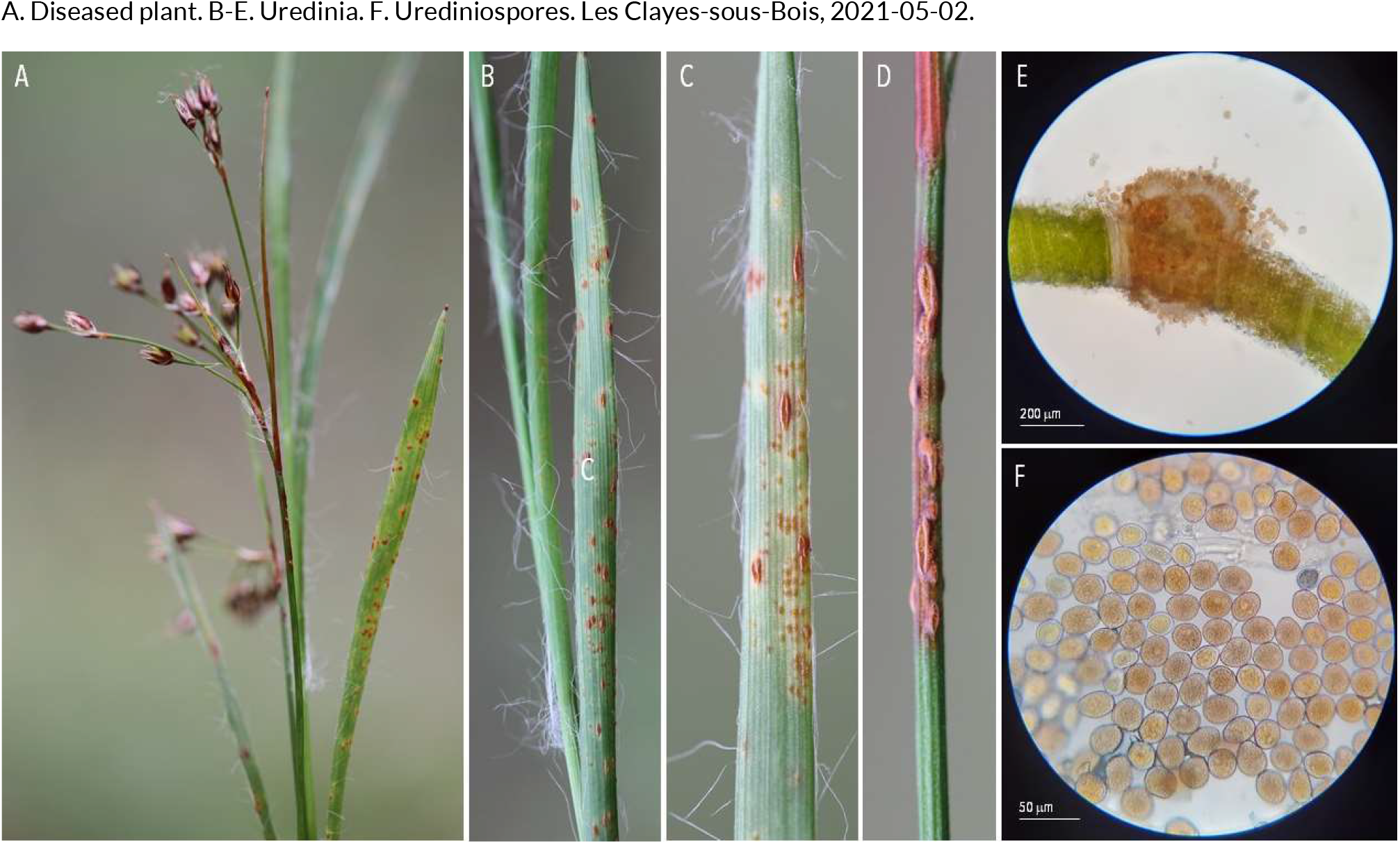
RUSTS | *Puccinia obscura* (?) ex *Luzula forsteri* (southern wood-rush)

**65.**
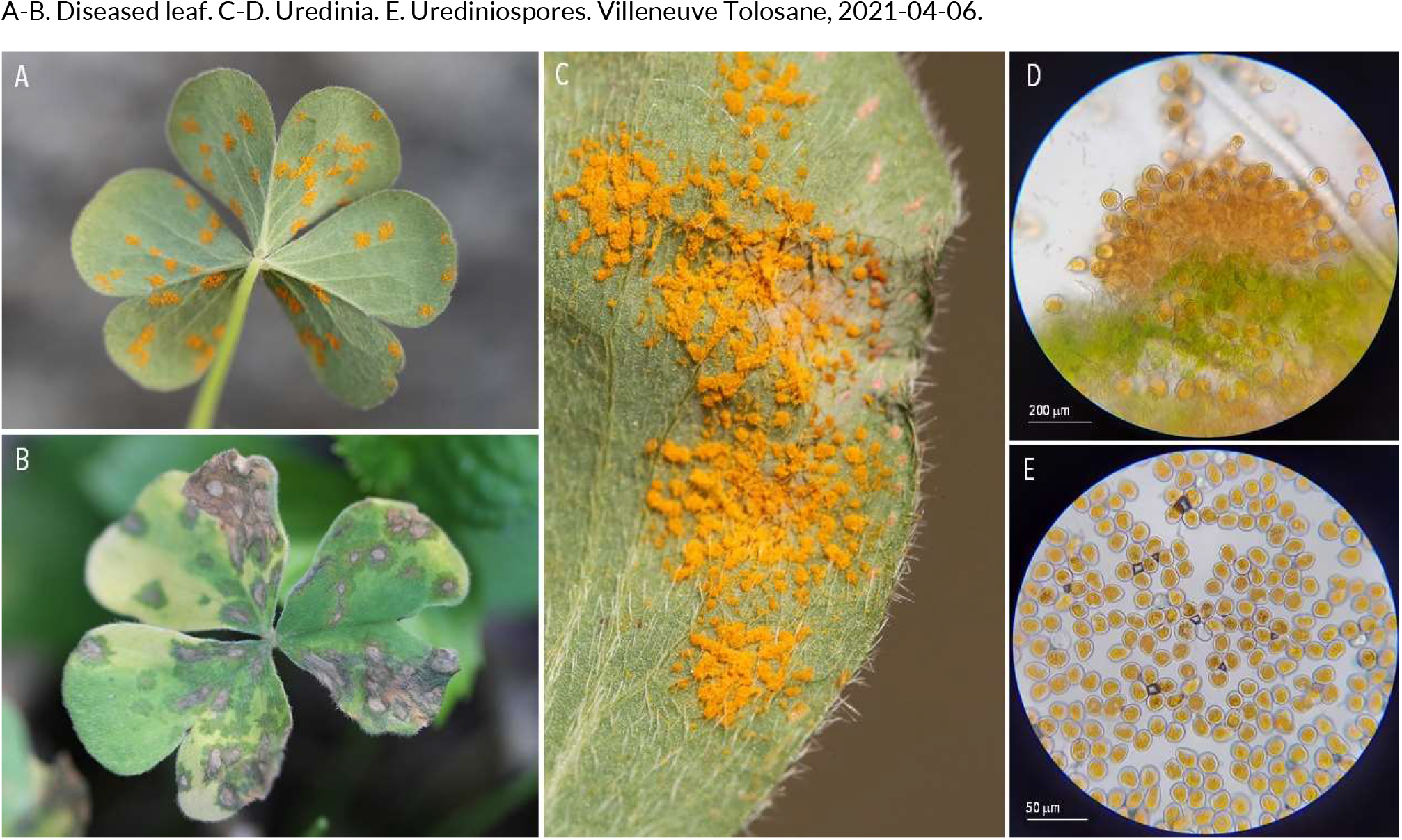
RUSTS | *Puccinia oxalidis* ex *Oxalis articulata* (pink wood sorrel)

**66.**
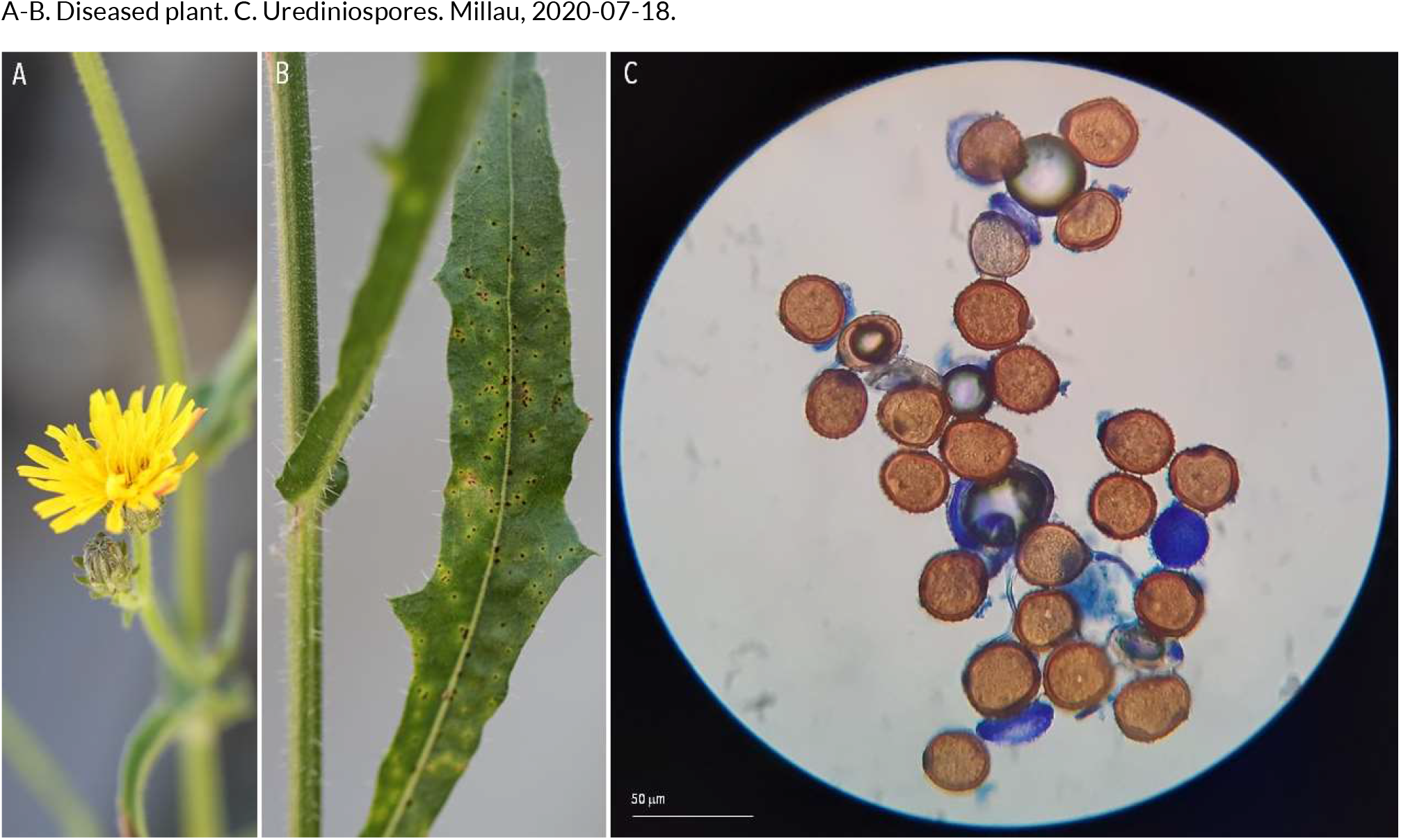
RUSTS | *Puccinia picridis* ex *Picris hieracioides* (water mind)

**67.**
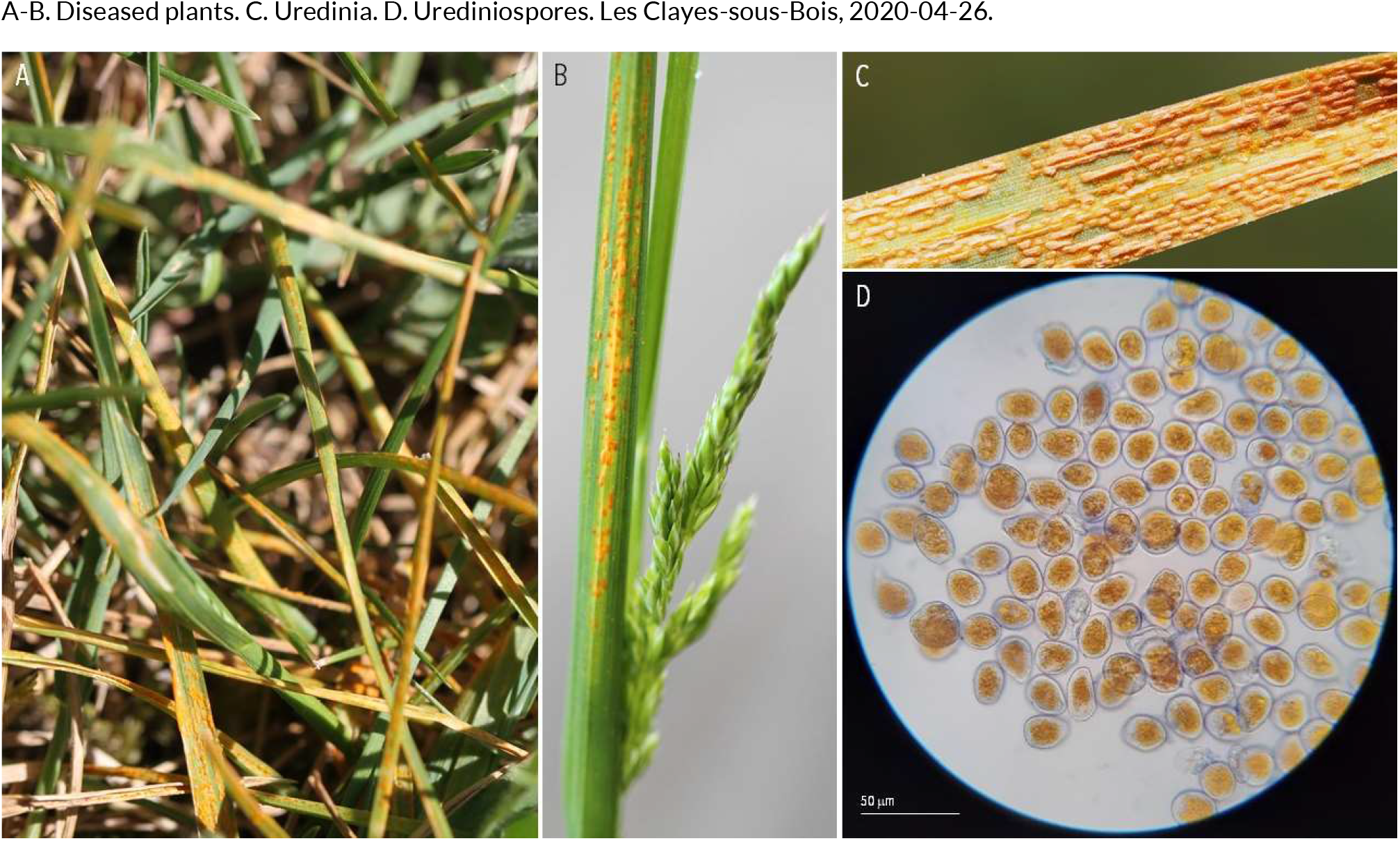
RUSTS | *Puccinia poae-nemoralis* or *Puccinia pseudostriiformis* (?) ex *Poa annua* (annual meadow grass)

**68.**
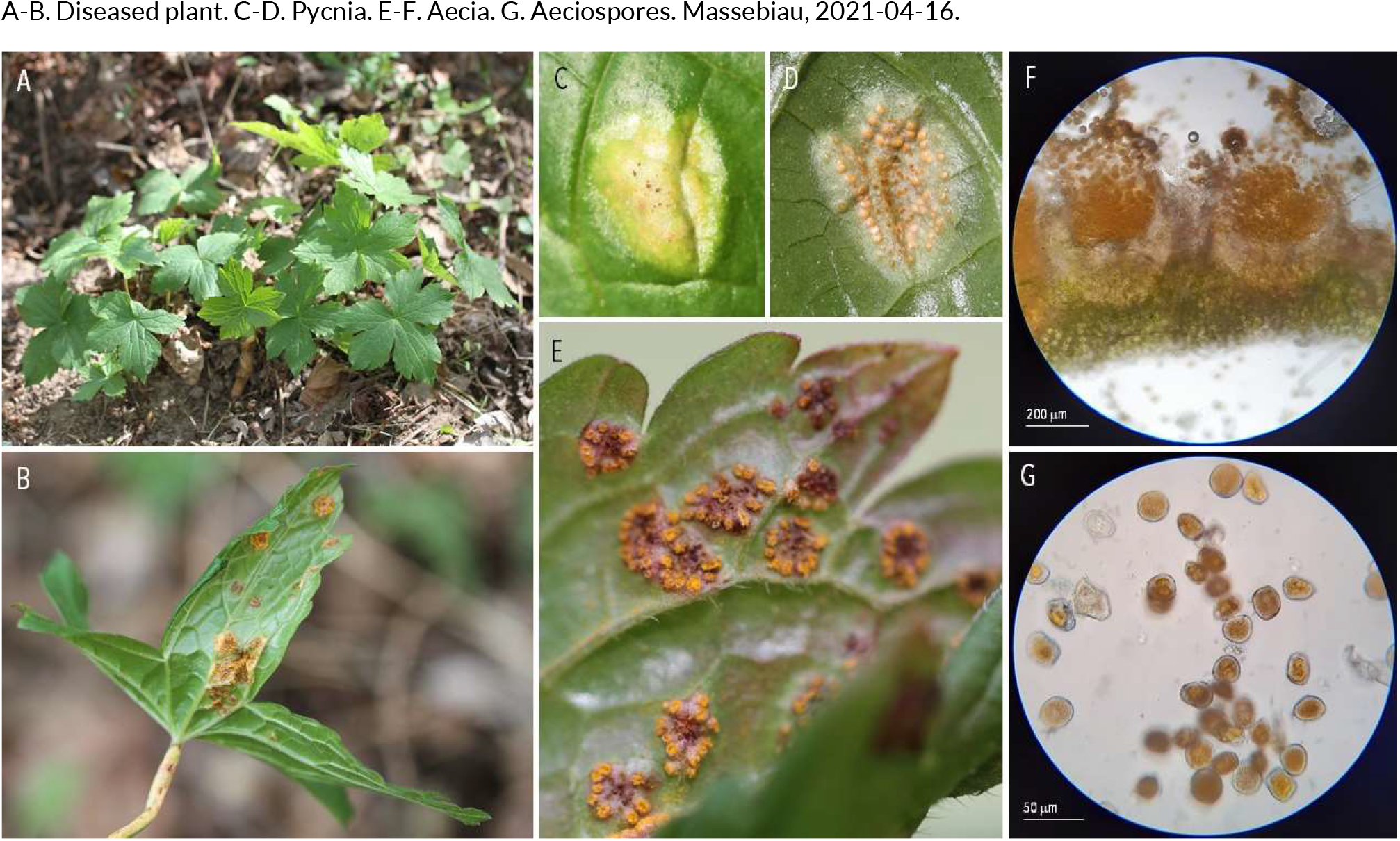
RUSTS | *Puccinia polygoni* ex *Geranium nodosum* (knotted cranesbill)

**69.**
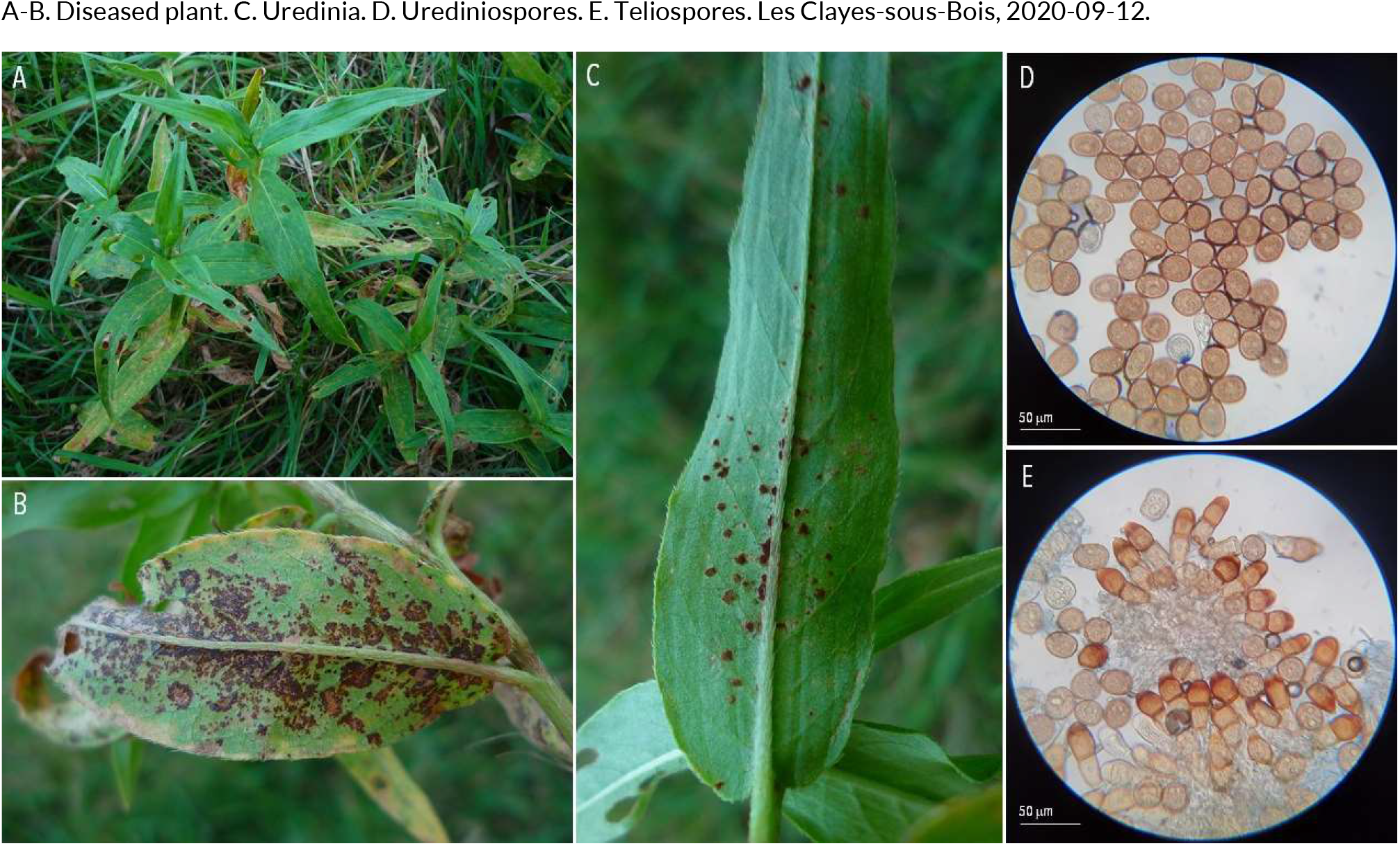
RUSTS | *Puccinia polygoni-amphibii* ex *Persicaria maculosa* (lady’s thumb)

**70.**
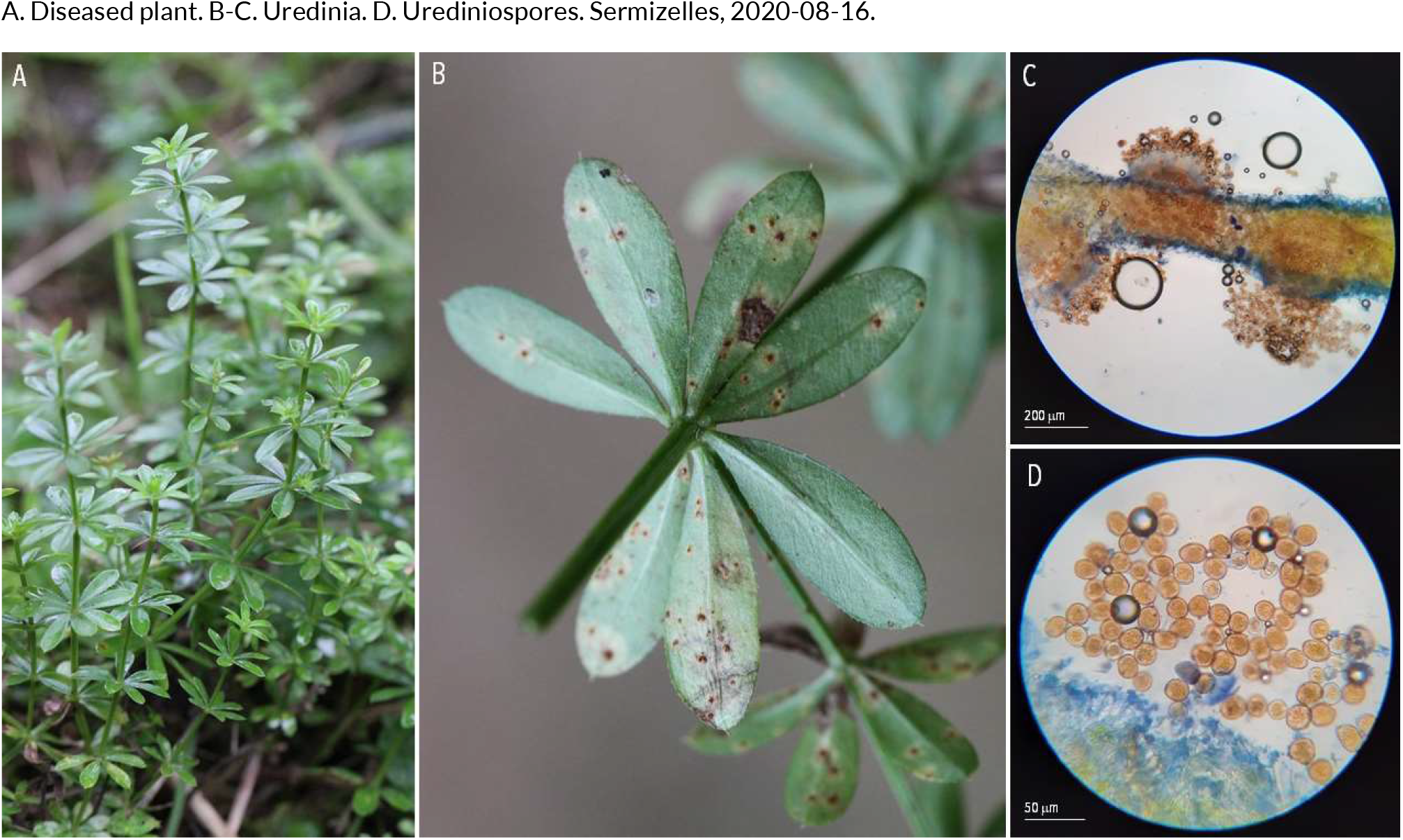
RUSTS | *Puccinia punctata* ex *Galium mollugo* (sweet woodruff)

**71.**
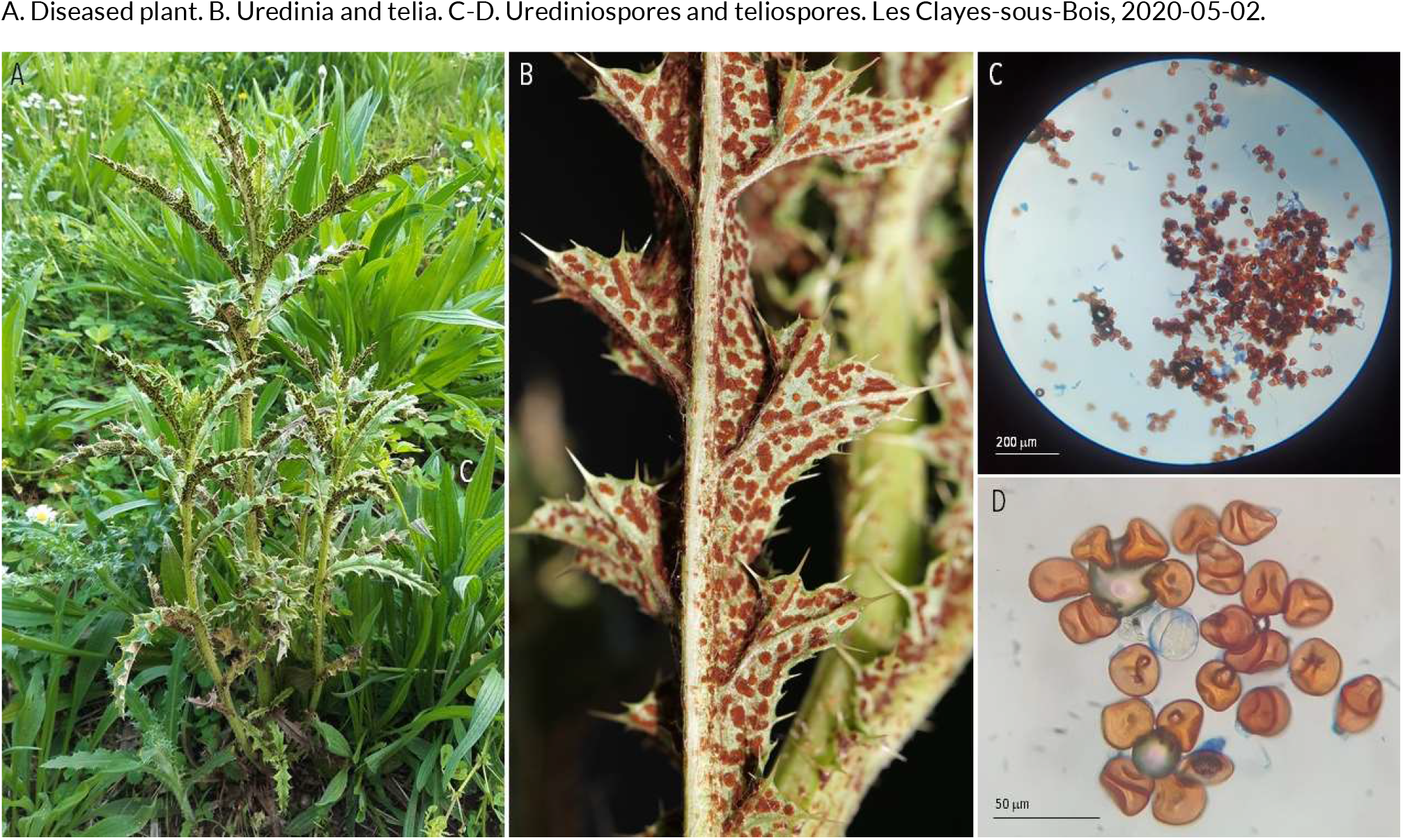
RUSTS | *Puccinia punctiformis* ex *Cirsium arvense* (creeping thistle)

**72.**
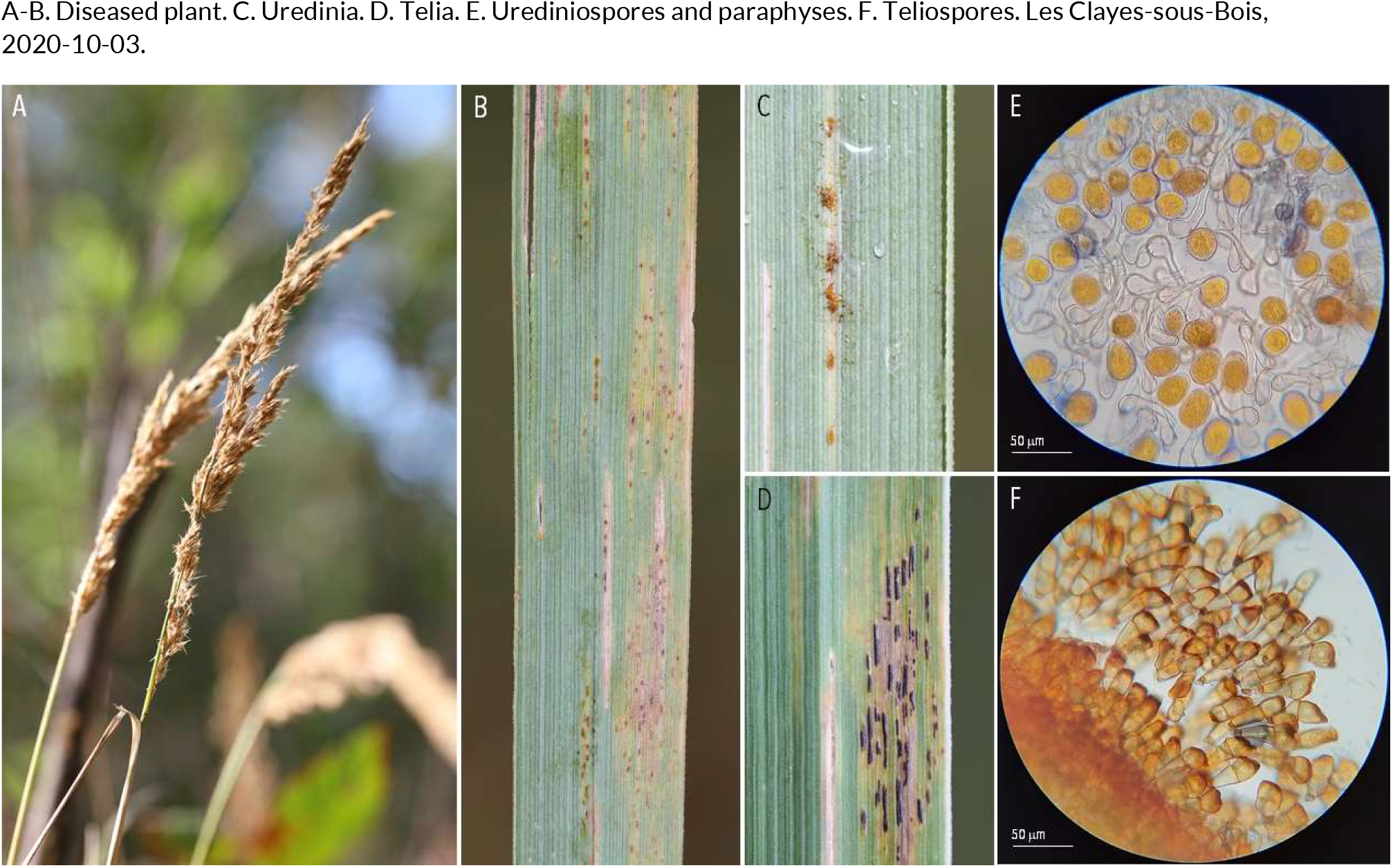
RUSTS | *Puccinia pygmaea* ex *Calamagrostis epigejos* (bushgrass)

**73.**
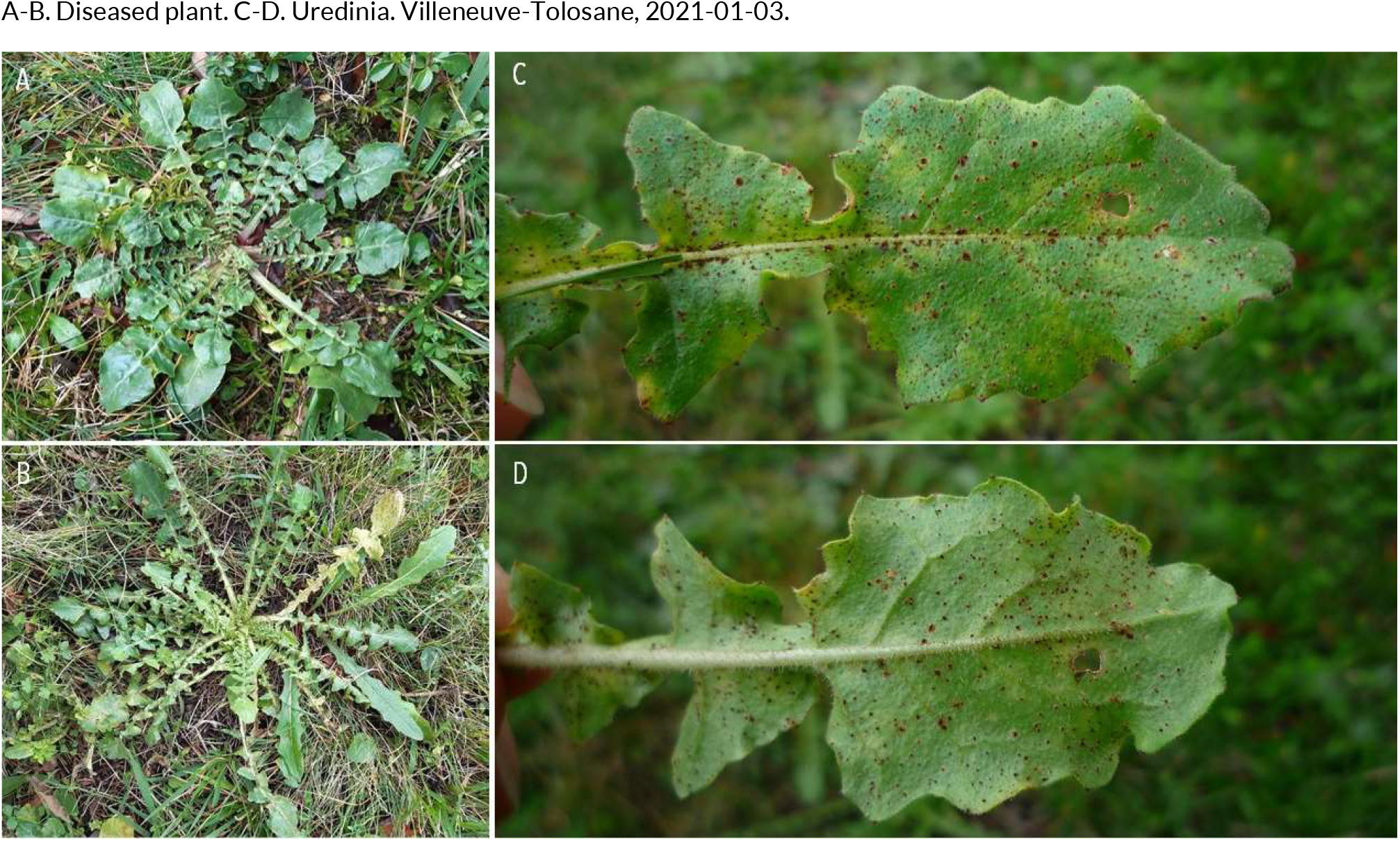
RUSTS | *Puccinia scaliana* ex *Crepis bursifolia* (Italian hawksbeard)

**74.**
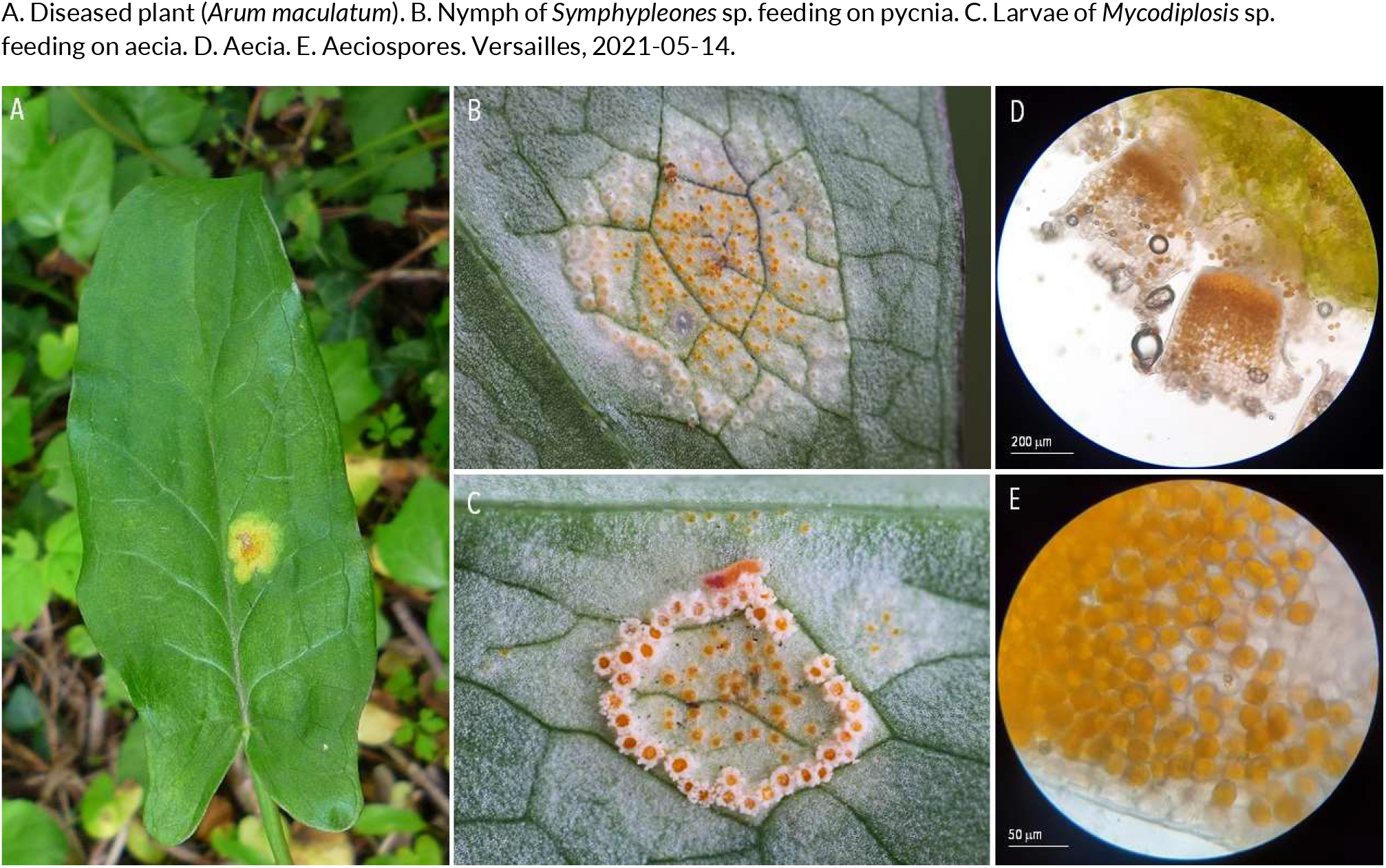
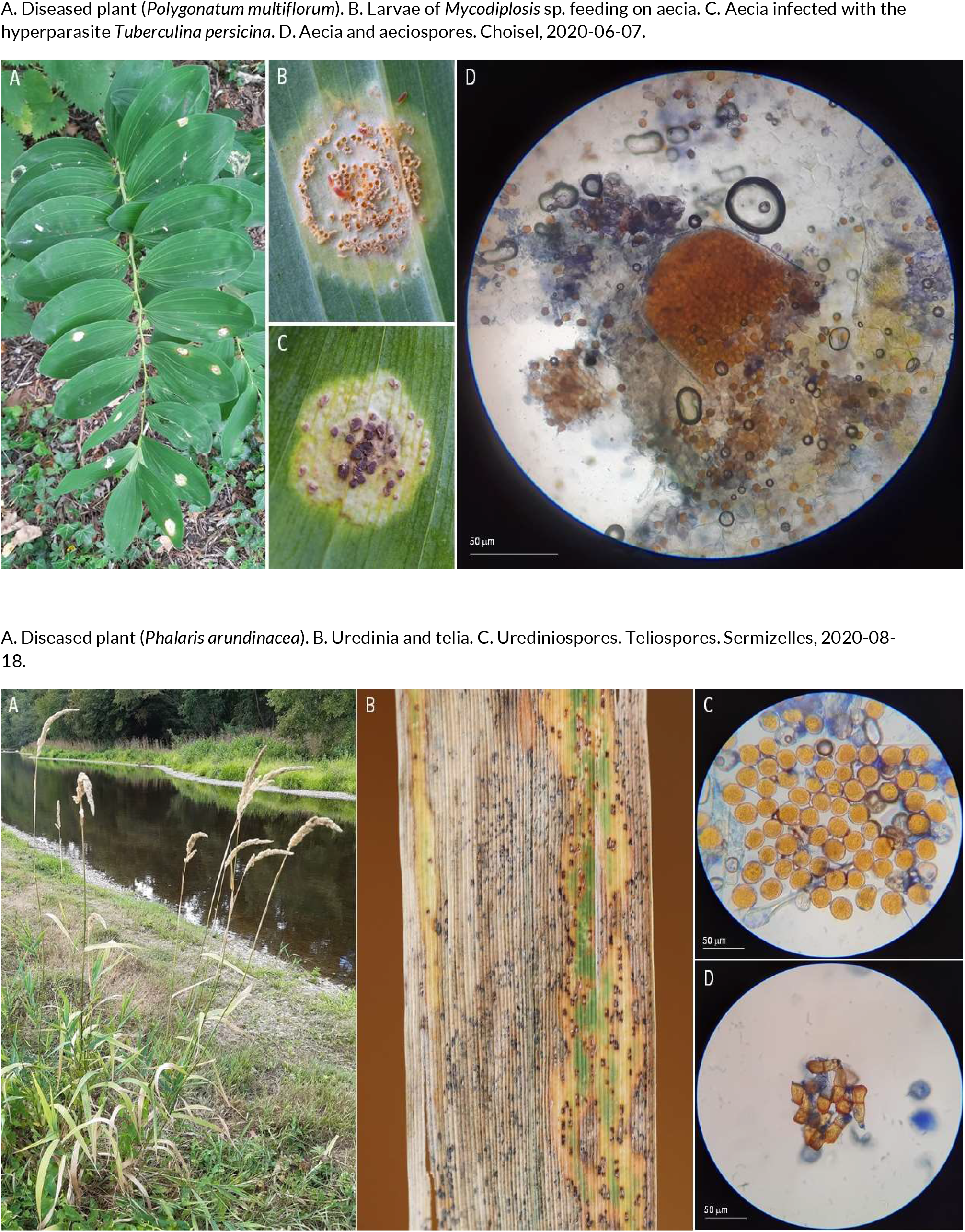
RUSTS | *Puccinia sessilis* ex *Arum maculatum* (cuckoo pint), *Polygonatum multiflorum* (Solomon’s seal) and *Phalaris arundinacea* (reed canary grass)

**75.**
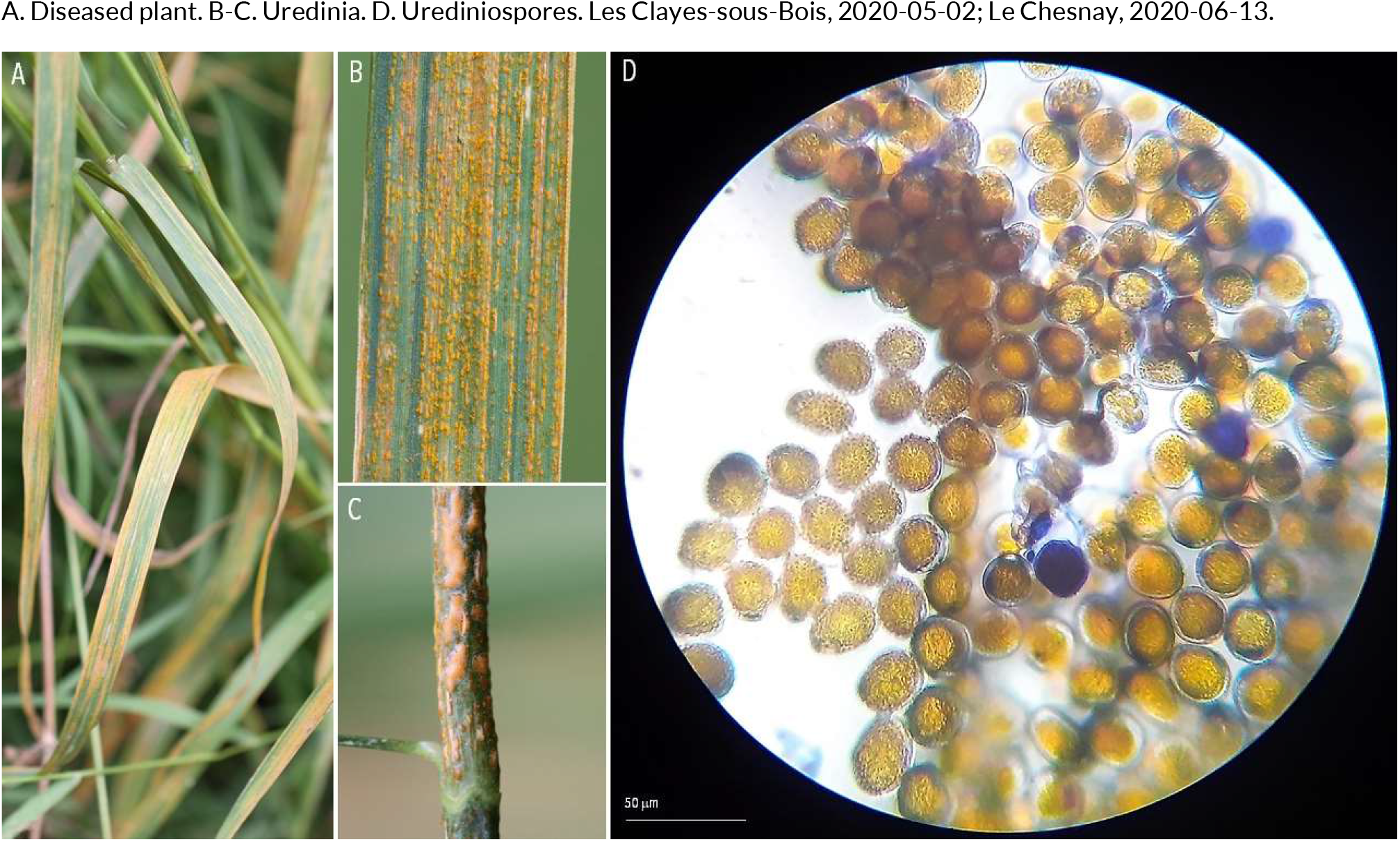
RUSTS | *Puccinia striiformis* f. sp. *dactylidis* ex *Dactylis glomerata* (cock’s-foot)

**76.**
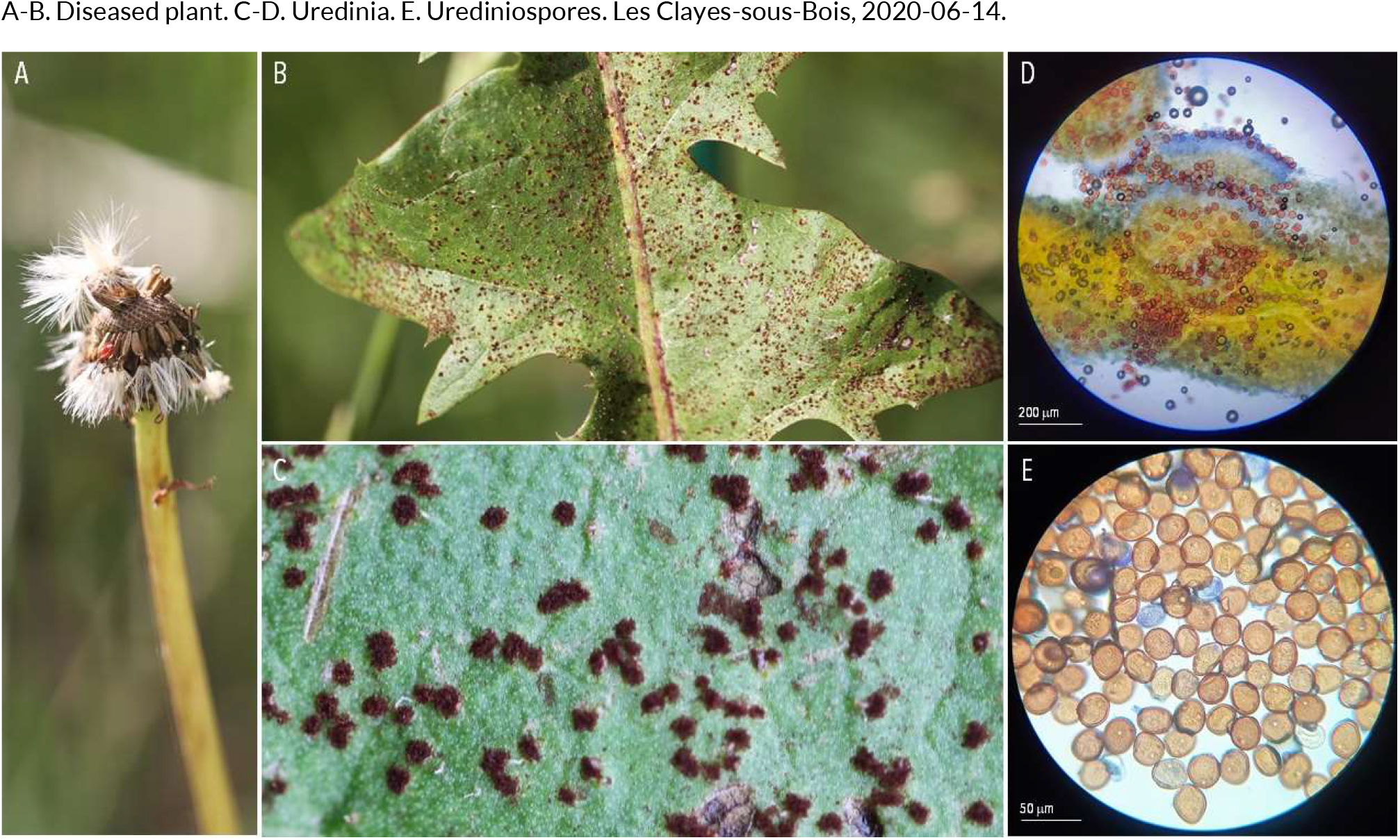
RUSTS | *Puccinia taraxaci* ex *Taraxacum officinale* (common dandelion)

**77.**
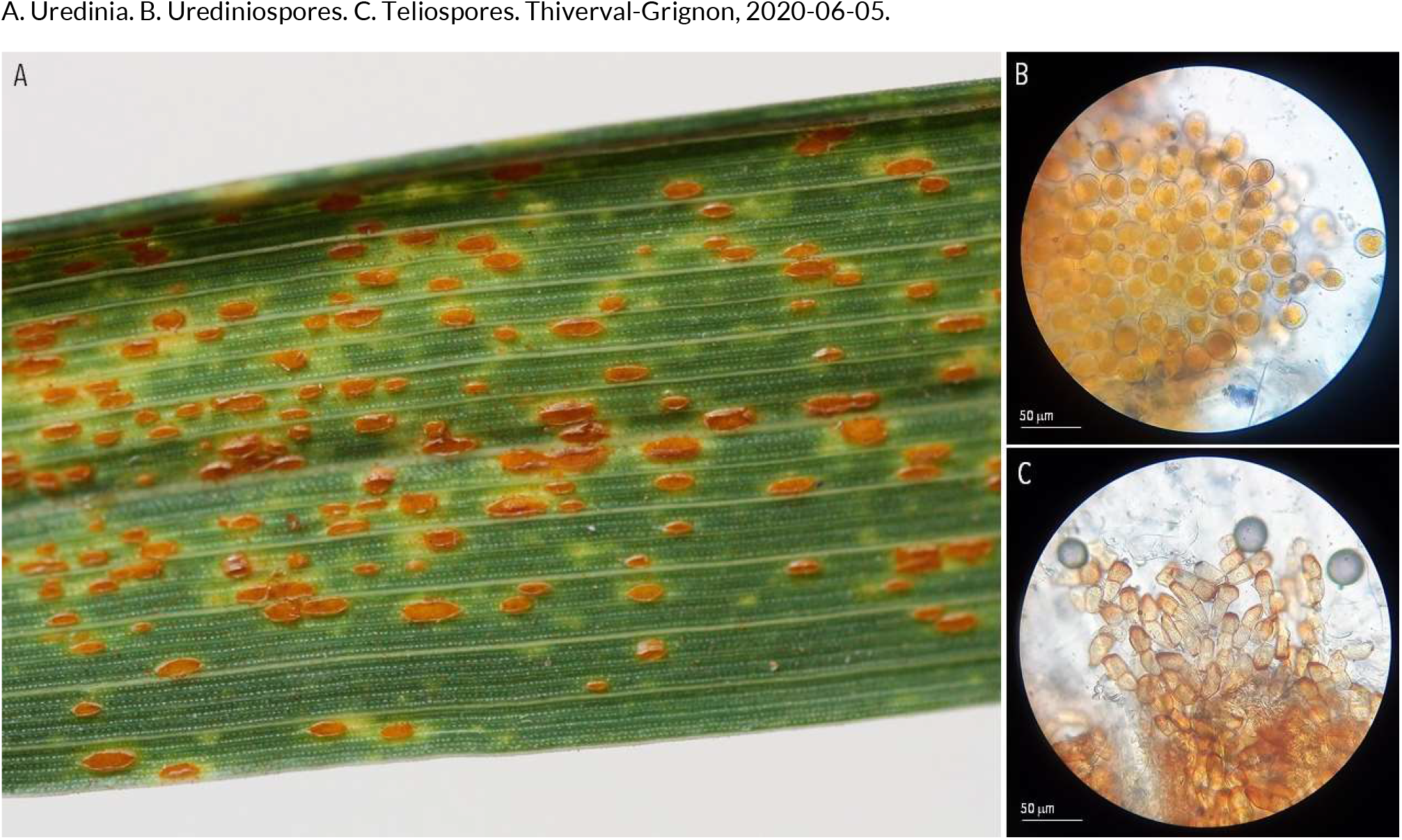
RUSTS | *Puccinia triticina* ex *Triticum aestivum* (bread wheat)

**78.**
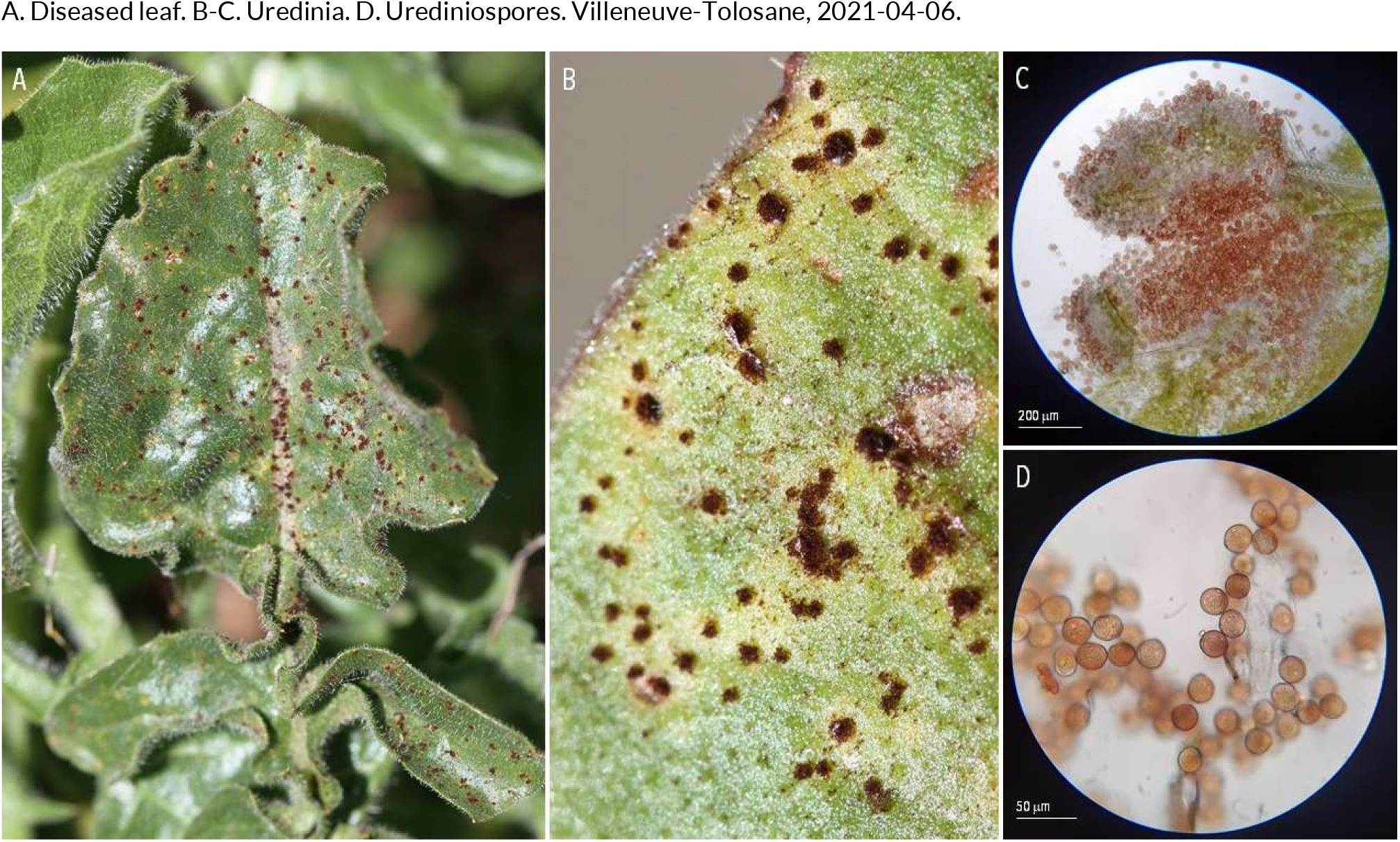
RUSTS | *Puccinia urospermi* ex *Urospermum dalechampii* (smooth golden fleece)

**79.**
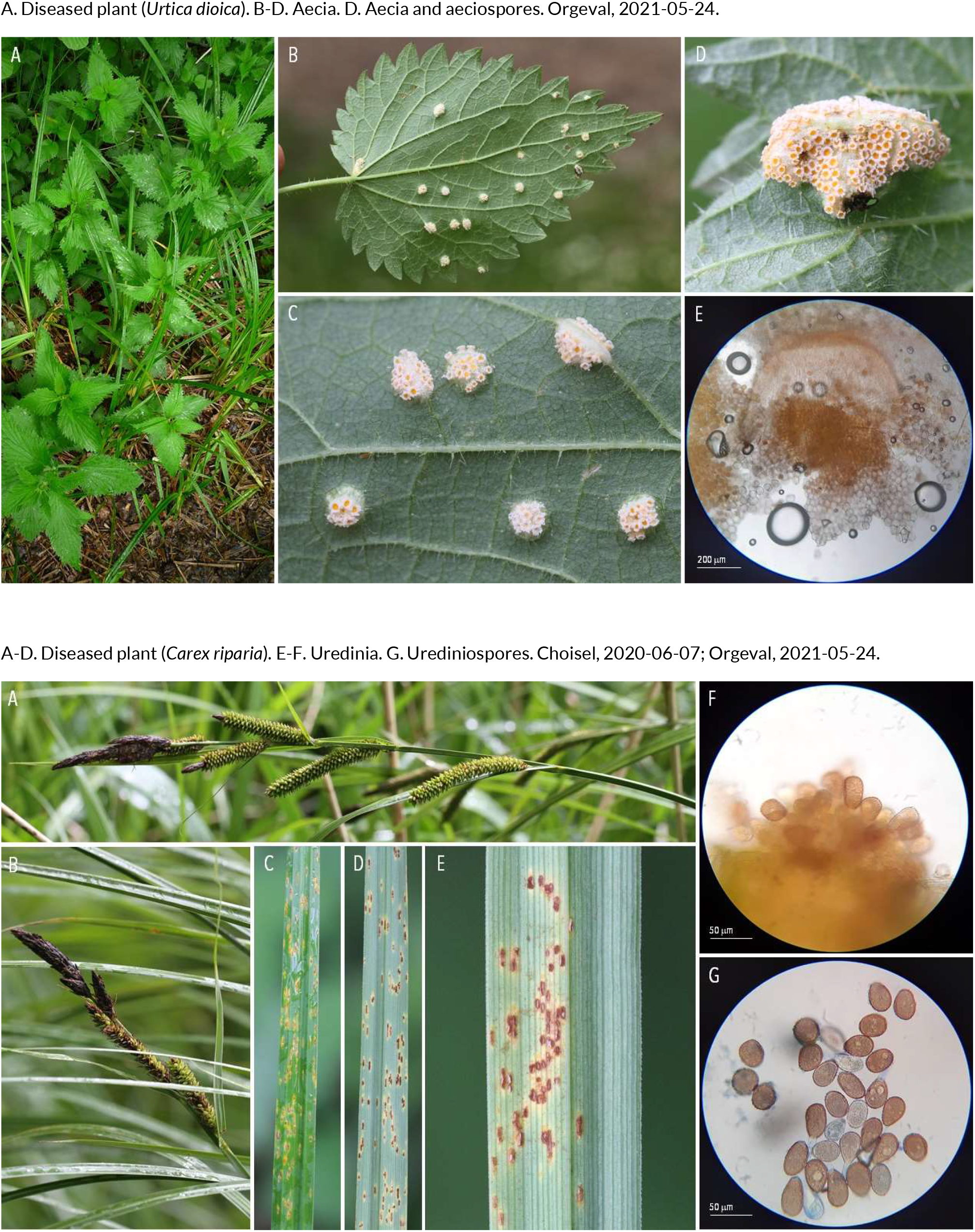
RUSTS | *Puccinia urticae-acutiformis* ex *Urtica dioca* (nettle) and *Carex riparia* (greater pond sedge)

**80.**
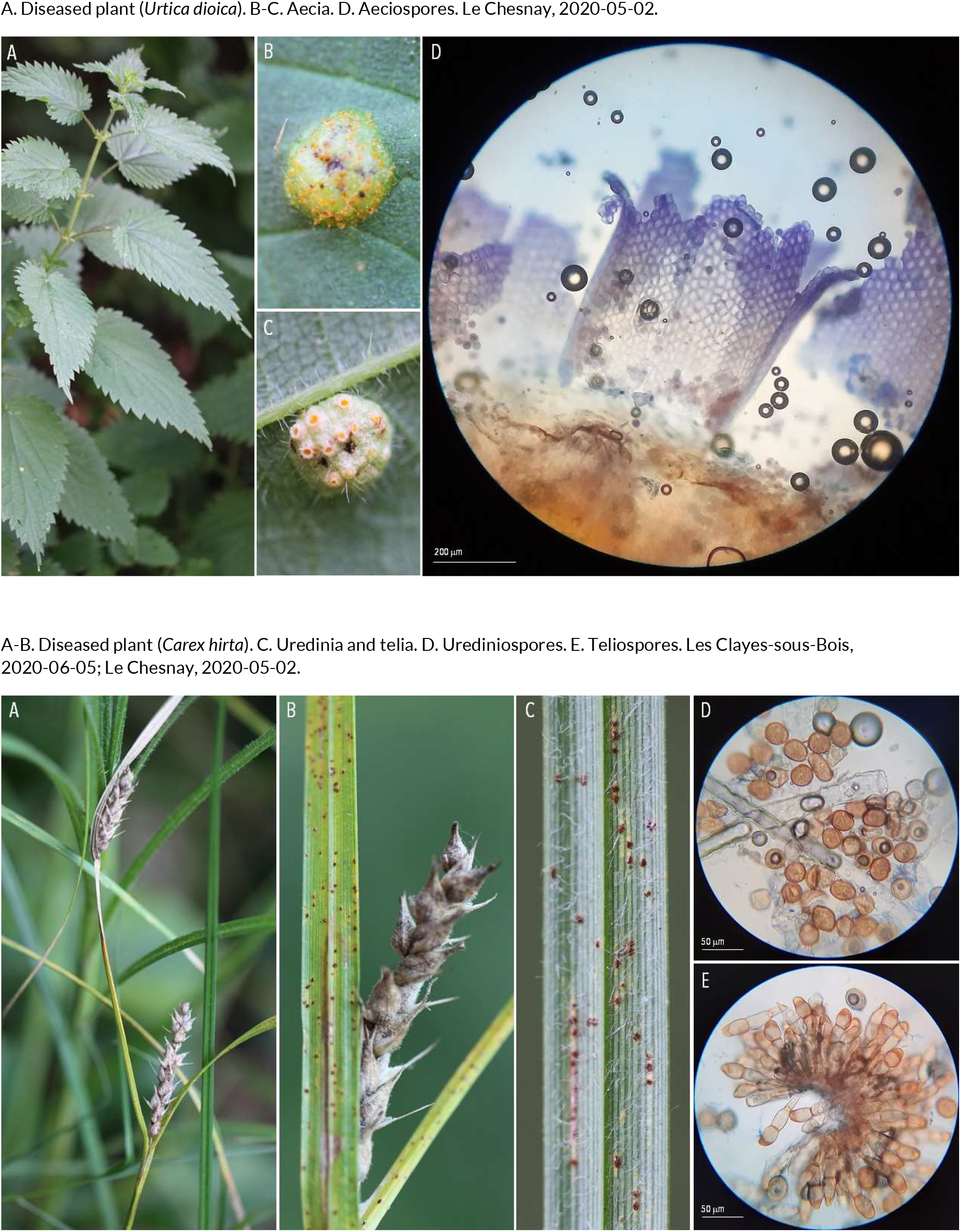
RUSTS | *Puccinia urticae-hirtae* ex *Urtica dioica* (nettle) and *Carex hirta* (hairy sedges)

**81.**
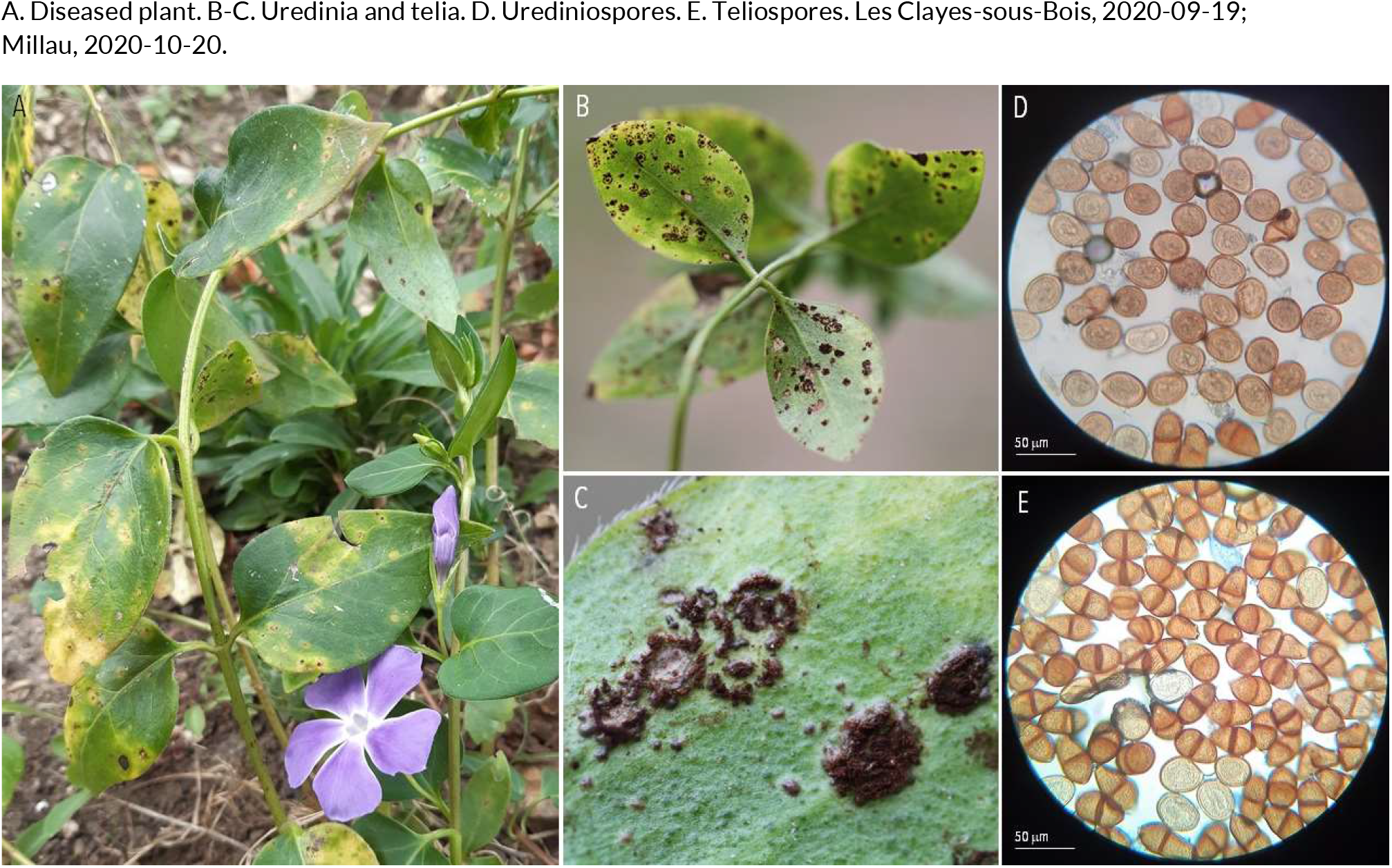
RUSTS | *Puccinia vincae* ex *Vinca major* (bigleaf periwinkle)

**82.**
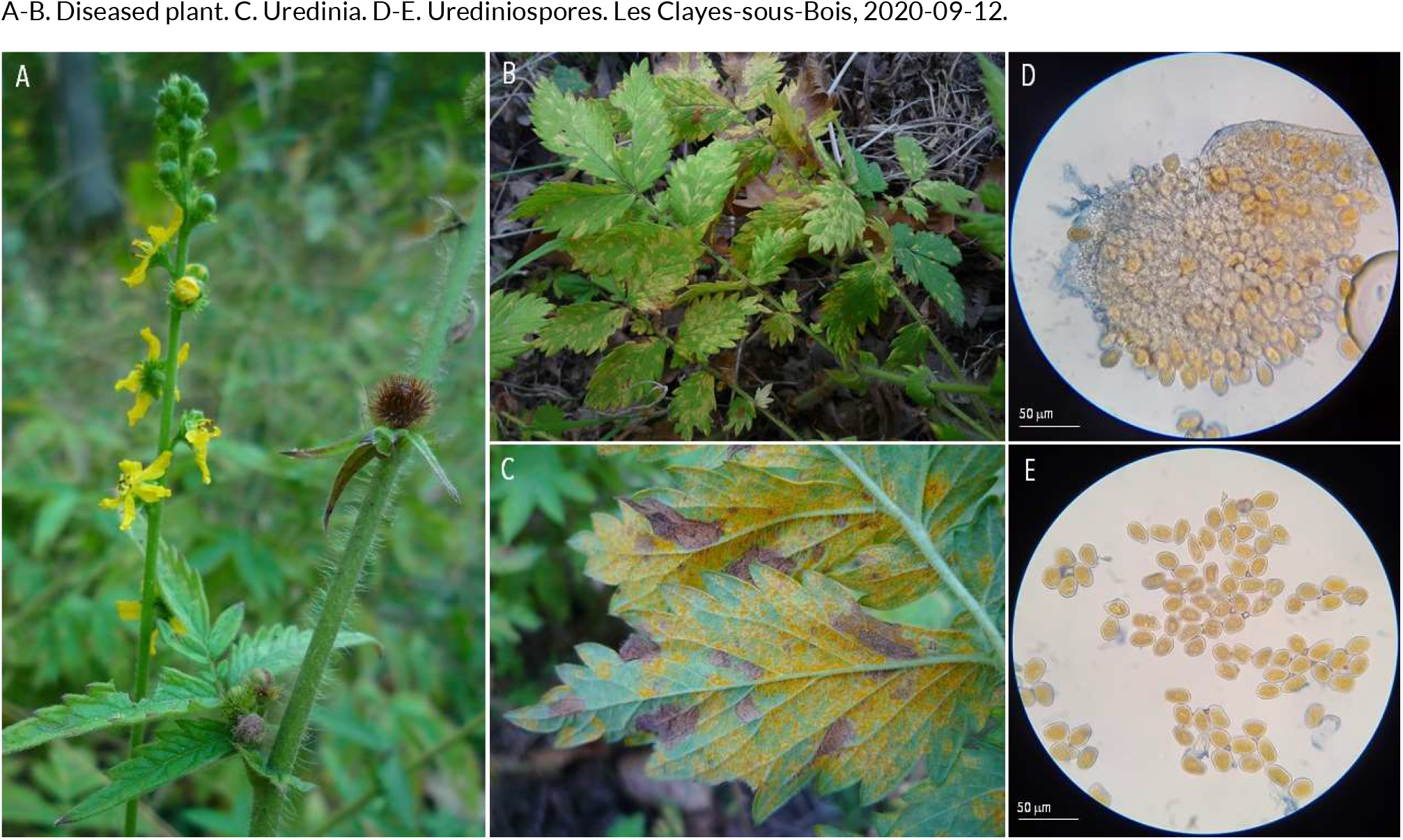
RUSTS | *Pucciniastrum agrimoniae* ex *Agrimonia eupatoria* (agrimony)

**83.**
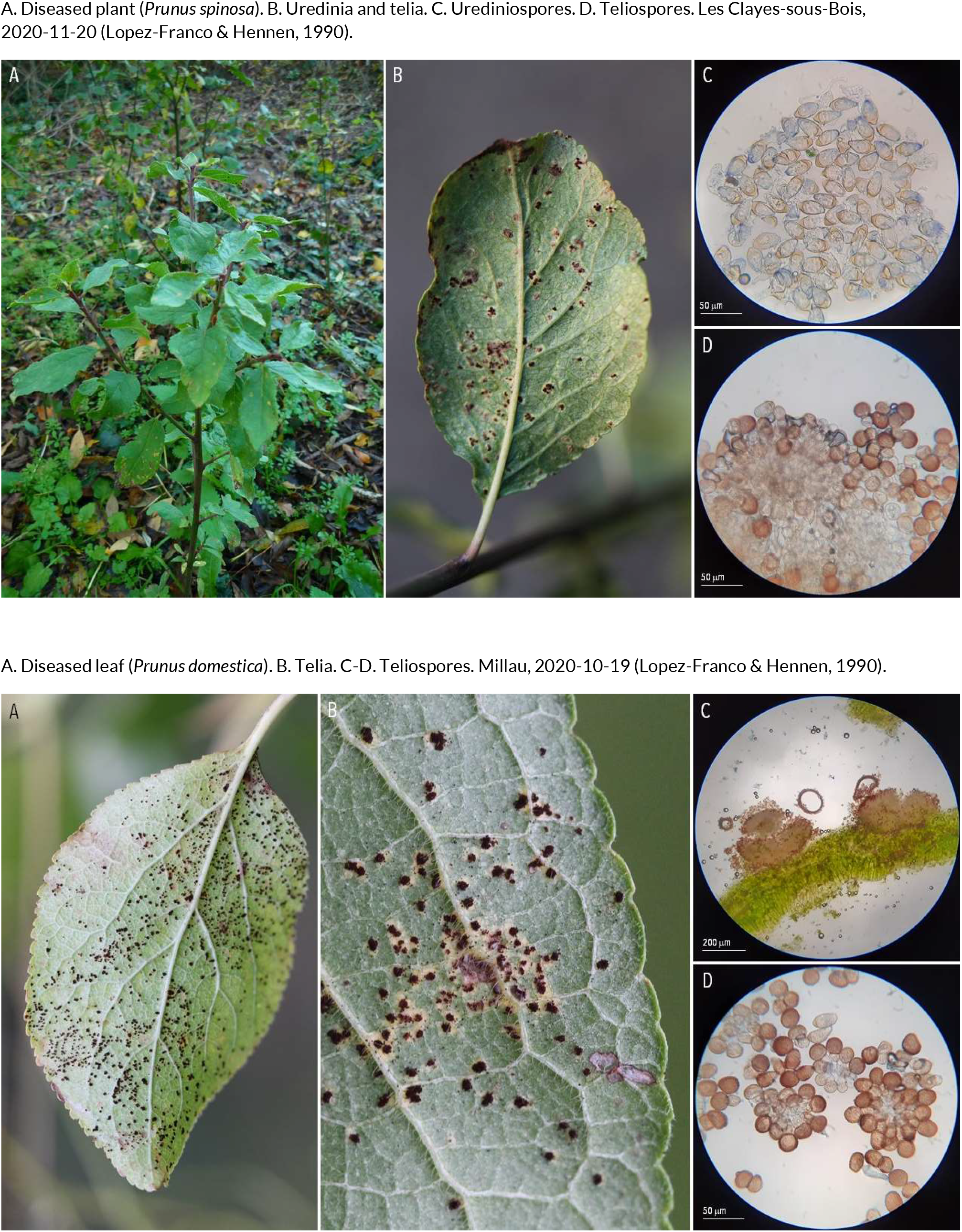
RUSTS | *Tranzschelia discolor* ex *Prunus spinosa* (blackthorn)

**84.**
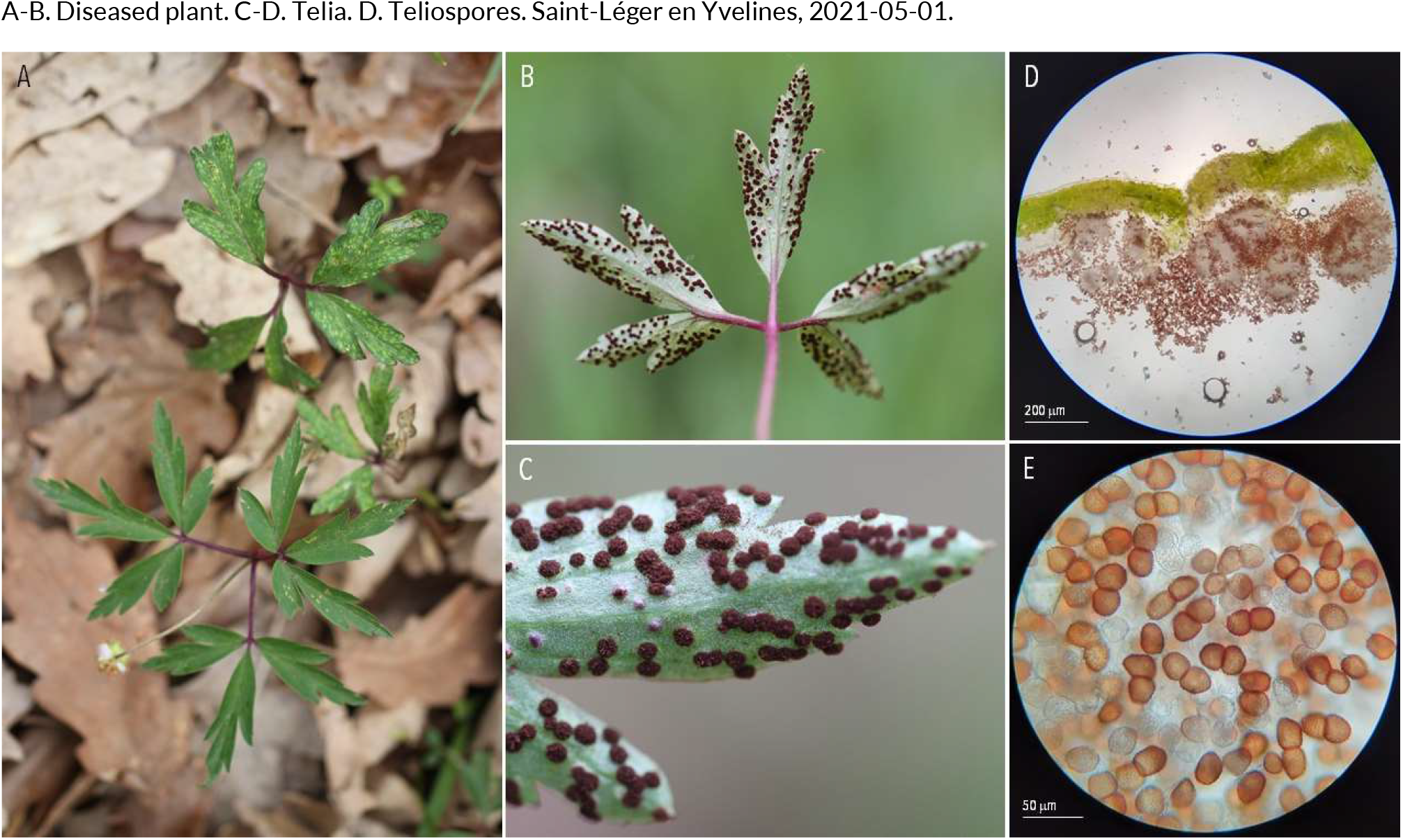
RUSTS | *Tranzschelia fusca* ex *Anemone nemorosa* (wood anemone)

**85.**
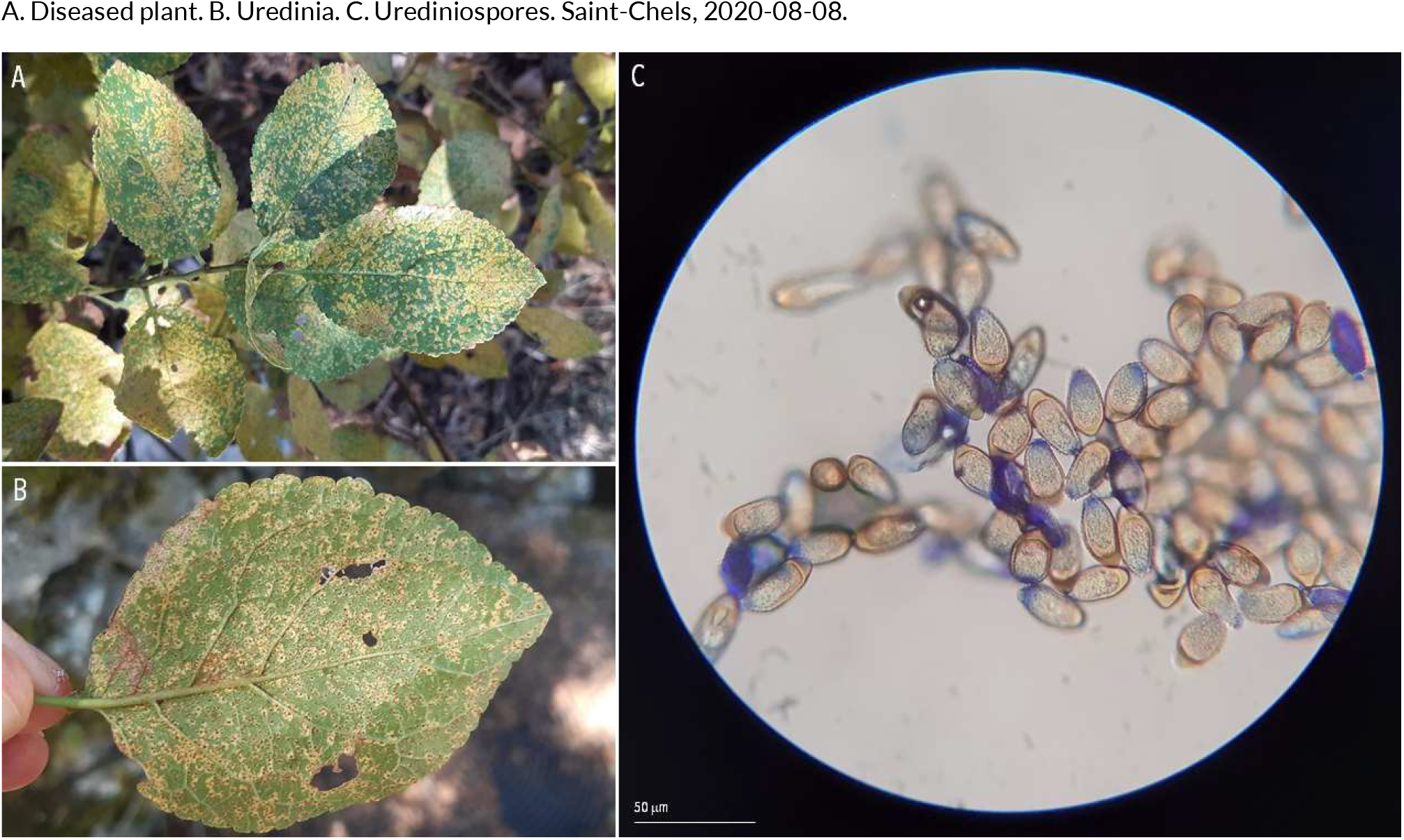
RUSTS | *Tranzschelia pruni-spinosae* (?) ex *Prunus sp*. (plum)

**86.**
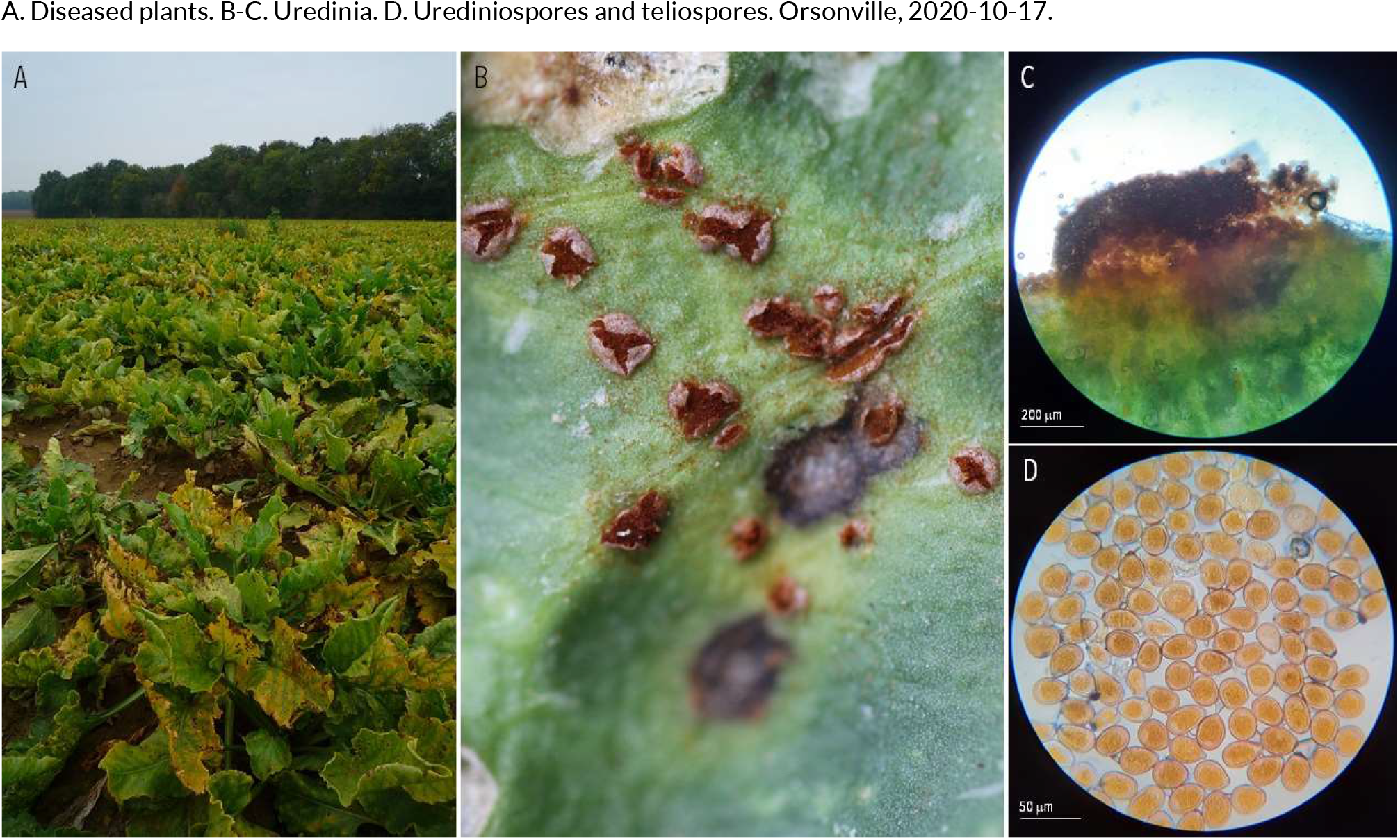
RUSTS | *Uromyces beticola* ex *Beta vulgaris* (beet)

**87.**
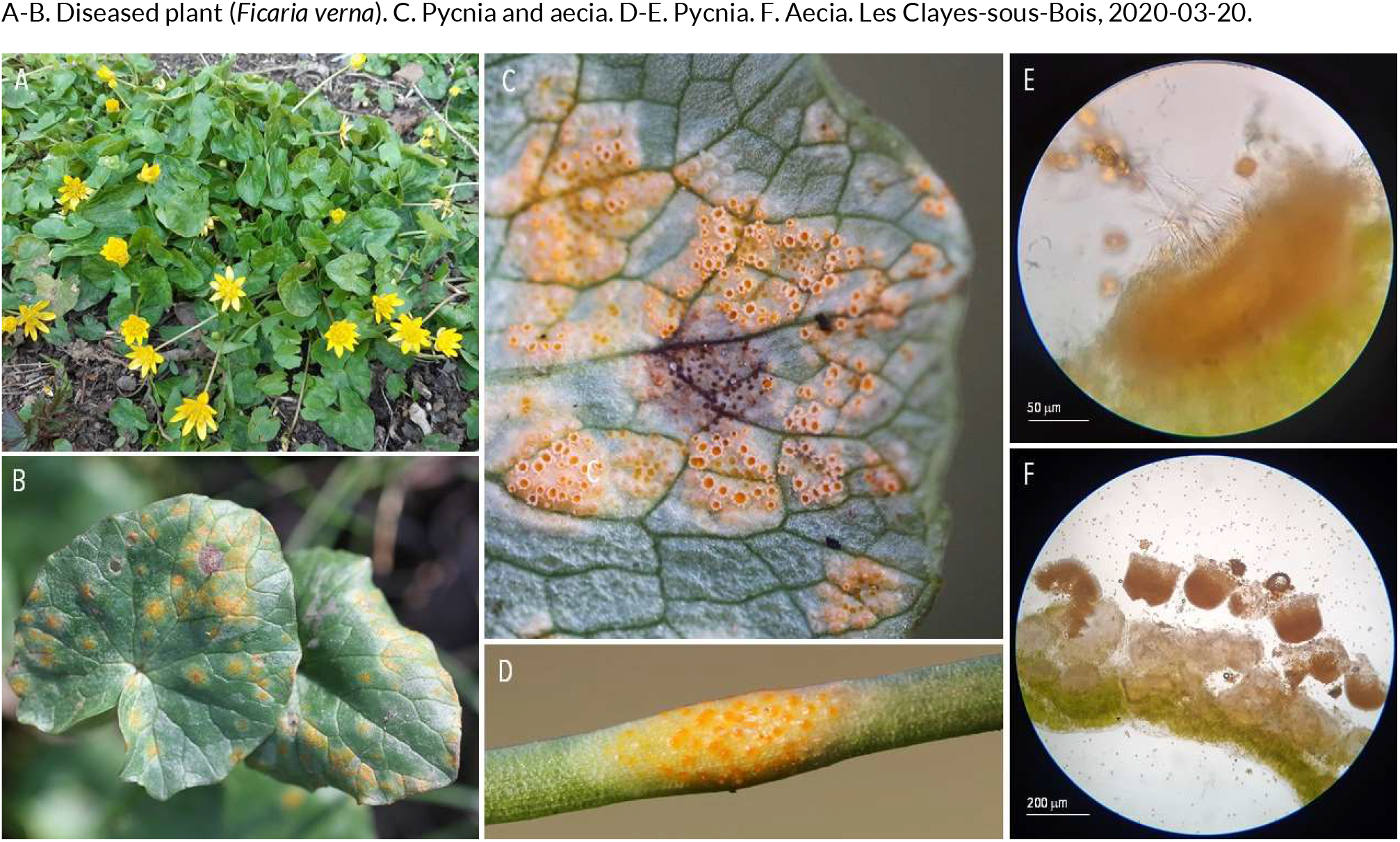
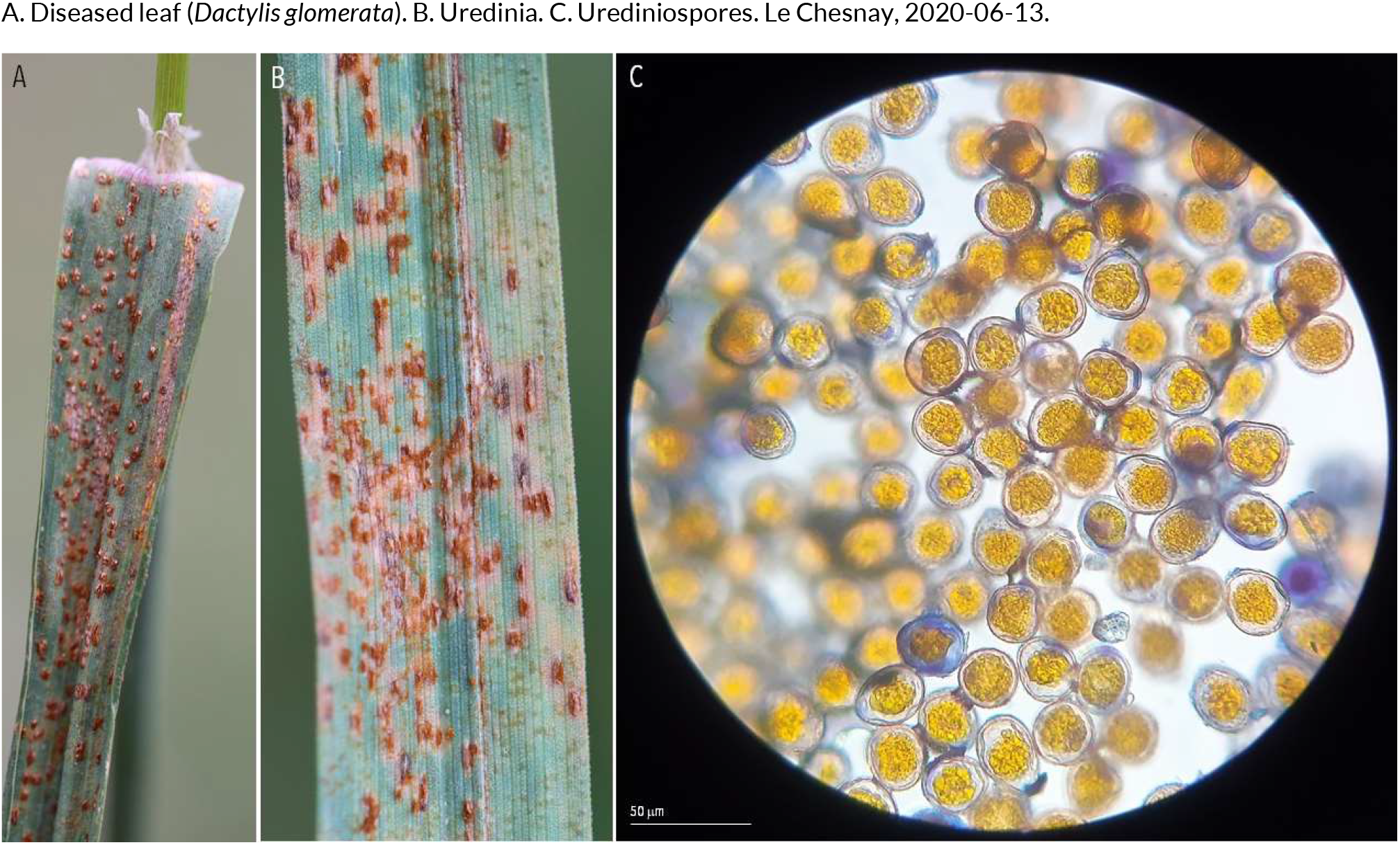
RUSTS | *Uromyces dactylidis* ex *Ficaria verna* (lesser celandine) and *Dactylis glomerata* (cock’s-foot)

**88.**
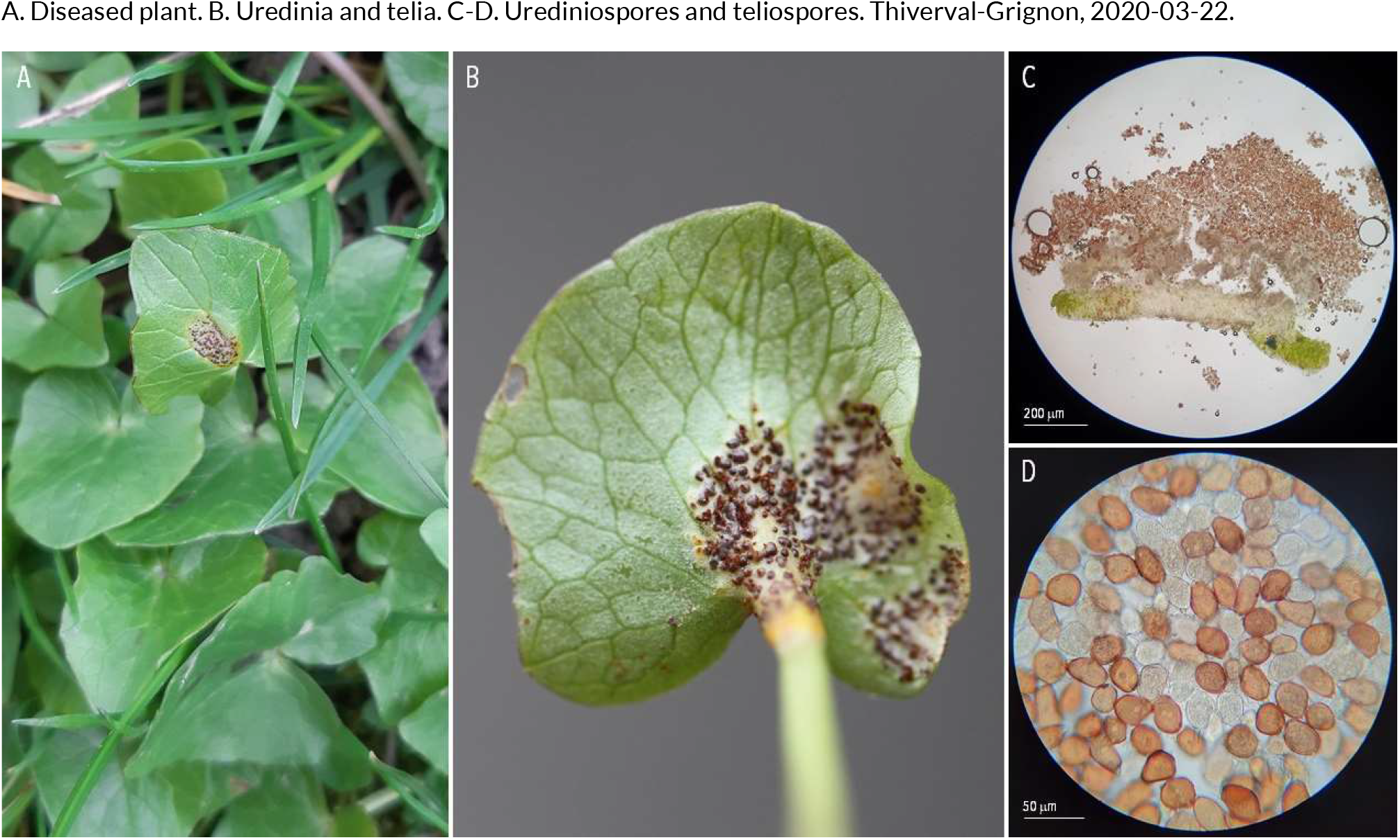
RUSTS | *Uromyces ficariae* ex *Ficaria verna* (lesser celandine)

**89.**
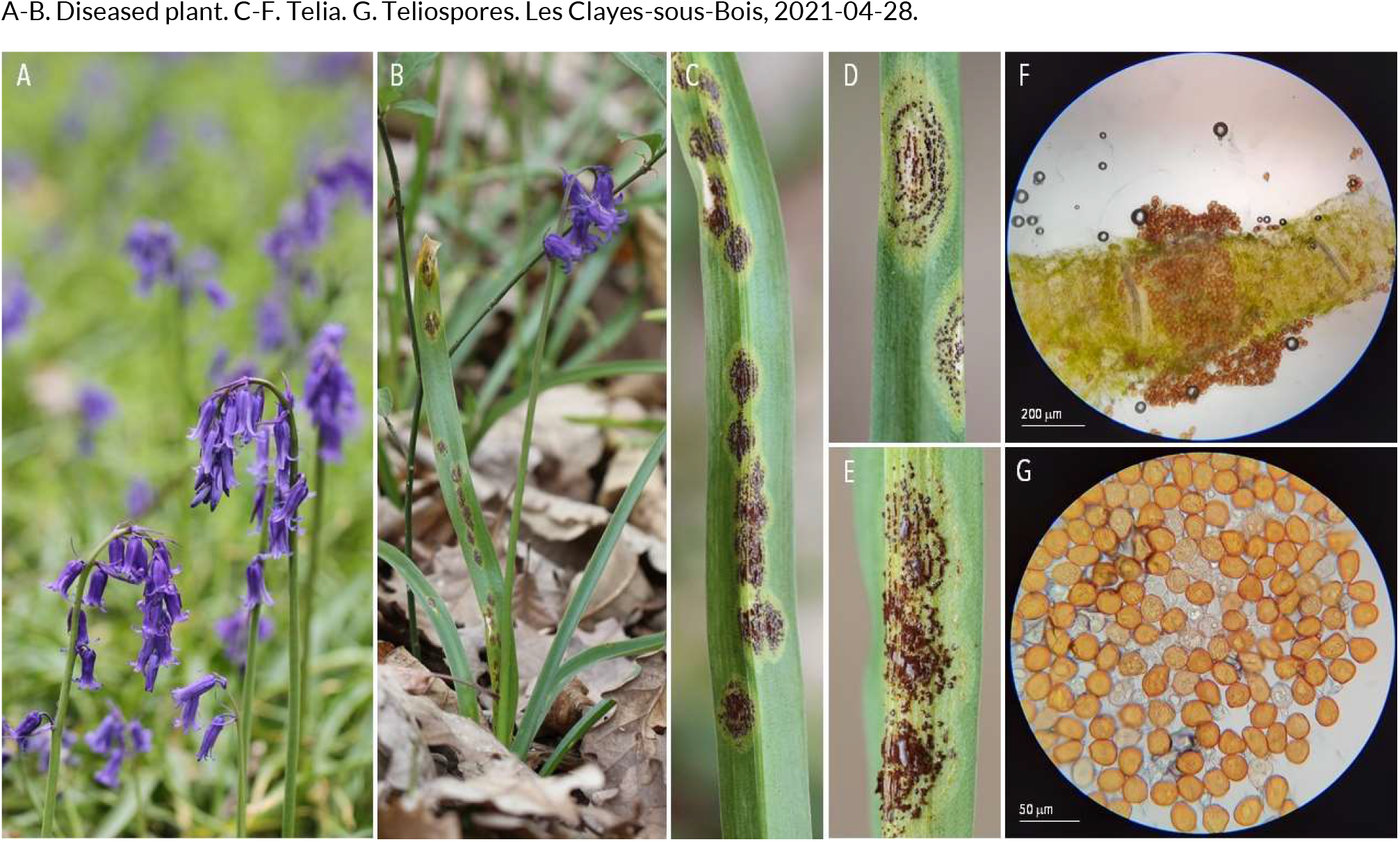
RUSTS | *Uromyces hyacinthi* ex *Hyacinthoides non-scripta* (common bluebell)

**90.**
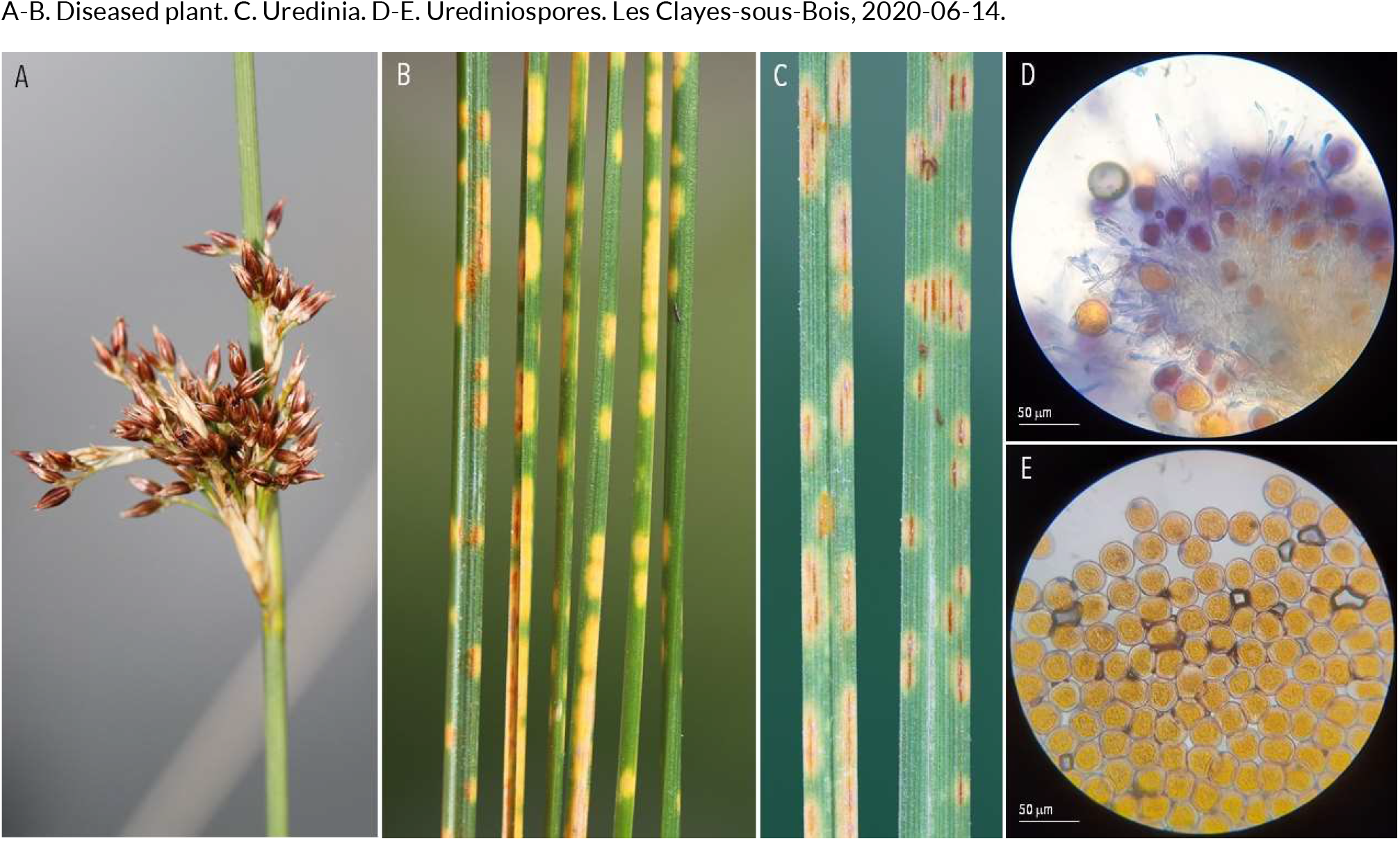
RUSTS | *Uromyces junci* ex *Juncus inflexus* (rush)

**91.**
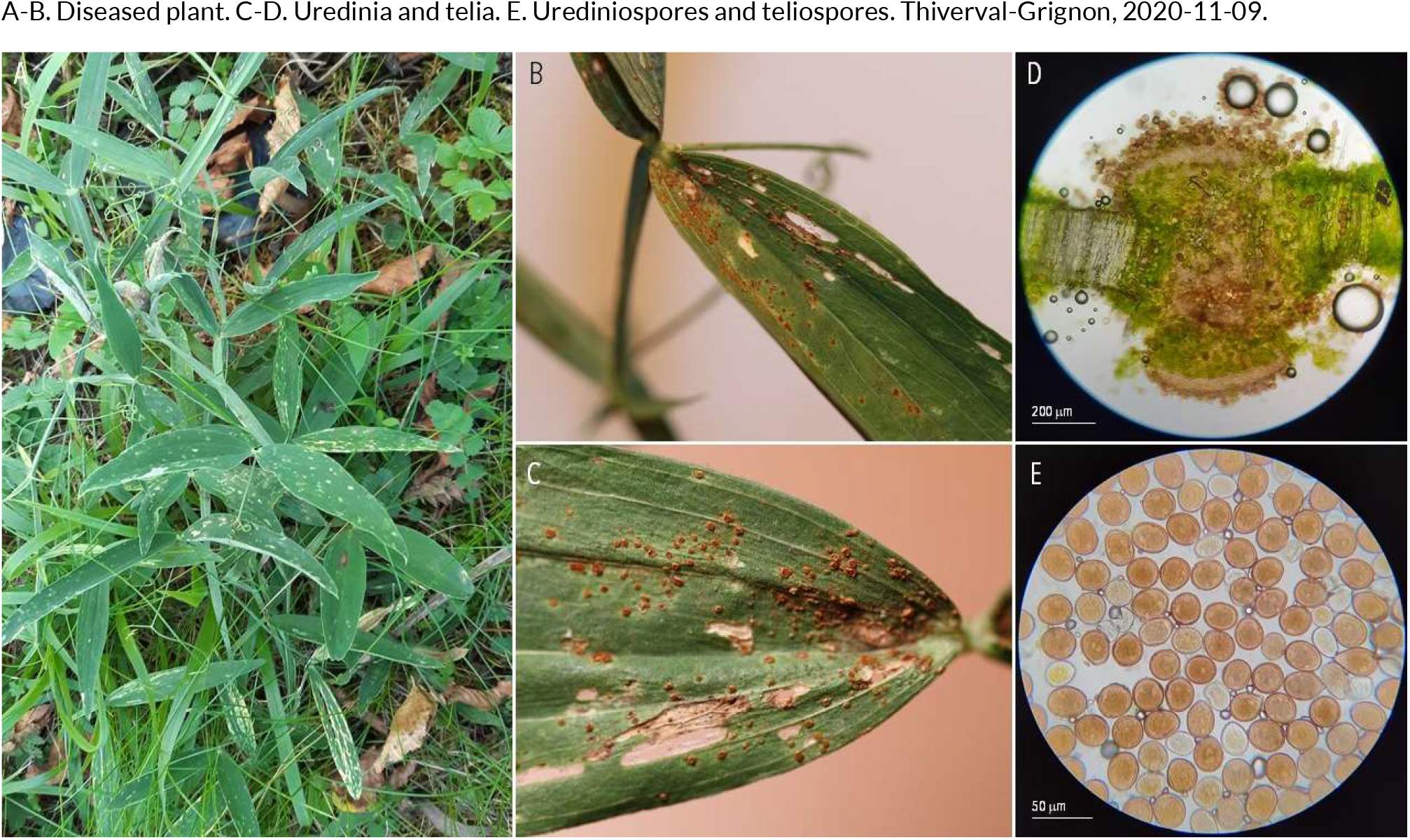
RUSTS | *Uromyces lathyri-latifoli* ex *Lathyrus latifolius* (everlasting pea)

**92.**
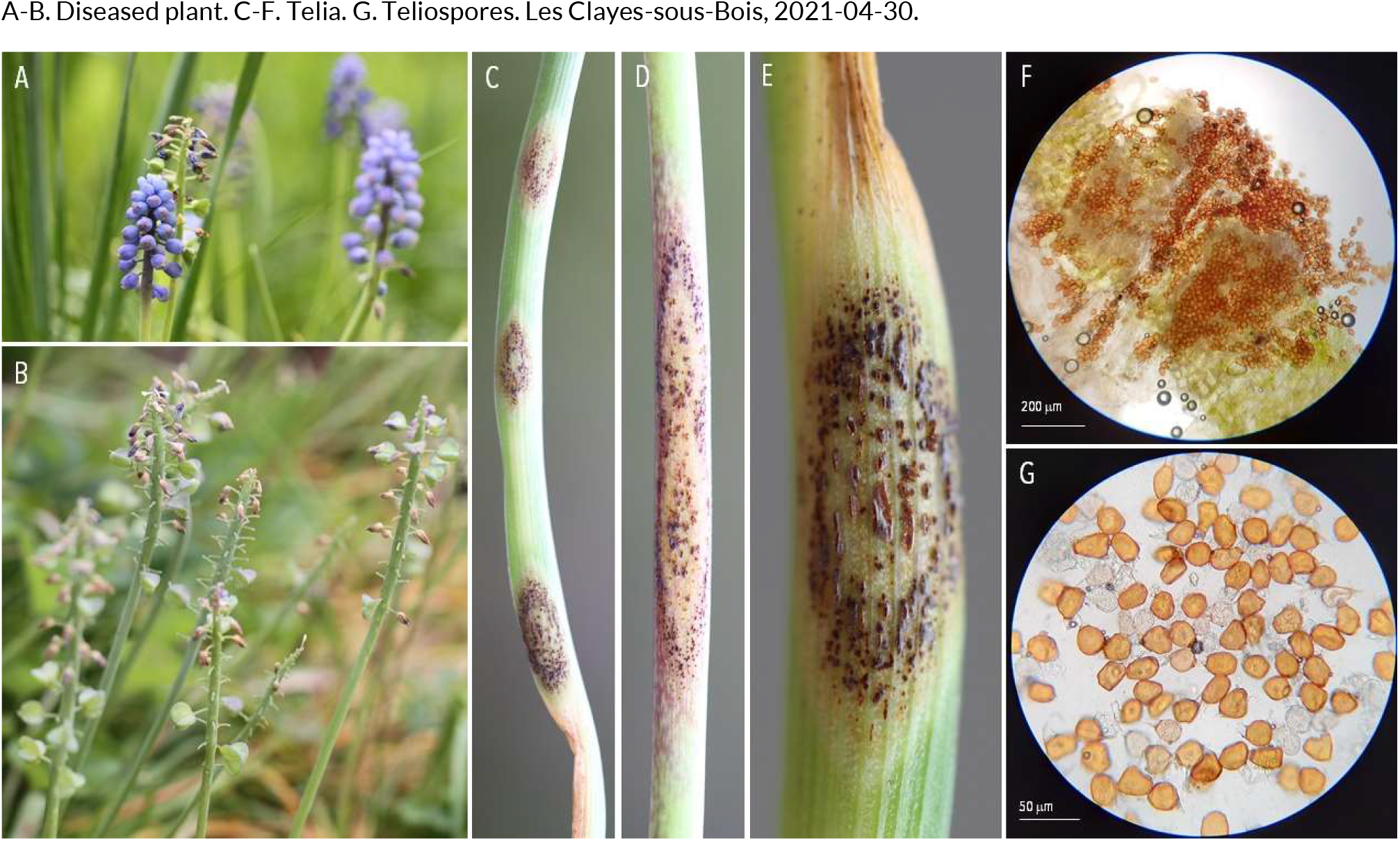
RUSTS | *Uromyces muscari* ex *Muscari armeniacum* (grape hyacinth)

**93.**
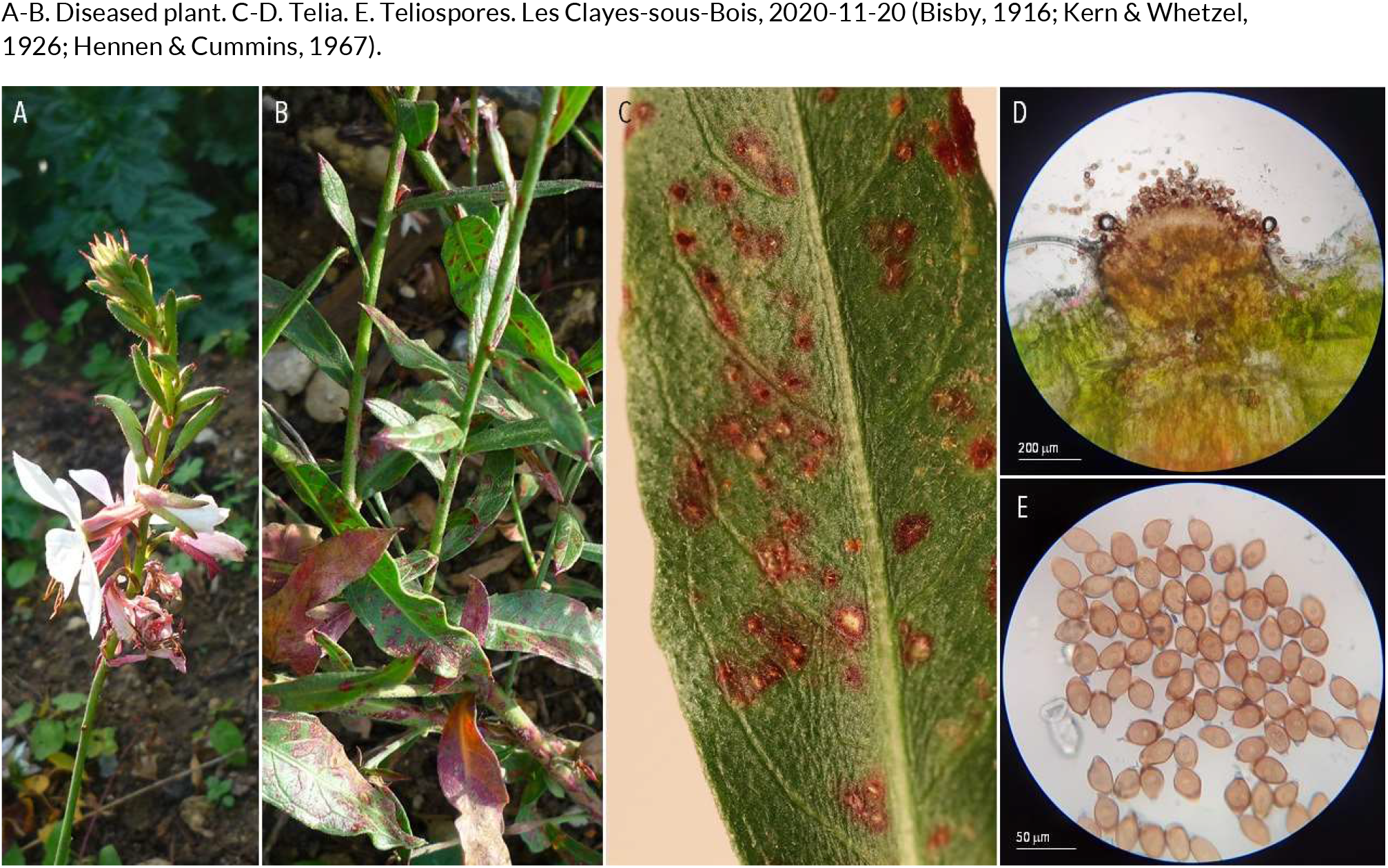
RUSTS | *Uromyces plumbarius* ex *Oenothera gaura syn. Gaura biennis* (biennal gaura)

**94.**
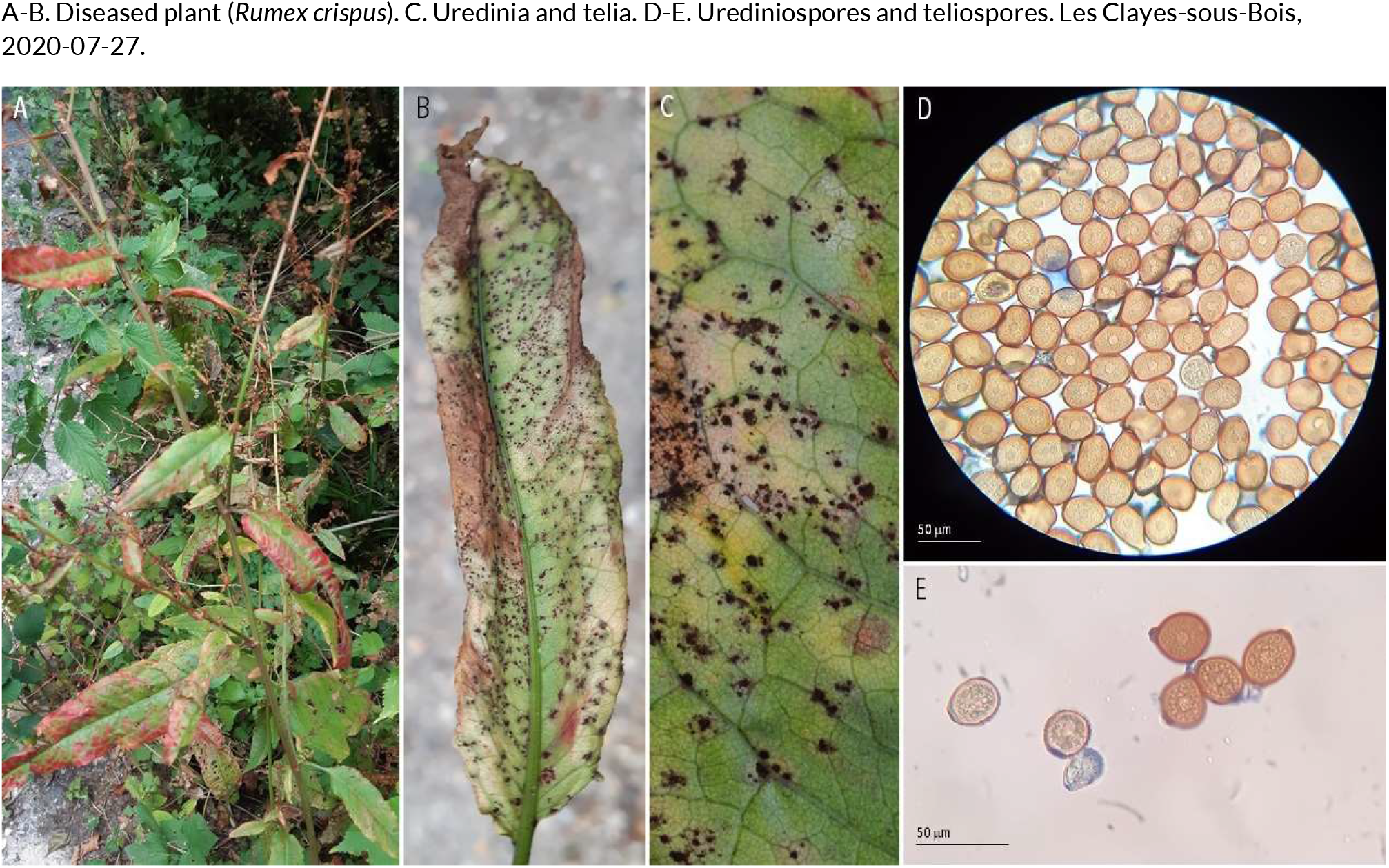
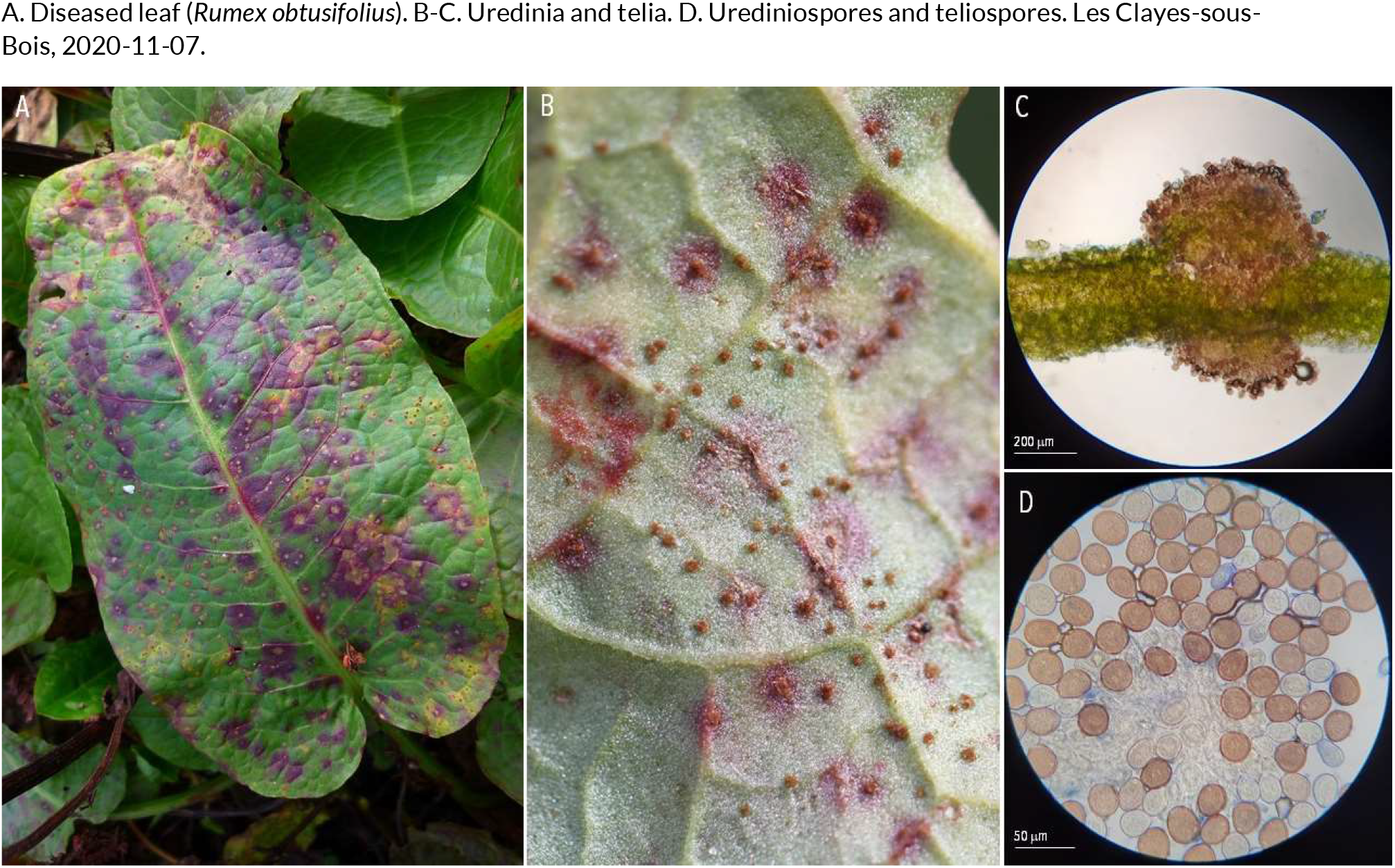
RUSTS | *Uromyces rumicis* ex *Rumex crispus* (curly dock) and *Rumex obtusifolius* (bitter dock)

**95.**
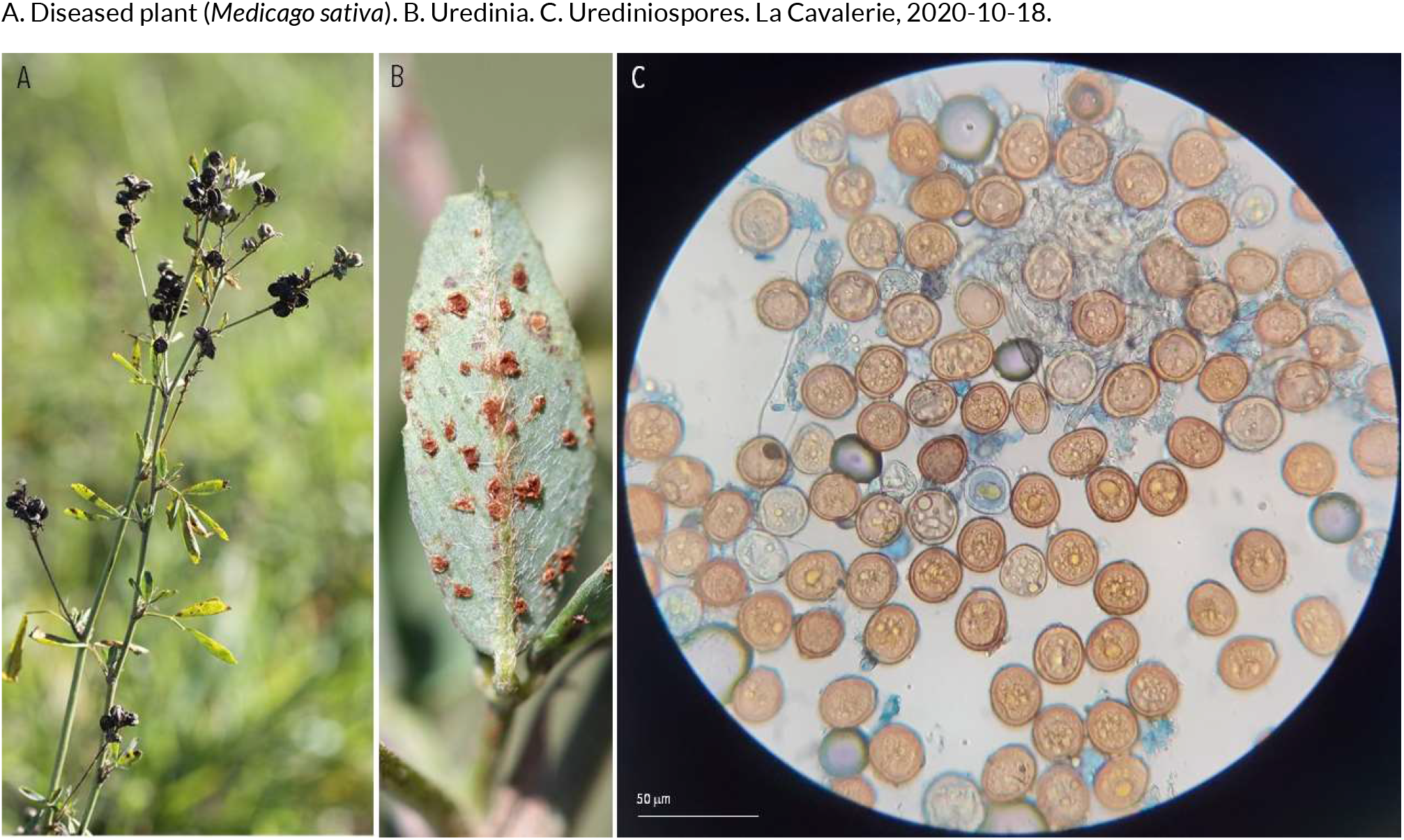
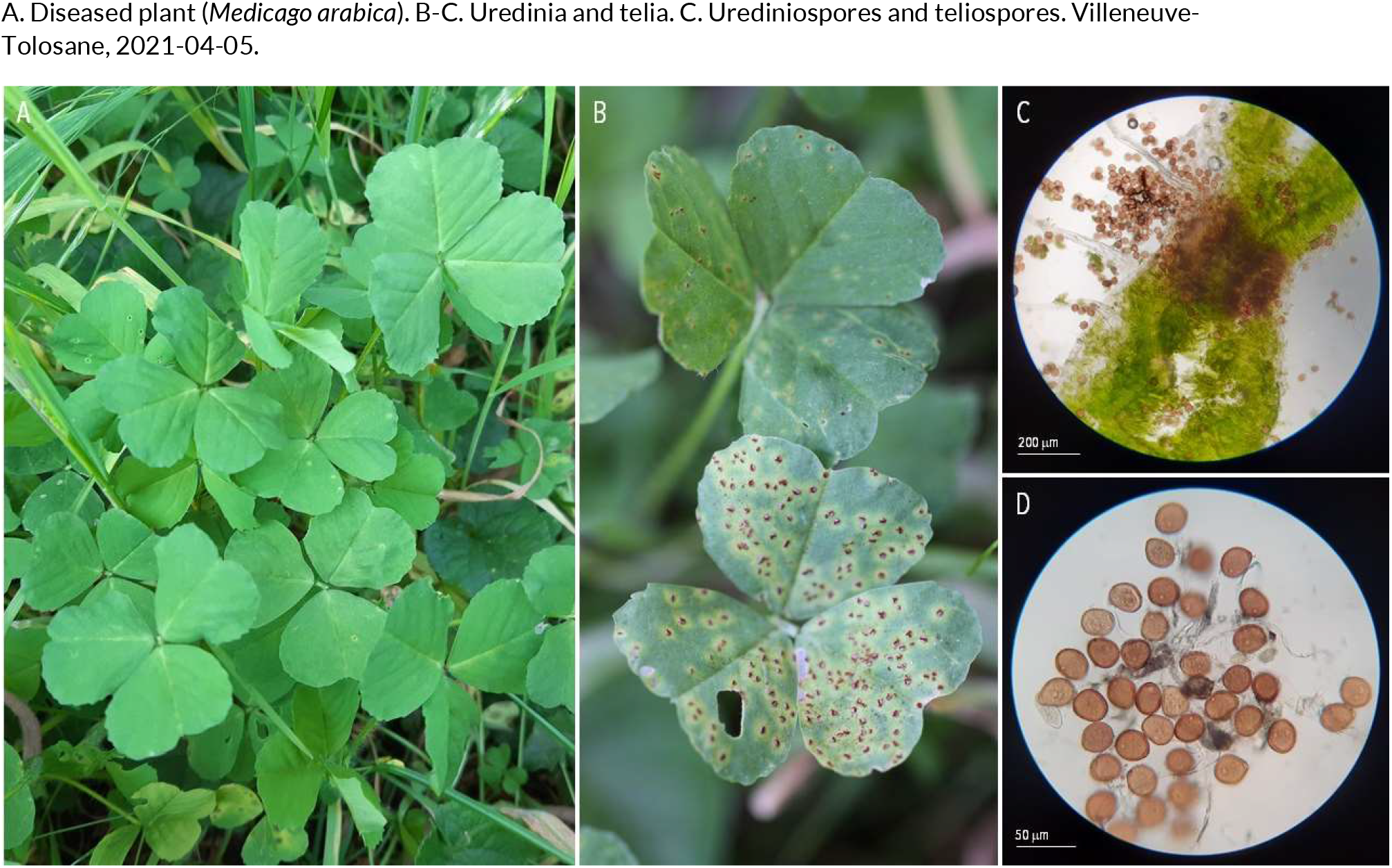
RUSTS | *Uromyces striatus* ex *Medicago sativa* (alfalfa) and *Medicago arabica* (spotted medick)

**96.**
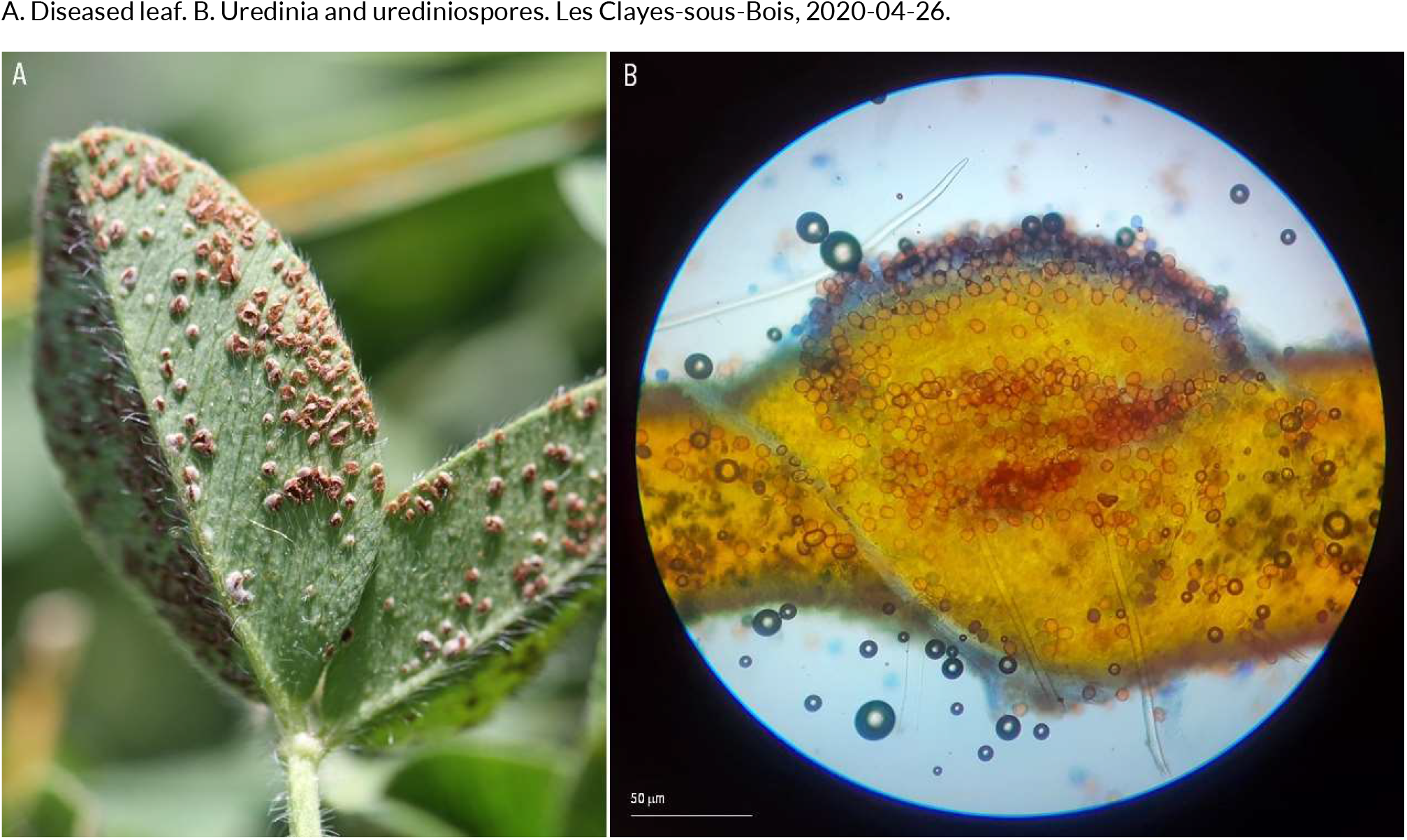
RUSTS | *Uromyces trifolii-repentis* ex *Trifolium pratense* (red clover)

**97.**
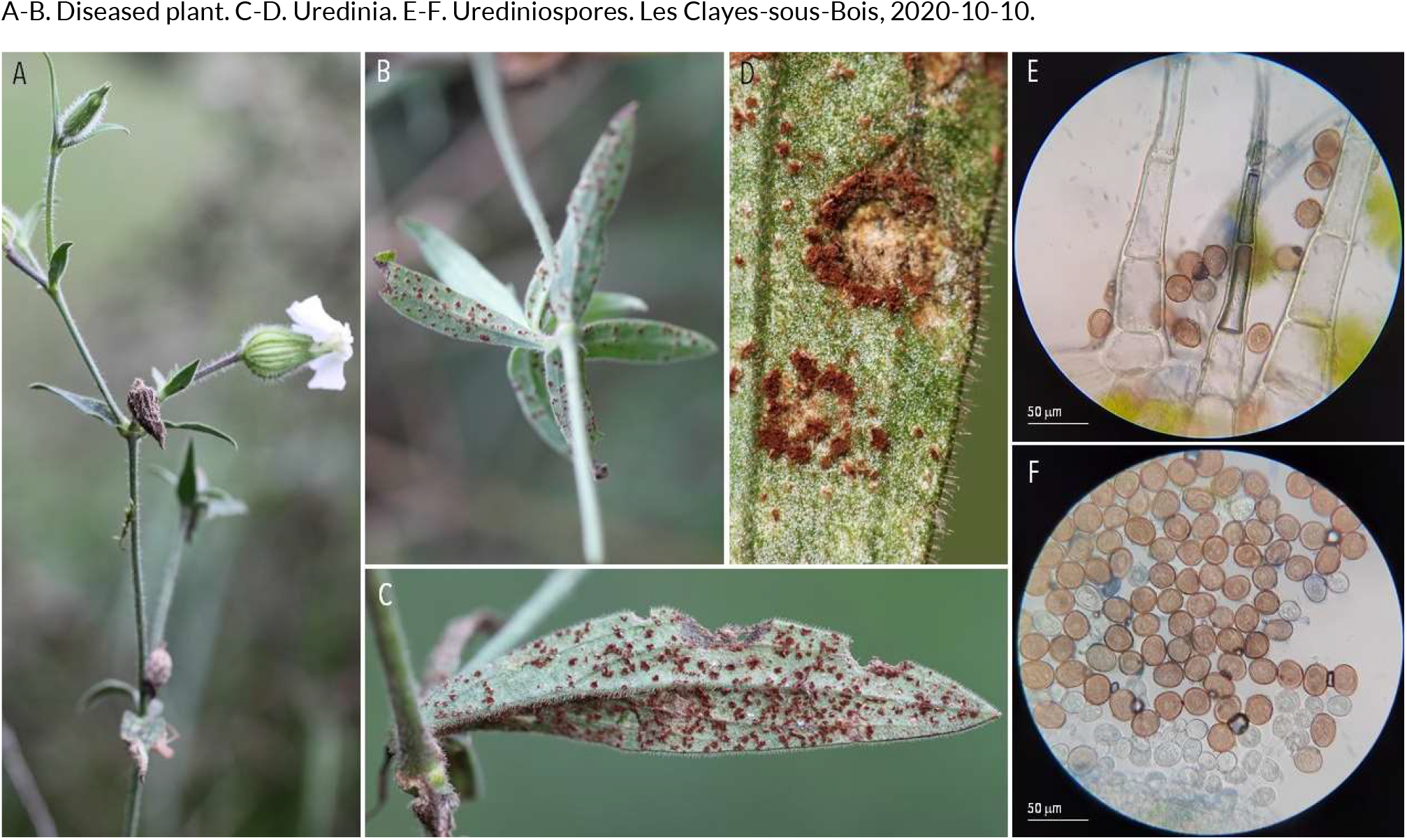
RUSTS | *Uromyces verruculosus* ex *Silene latifolia* (white campion)

**98.**
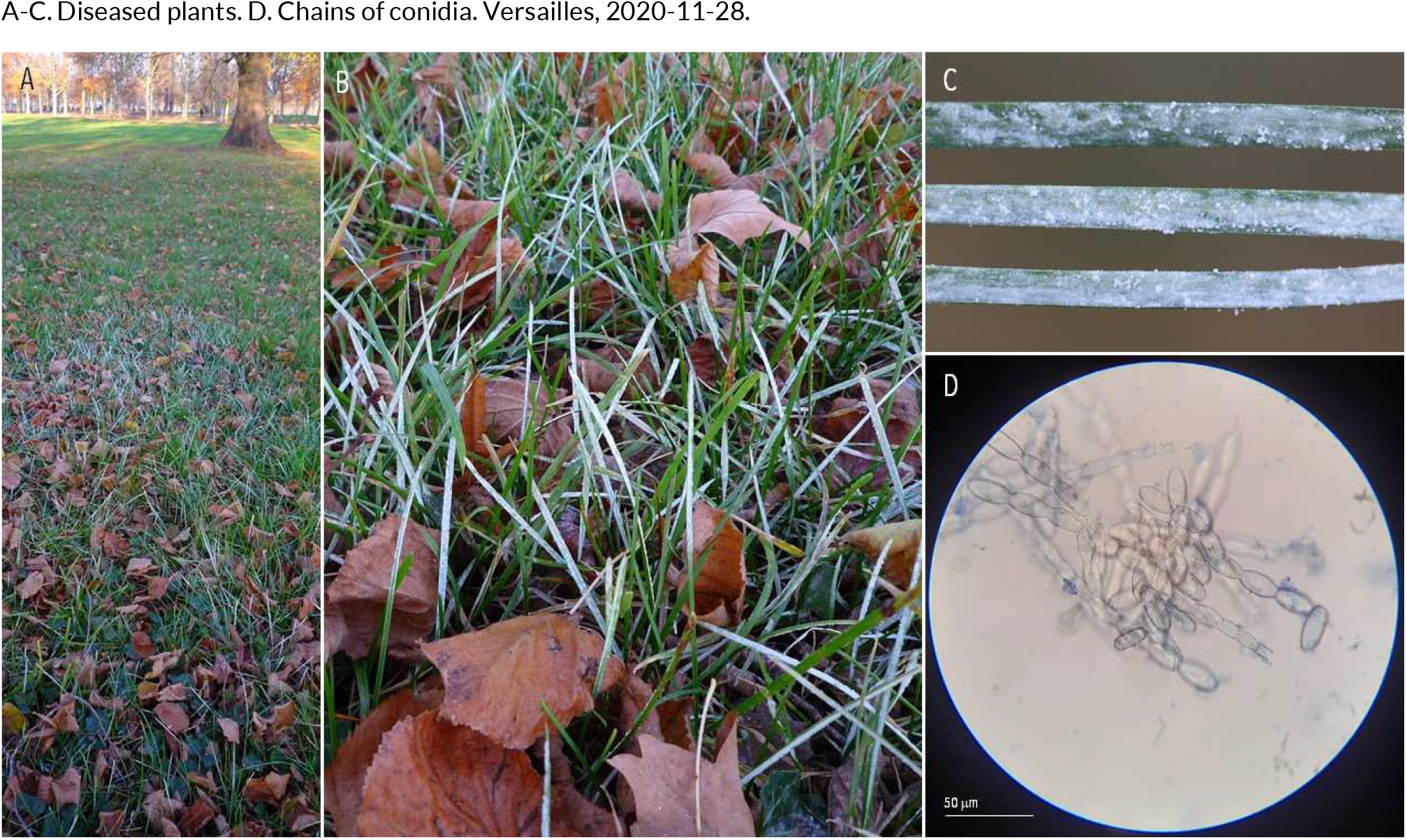
POWDERY MILDEW | *Blumeria graminis* sp. (??) ex *Poaceae sp*. (unidentified grass)

**99.**
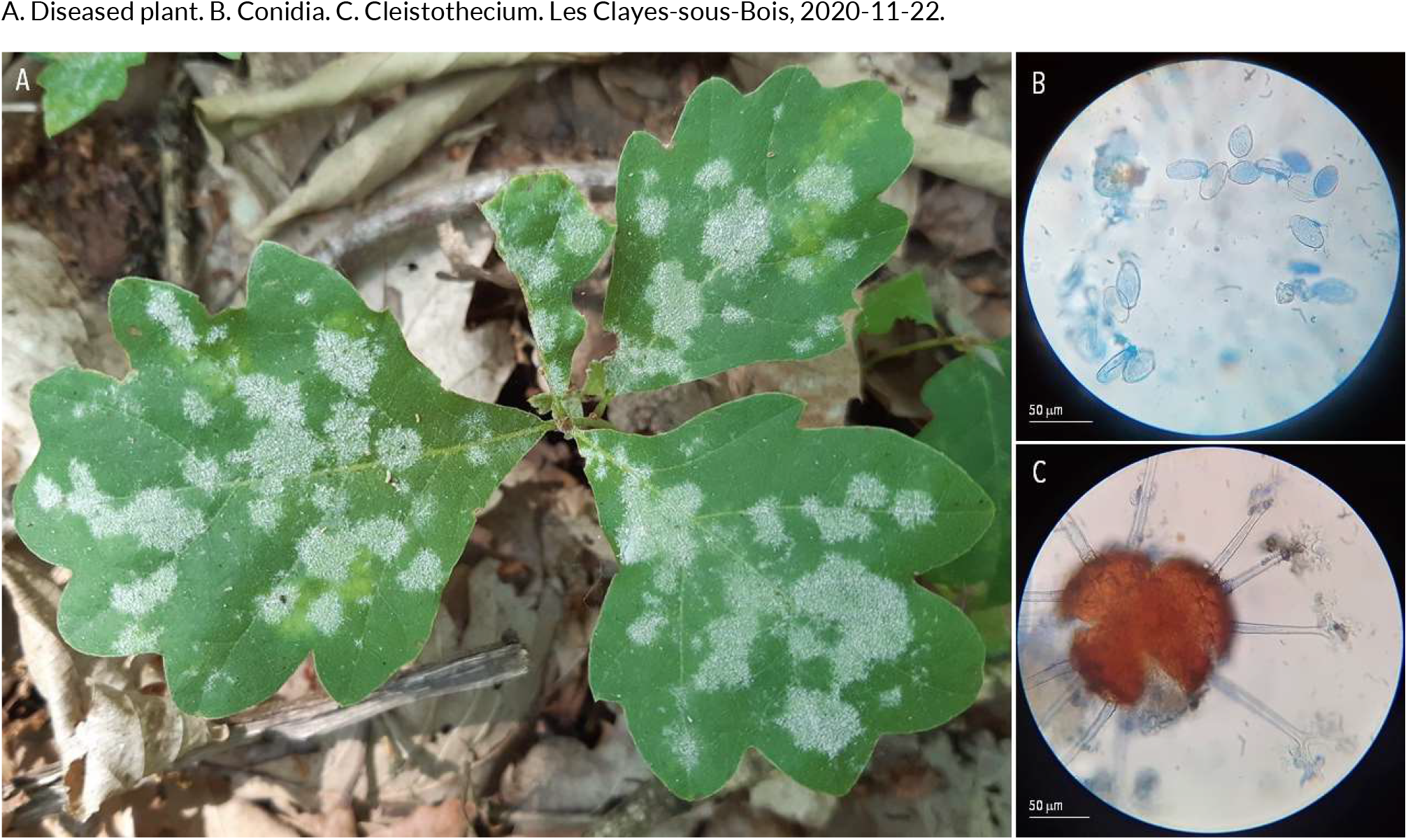
POWDERY MILDEW | *Erysiphe alphitoides* (?) ex *Quercus robur* (pedunculate oak)

**100.**
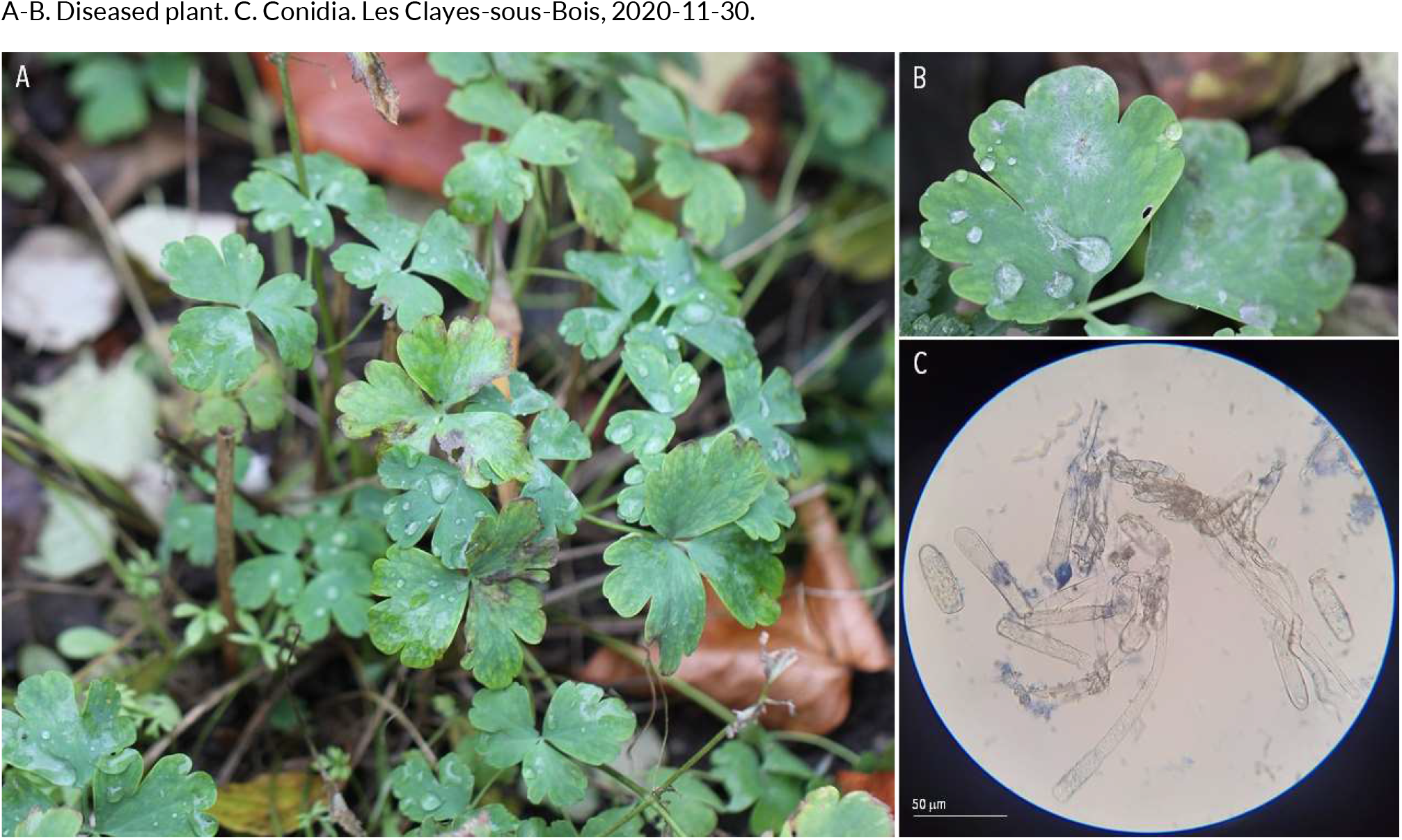
POWDERY MILDEW | *Erysiphe capreae* ex *Aquilegia vulgaris* (common columbine)

**101.**
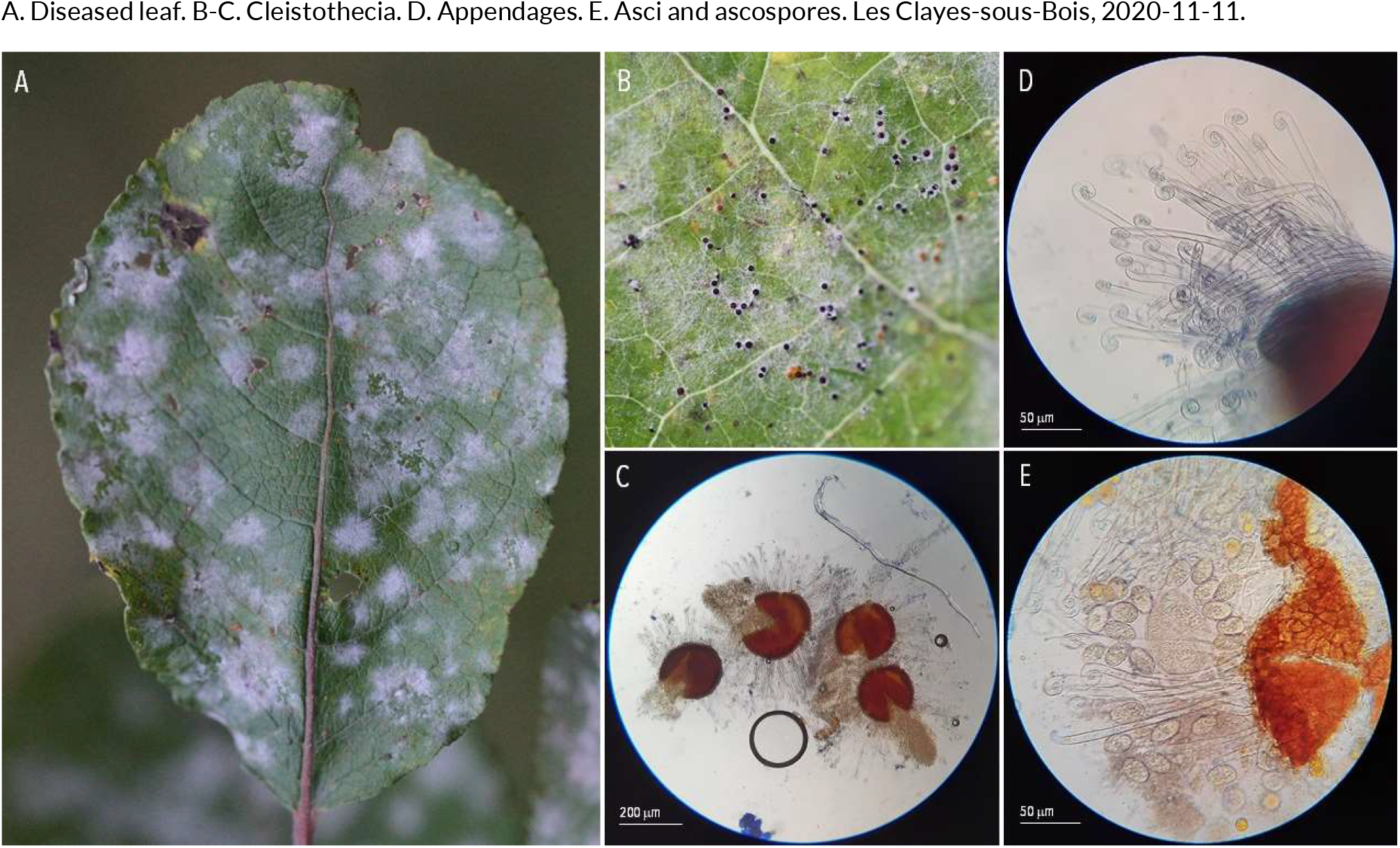
POWDERY MILDEW | *Erysiphe capreae* ex *Salix caprea* (goat willow)

**102.**
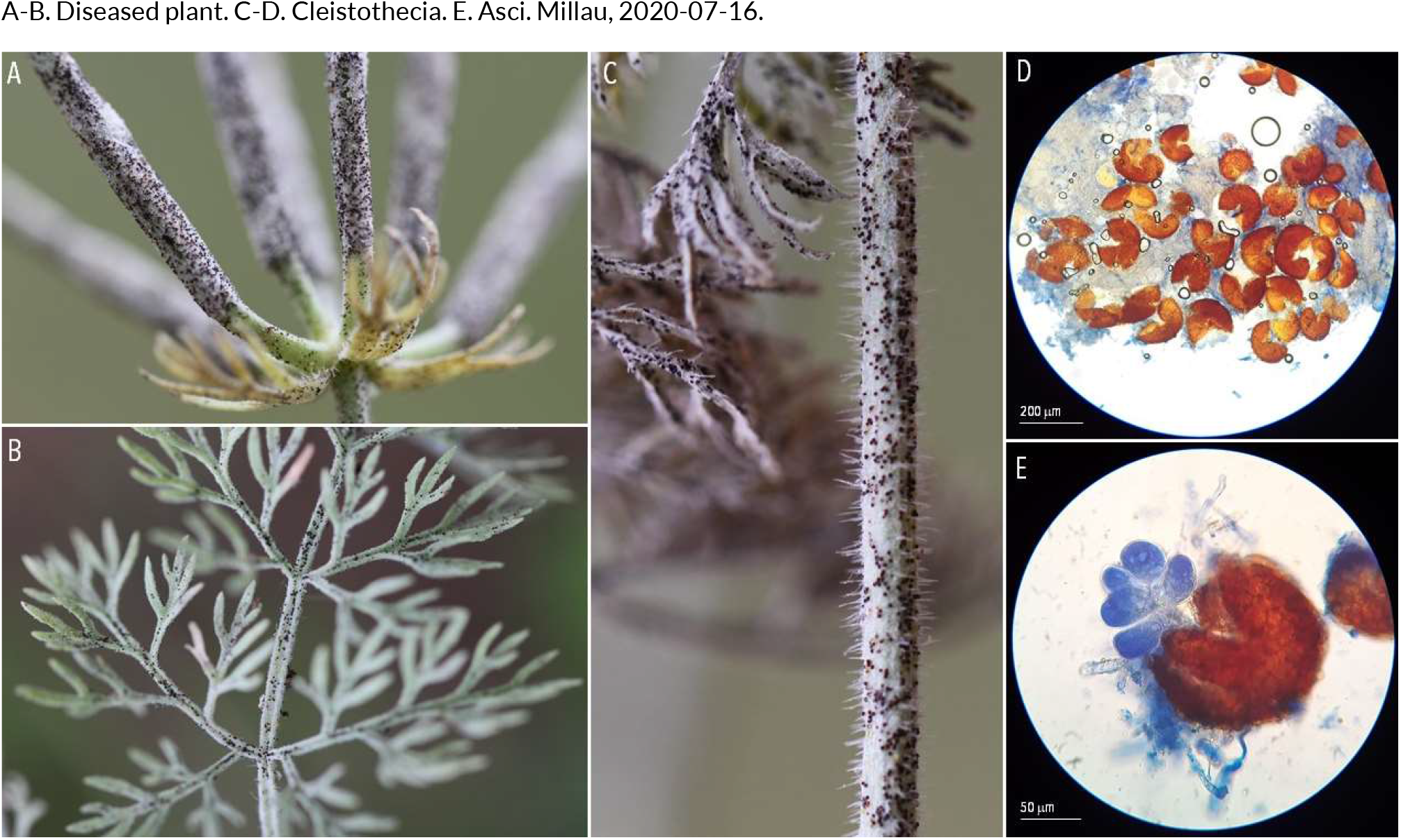
POWDERY MILDEW | *Erysiphe cruciferarum* ex *Eschscholzia californica* (California poppy)

**103.**
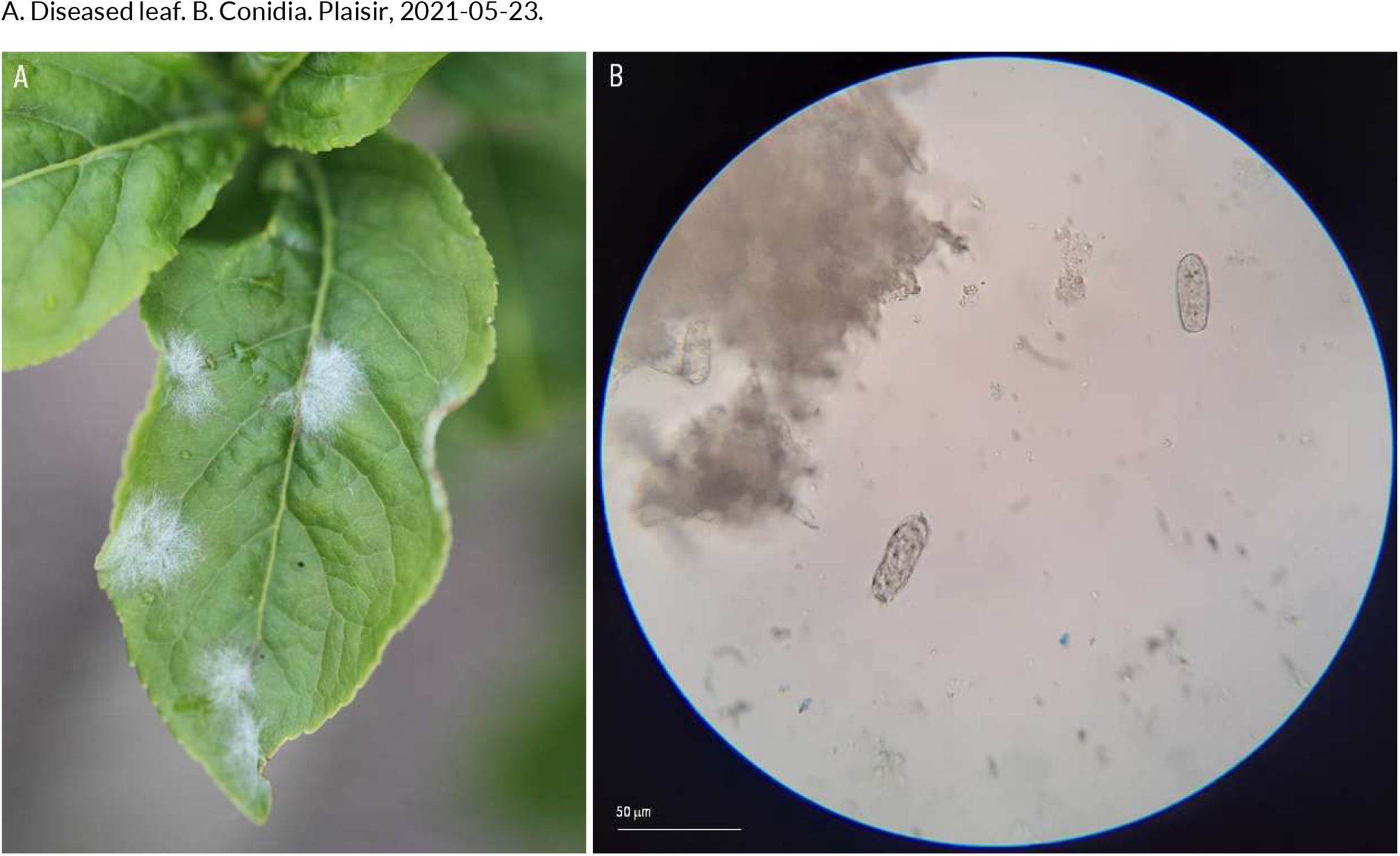
POWDERY MILDEW | *Erysiphe euonymi* ex *Euonymus europaeus* (common spindle)

**104.**
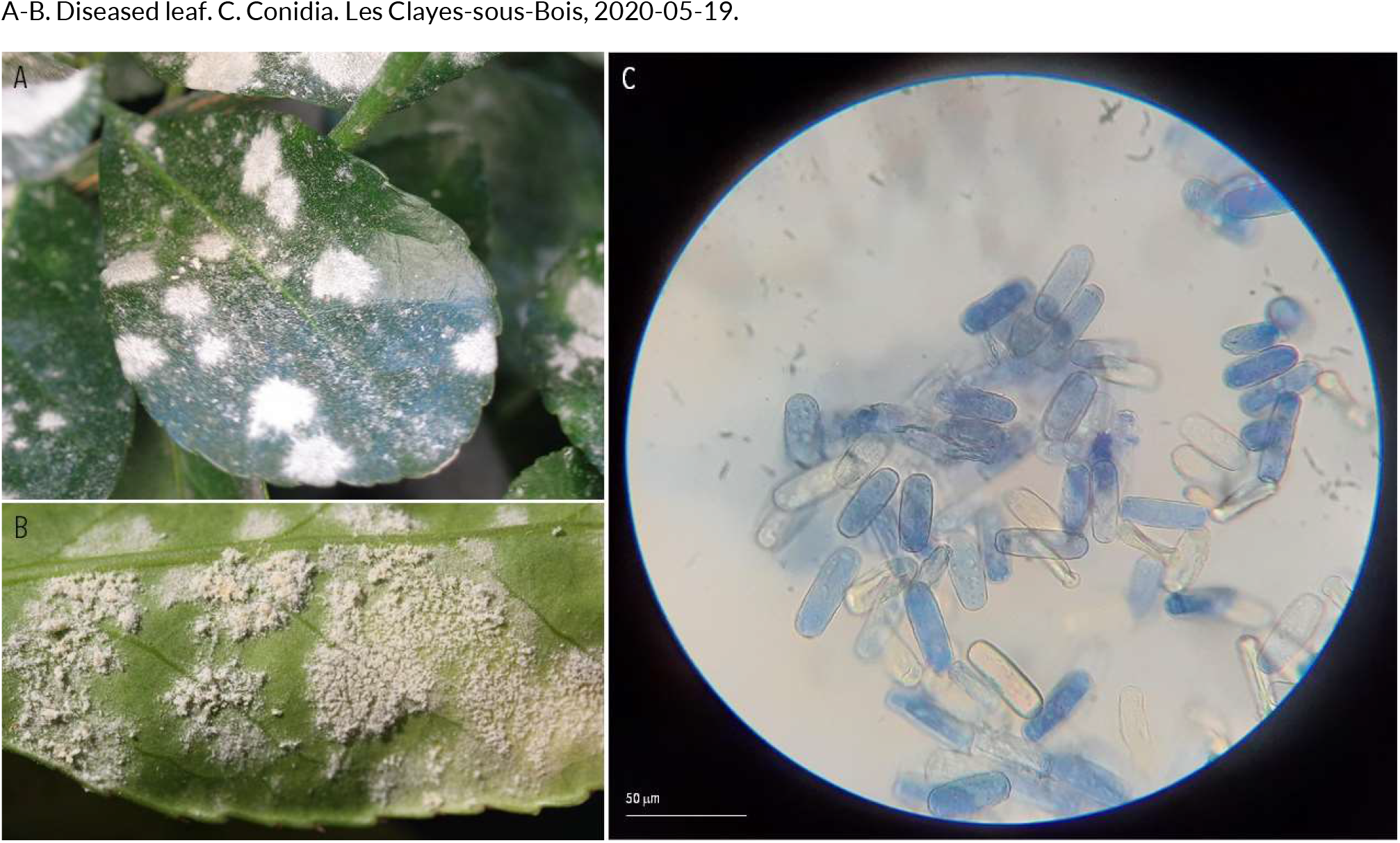
POWDERY MILDEW | *Erysiphe euonymi-japonici* ex *Euonymus japonicus* (Japanese spindle tree)

**105.**
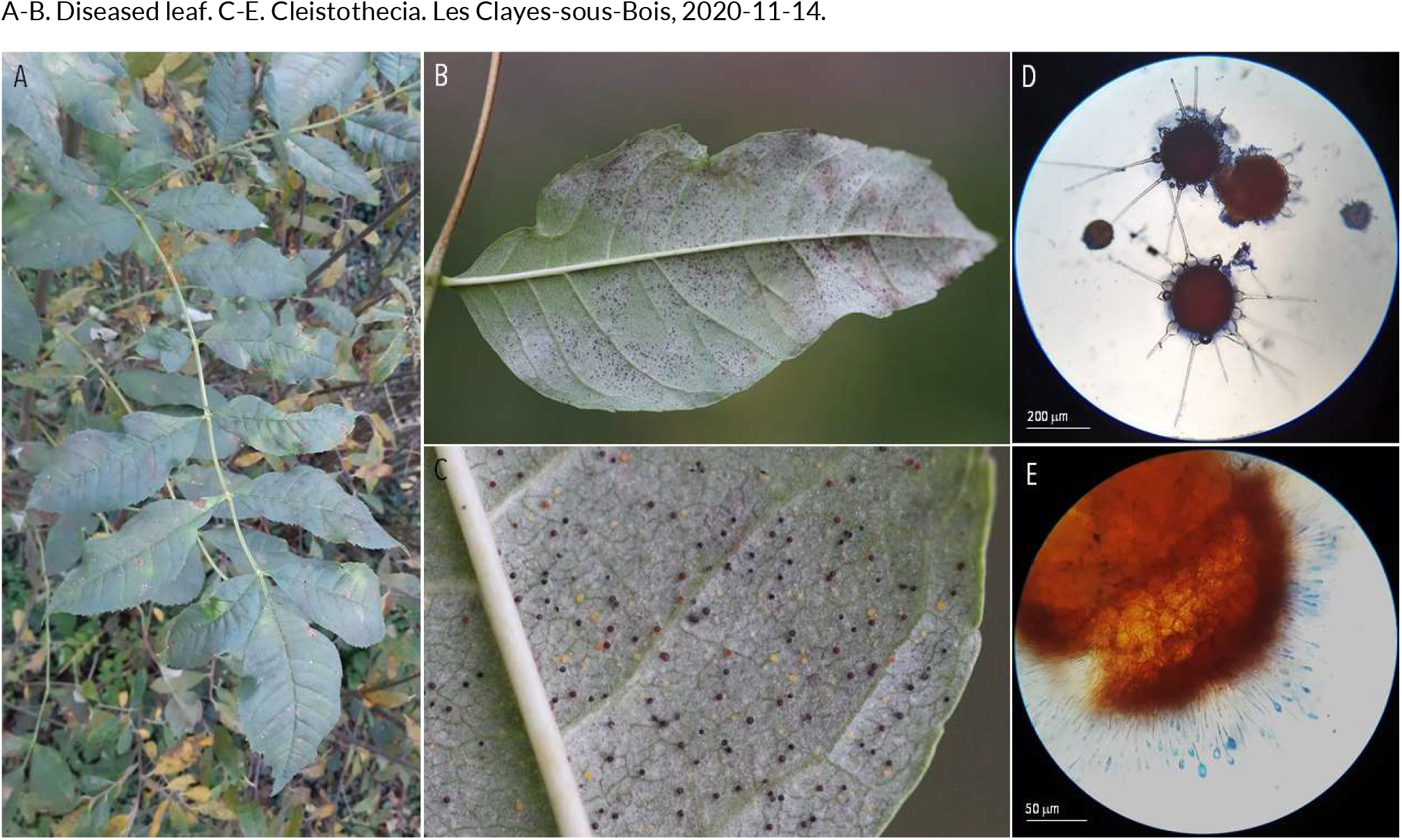
POWDERY MILDEW | *Erysiphe fraxini* (syn. *Phyllactinia fraxini*) ex *Fraxinus excelsior* (common ash)

**106.**
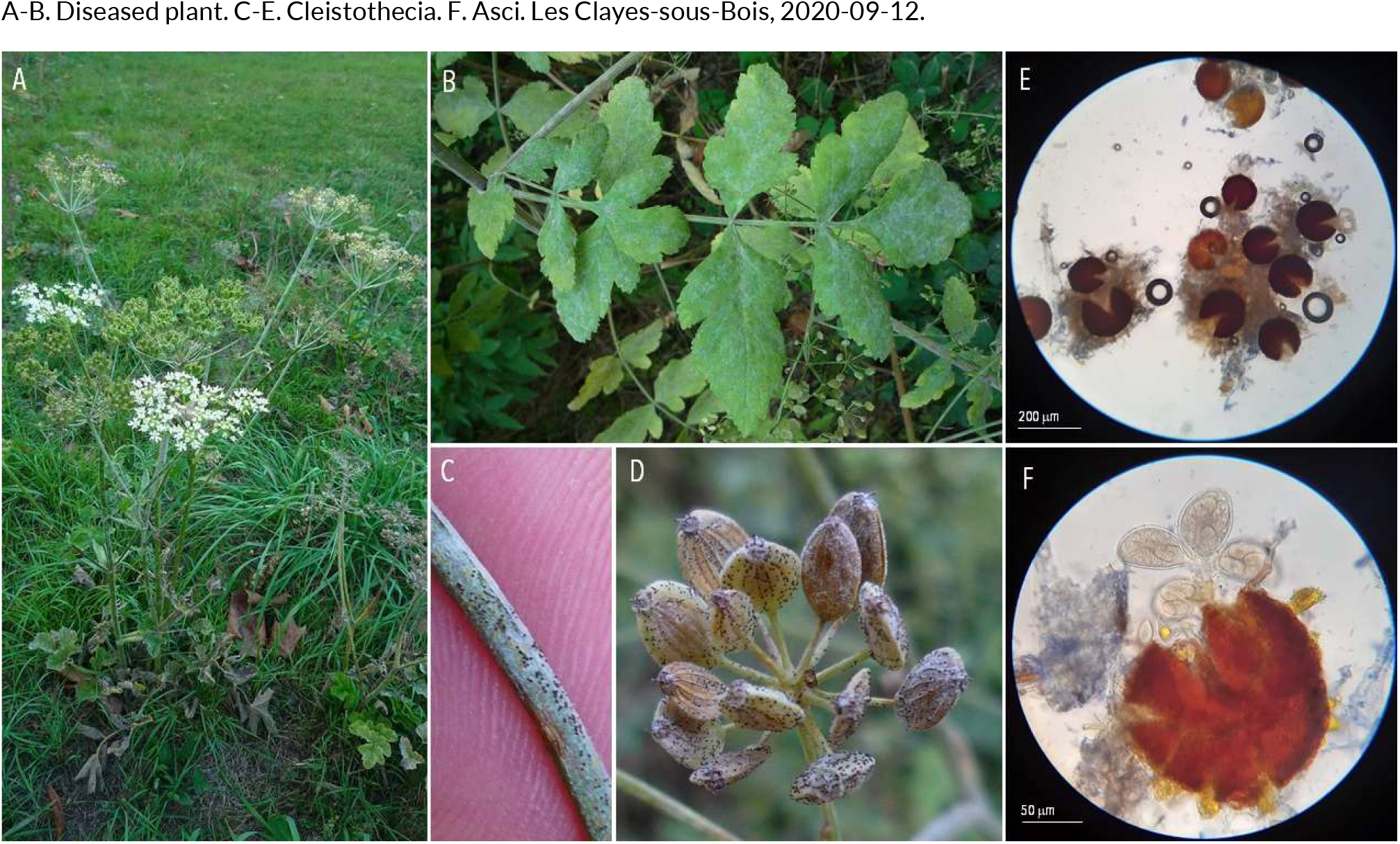
POWDERY MILDEW | *Erysiphe heraclei* ex *Heracleum sphondylium* (common hogweed)

**107.**
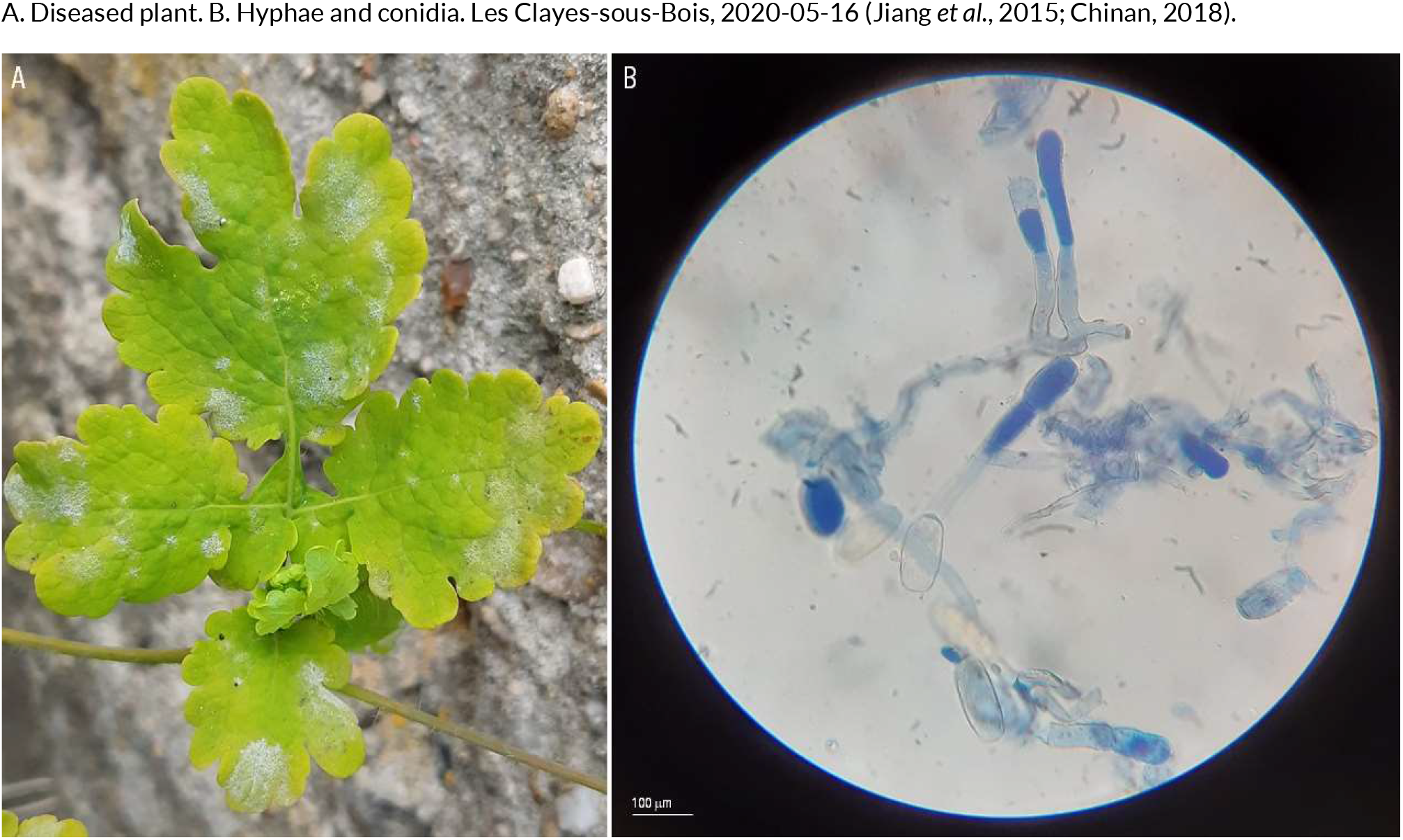
POWDERY MILDEW | *Erysiphe macleayae* ex *Chelidonium majus* (greater celandine)

**108.**
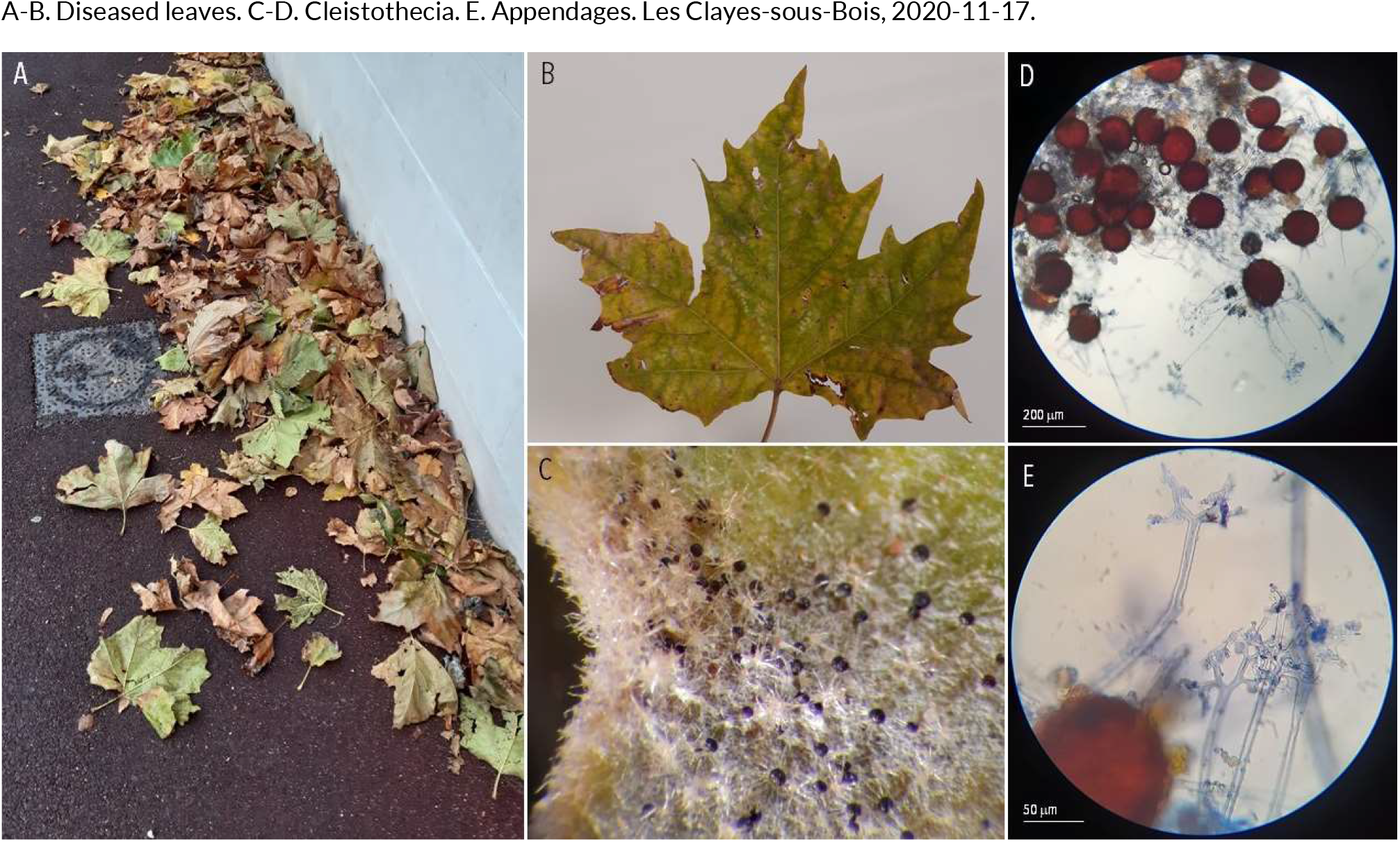
POWDERY MILDEW | *Erysiphe platani* ex *Platanus x hispanica* (London plane)

**109.**
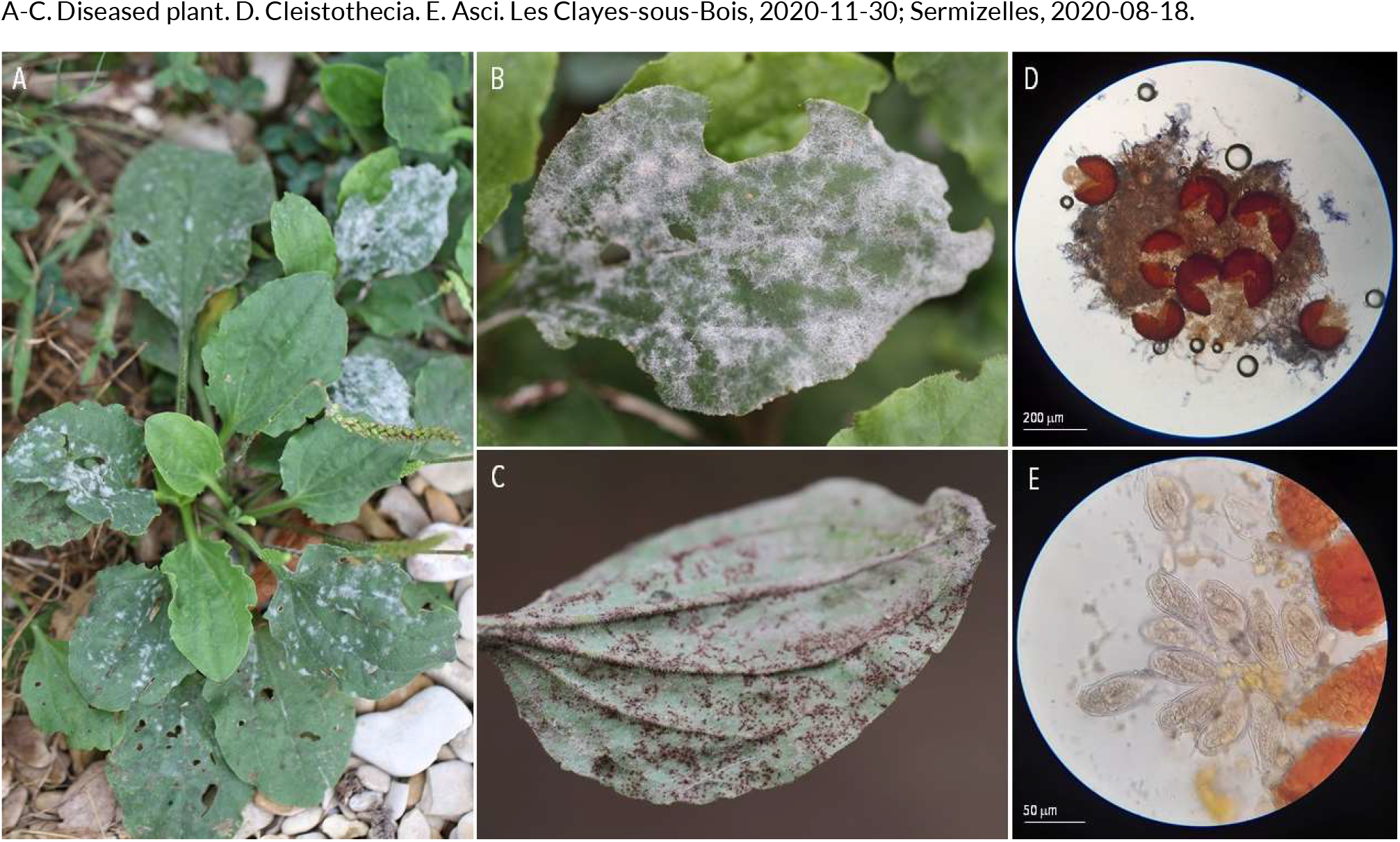
POWDERY MILDEW | *Erysiphe sordida* ex *Plantago major* (broadleaf plantain)

**110.**
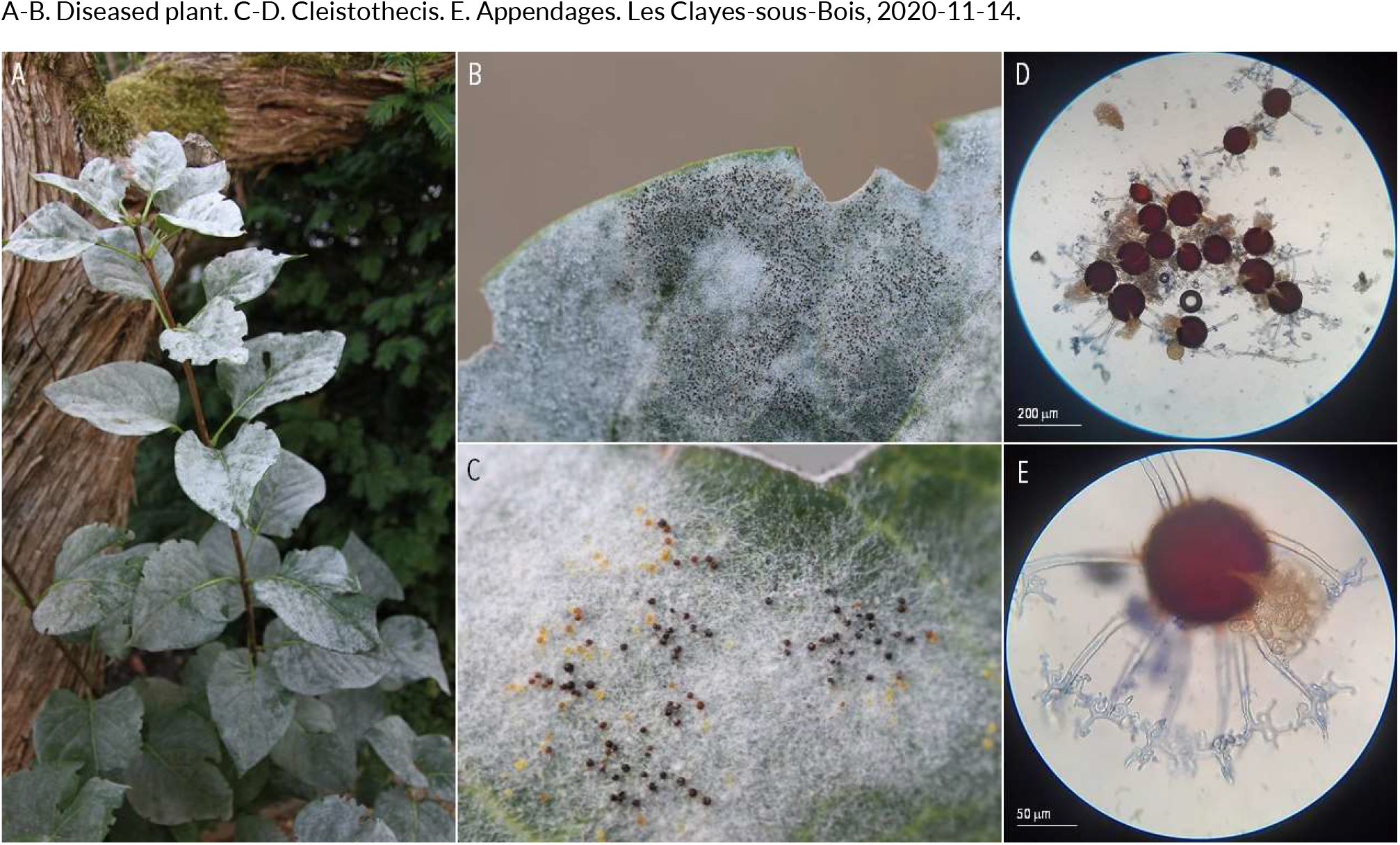
POWDERY MILDEW | *Erysiphe syringae-japonicae* ex *Syringa vulgaris* (common lilac)

**111.**
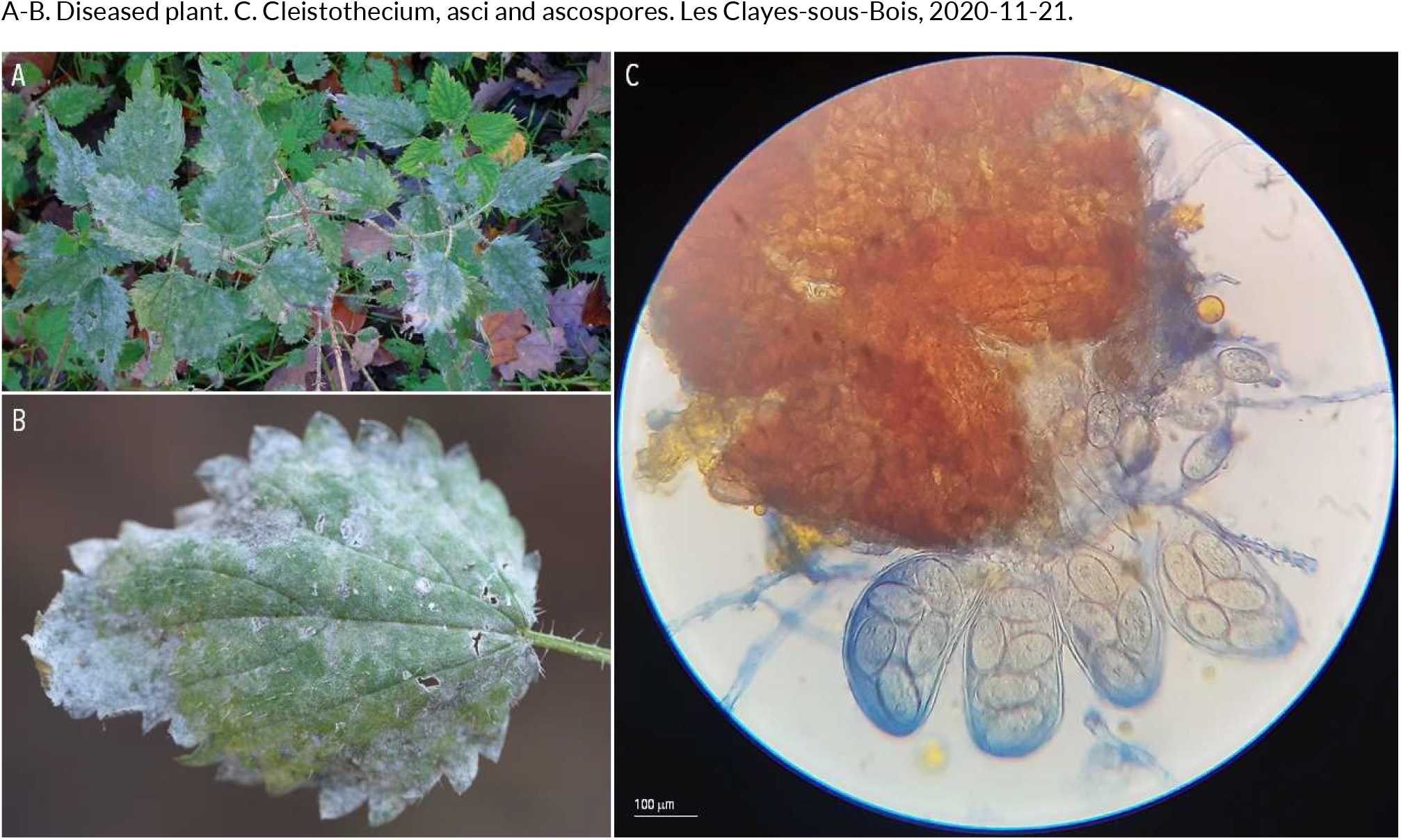
POWDERY MILDEW | *Erysiphe urticae* ex *Urtica dioica* (nettle)

**112.**
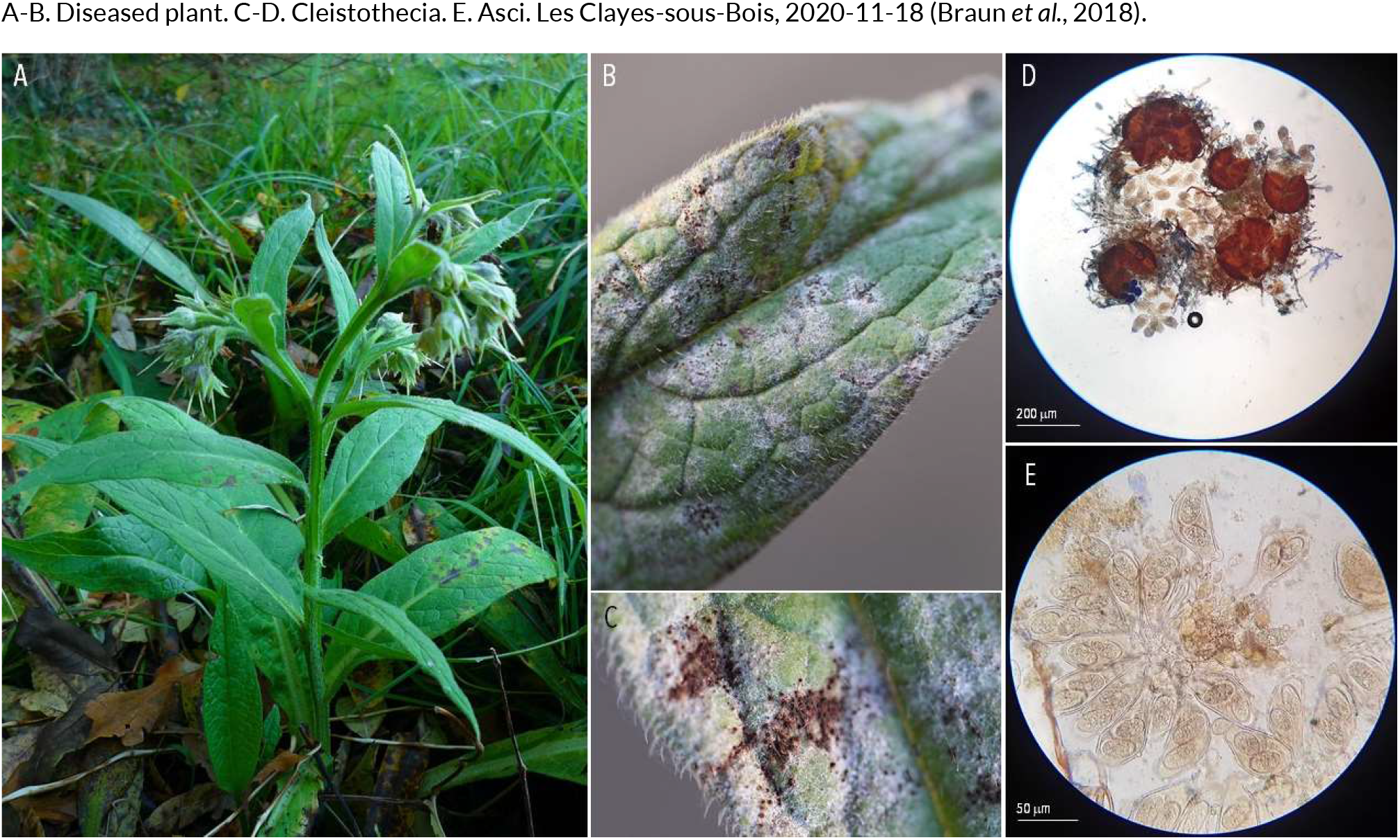
POWDERY MILDEW | *Golovinomyces asperifoliorum* (syn. Erysiphe asperifoliorum) ex *Symphytum officinale* (consound)

**113.**
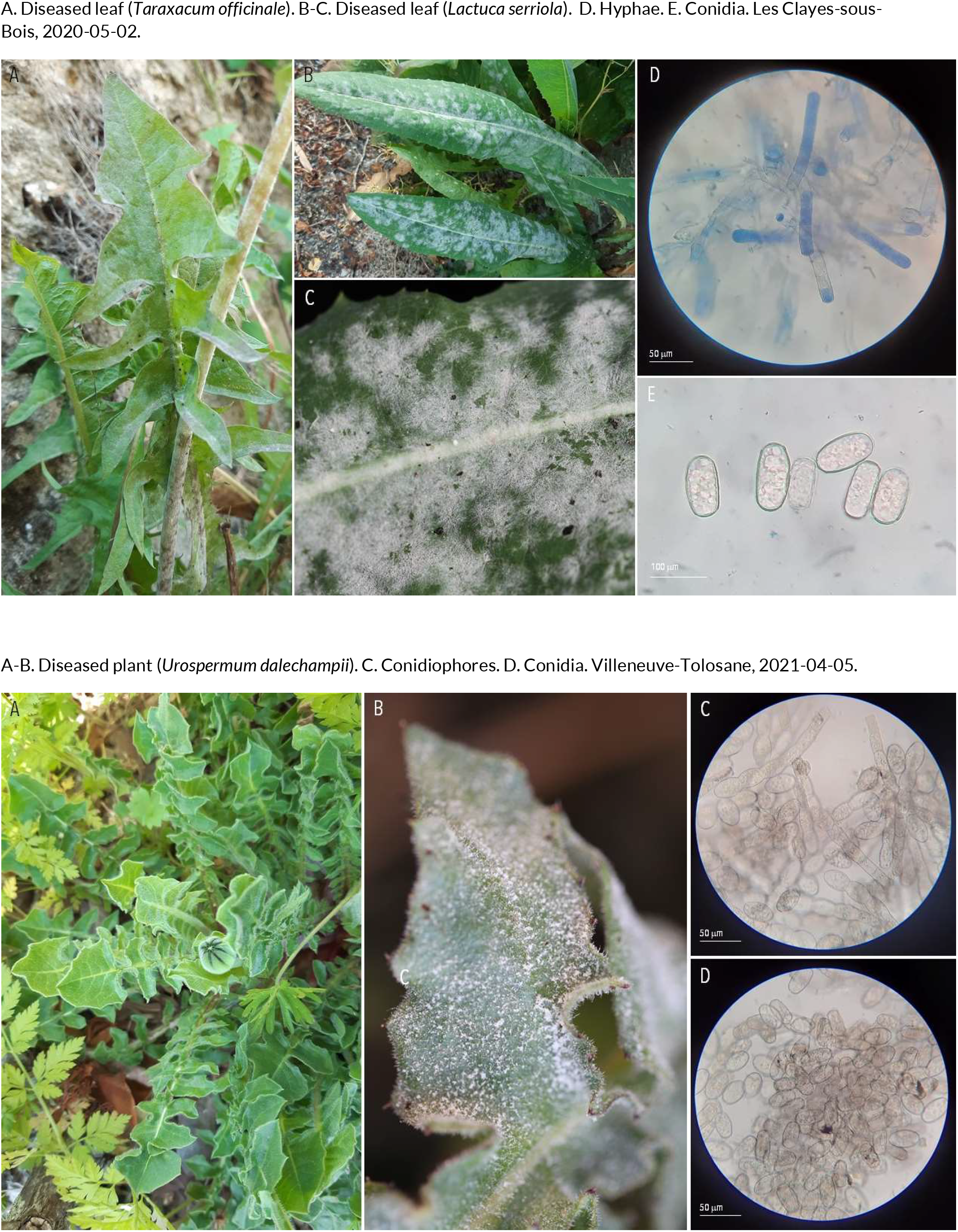
POWDERY MILDEW | *Golovinomyces cichoracearum* (syn. Erysiphe cichoracearum) ex *Taraxacum officinale* (common dandelion), *Lactuca serriola* (wild lettuce) and *Urospermum dalechampii* (smooth golden fleece)

**114.**
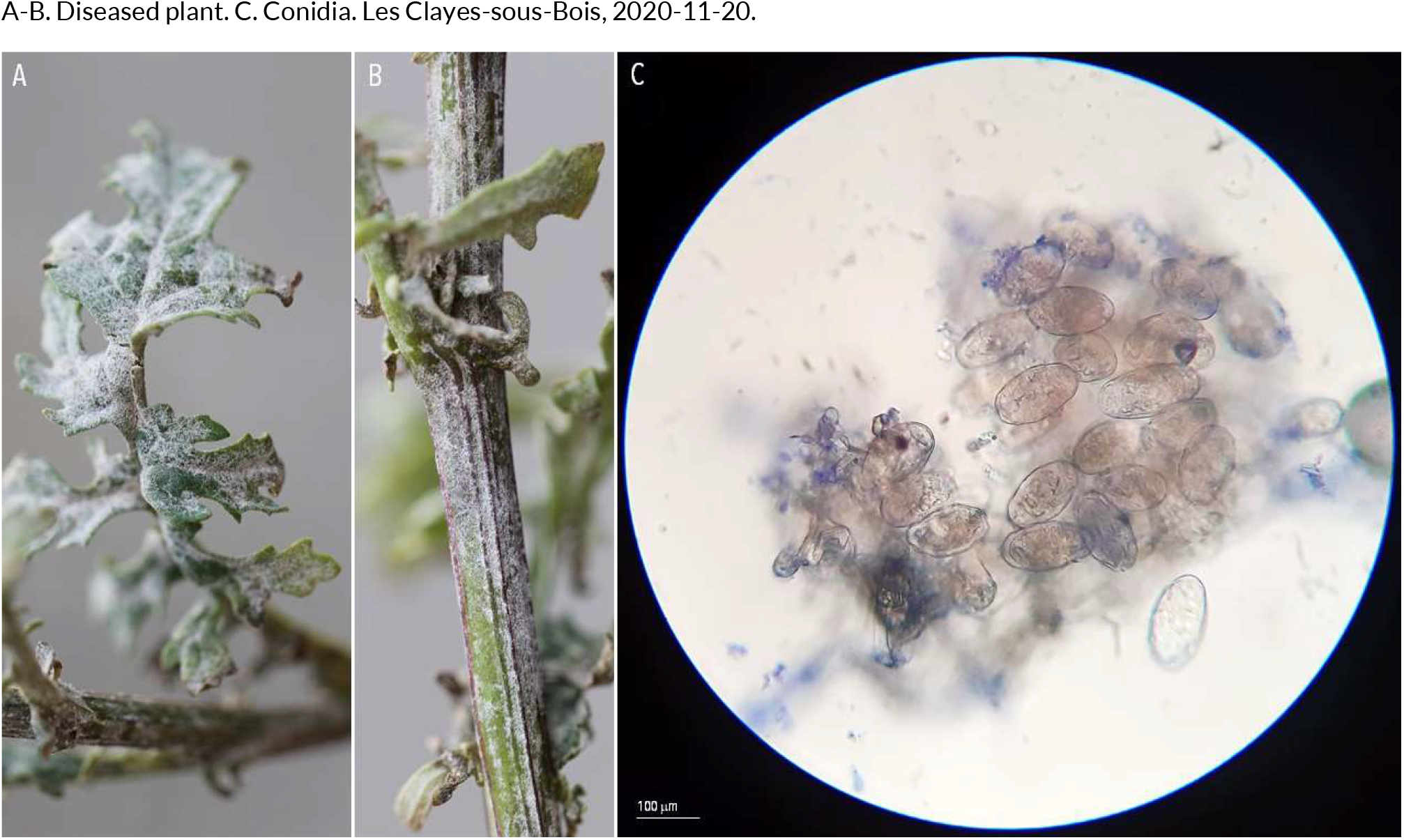
POWDERY MILDEW | *Golovinomyces fischeri* (?) (syn. Erysiphe fischeri) ex *Senecio vulgaris* (common groundsel)

**115.**
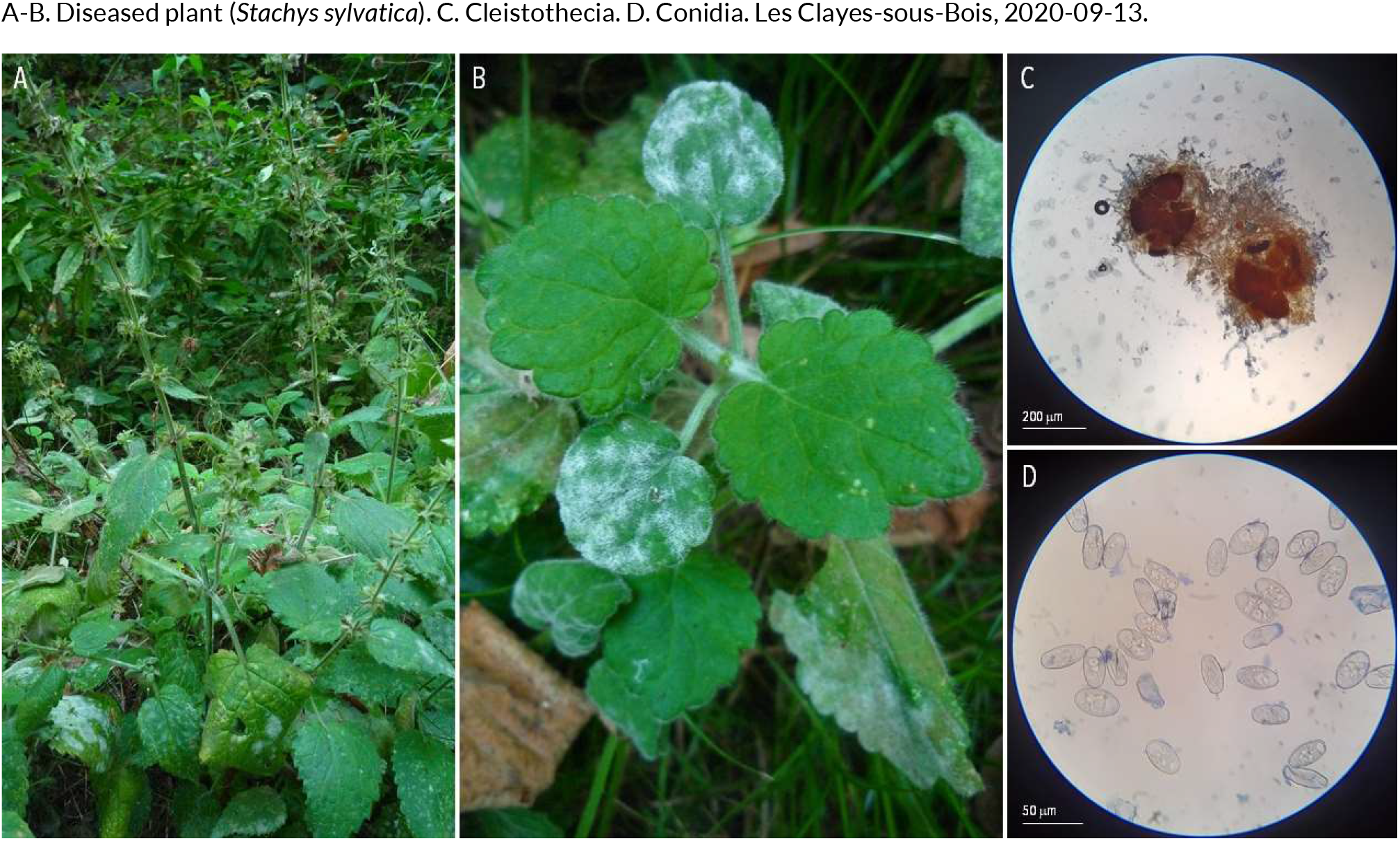
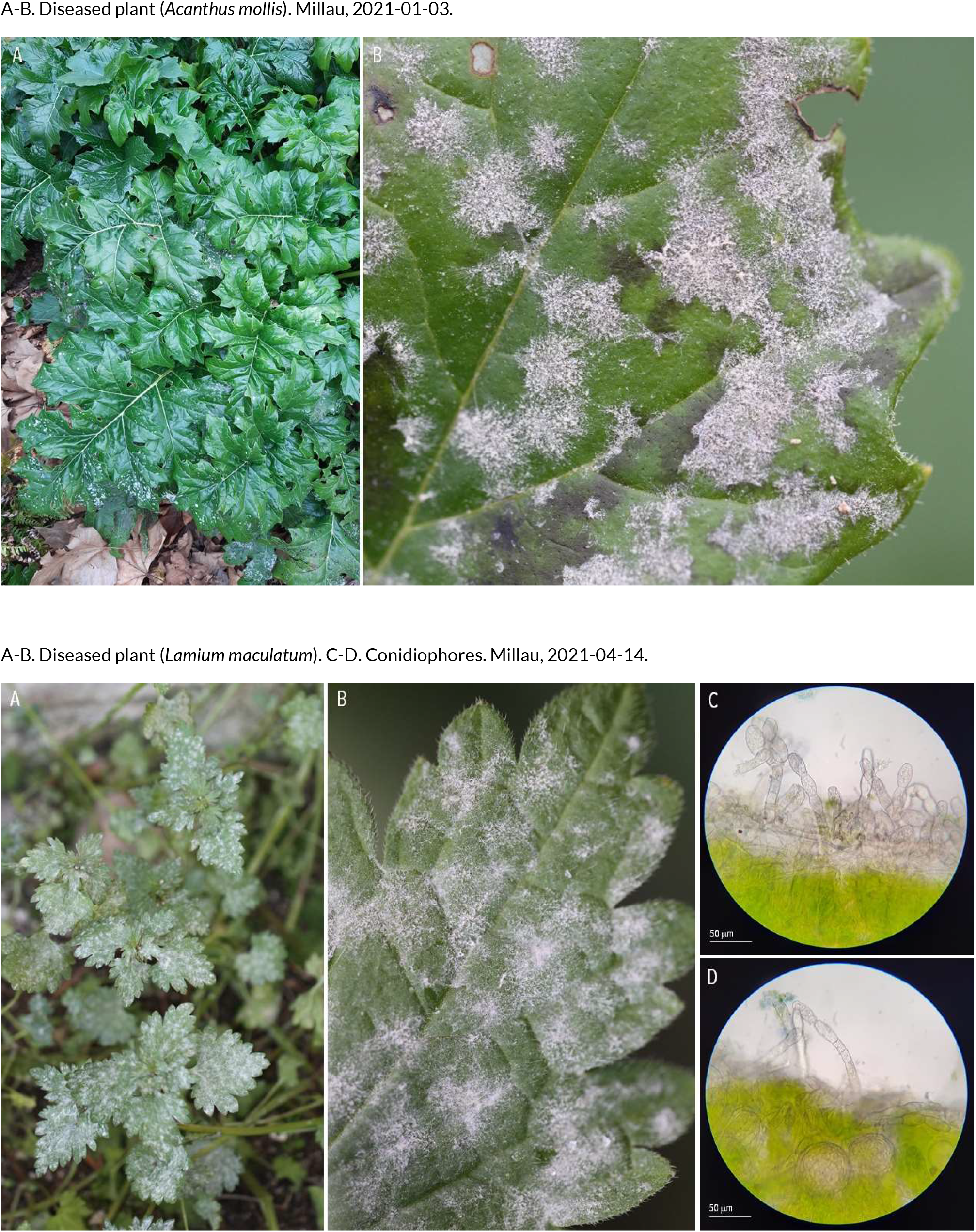
POWDERY MILDEW | *Neoerysiphe galeopsidis* (?) ex *Stachys sylvatica* (hedge woundwort), *Acanthus mollis* (sea-dock) and *Lamium maculatum* (spotted dead-nettle)

**116.**
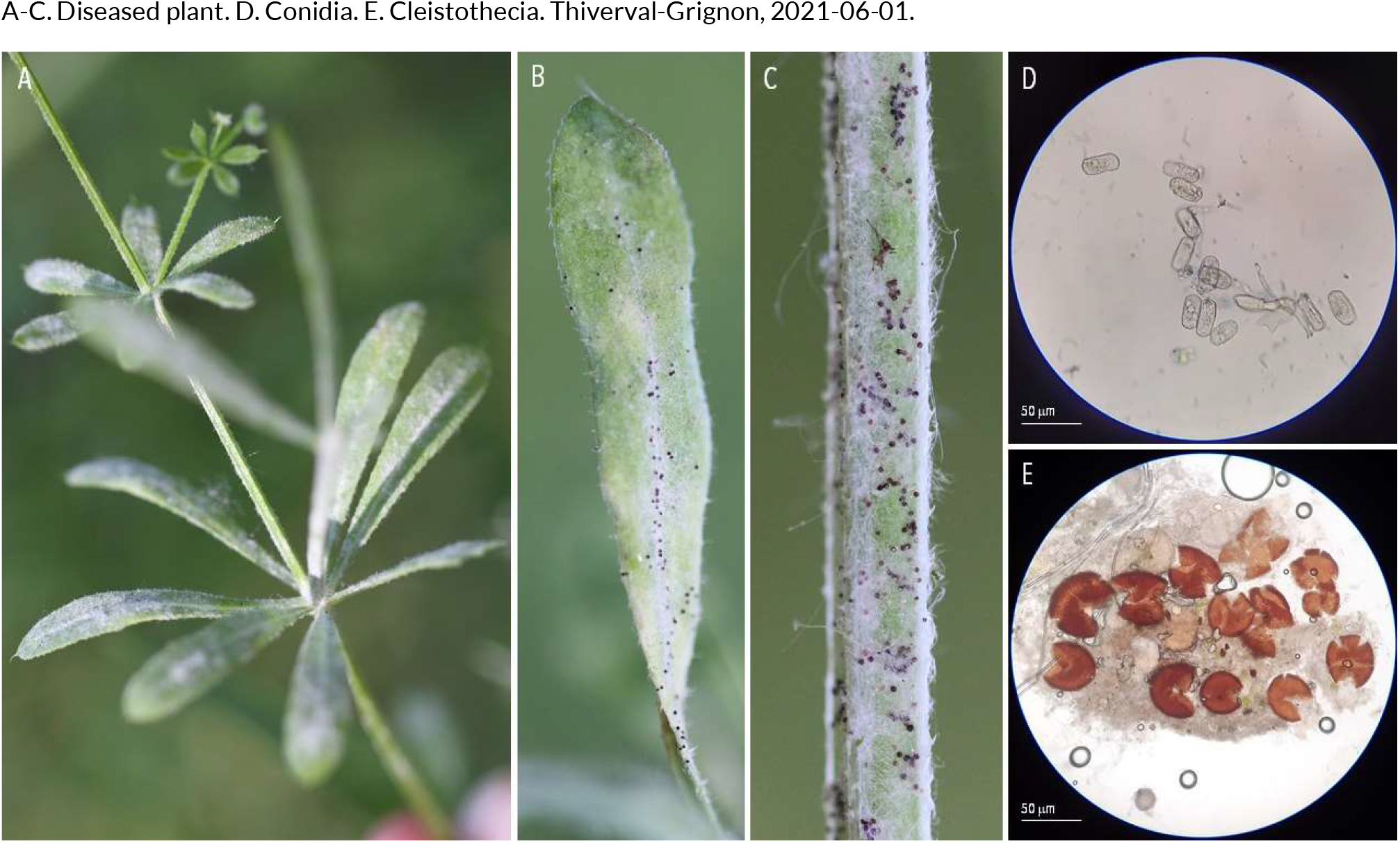
POWDERY MILDEW | *Neoerysiphe galii* ex *Galium aparine* (catchweed bedstraw)

**117.**
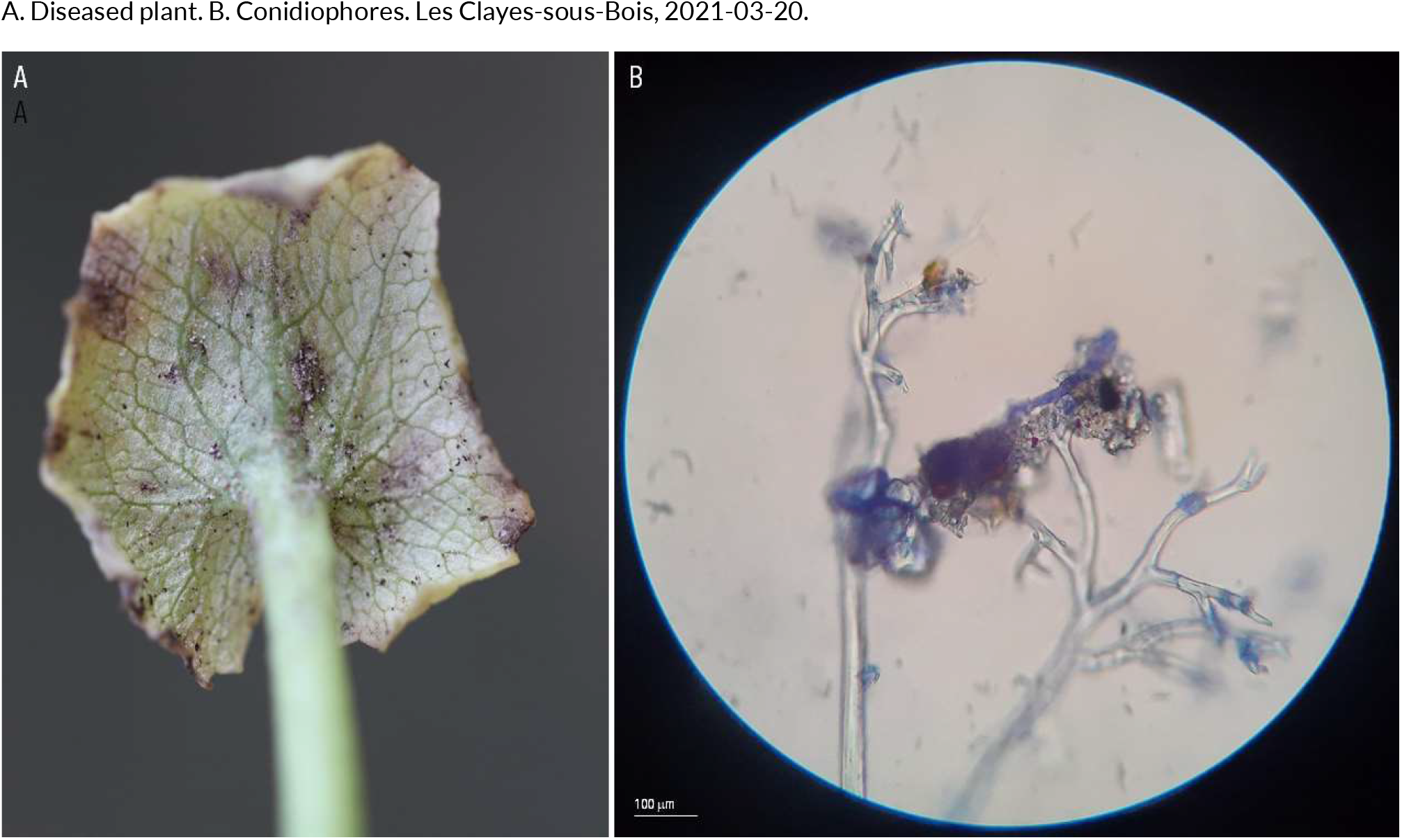
POWDERY MILDEW | *Peronospora ficaria* ex *Ficaria verna* (lesser celandine)

**118.**
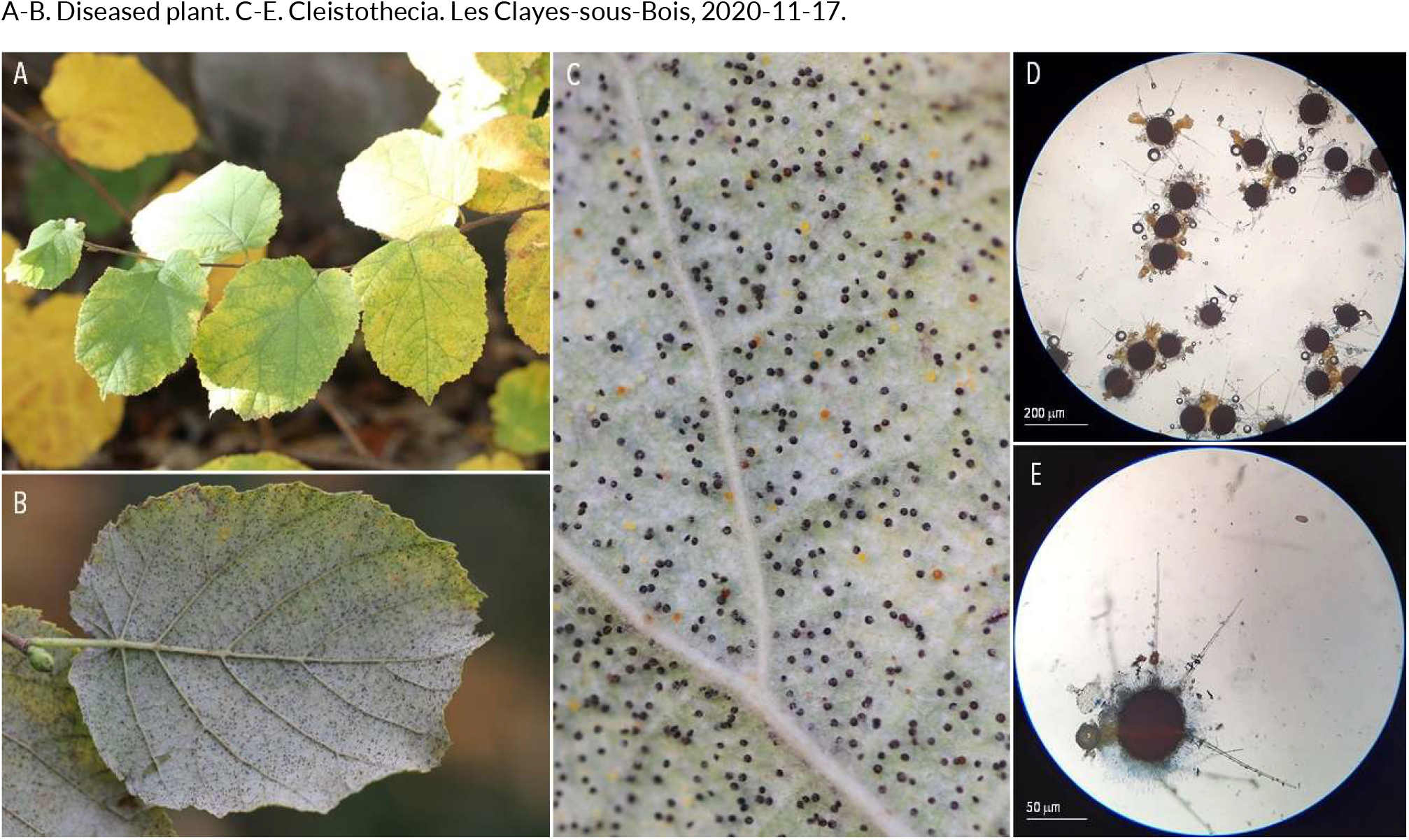
POWDERY MILDEW | *Phyllactinia guttata* (syn. *Erysiphe coryli*) ex *Corylus avellana* (common hazel)

**119.**
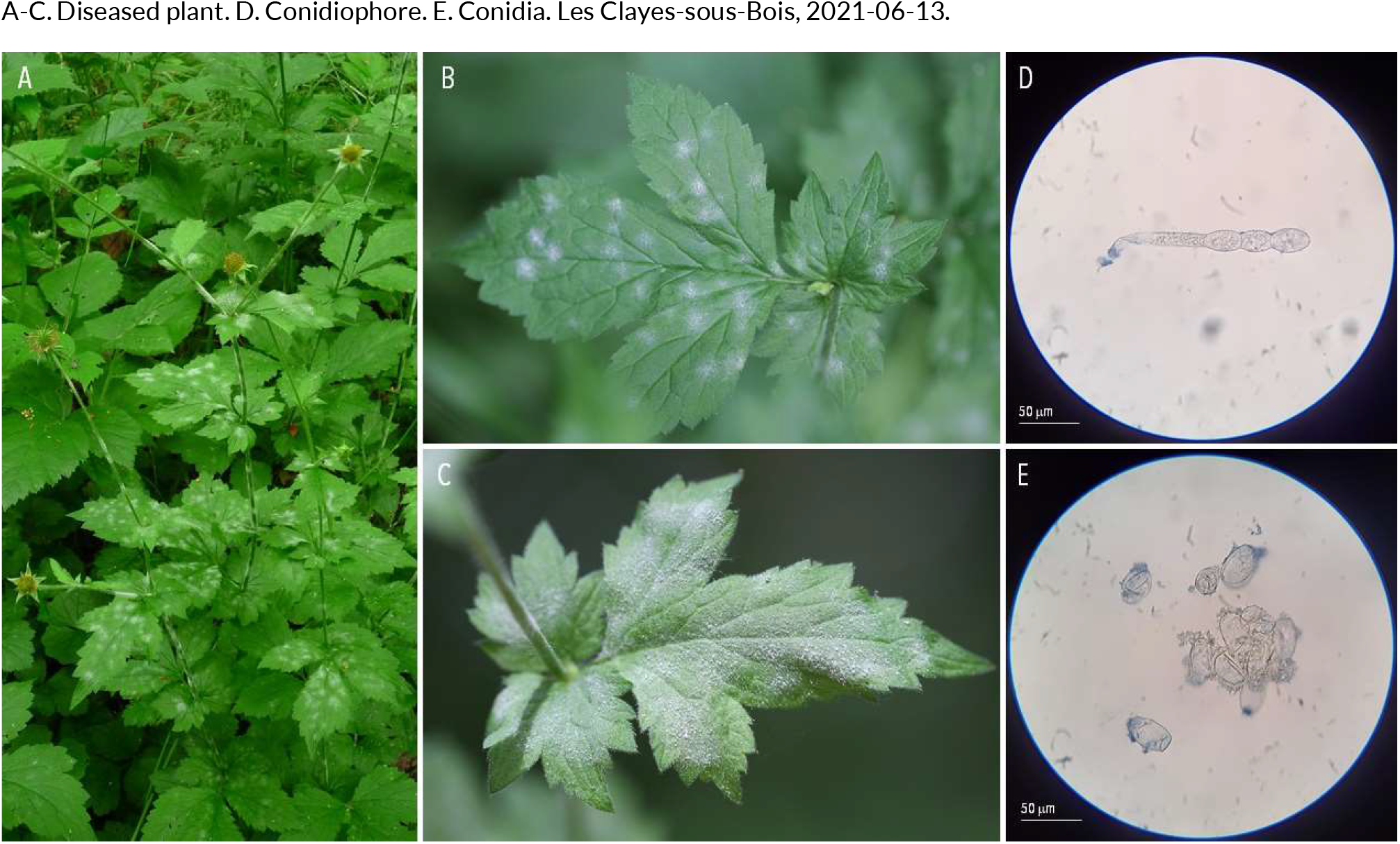
POWDERY MILDEW | *Podosphaera aphanis* ex *Geum urbanum* (wood avens)

**120.**
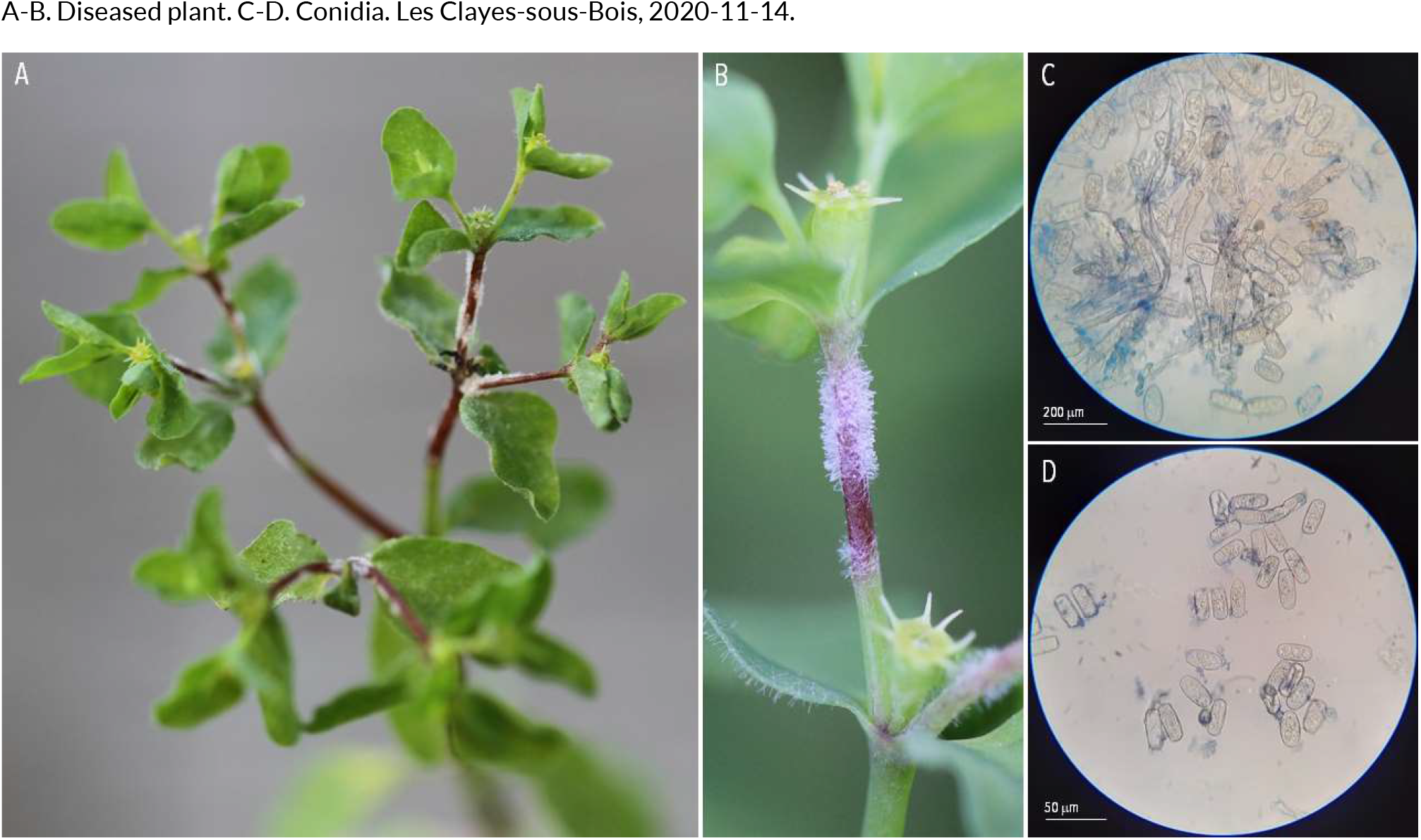
POWDERY MILDEW | *Podosphaera euphorbiae* ex *Euphorbia peplus* (petty spurge)

**121.**
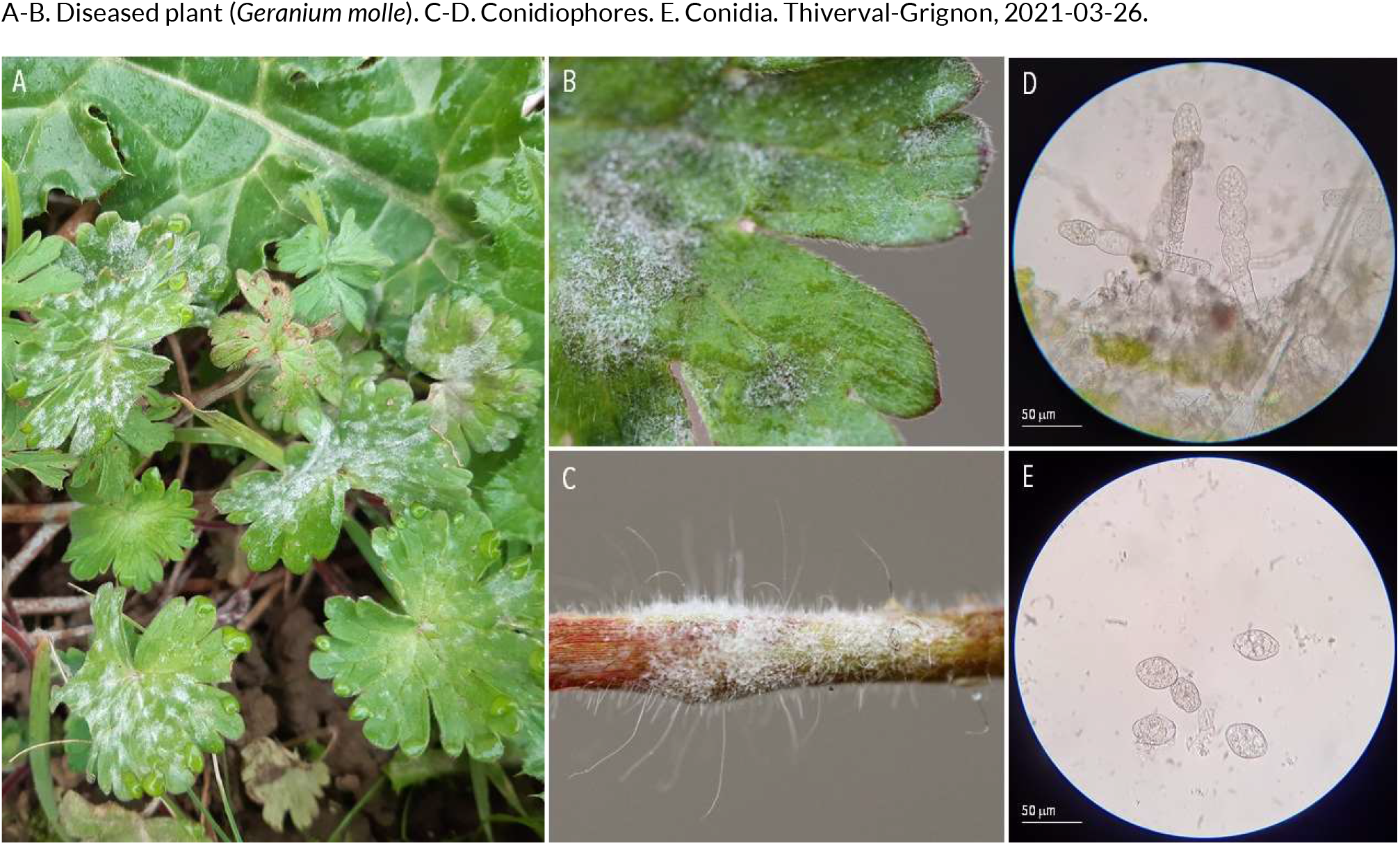
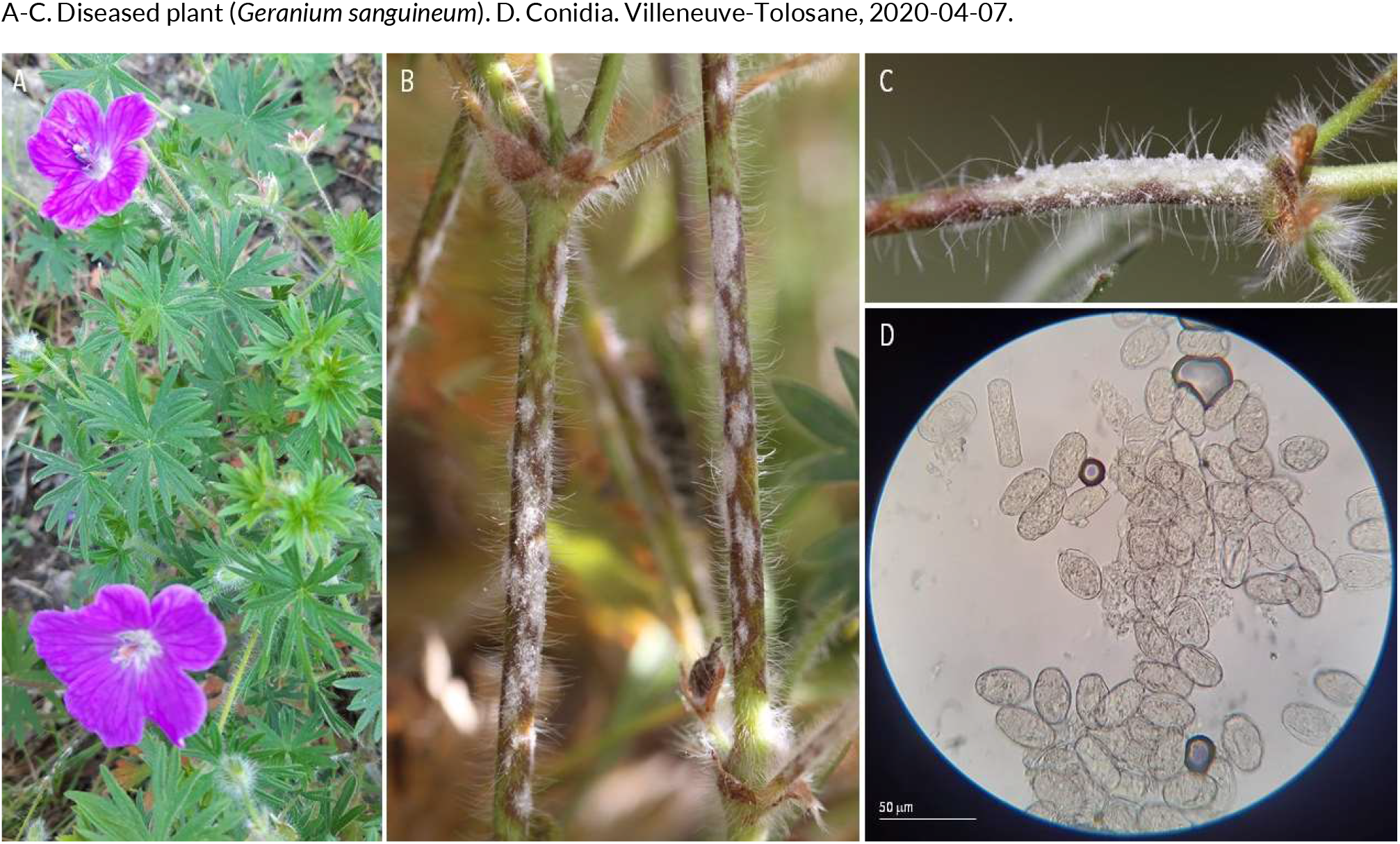
POWDERY MILDEW | *Podosphaera fugax* ex *Geranium molle* (dove’s-foot crane’s-bill) and *Geranium sanguineum* (bloody crane’s-bill)

**122.**
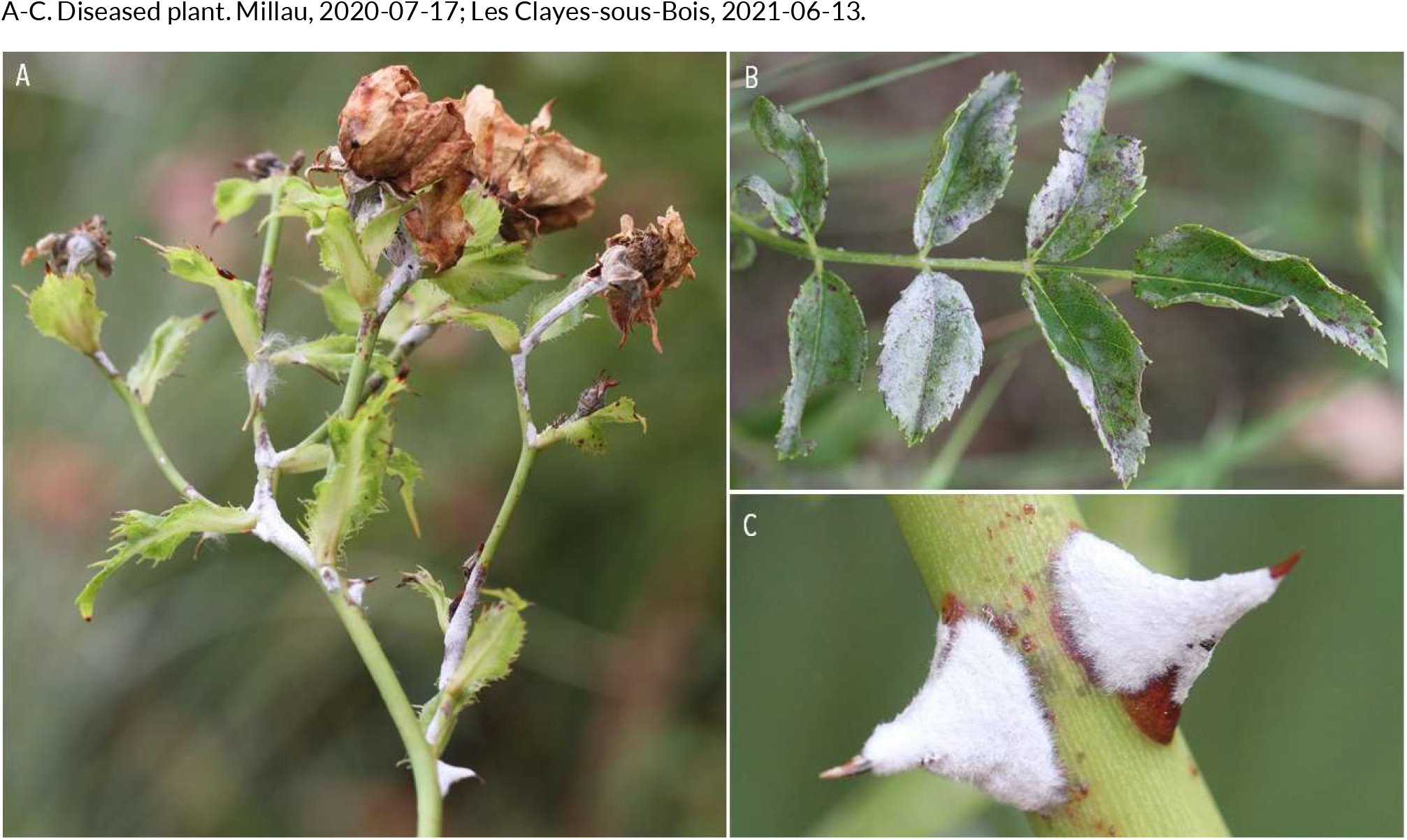
POWDERY MILDEW | *Podosphaera pannosa* ex *Rosa sp*. (rose)

**123.**
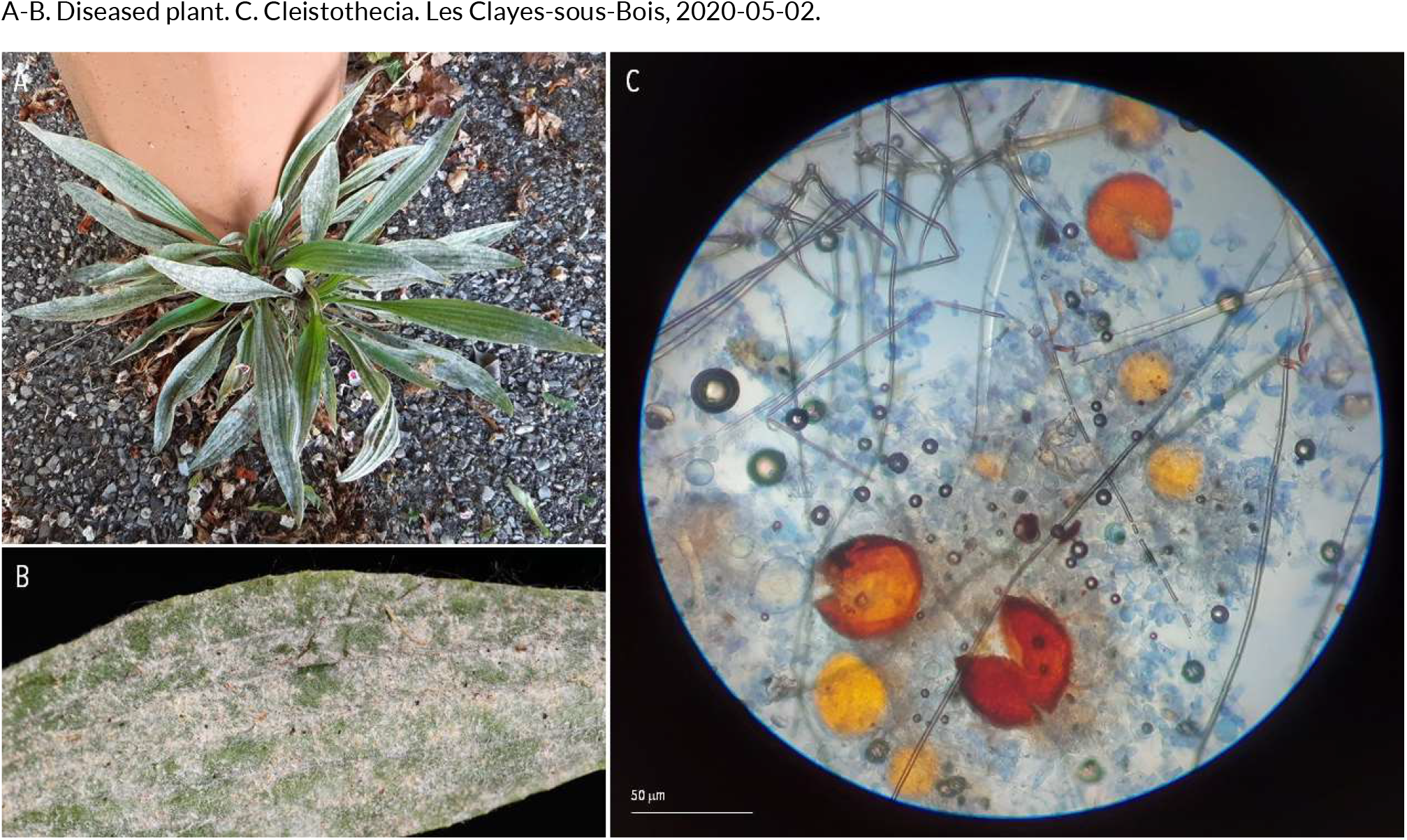
POWDERY MILDEW | *Podosphaera plantaginis* ex *Plantago lanceolata* (ribwort plantain)

**124.**
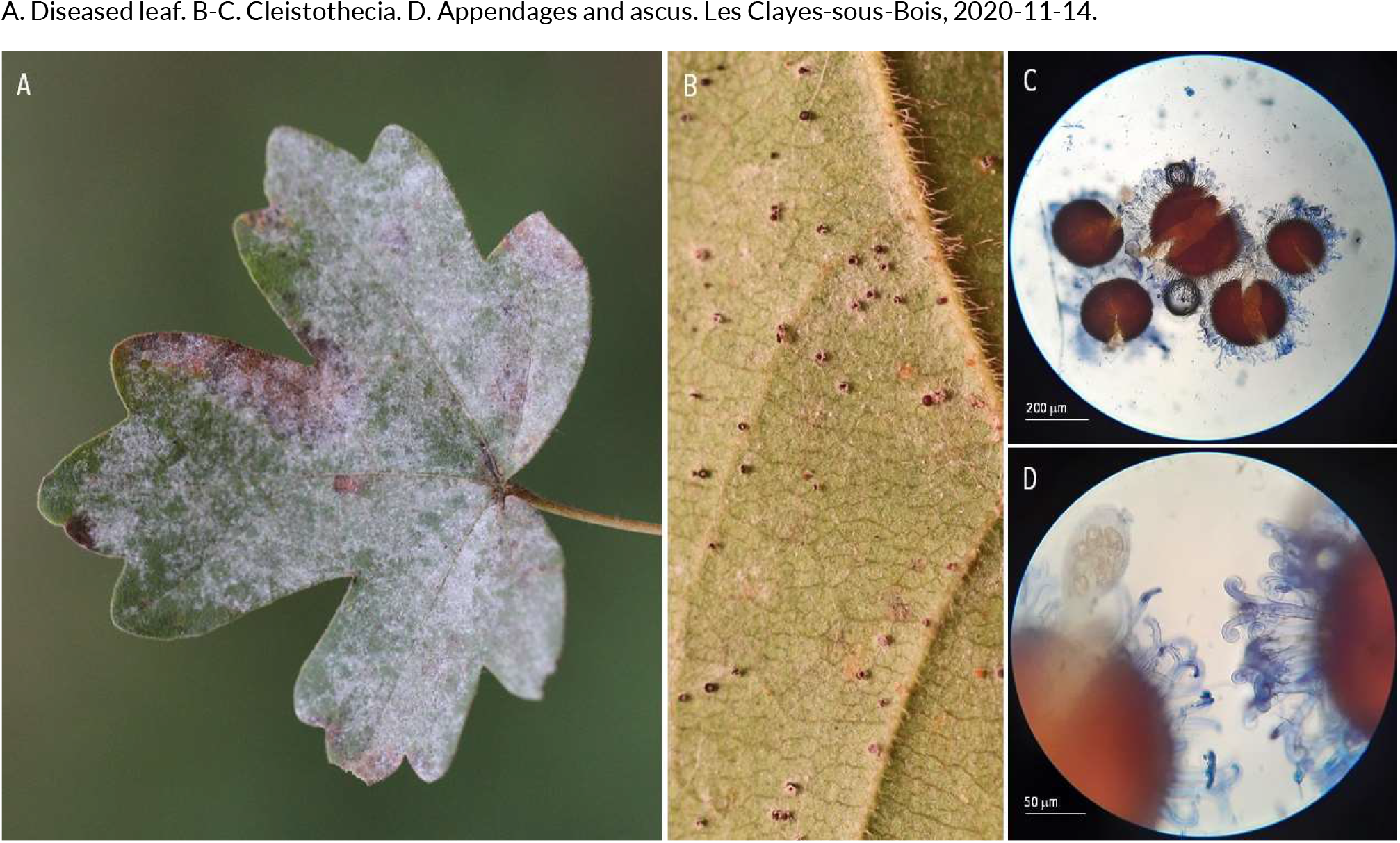
POWDERY MILDEW | *Sawadaea bicornis* ex *Acer campestre* (field maple)

**125.**
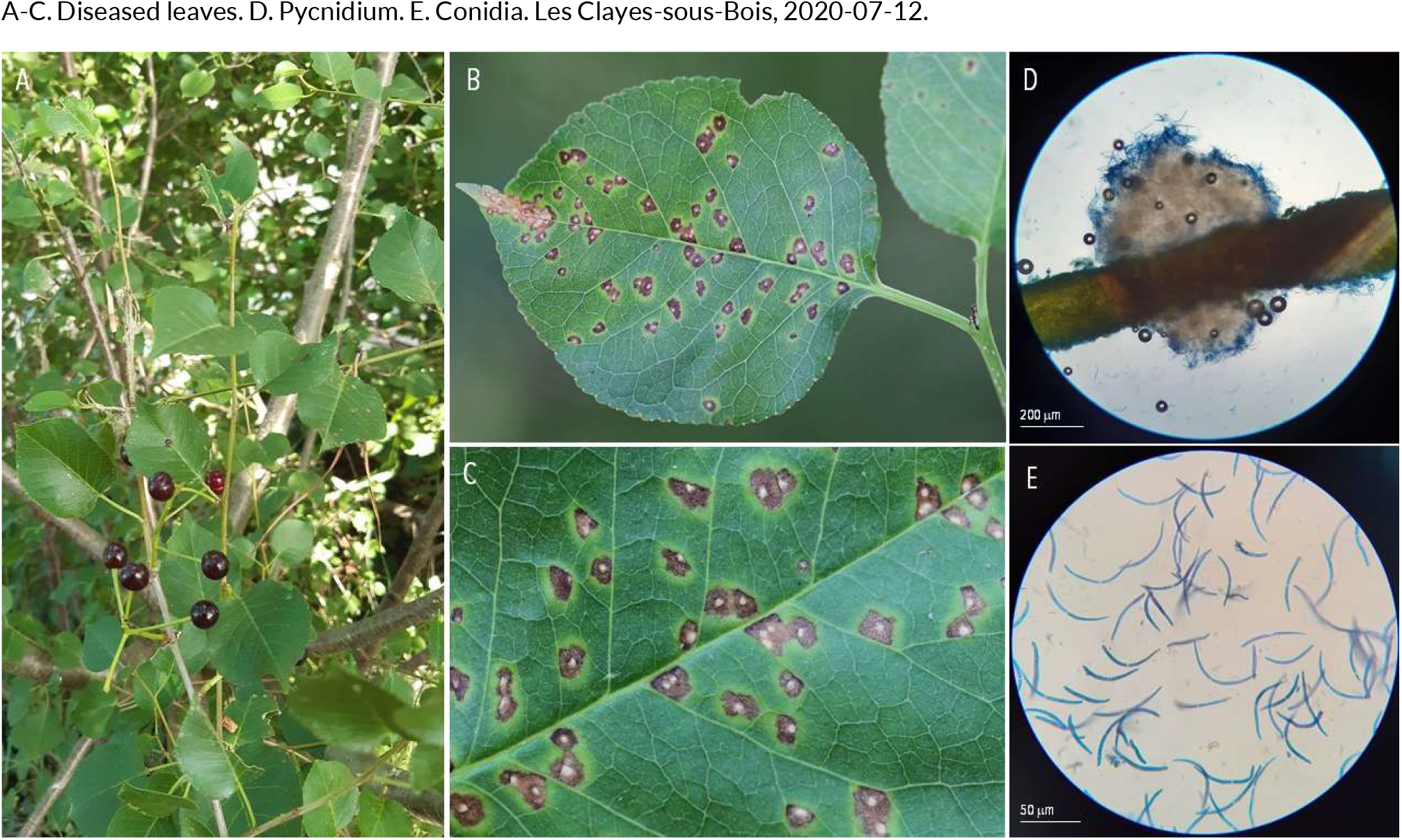
SEPTORIA-LIKE DISEASES | *Cylindrosporium padi* ex *Cerasus mahaleb* (St. Lucie cherry)

**126.**
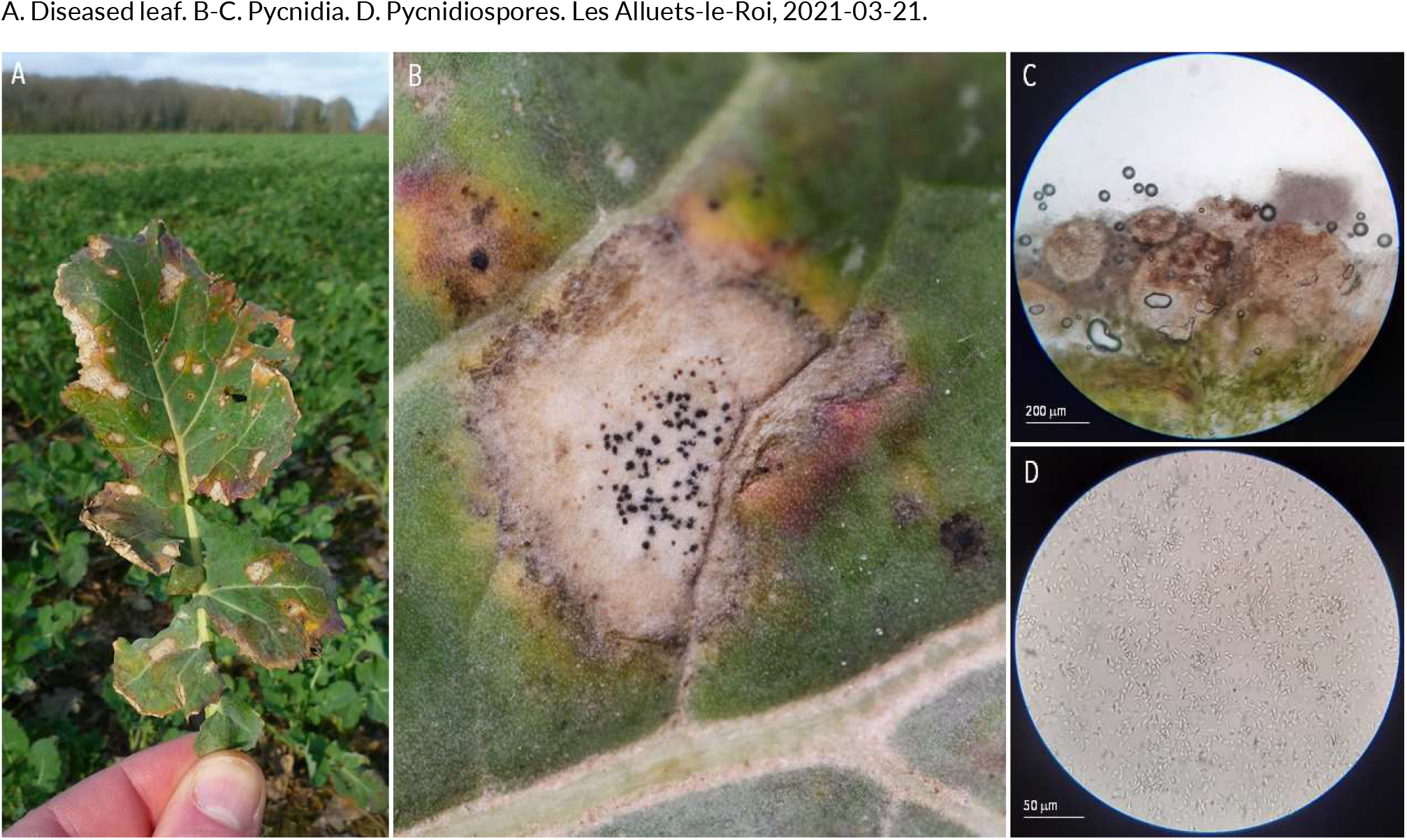
SEPTORIA-LIKE DISEASES | *Leptosphaeria maculans* ex *Brassica napus* (oilseed rape)

**127.**
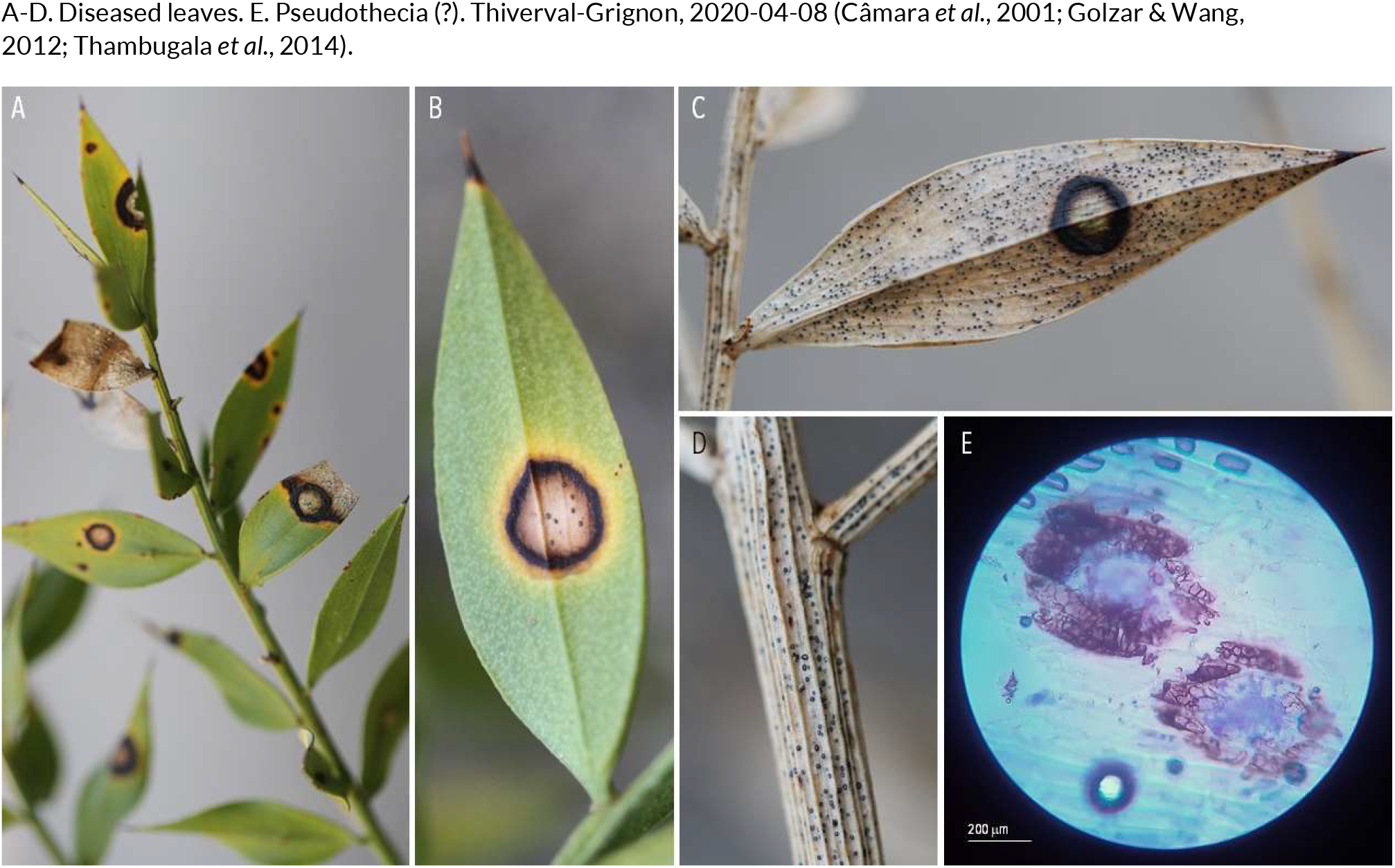
SEPTORIA-LIKE DISEASES | *Leptosphaeria rusci* or *Phyllosticta ruscigena* (??) ex *Ruscus aculeatus* (butcher’s broom)

**128.**
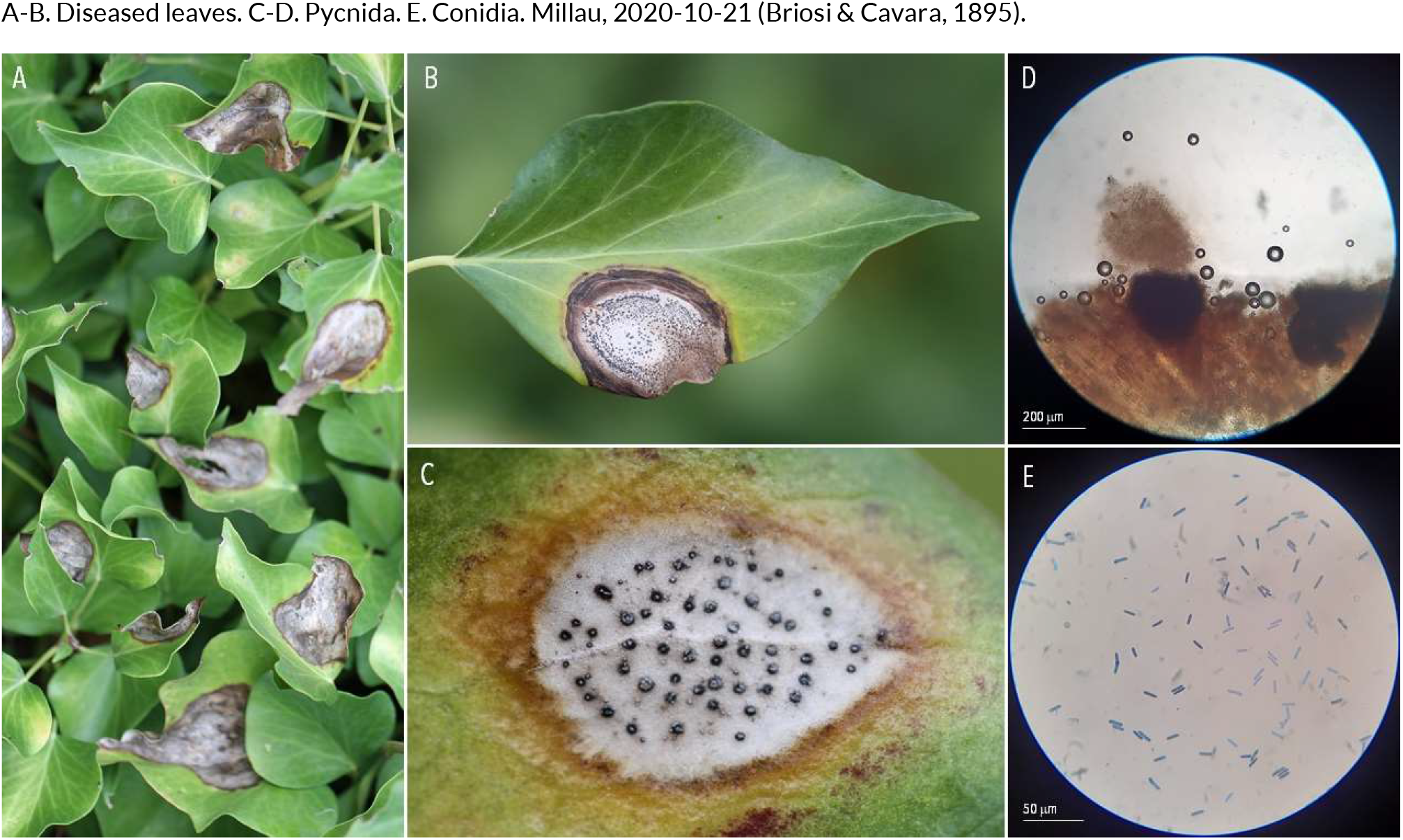
SEPTORIA-LIKE DISEASES | *Mycosphaerella hedericola* ex *Hedera helix* (common ivy.

**129.**
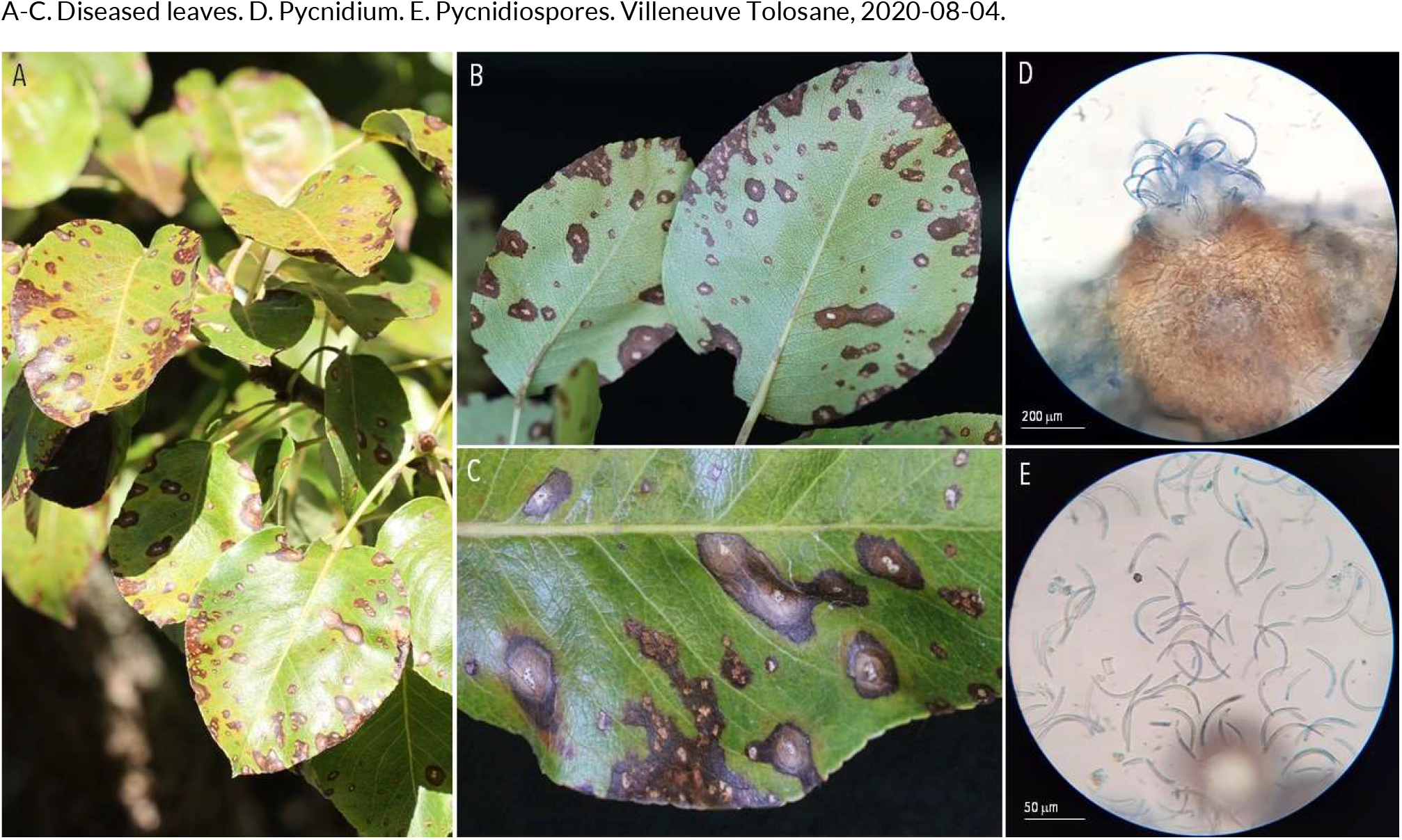
SEPTORIA-LIKE DISEASES | *Mycosphaerella pyri* ex *Pyrus communis* (pear.

**130.**
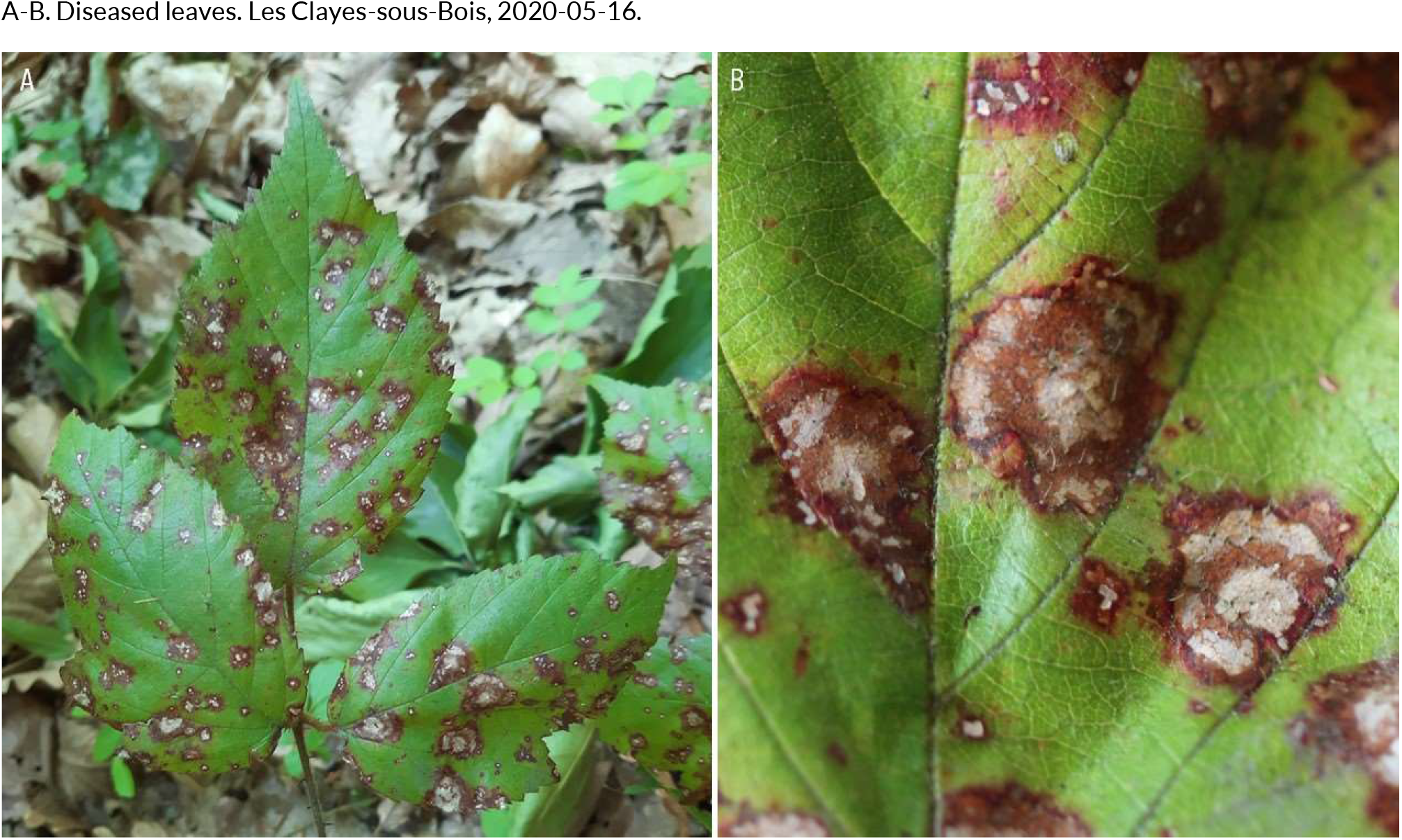
SEPTORIA-LIKE DISEASES | *Mycosphaerella rubi* ex *Rubus fruticosus* (bramble)

**131.**
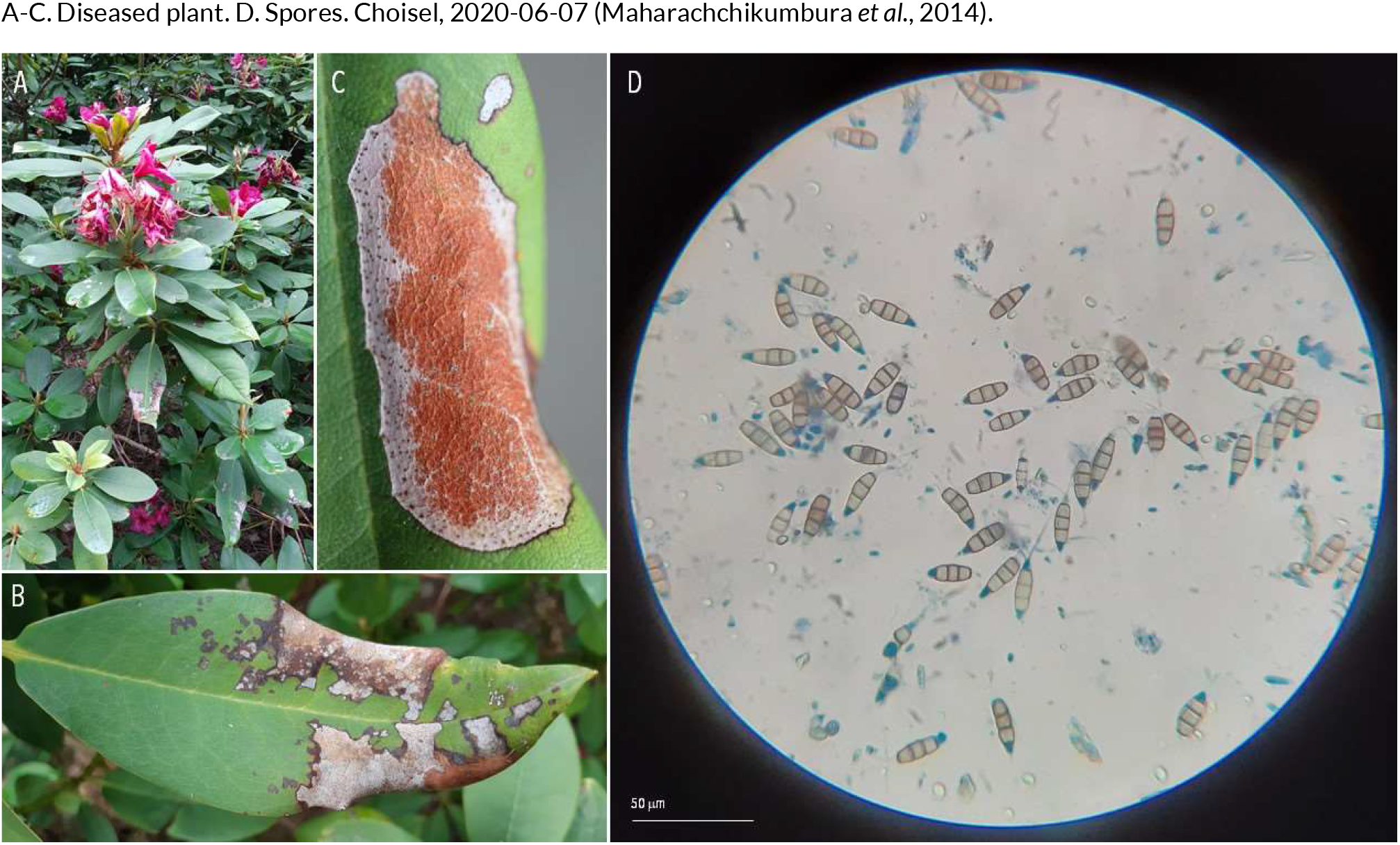
SEPTORIA-LIKE DISEASES | *Pestalotiopsis guepinii* ex *Rhododendron sp*. (rhododendron)

**132.**
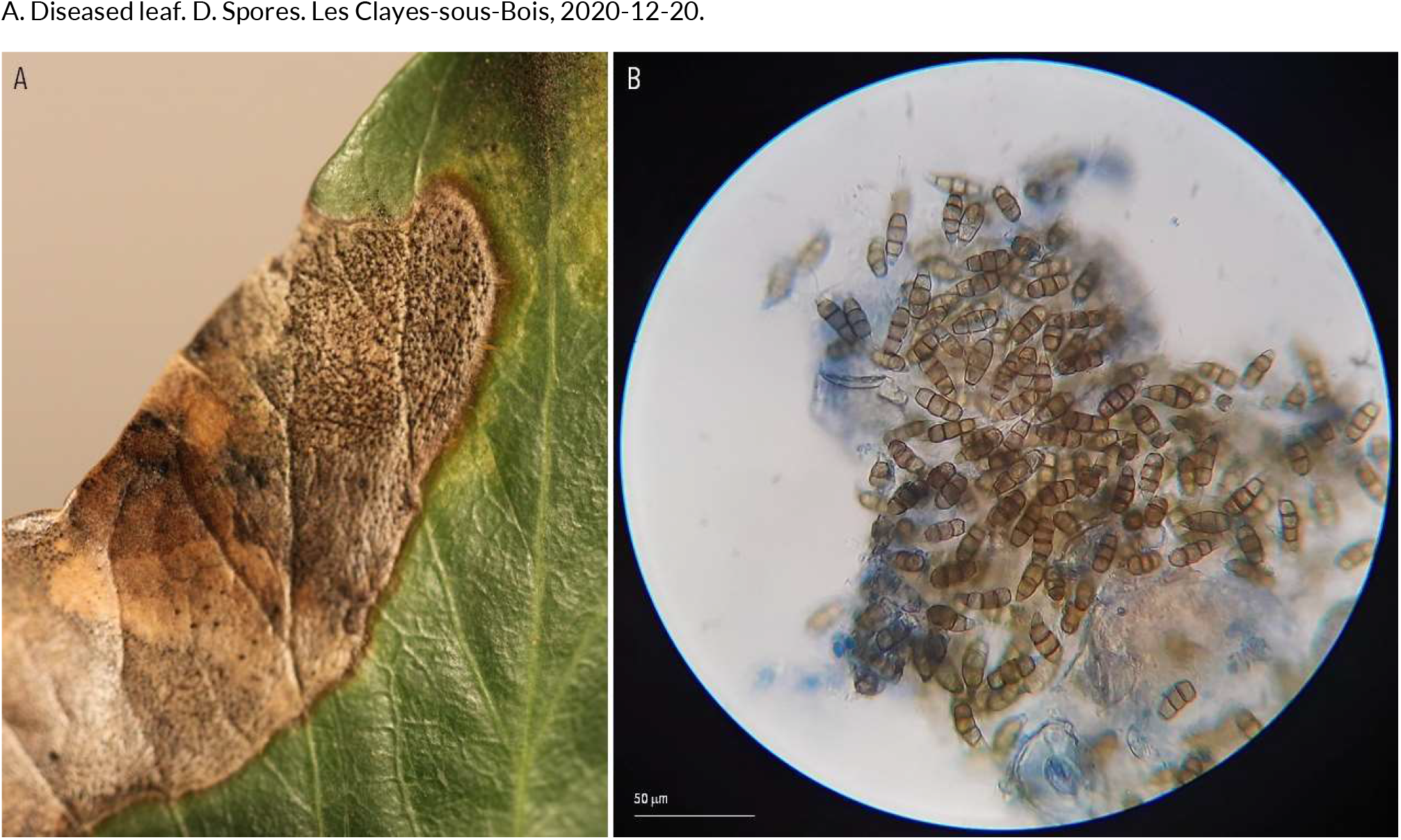
SEPTORIA-LIKE DISEASES | *Pestalotiopsis* sp. (??) ex *Hedera helix* (common ivy)

**133.**
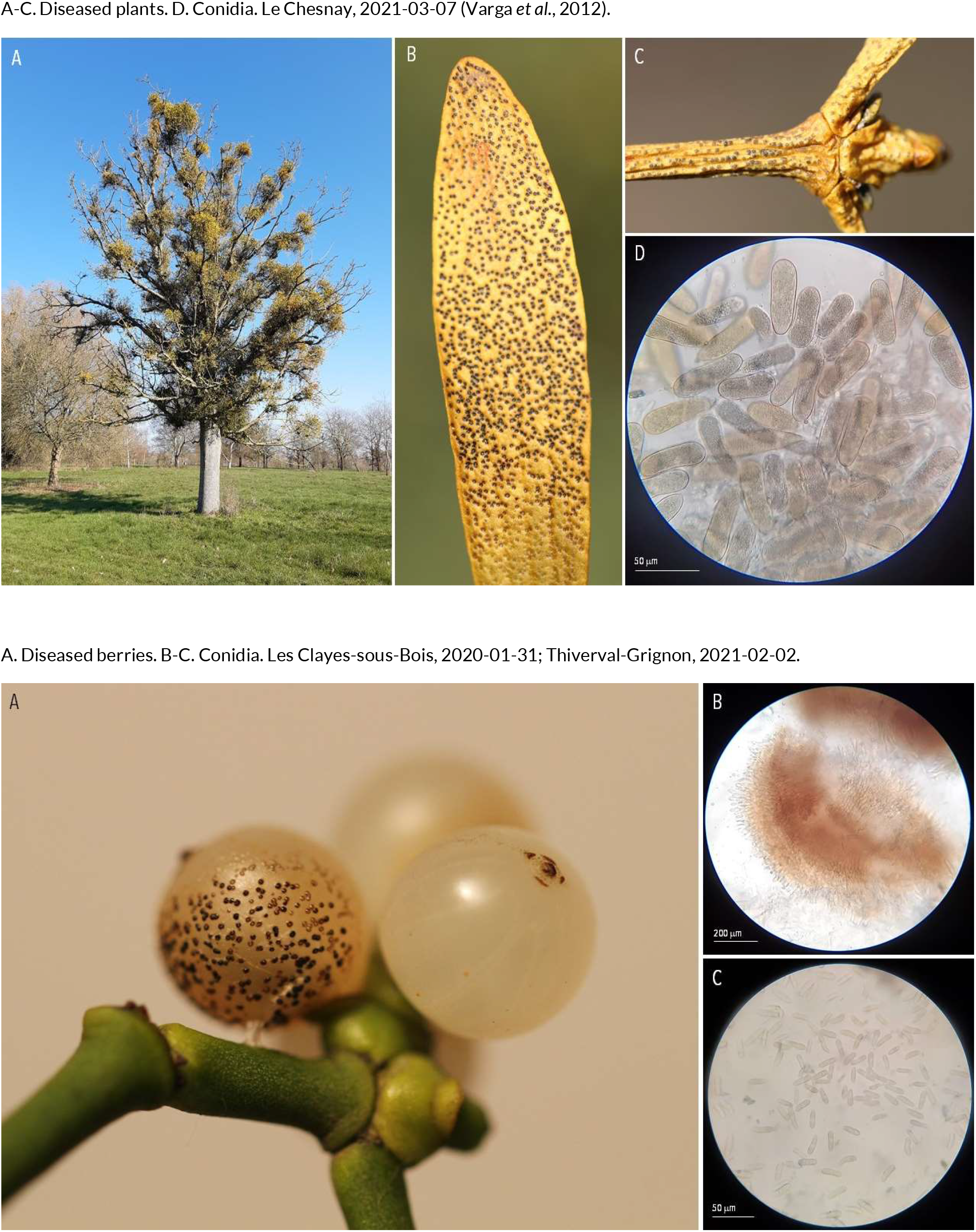
SEPTORIA-LIKE DISEASES | *Phaeobotryosphaeria visci* ex *Viscum album* (European mistletoe)

**134.**
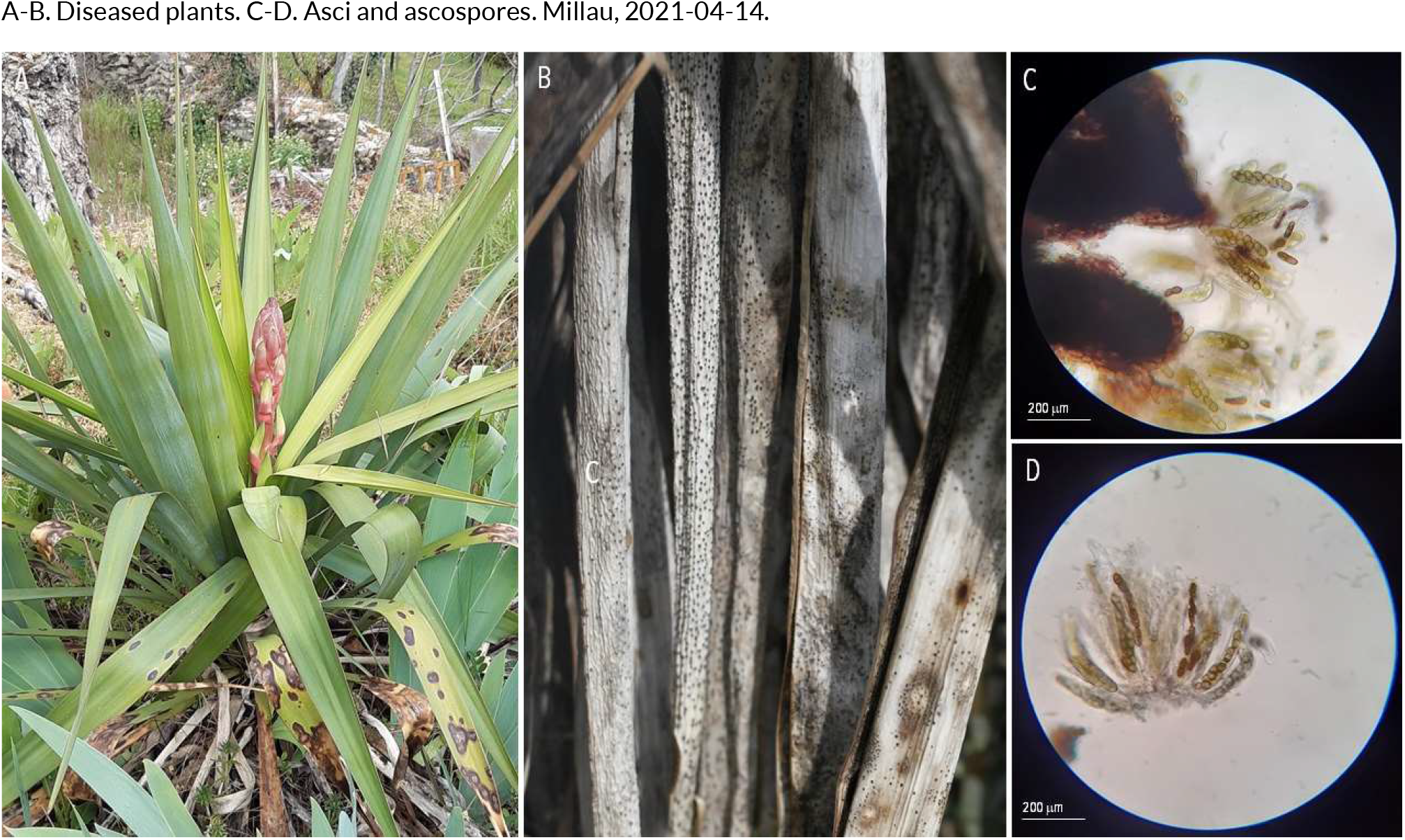
SEPTORIA-LIKE DISEASES | *Phaeobotryosphaeria* sp. (??) ex *Yucca gloriosa* (Adam’s needle)

**135.**
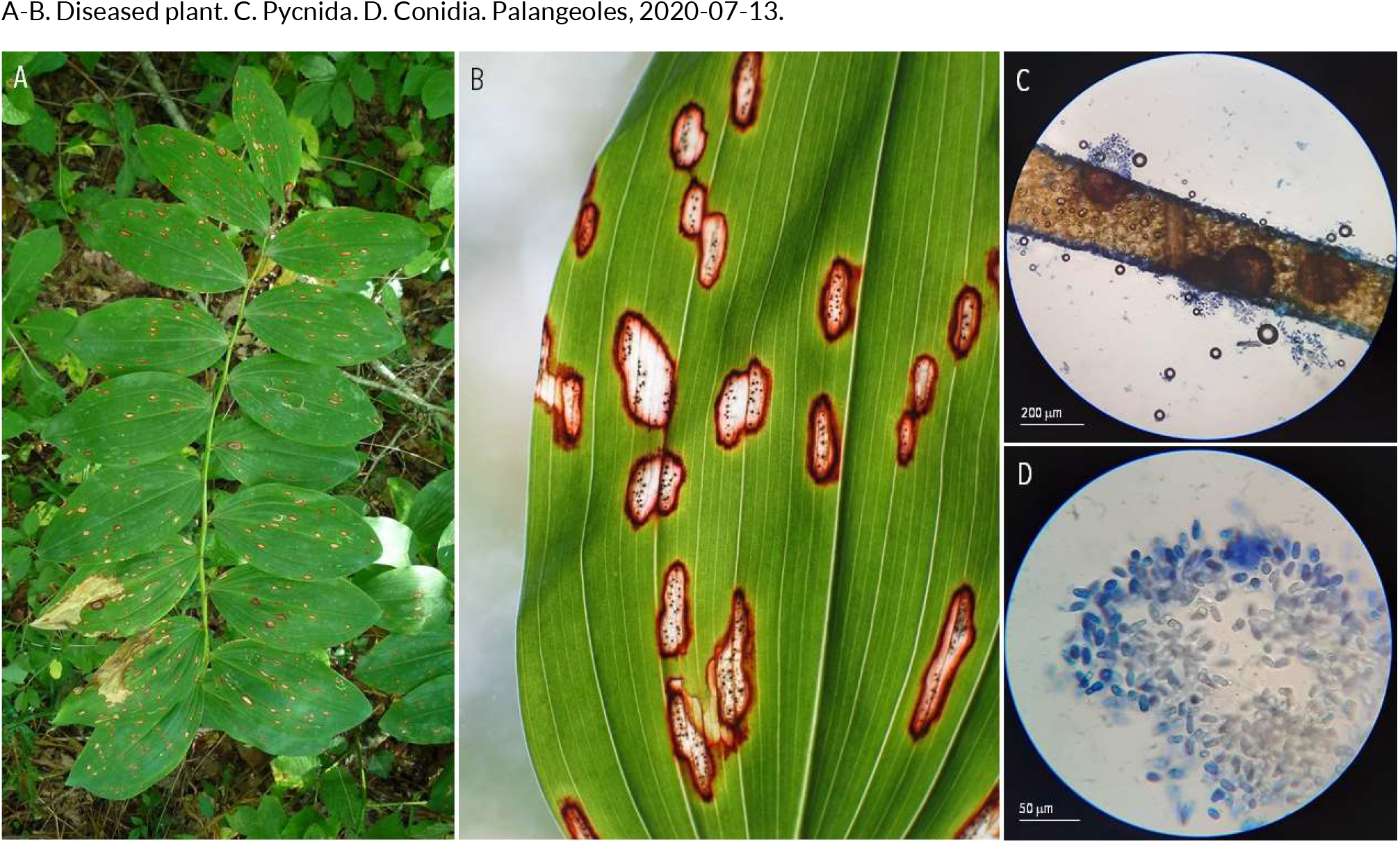
SEPTORIA-LIKE DISEASES | *Phyllosticta cruenta* ex *Polygonatum multiflorum* (Solomon’s seal)

**136.**
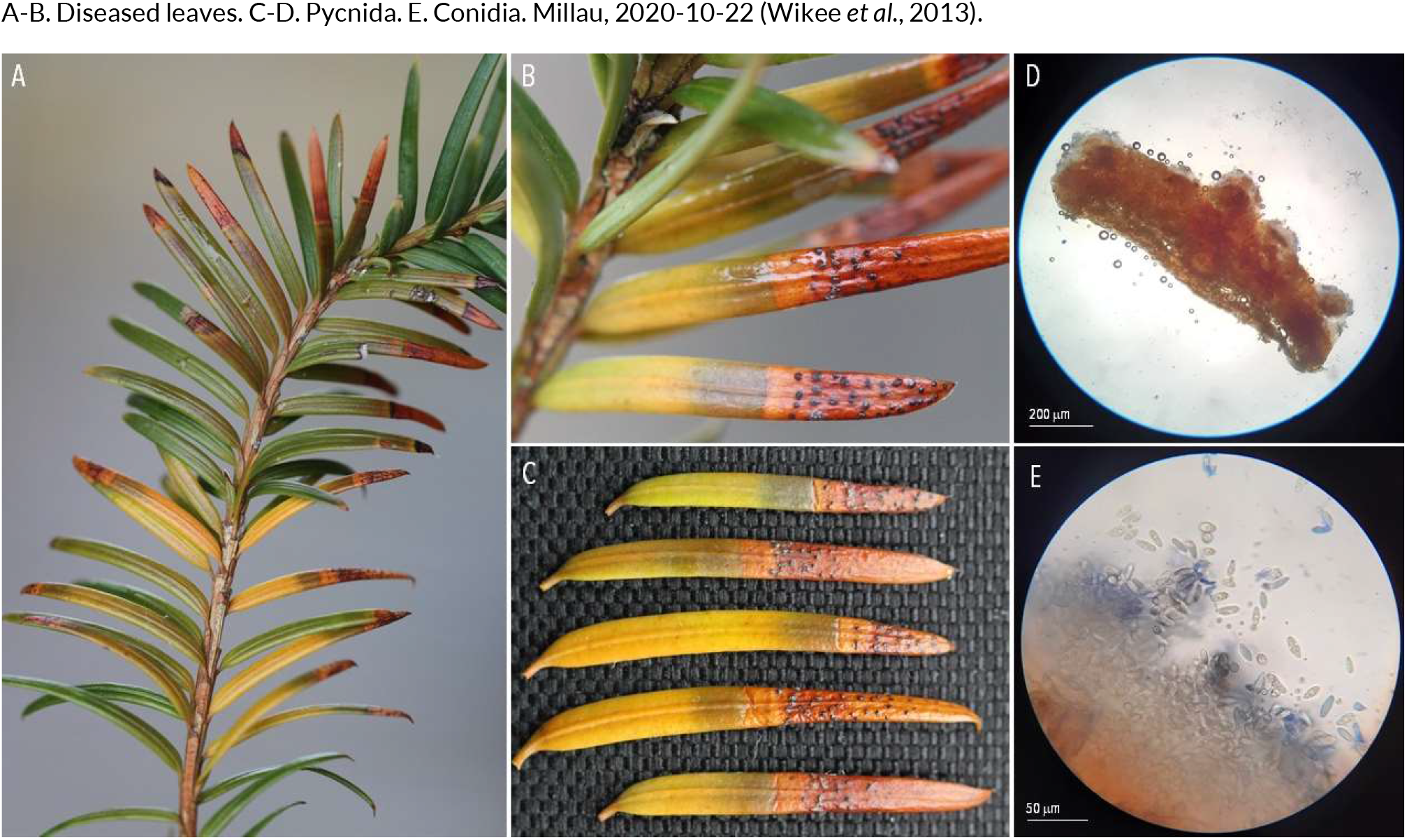
SEPTORIA-LIKE DISEASES | *Phyllosticta foliorum* or *Phyllosticta philoprina* (??) ex *Taxus baccata* (common yew)

**137.**
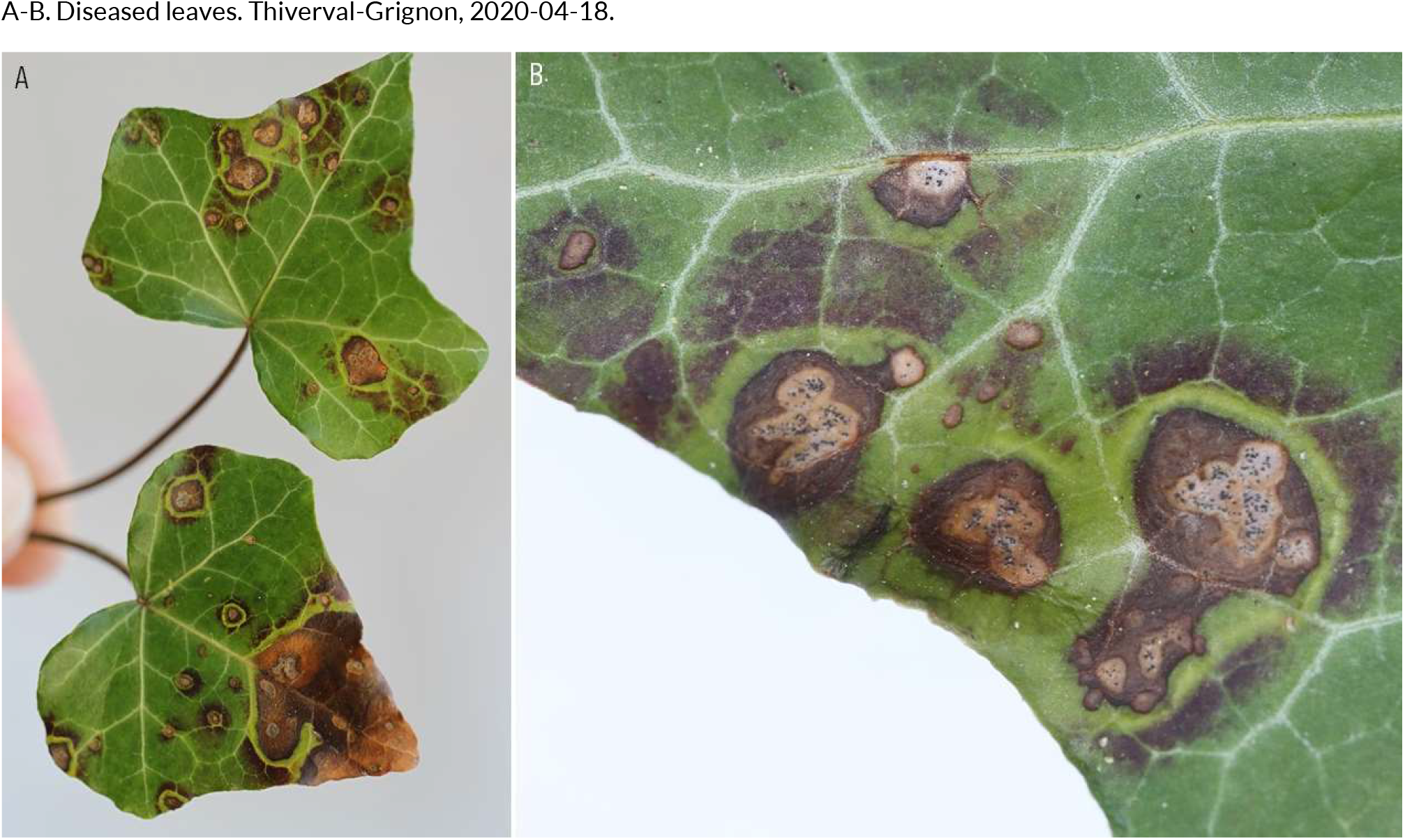
SEPTORIA-LIKE DISEASES | *Phyllosticta hederae* (?) ex *Hedera helix* (common ivy)

**138.**
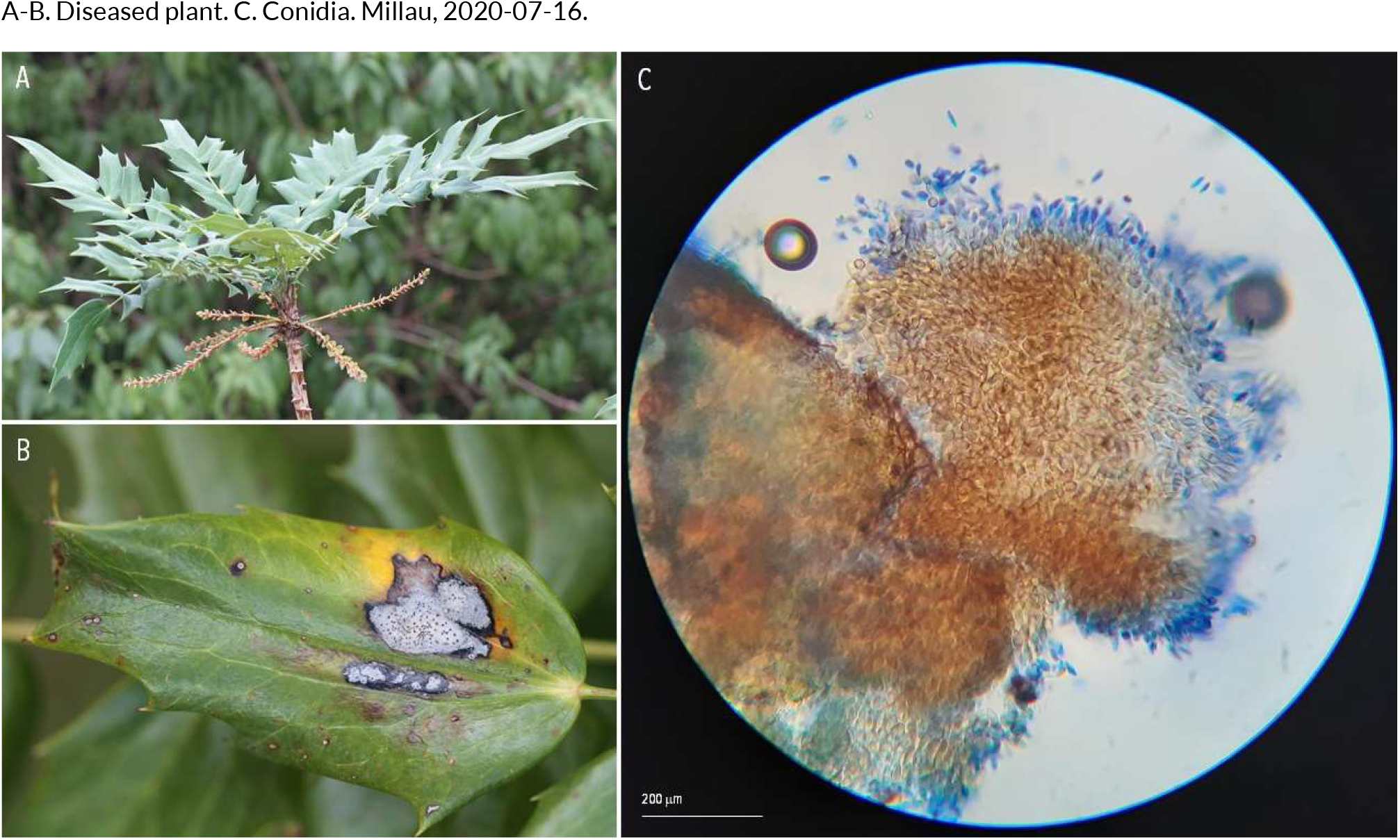
SEPTORIA-LIKE DISEASES | *Phyllosticta mahoniae* ex *Mahonia japonica* (Japanese mahonia)

**139.**
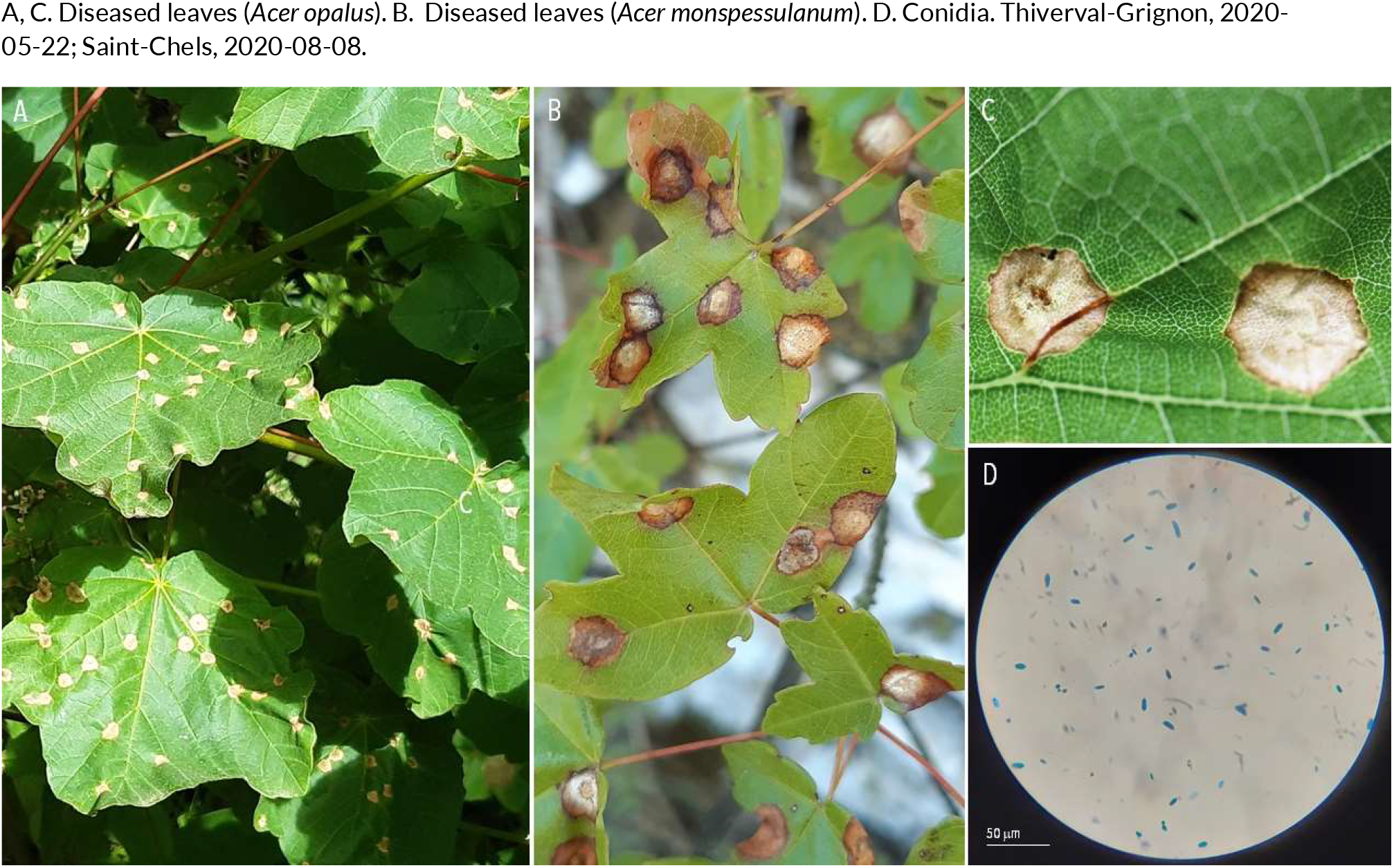
SEPTORIA-LIKE DISEASES | *Phyllosticta minima* (??) ex *Acer oplaus* (Italian maple) and *Acer monspessulanum* (Montpellier maple)

**140.**
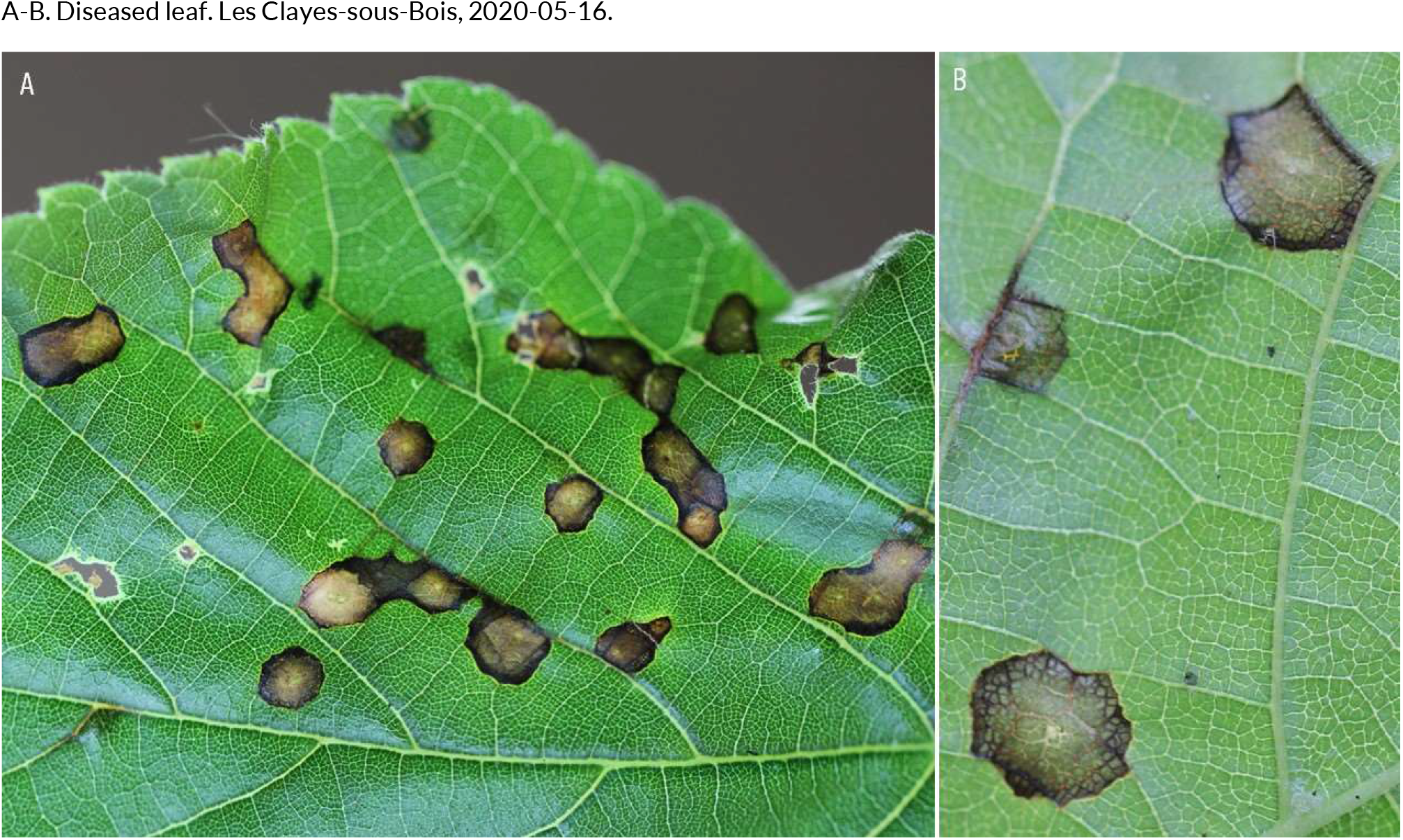
SEPTORIA-LIKE DISEASES | *Phyllosticta tiliae* (??) ex *Tilia cordata* (linden.

**141.**
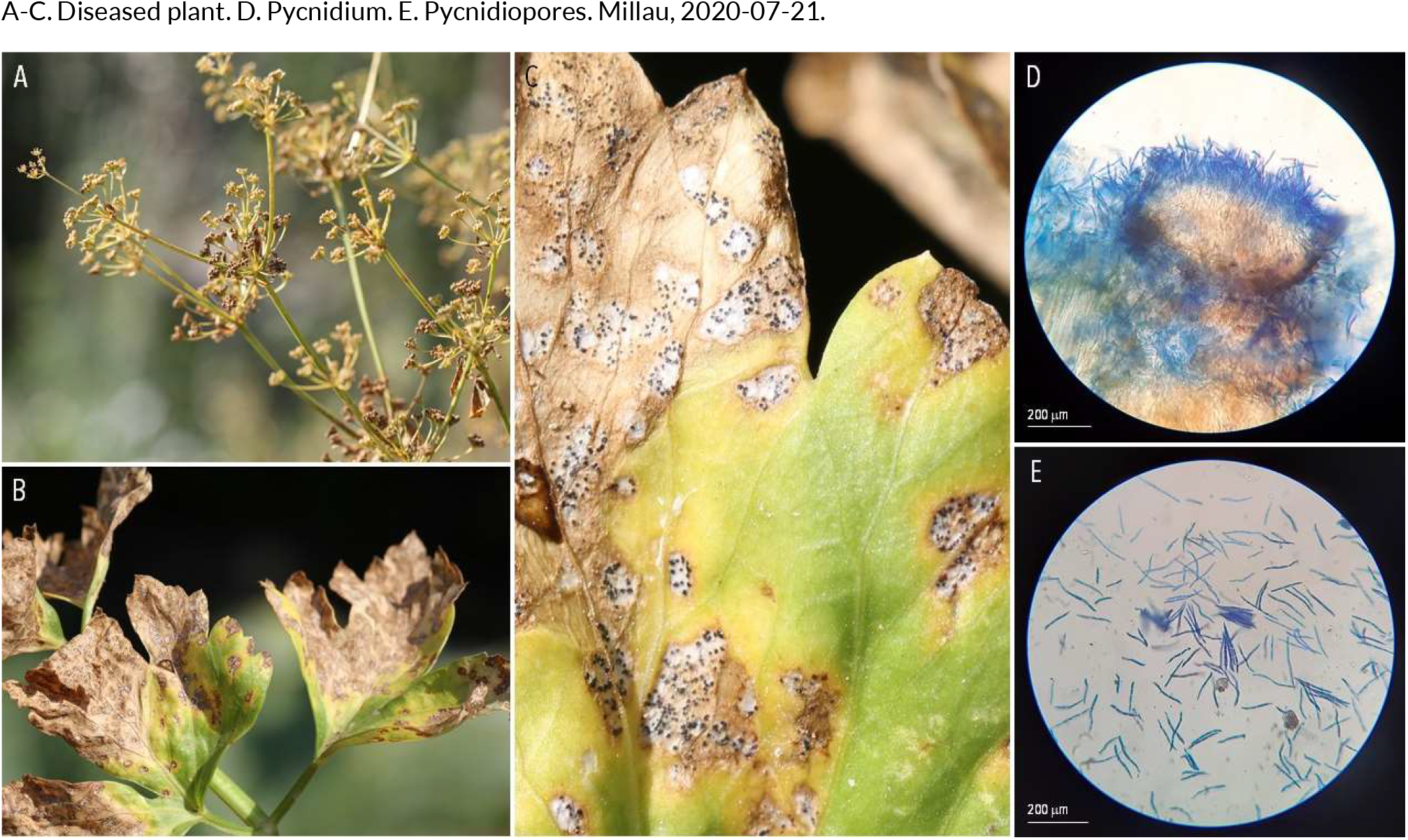
SEPTORIA-LIKE DISEASES | *Septoria apiicola* ex *Apium graveolens* (celery)

**142.**
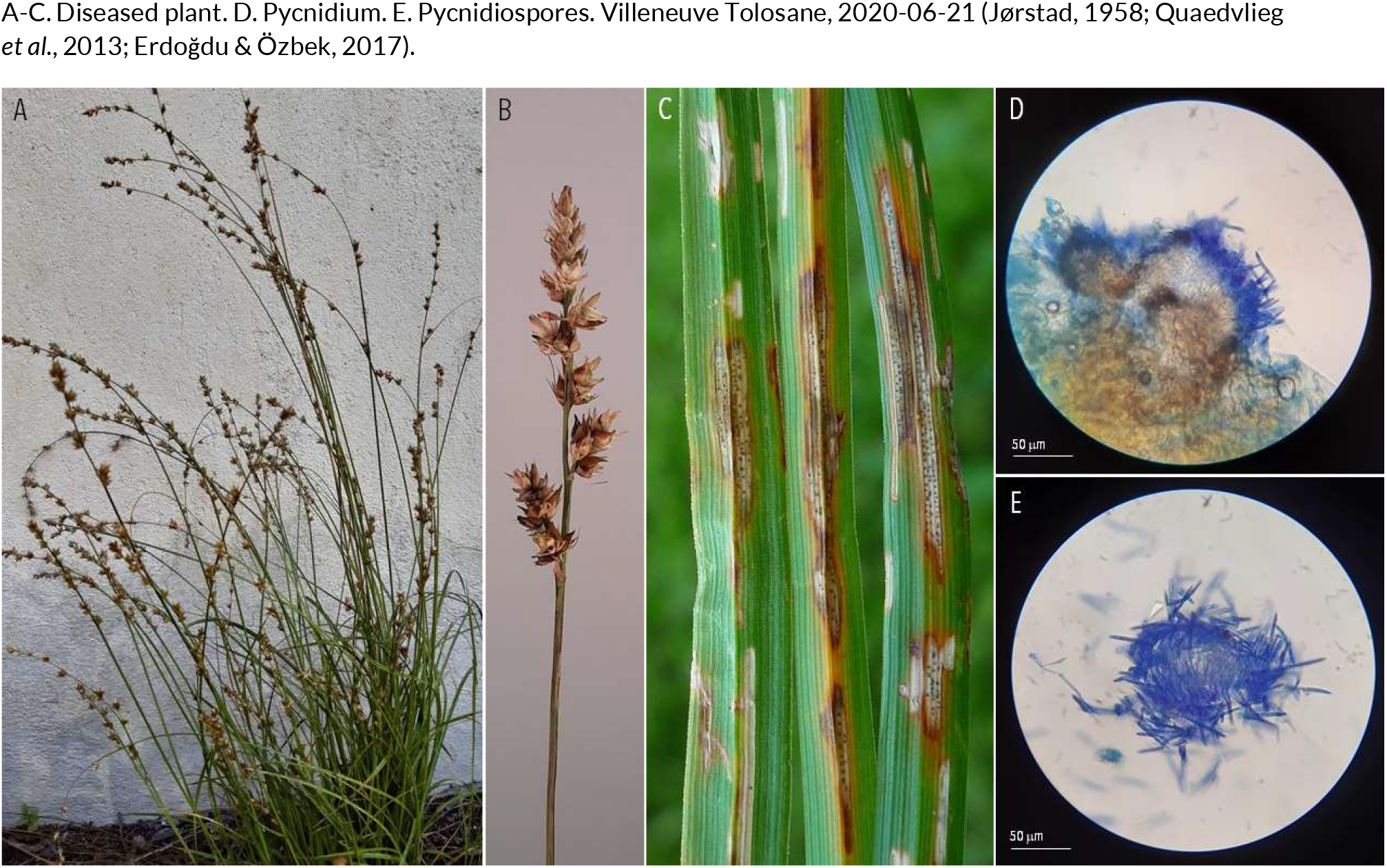
SEPTORIA-LIKE DISEASES | *Septoria caricis* ex *Carex divulsa subsp. divulsa* (grey sedge)

**143.**
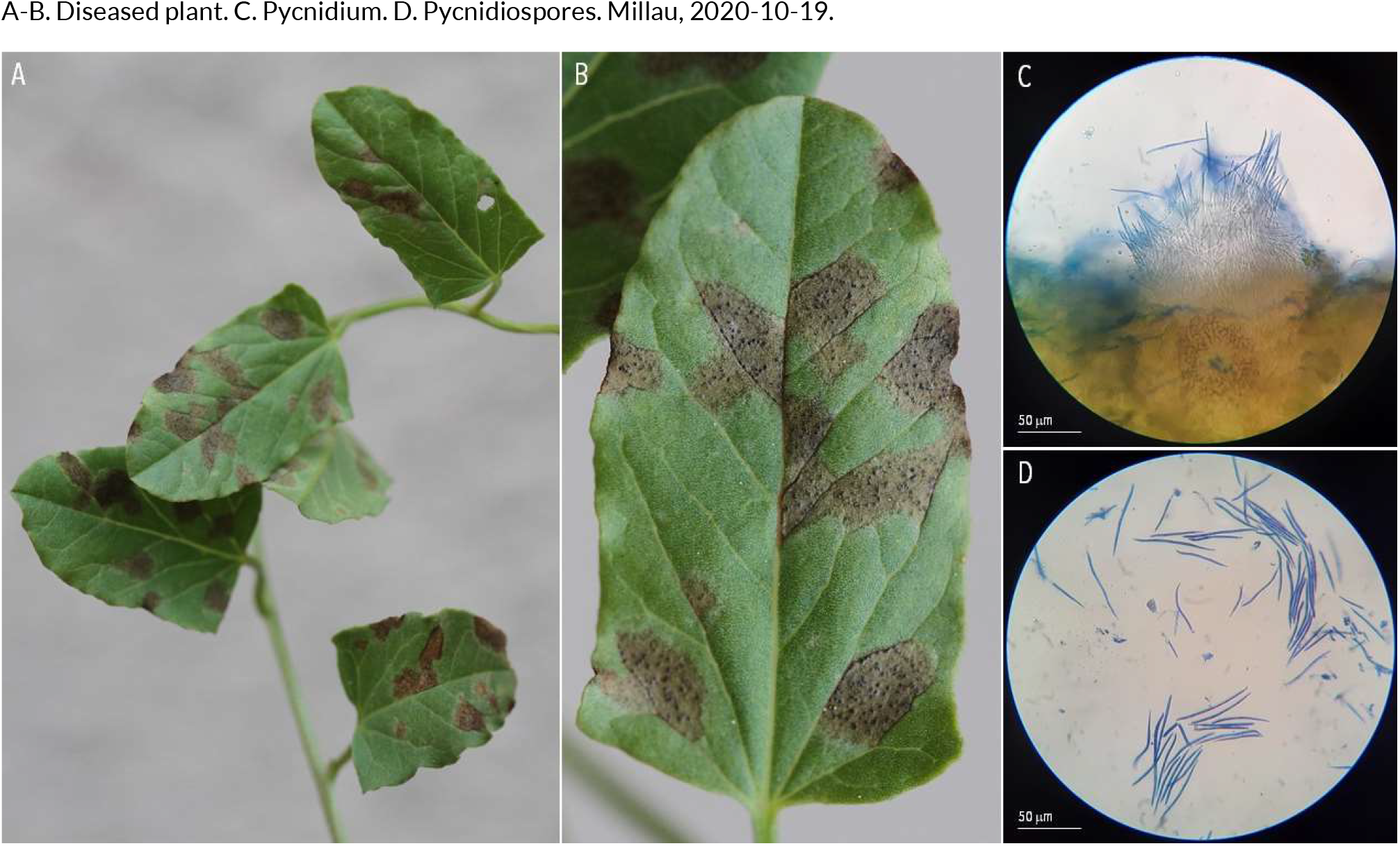
SEPTORIA-LIKE DISEASES | *Septoria convolvuli* ex *Convolvulus arvensis* (field bindweed)

**144.**
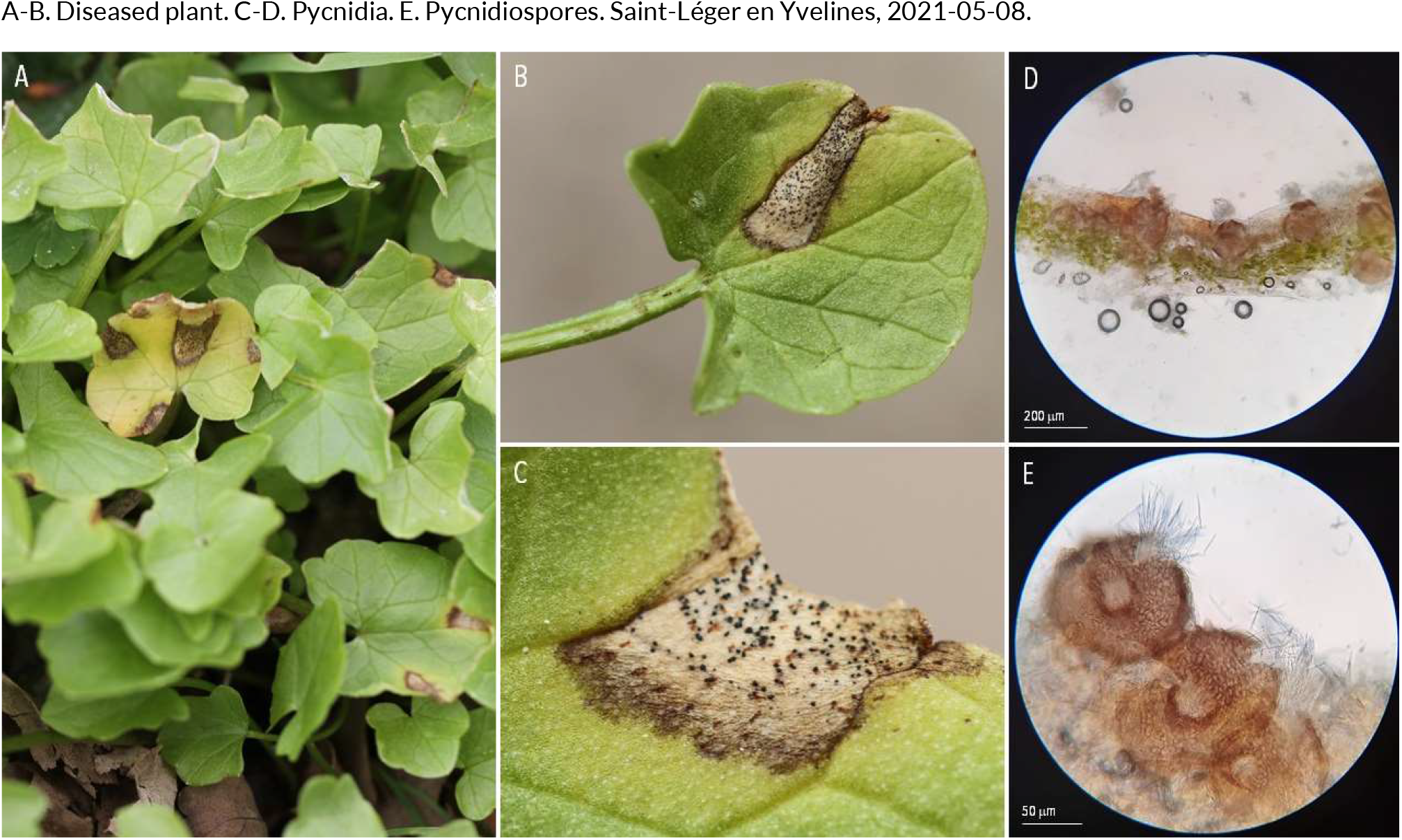
SEPTORIA-LIKE DISEASES | *Septoria ficariae* ex *Ficaria verna* (lesser celandine)

**145.**
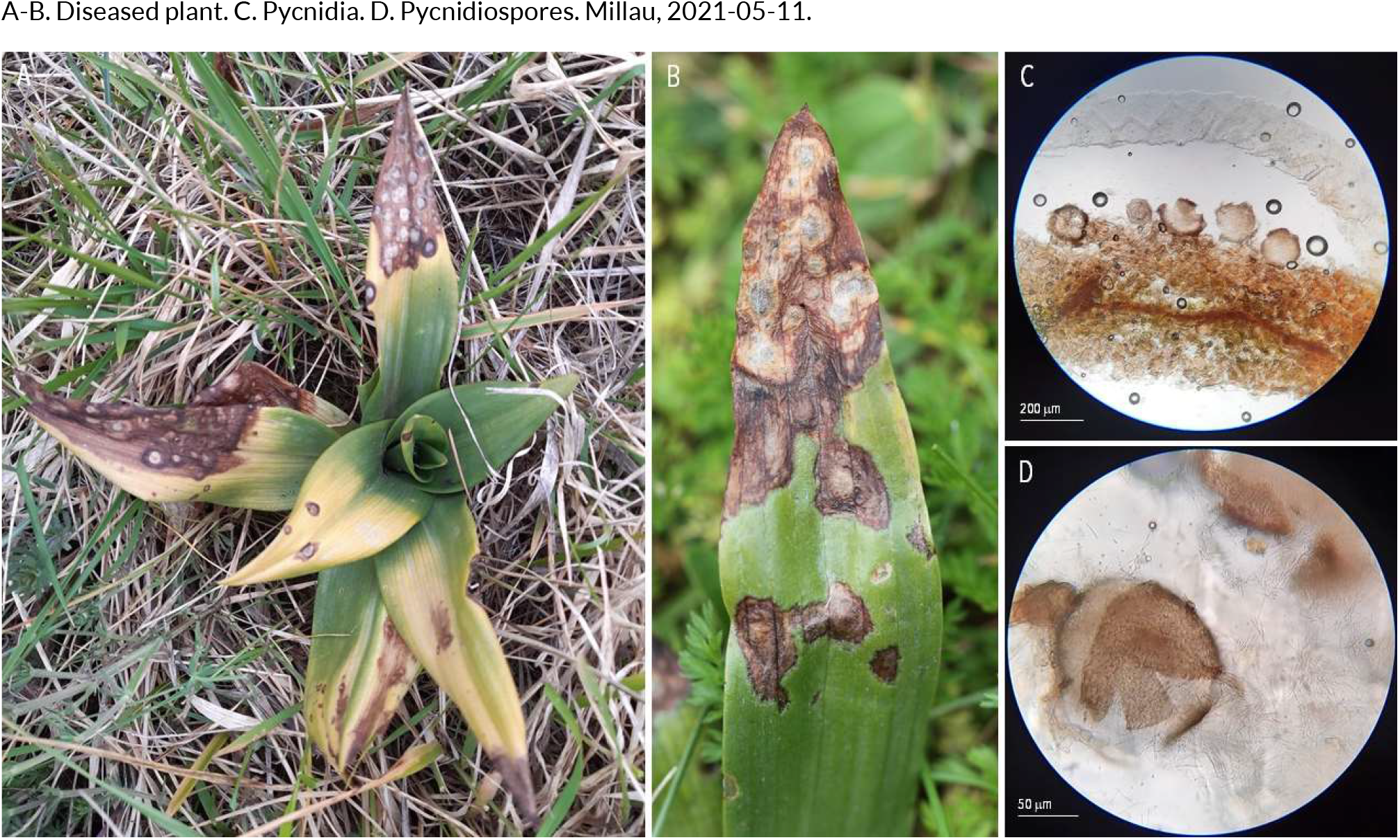
SEPTORIA-LIKE DISEASES | *Septoria orchidearum* (?) ex *Orchis pyramidalis* (pyramidal orchid.

**146.**
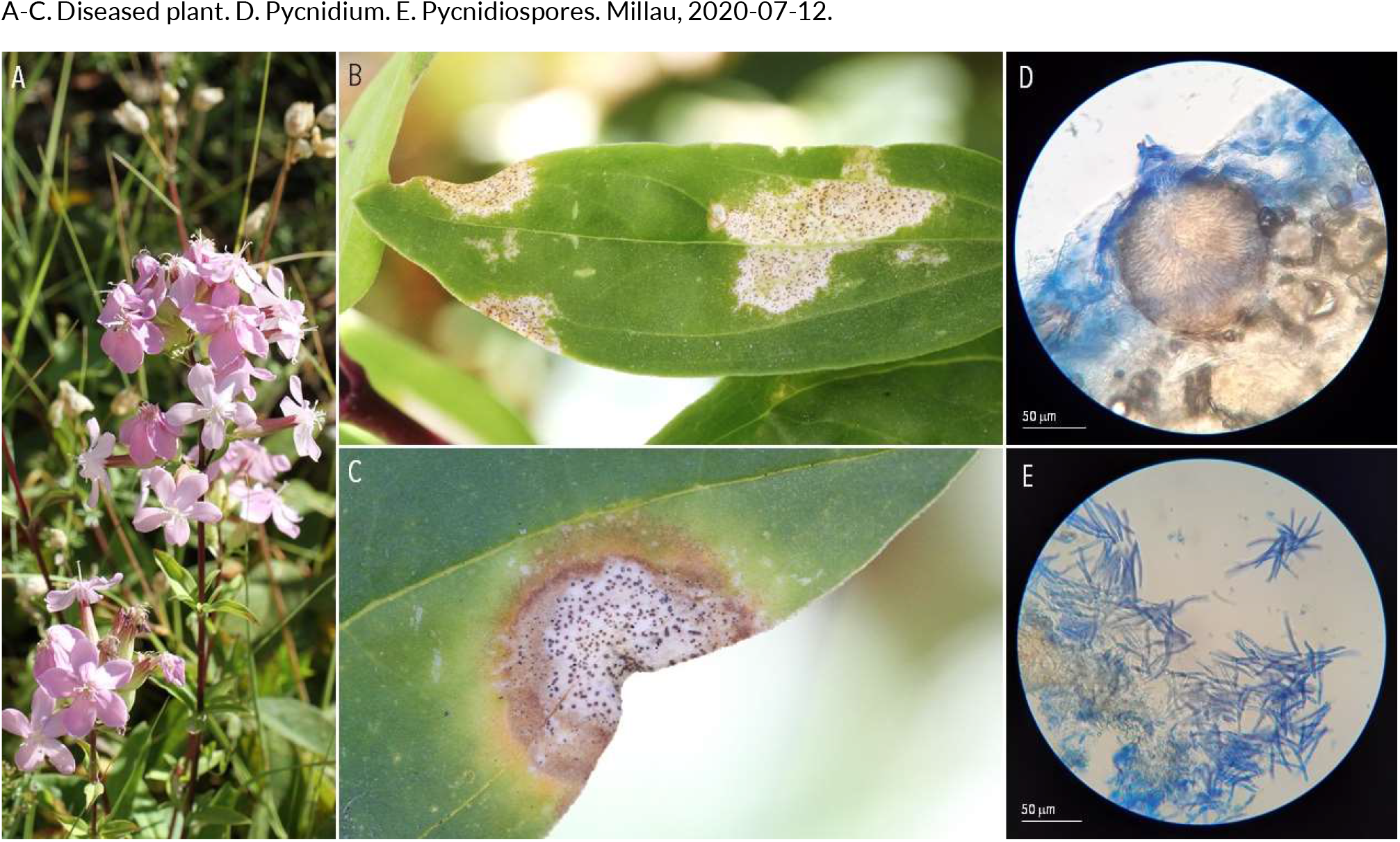
SEPTORIA-LIKE DISEASES | *Septoria saponariae* ex *Saponaria officinalis* (common soapwort.

**147.**
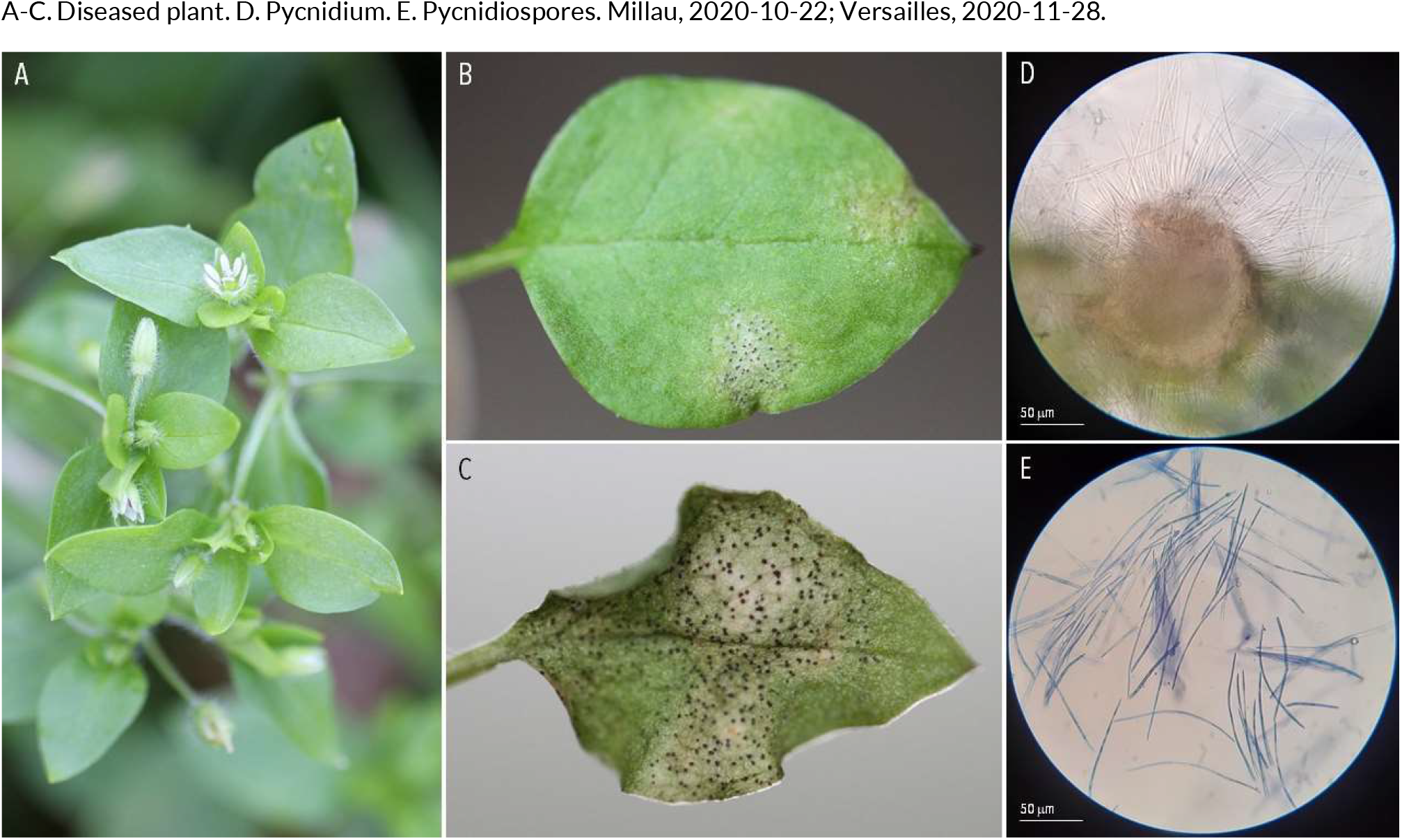
SEPTORIA-LIKE DISEASES | *Septoria stellariae* ex *Stellaria media* (chickweed.

**148.**
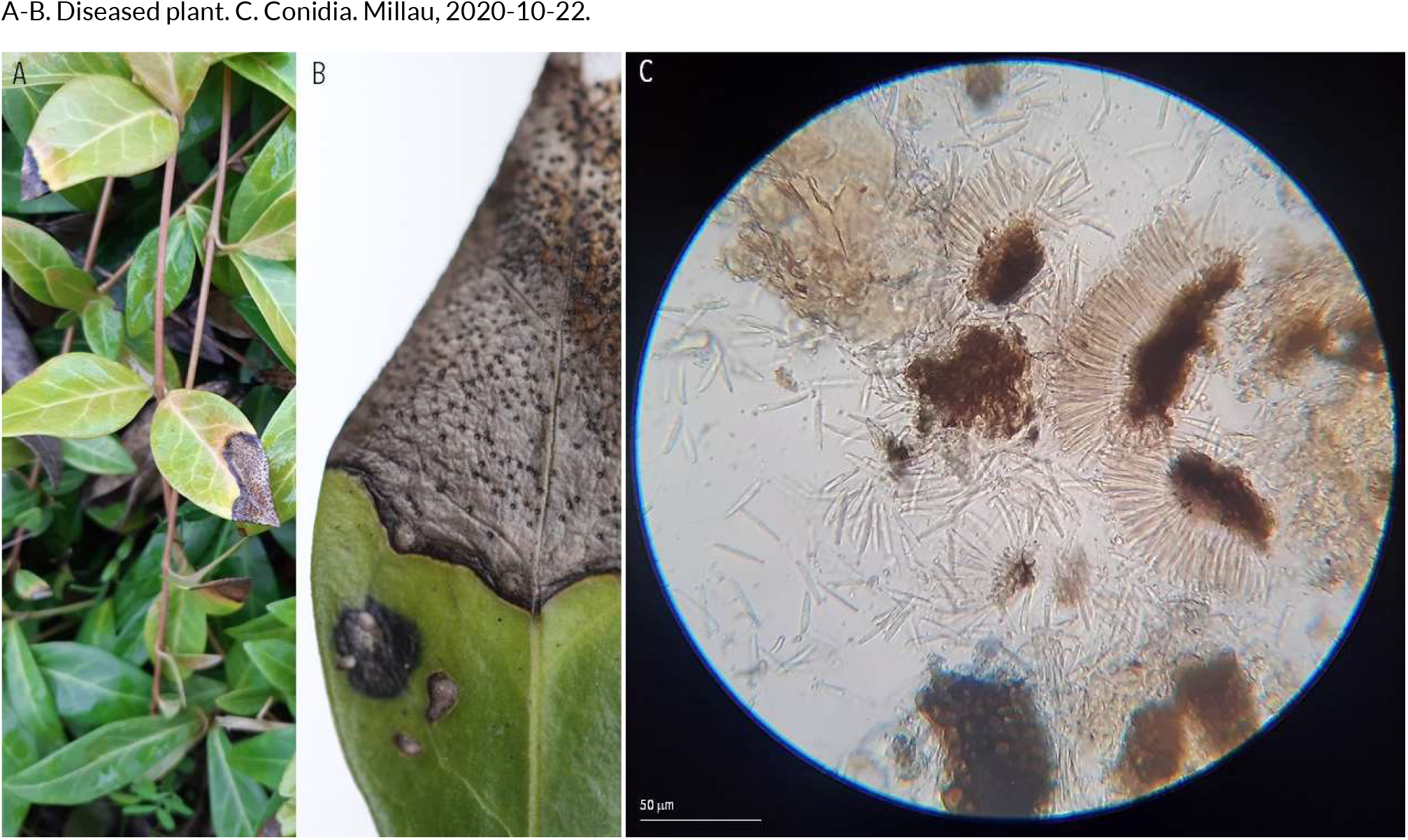
SEPTORIA-LIKE DISEASES | *Septoria vincae* ex *Vinca minor* (lesser periwinkle)

**149.**
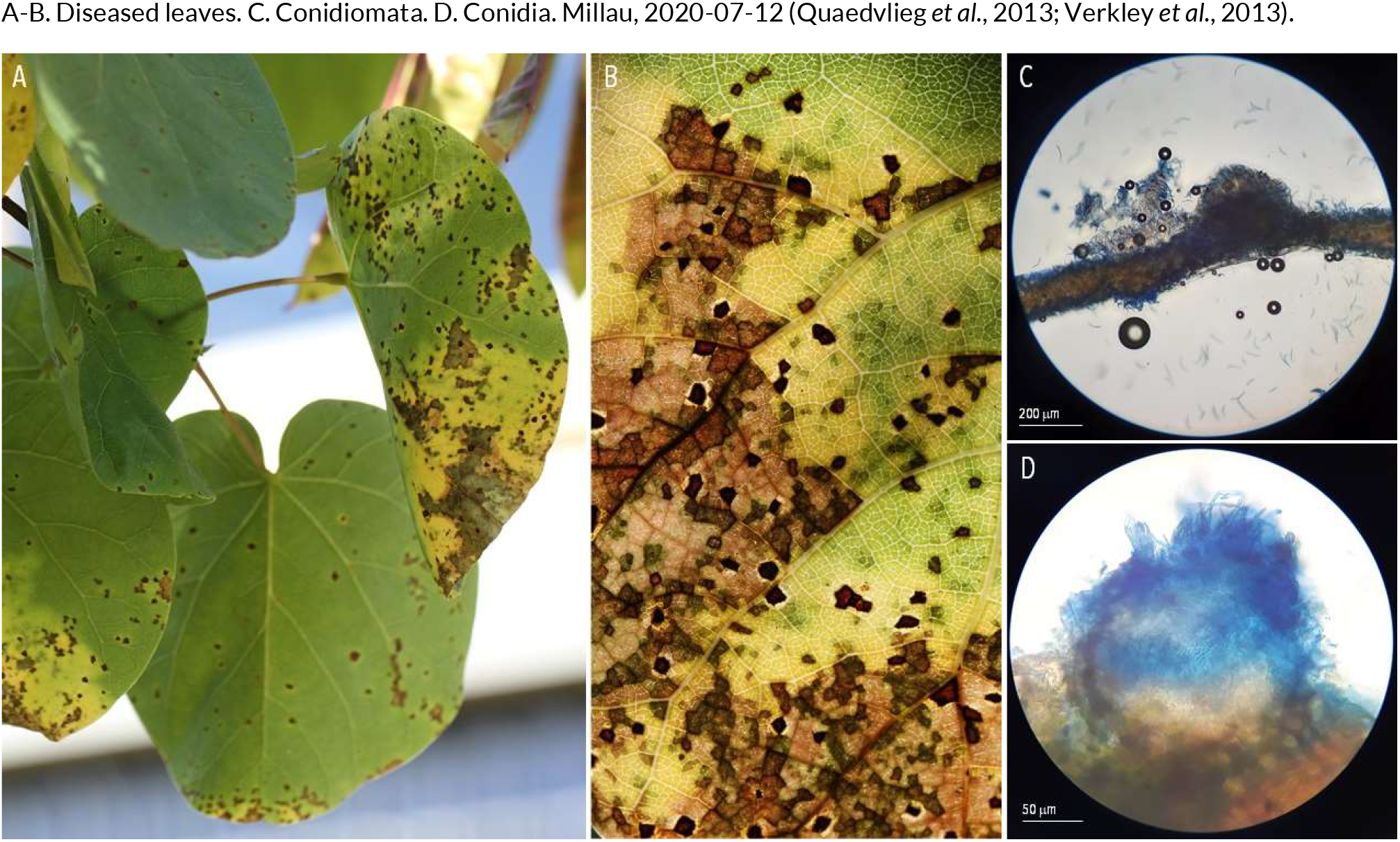
SEPTORIA-LIKE DISEASES | *Sphaerulina cercidis* (syn. Septoria provencialis) ex *Cercis siliquastrum* (Judas tree)

**150.**
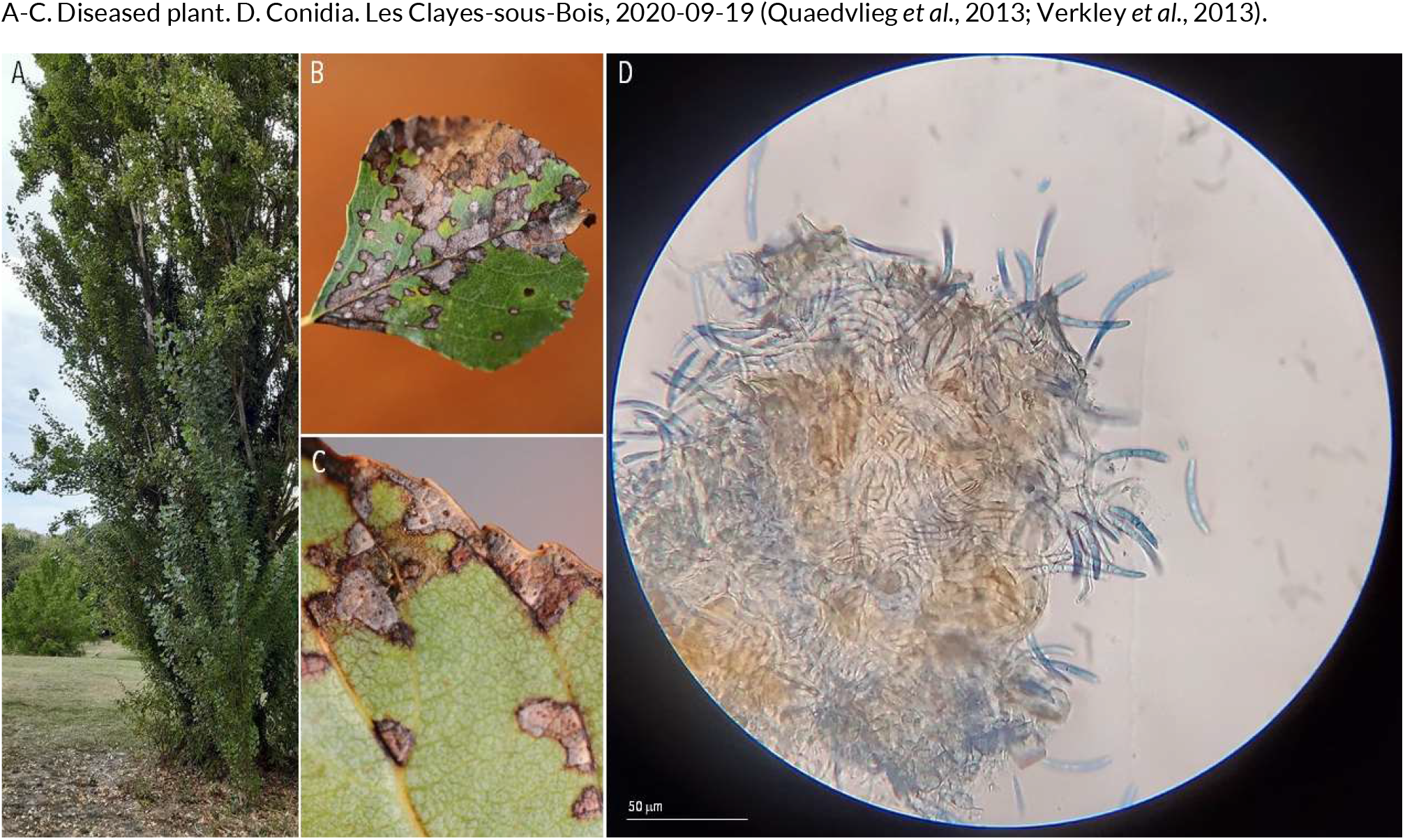
SEPTORIA-LIKE DISEASES | *Sphaerulina frondicola* (syn. *Septoria populi*) ex *Populus nigra* (black poplar)

**151.**
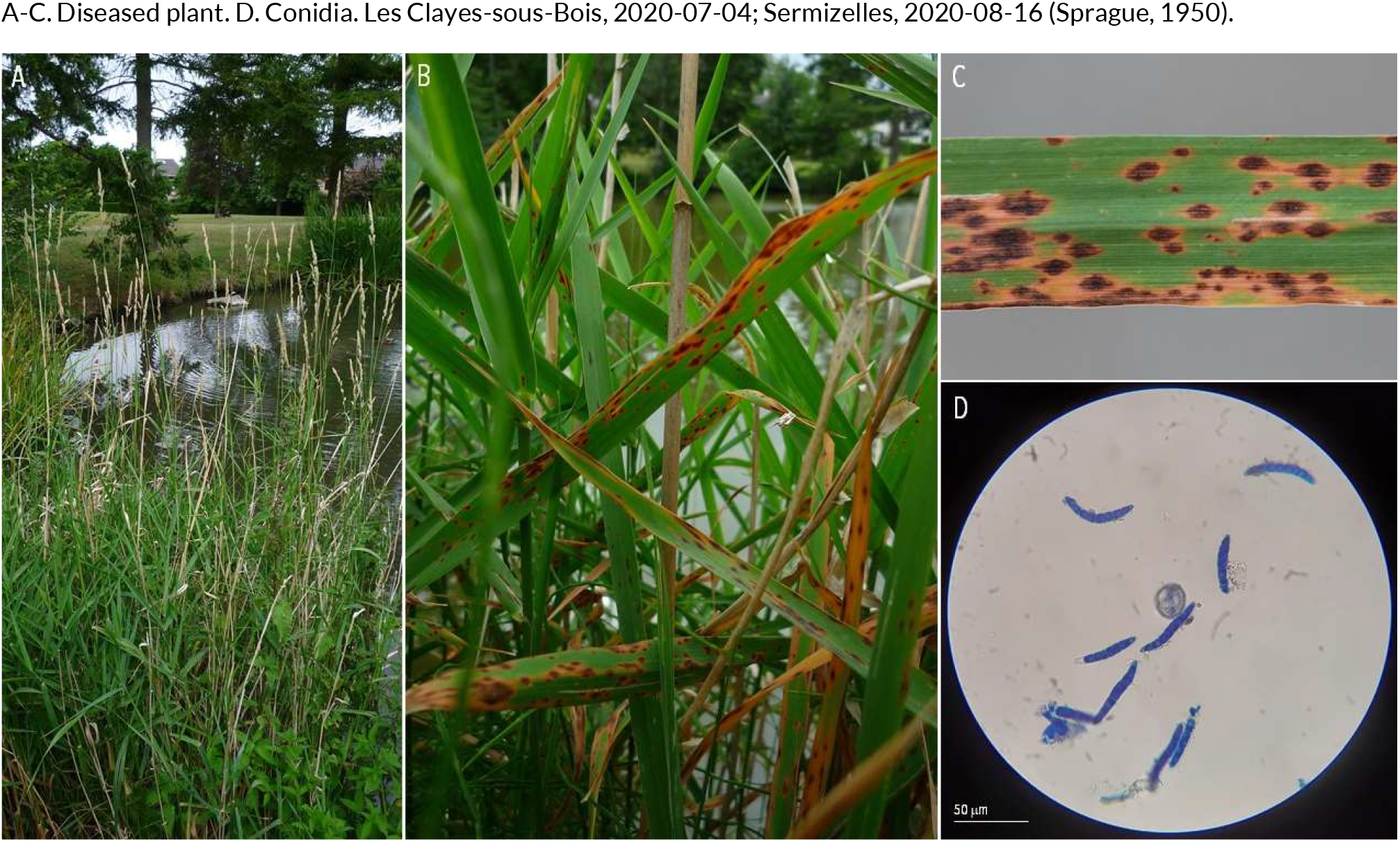
SEPTORIA-LIKE DISEASES | *Stagonospora foliicola* ex *Phalaris arundinacea* (reed canarygrass)

**152.**
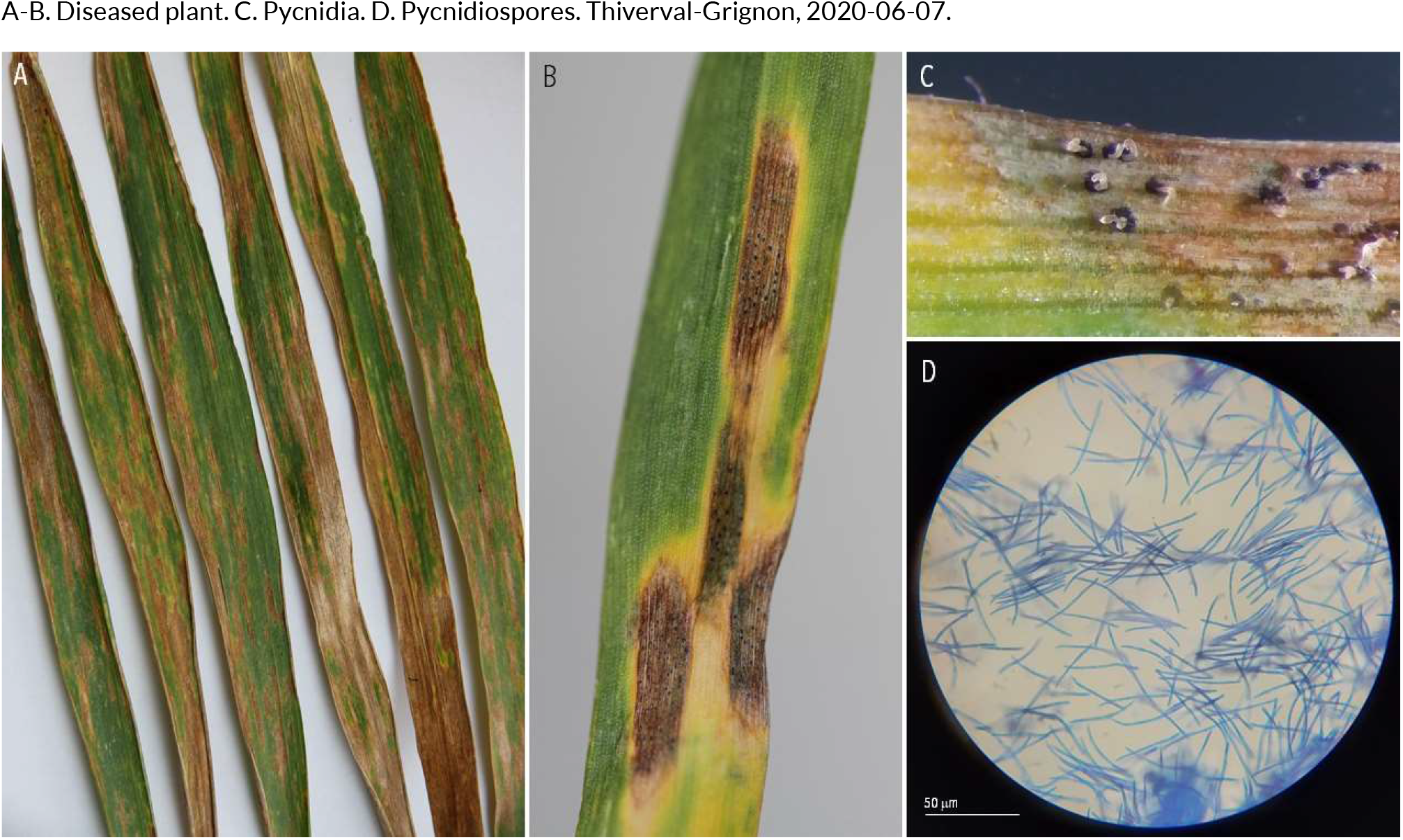
SEPTORIA-LIKE DISEASES | *Zymoseptoria tritici* ex *Triticum aestivum* (bread wheat.

**153.**
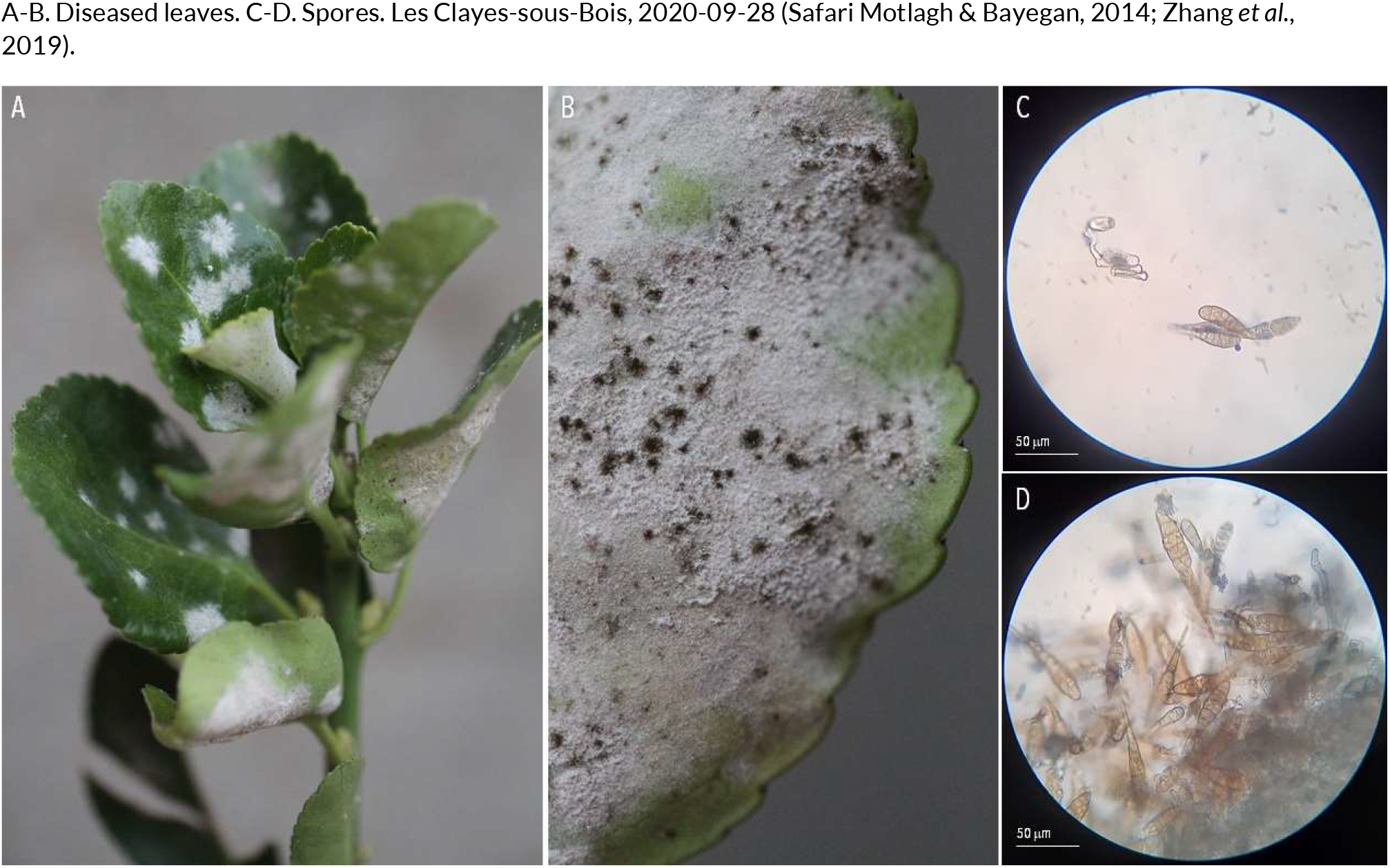
OTHER FUNGAL LEAF DISEASES | *Alternaria tenuissima* (?) ex *Euonymus japonicus* (Japanese spindle)

**154.**
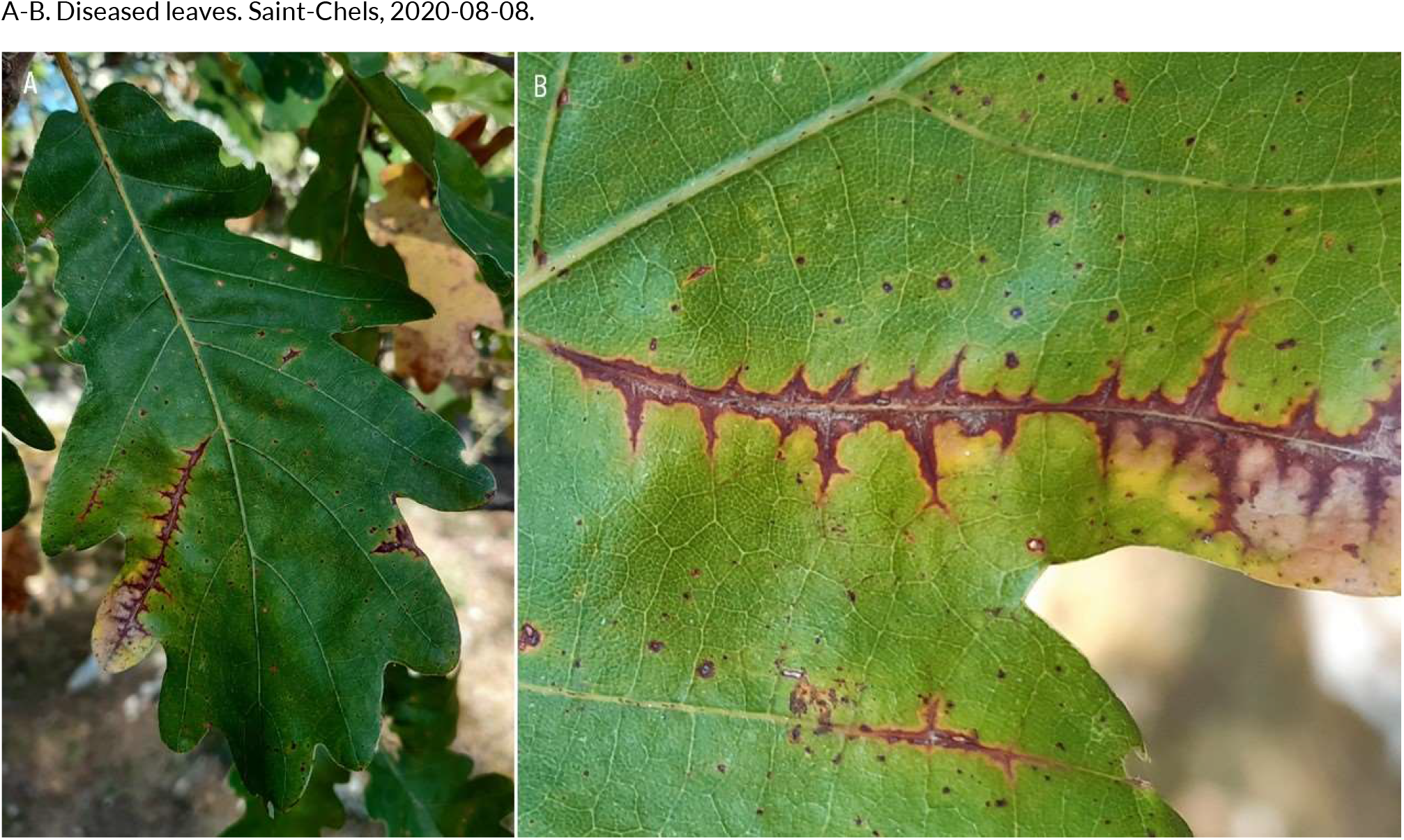
OTHER FUNGAL LEAF DISEASES | *Apiognomonia errabunda* ex *Querçus robur* (pedonculate oak)

**155.**
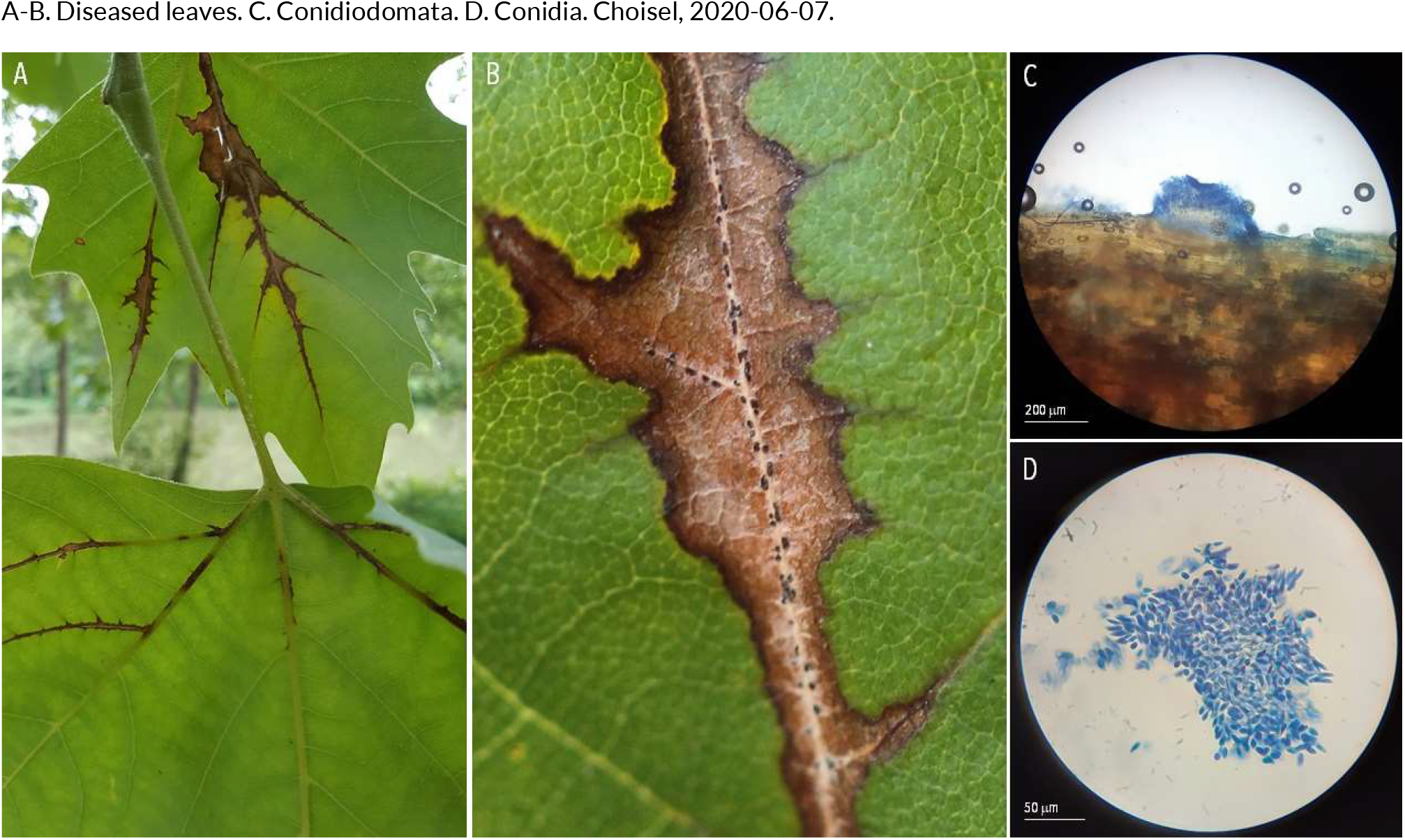
OTHER FUNGAL LEAF DISEASES | *Apiognomonia veneta* ex *Platanus x hispanica* (London plane)

**156.**
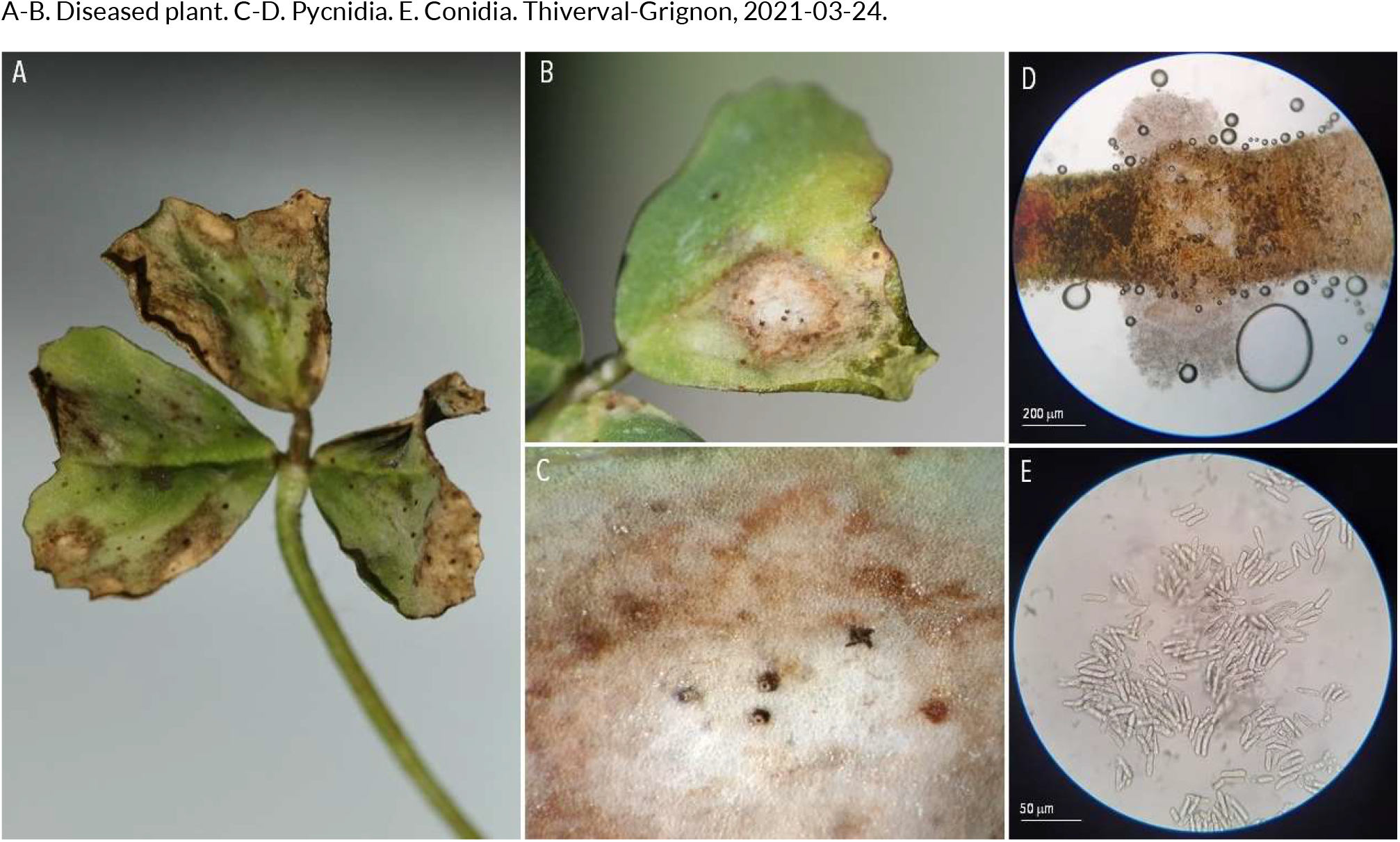
OTHER FUNGAL LEAF DISEASES | *Ascochyta trifolii* ex *Trifolium repens* (white clover)

**157.**
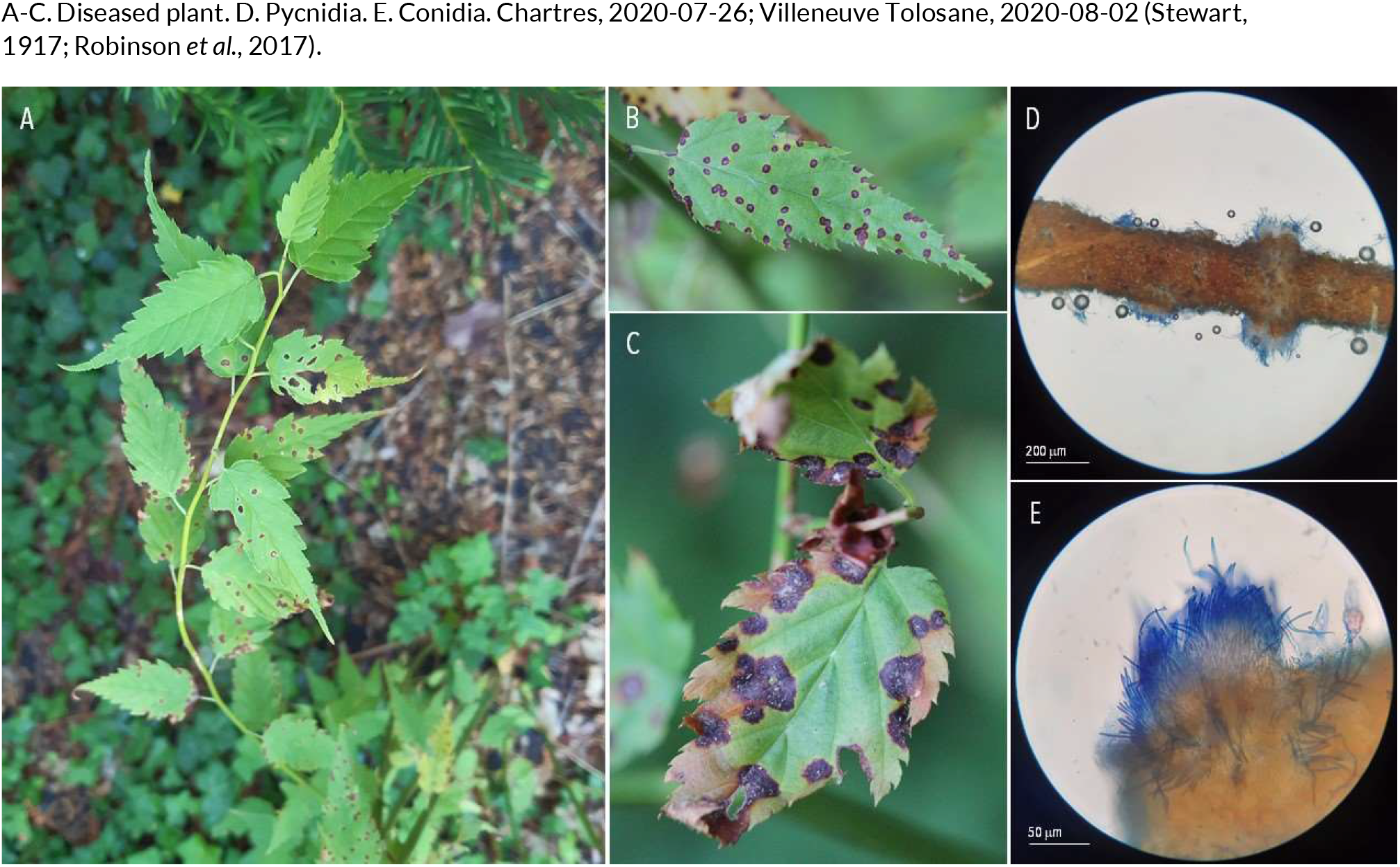
OTHER FUNGAL LEAF DISEASES | *Blumeriella kerriae* ex *Kerria japonica* (Japanese marigold bush)

**158.**
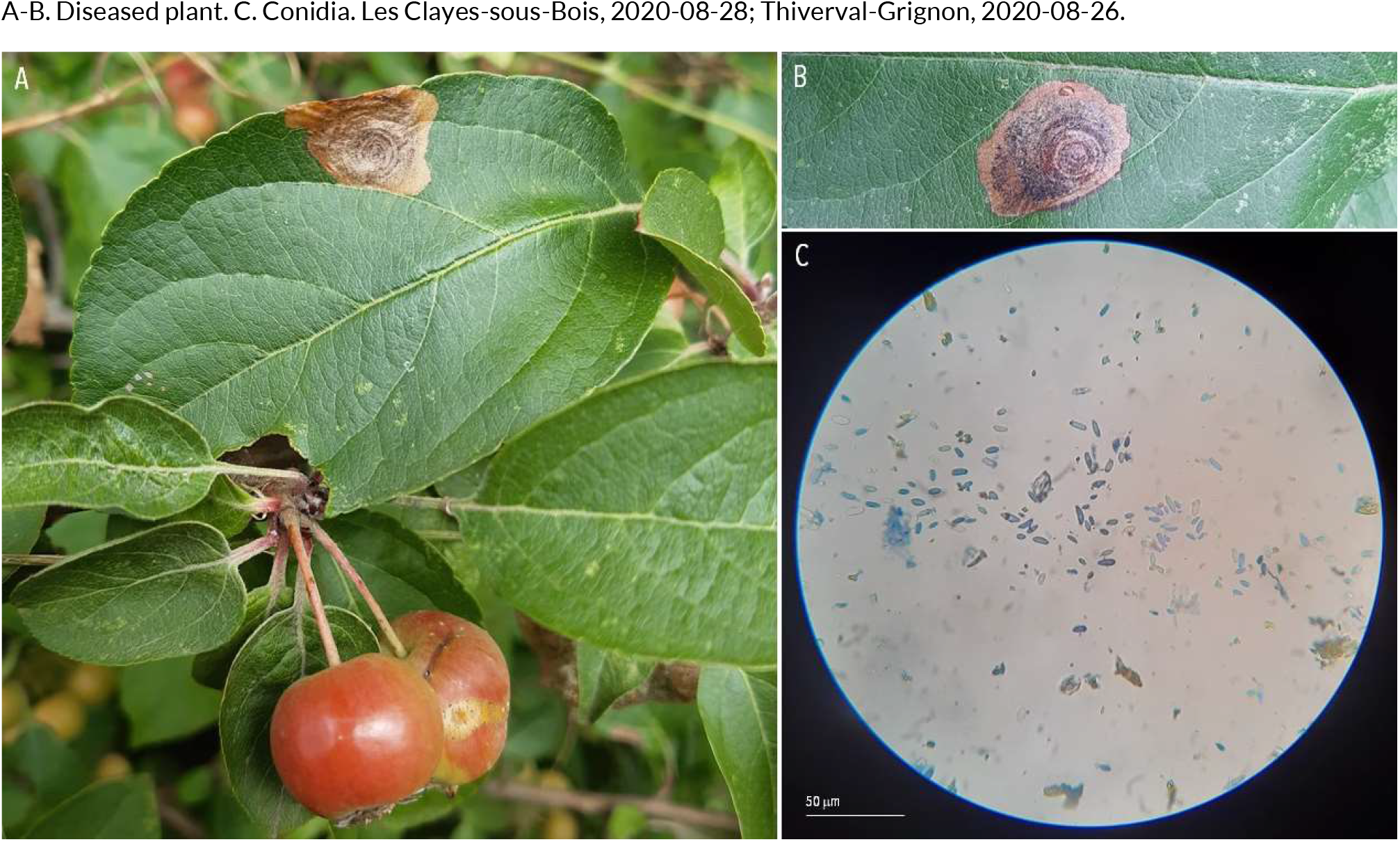
OTHER FUNGAL LEAF DISEASES | *Botryosphaeria obtusa* ex *Malus sylvestris* (European crab)

**159.**
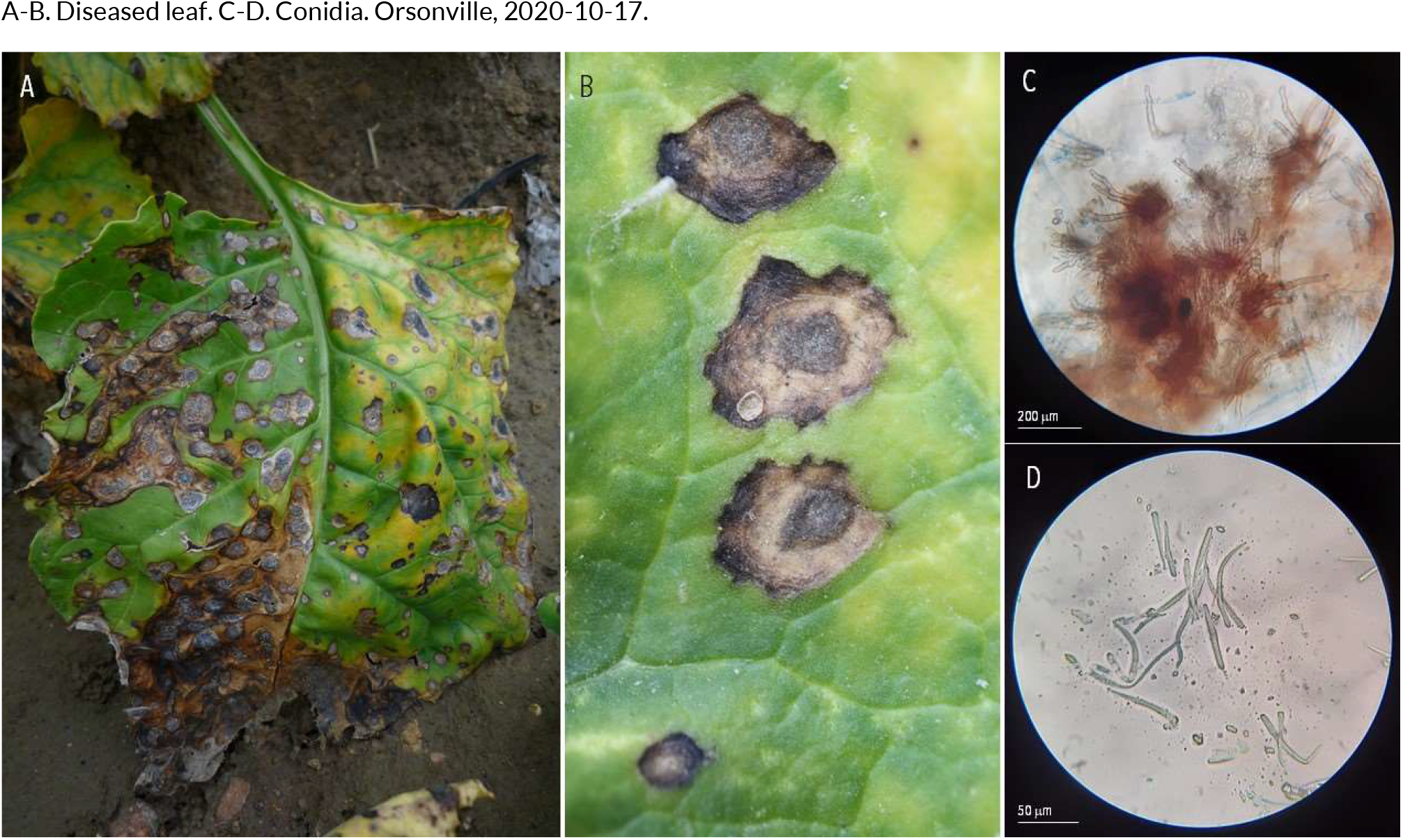
OTHER FUNGAL LEAF DISEASES | *Cercospora beticola* ex *Beta vulgaris* (beet)

**160.**
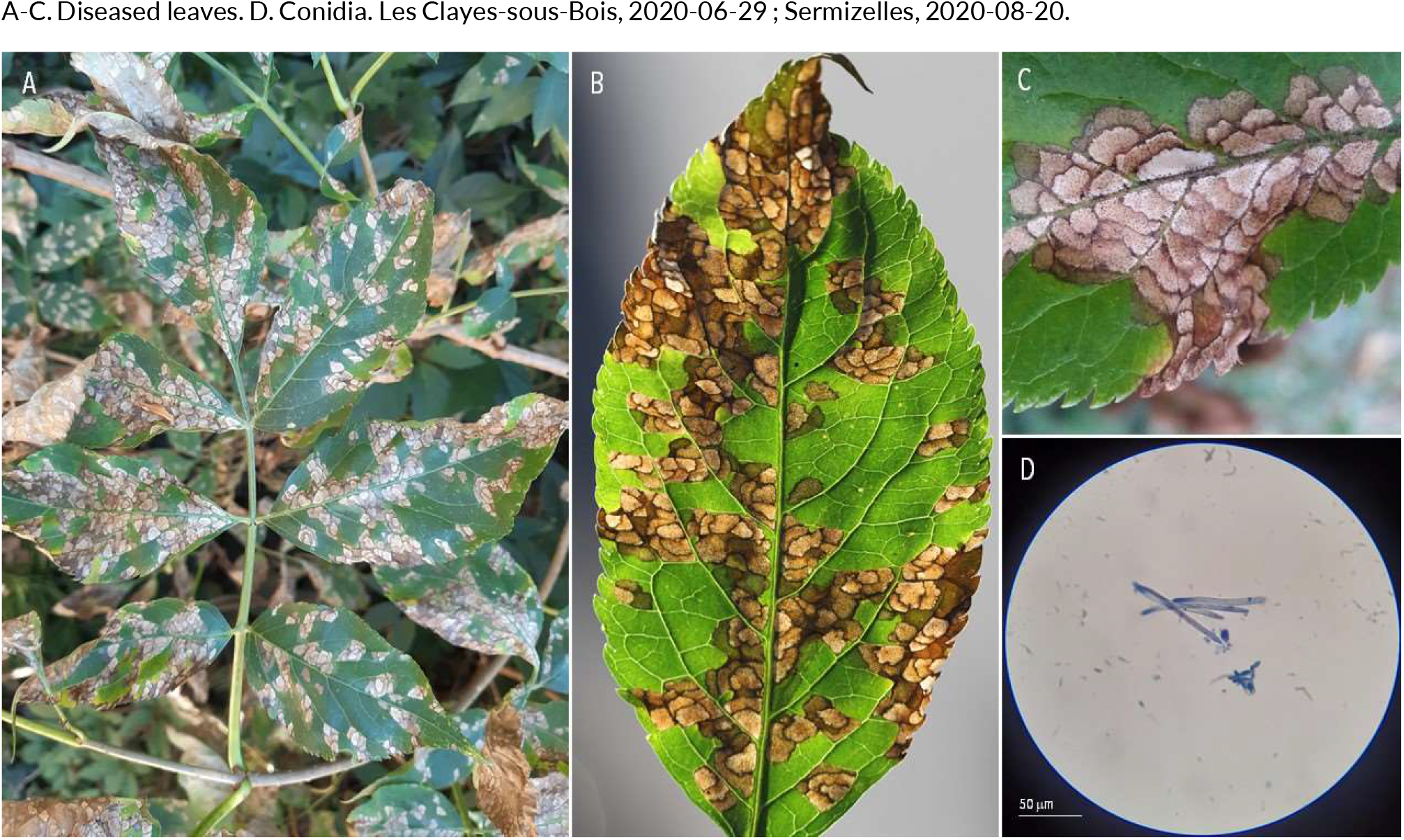
OTHER FUNGAL LEAF DISEASES | *Cercospora depazeoides* ex *Sambucus nigra* (black elder)

**161.**
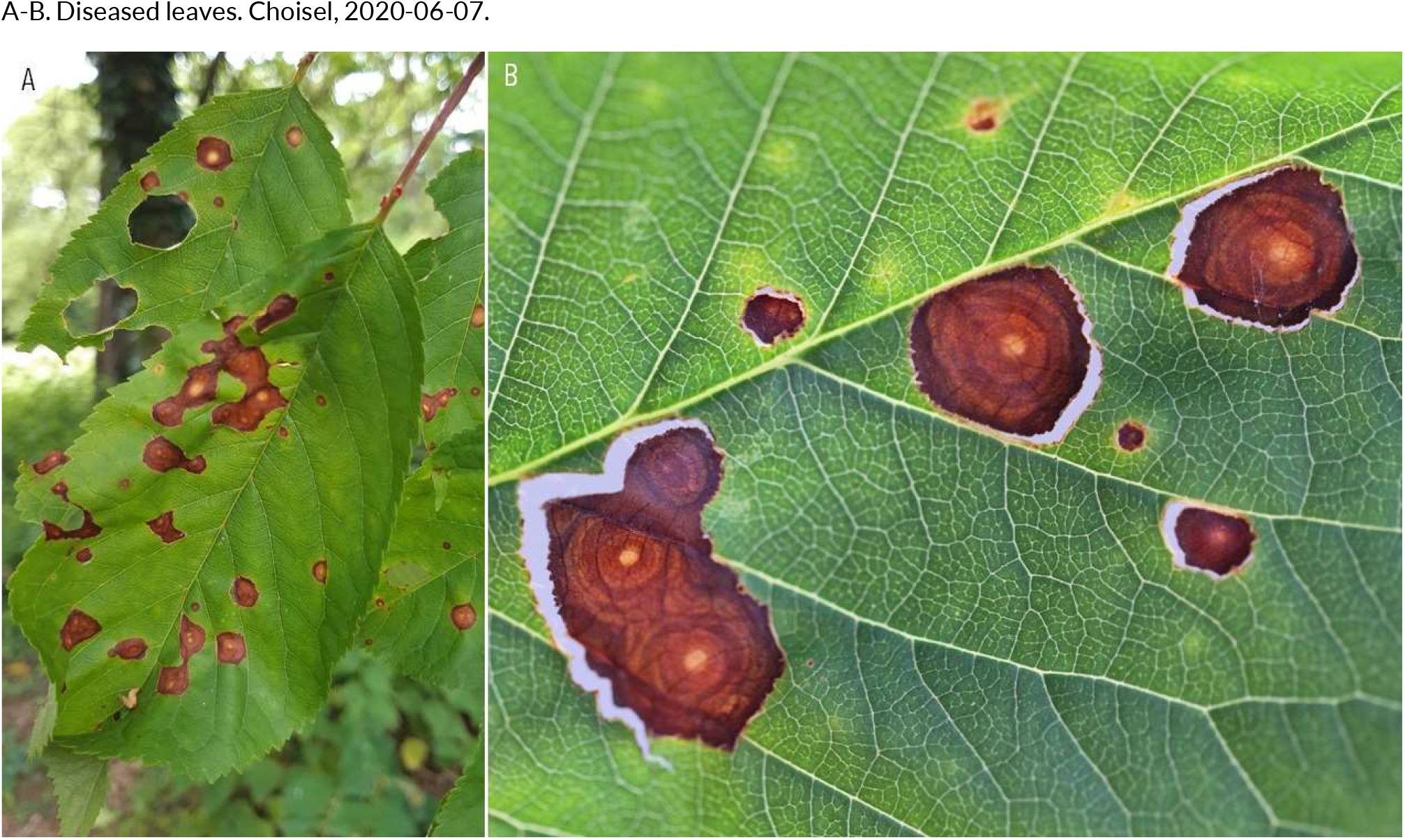
OTHER FUNGAL LEAF DISEASES | *Coryneum beijerinckii* ex *Prunus cerasus* (sour cherry)

**162.**
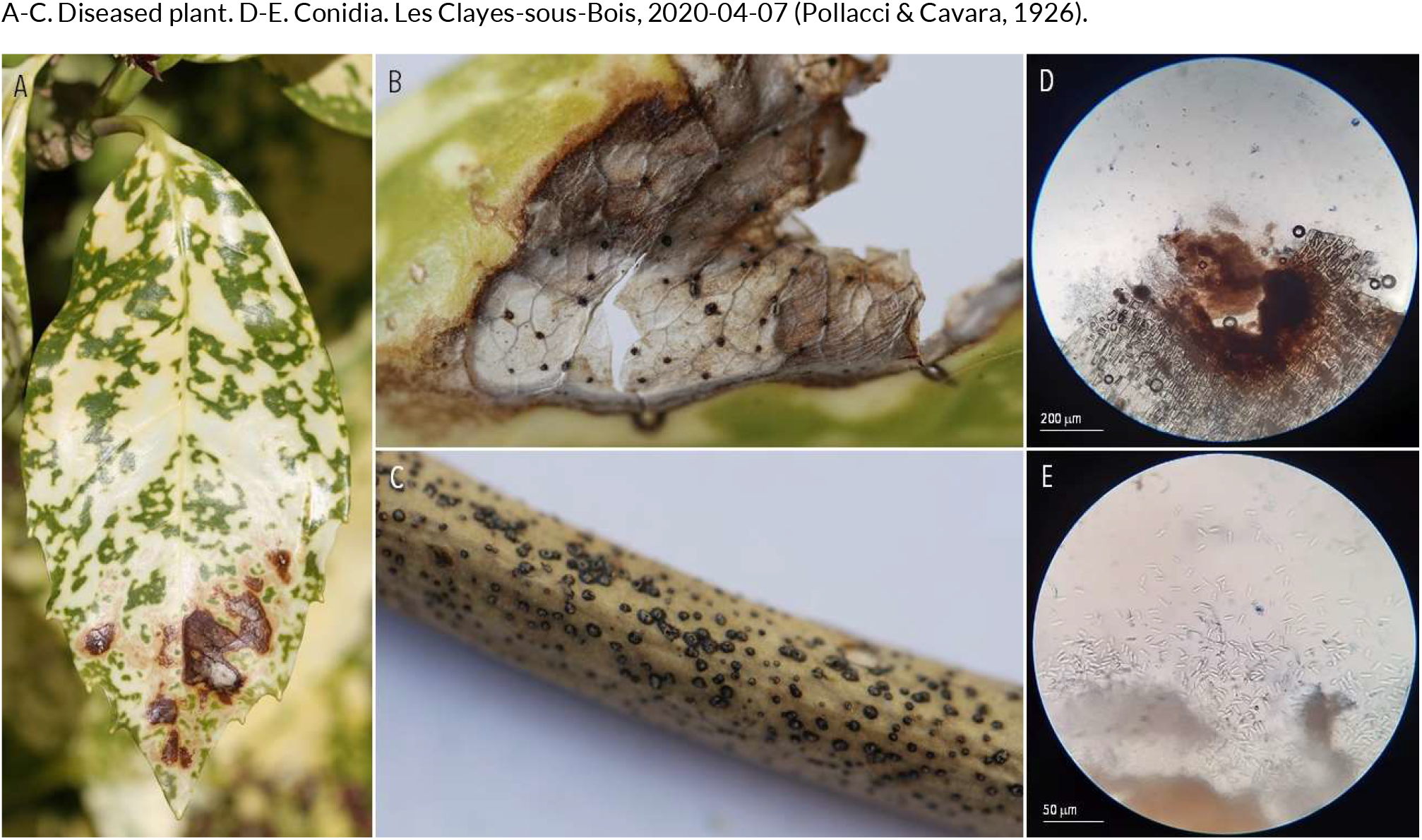
OTHER FUNGAL LEAF DISEASES | *Diaporthe aucubae* ex *Aucuba japonica* (spotted laurel)

**163.**
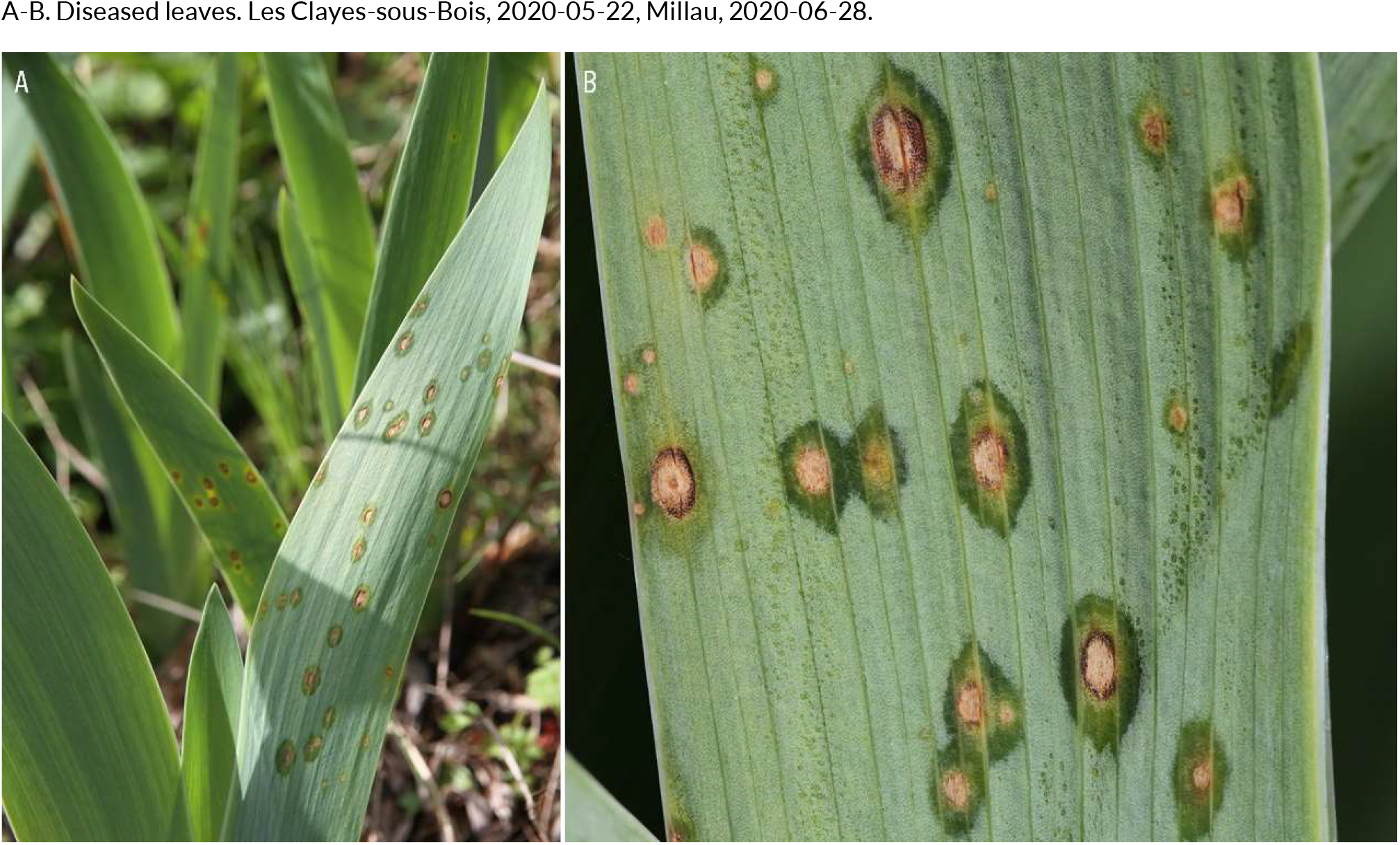
OTHER FUNGAL LEAF DISEASES | *Didymellina macrospora* ex *Iris germanica* (bearded iris)

**164.**
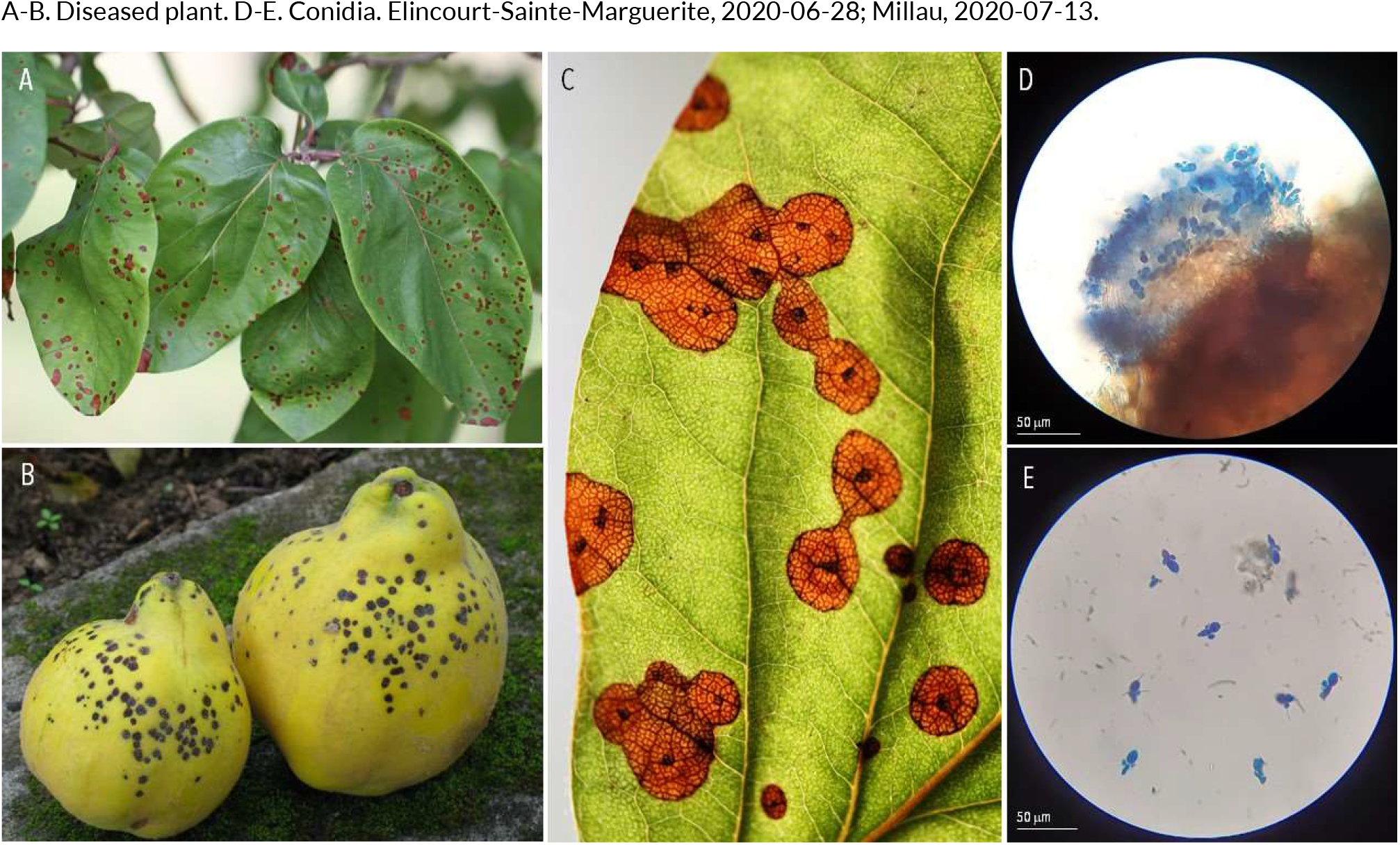
OTHER FUNGAL LEAF DISEASES | *Diplocarpon mespili* ex *Cydonia oblonga* (quince)

**165.**
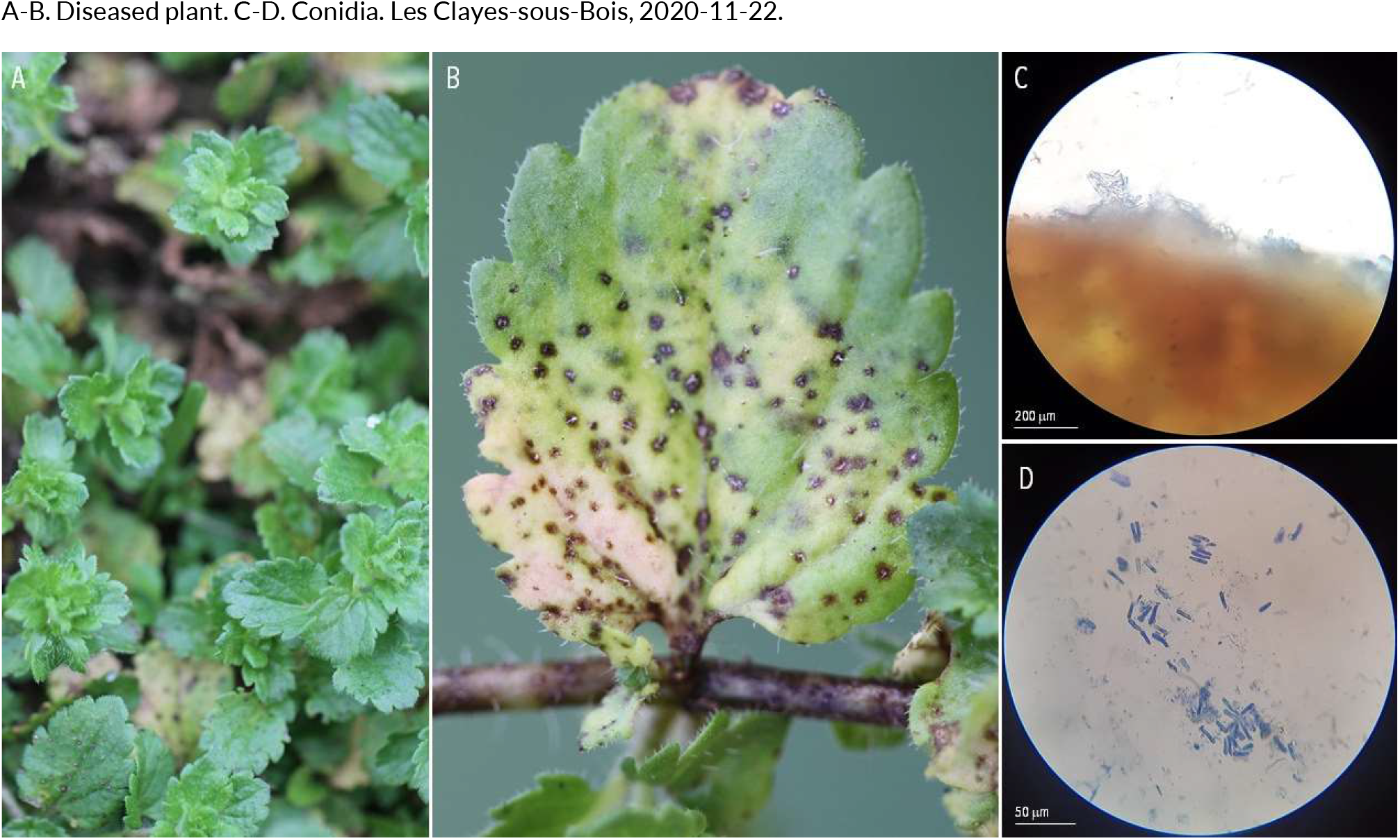
OTHER FUNGAL LEAF DISEASES | *Discogloeum veronicae* ex *Veronica persica* (bird’s eye)

**166.**
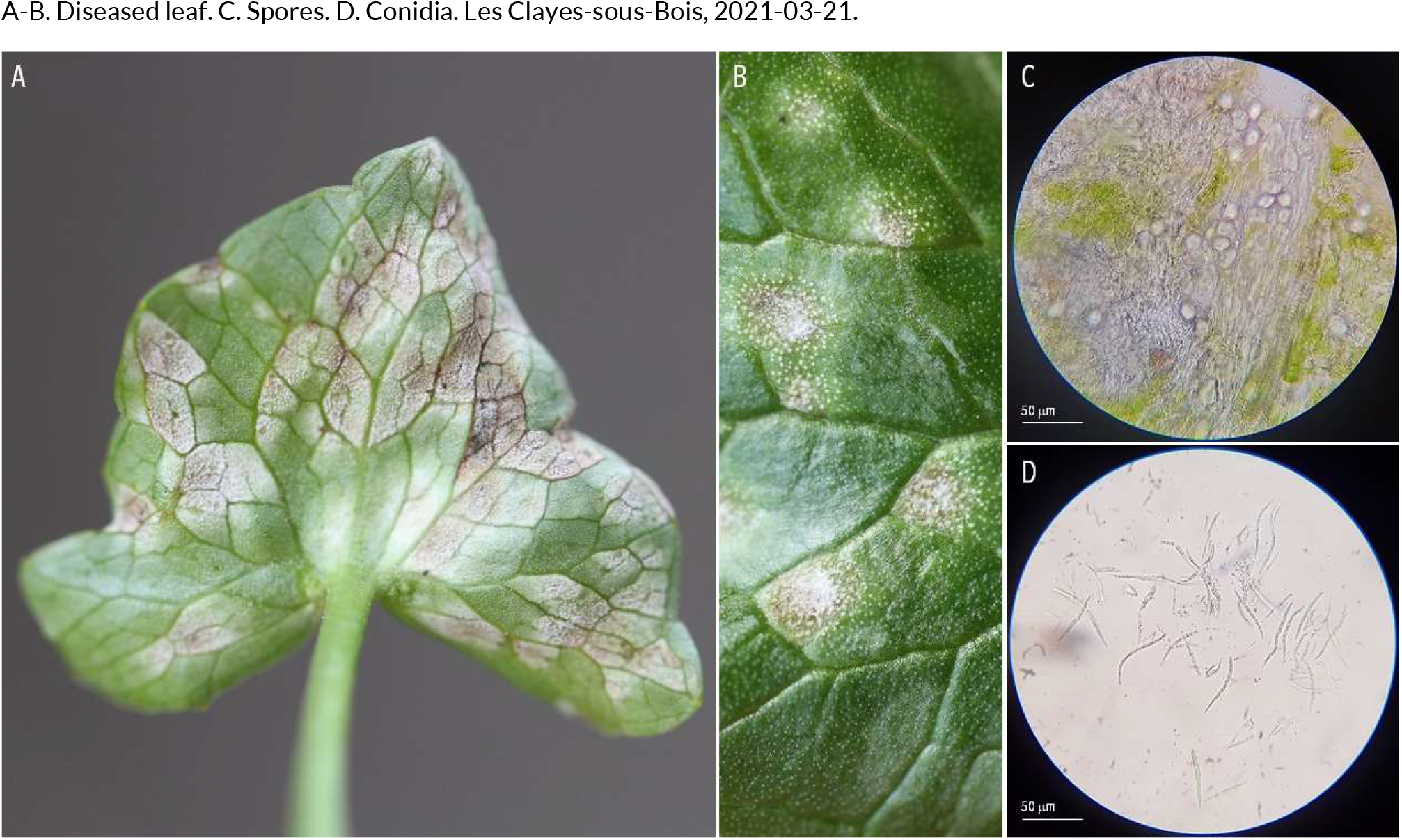
OTHER FUNGAL LEAF DISEASES | *Entyloma ficariae* ex *Ficaria verna* (lesser celandine)

**167.**
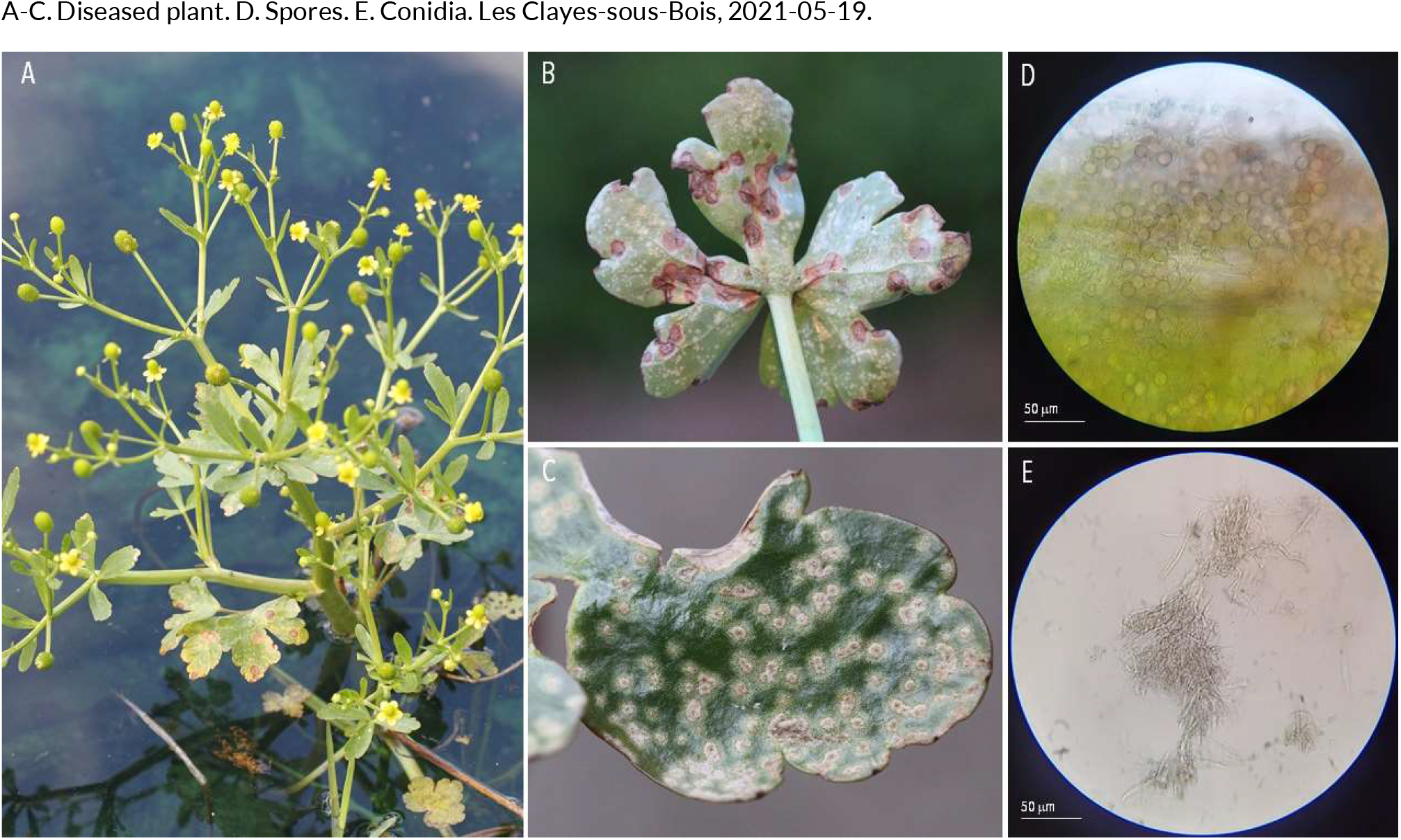
OTHER FUNGAL LEAF DISEASES | *Entyloma ranunculi-scelerati* ex *Ranunculus deceleratus* (cursed buttercup)

**168.**
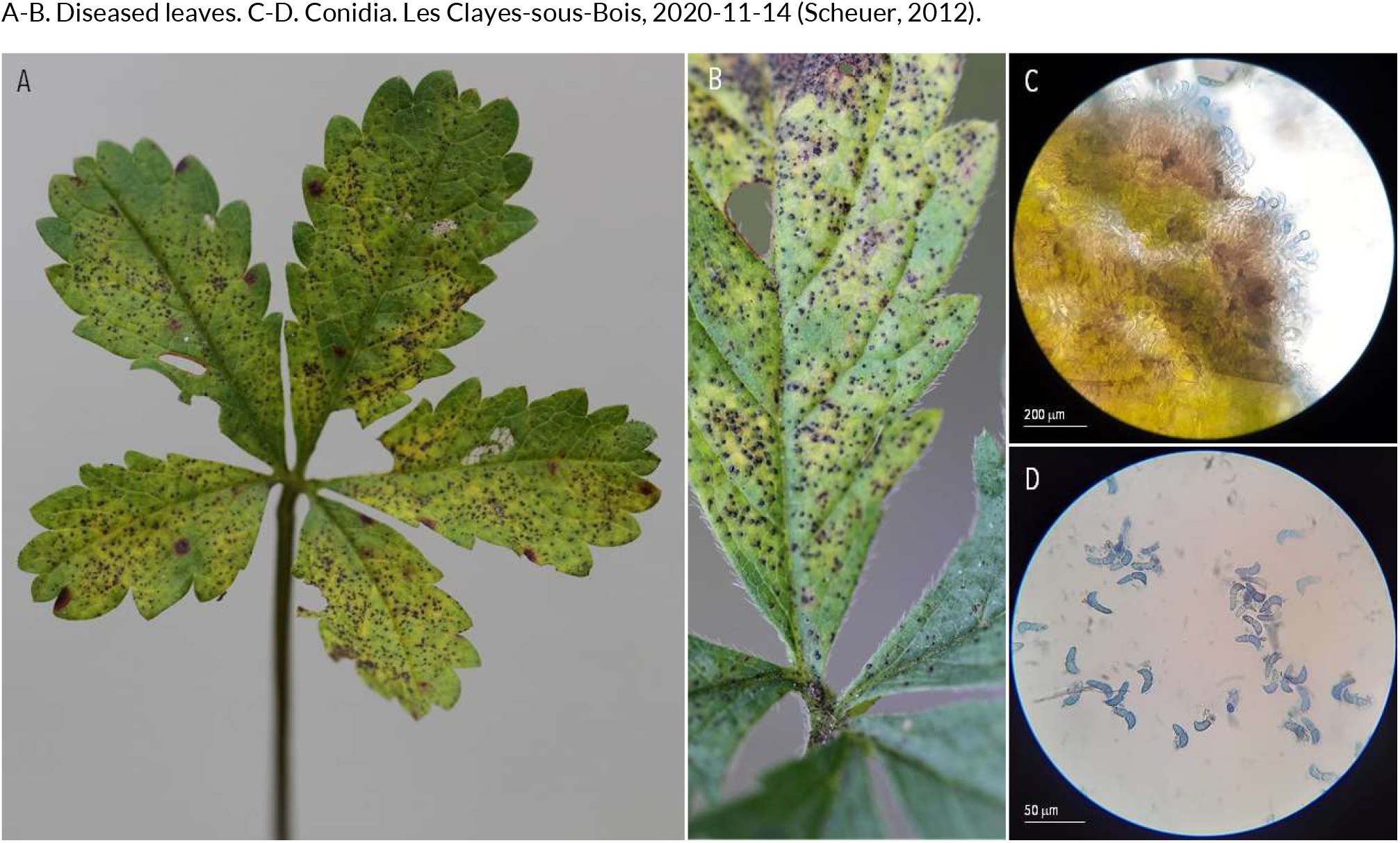
OTHER FUNGAL LEAF DISEASES | *Marssonina fragariae* ex *Potentilla reptans* (creeping cinquefoil)

**169.**
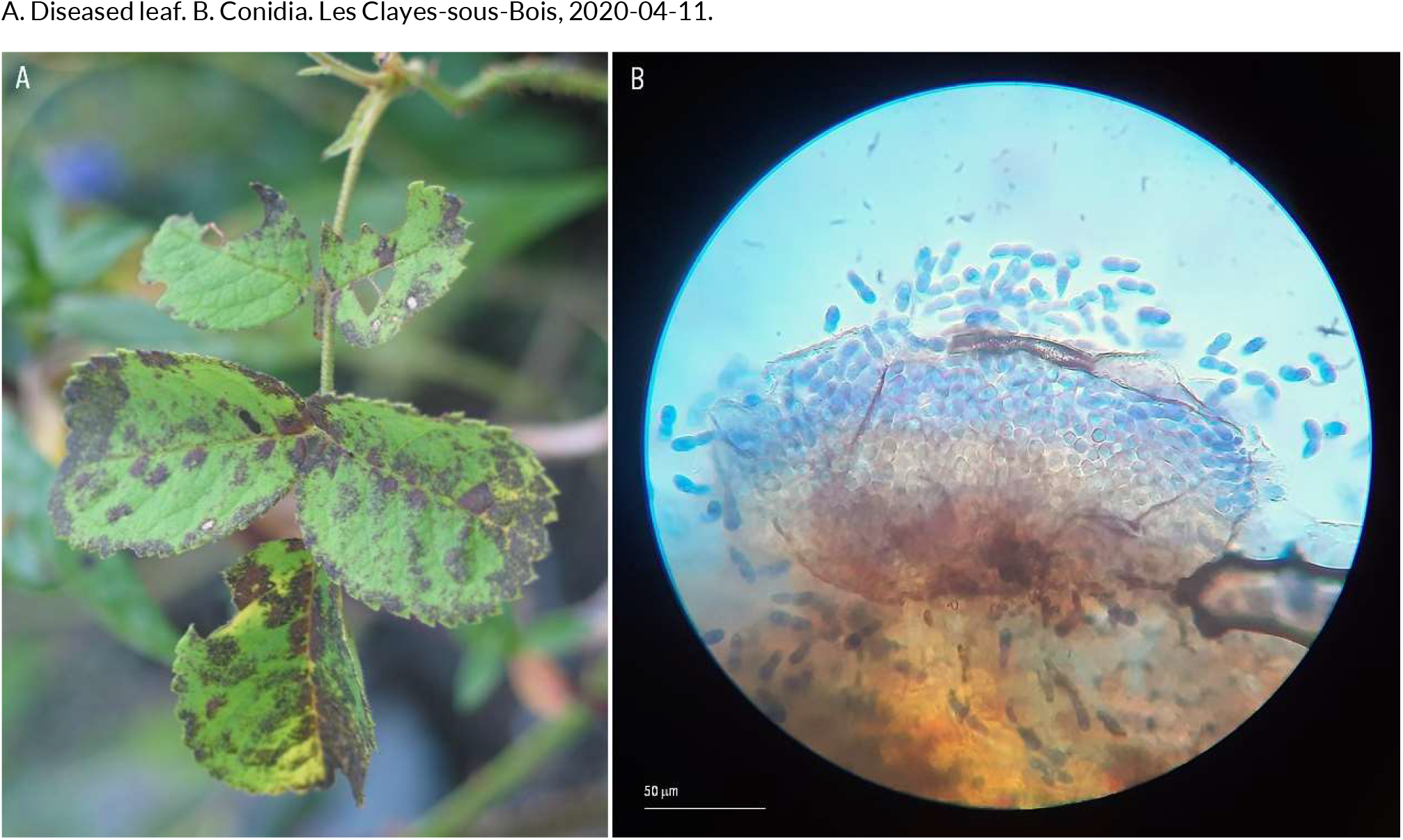
OTHER FUNGAL LEAF DISEASES | *Marssonina rosae* ex *Rosa sp*. (rose)

**170.**
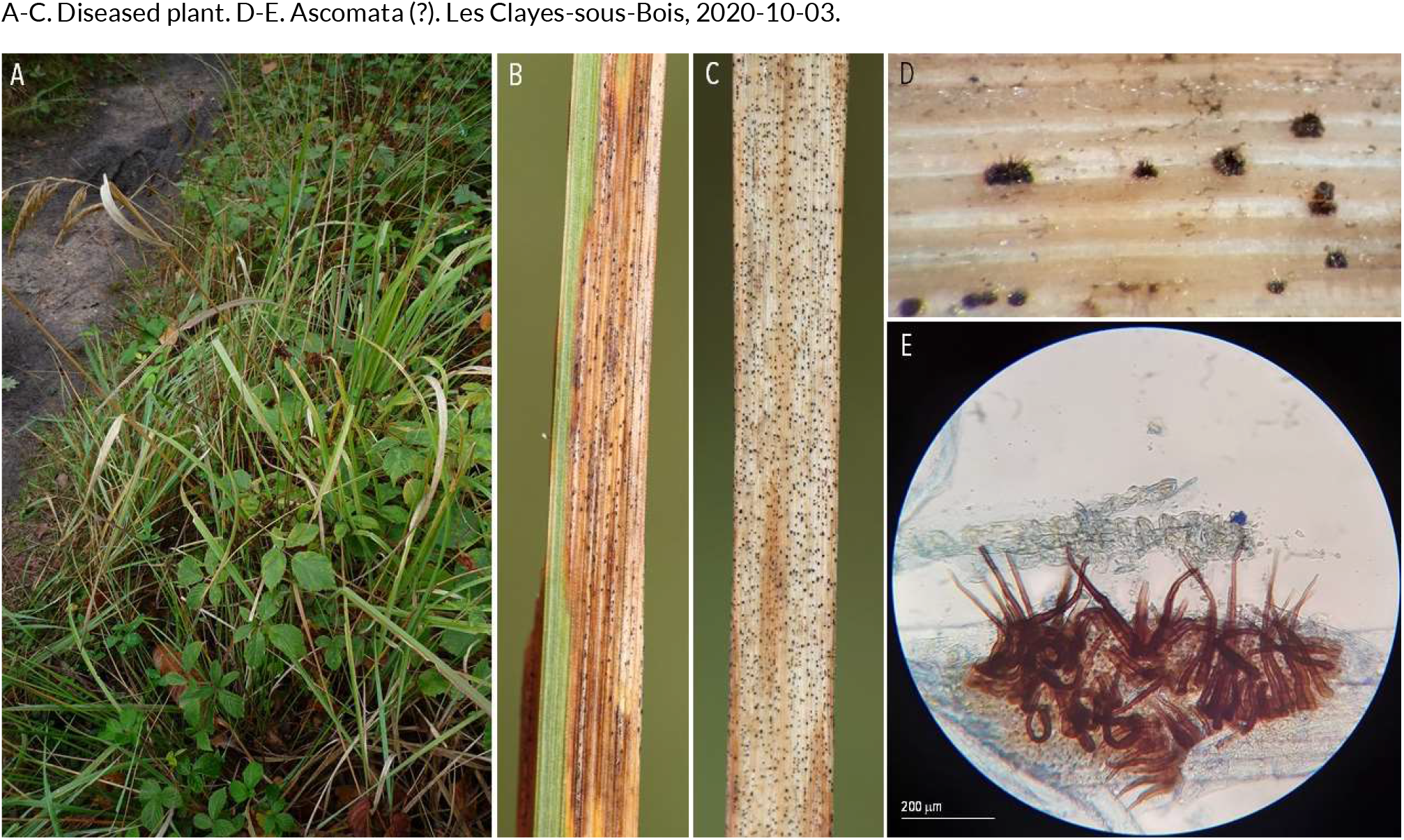
OTHER FUNGAL LEAF DISEASES | *Pyrenophora* sp. (??) ex *Calamagrostis epigejos* (bushgrass)

**171.**
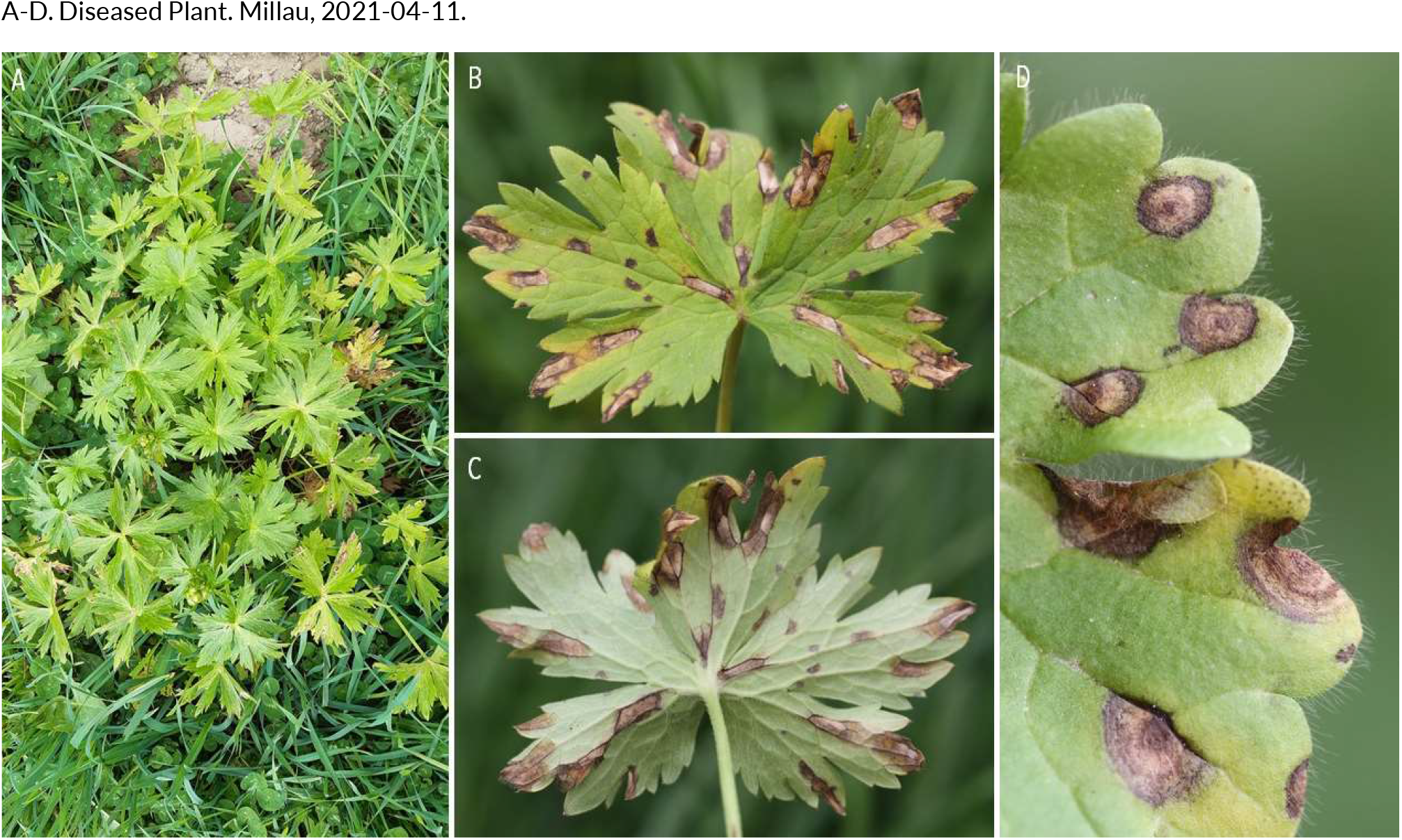
OTHER FUNGAL LEAF DISEASES | *Ramularia acris* ex *Ranunculus acris* (meadow buttercup)

**172.**
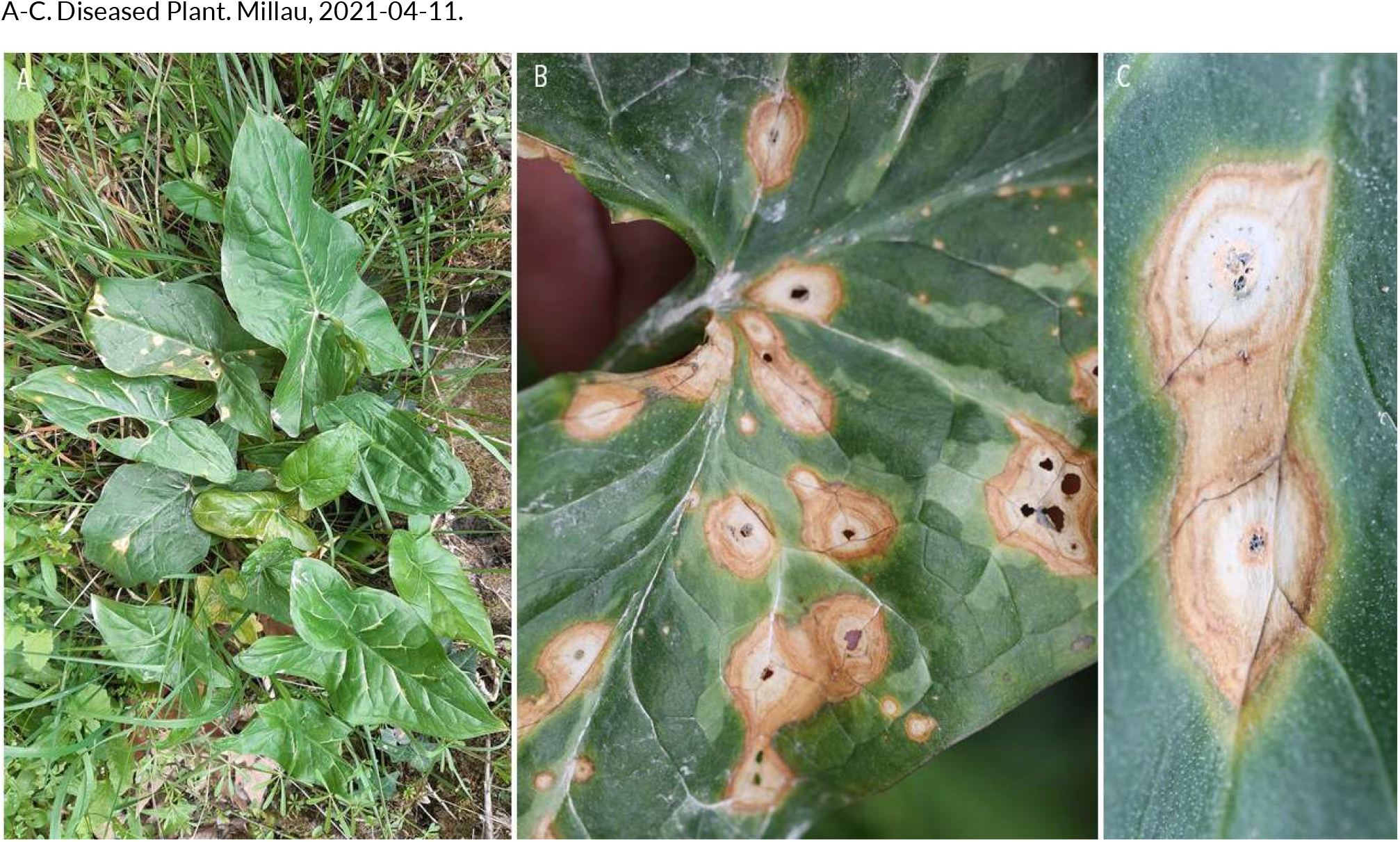
OTHER FUNGAL LEAF DISEASES | *Ramularia ari* ex *Arum maculata* (wild arum)

**173.**
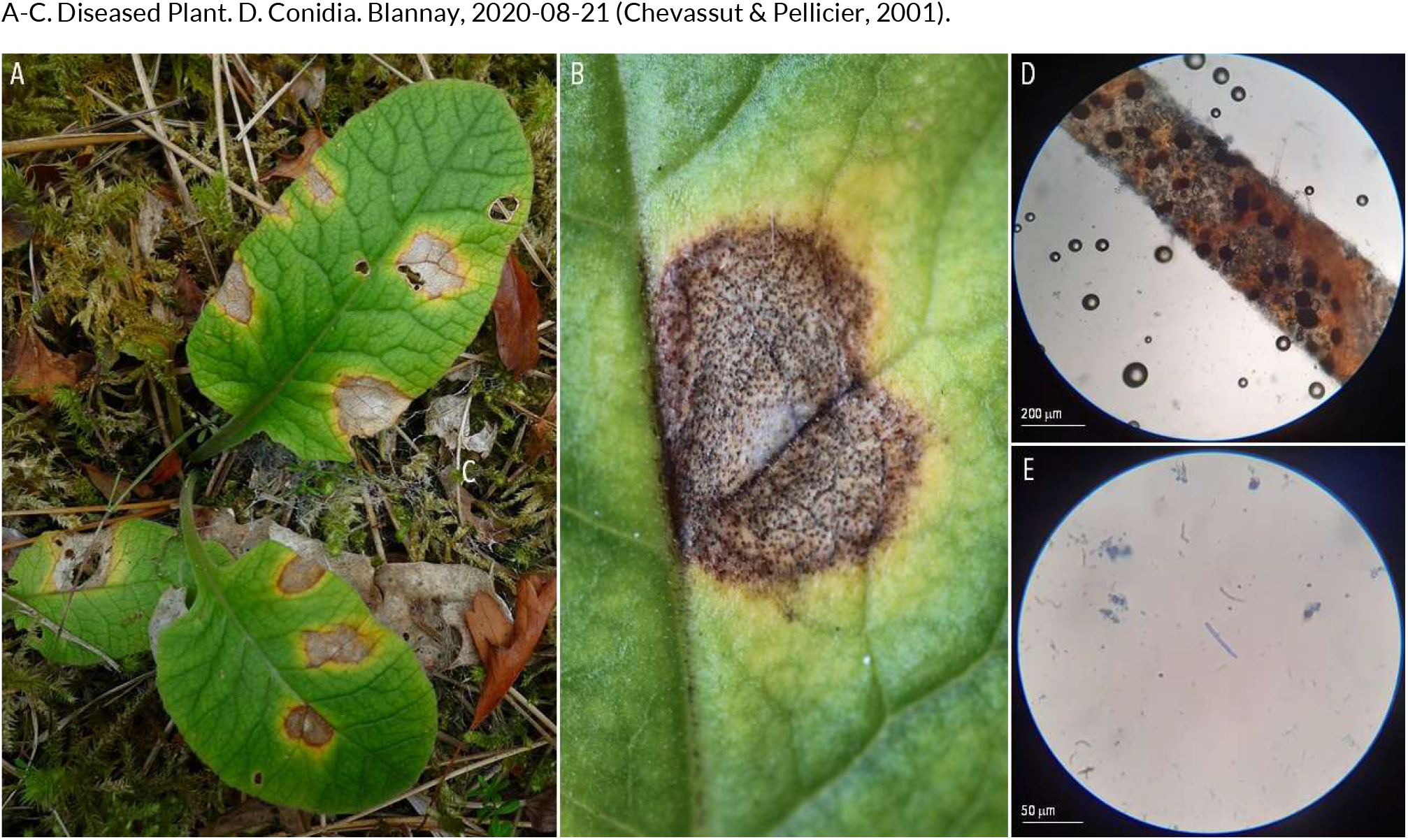
OTHER FUNGAL LEAF DISEASES | *Ramularia primulae* ex *Primula veris* (cowslip)

**174.**
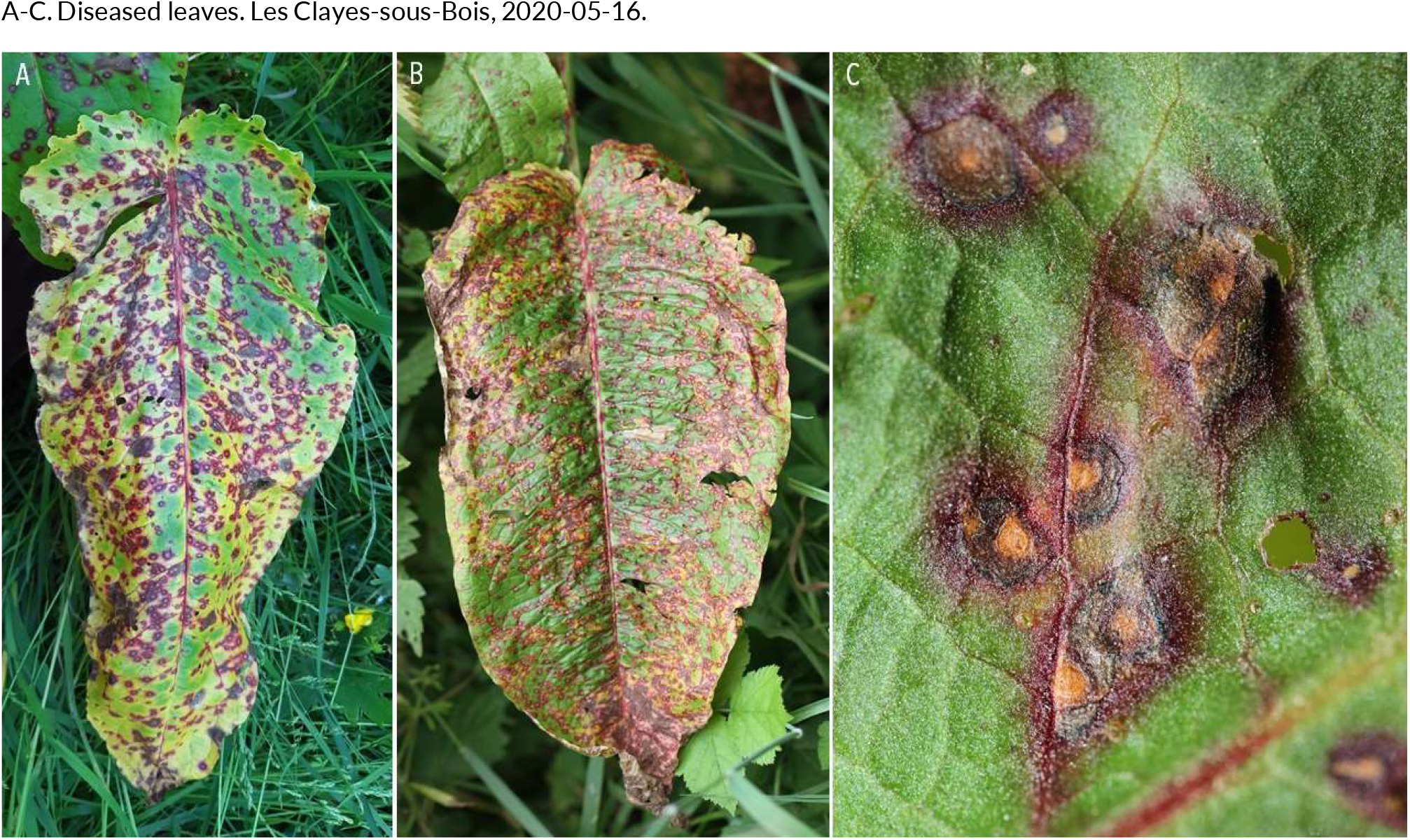
OTHER FUNGAL LEAF DISEASES | *Ramularia rubella* ex *Rumex obtusifolius* (broad-leaved dock)

**175.**
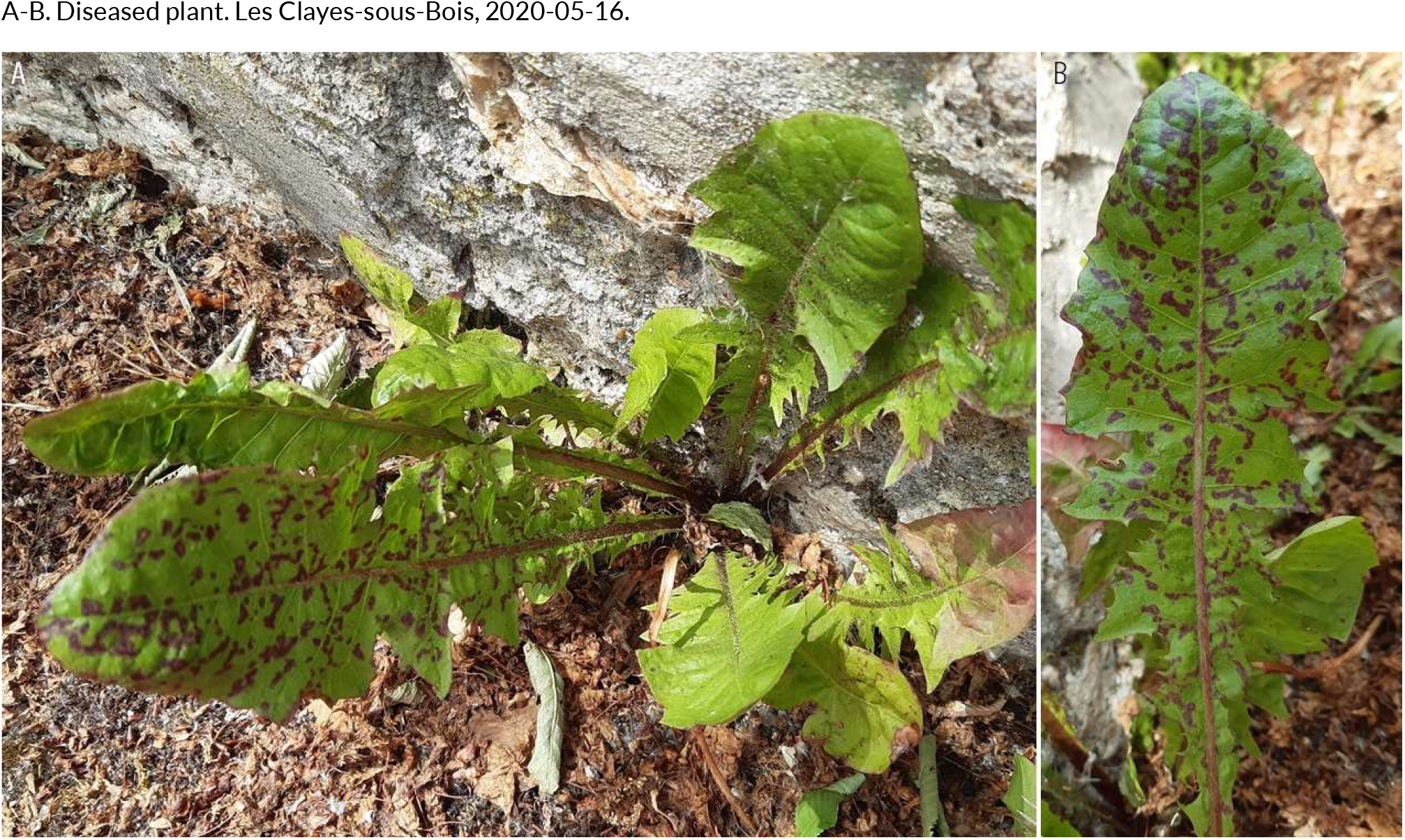
OTHER FUNGAL LEAF DISEASES | *Ramularia taraxaci* ex *Taraxacum officinale* (common dandelion)

**176.**
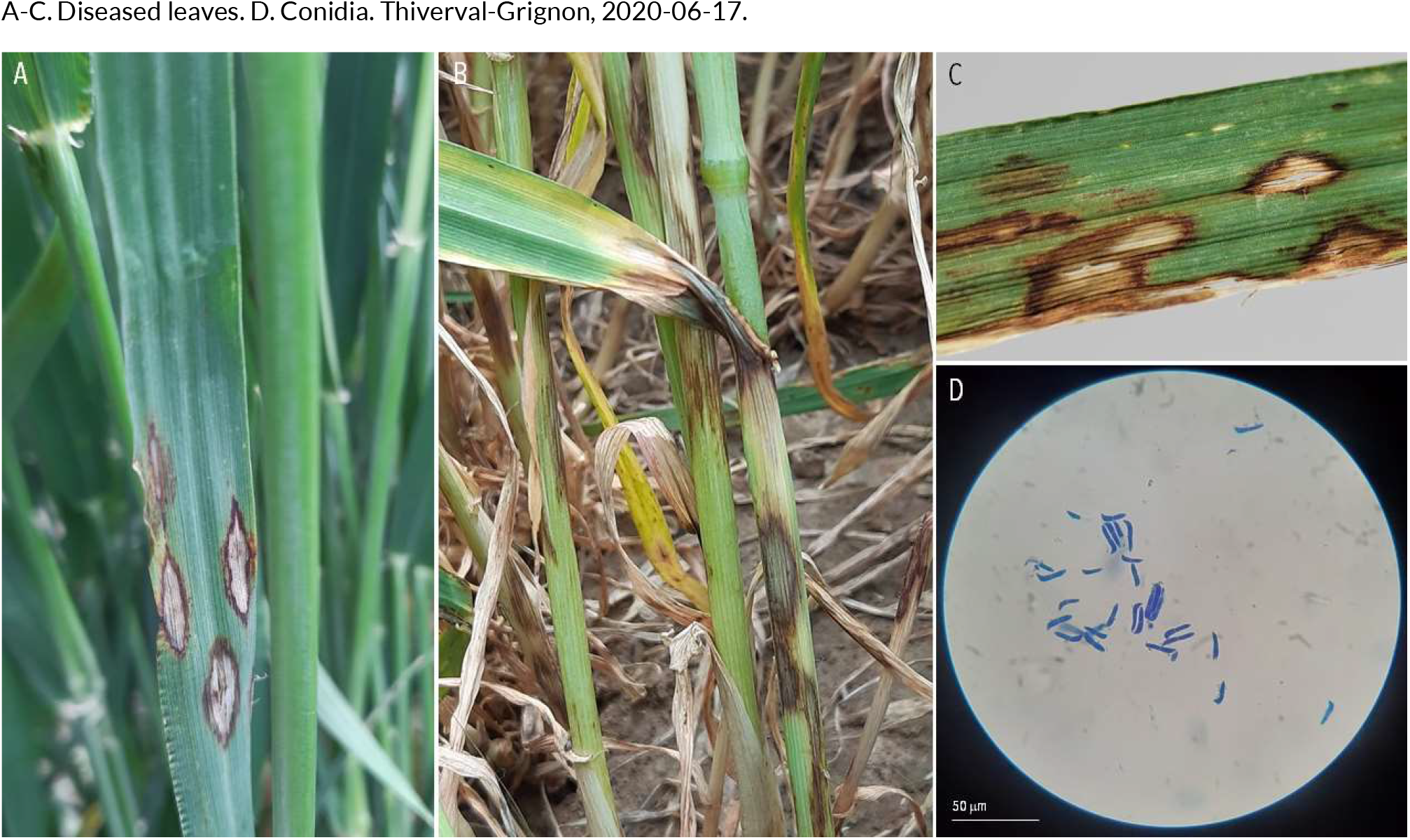
OTHER FUNGAL LEAF DISEASES | *Rhynchosporium secalis* ex *Hordeum vulgare* (barley)

**177.**
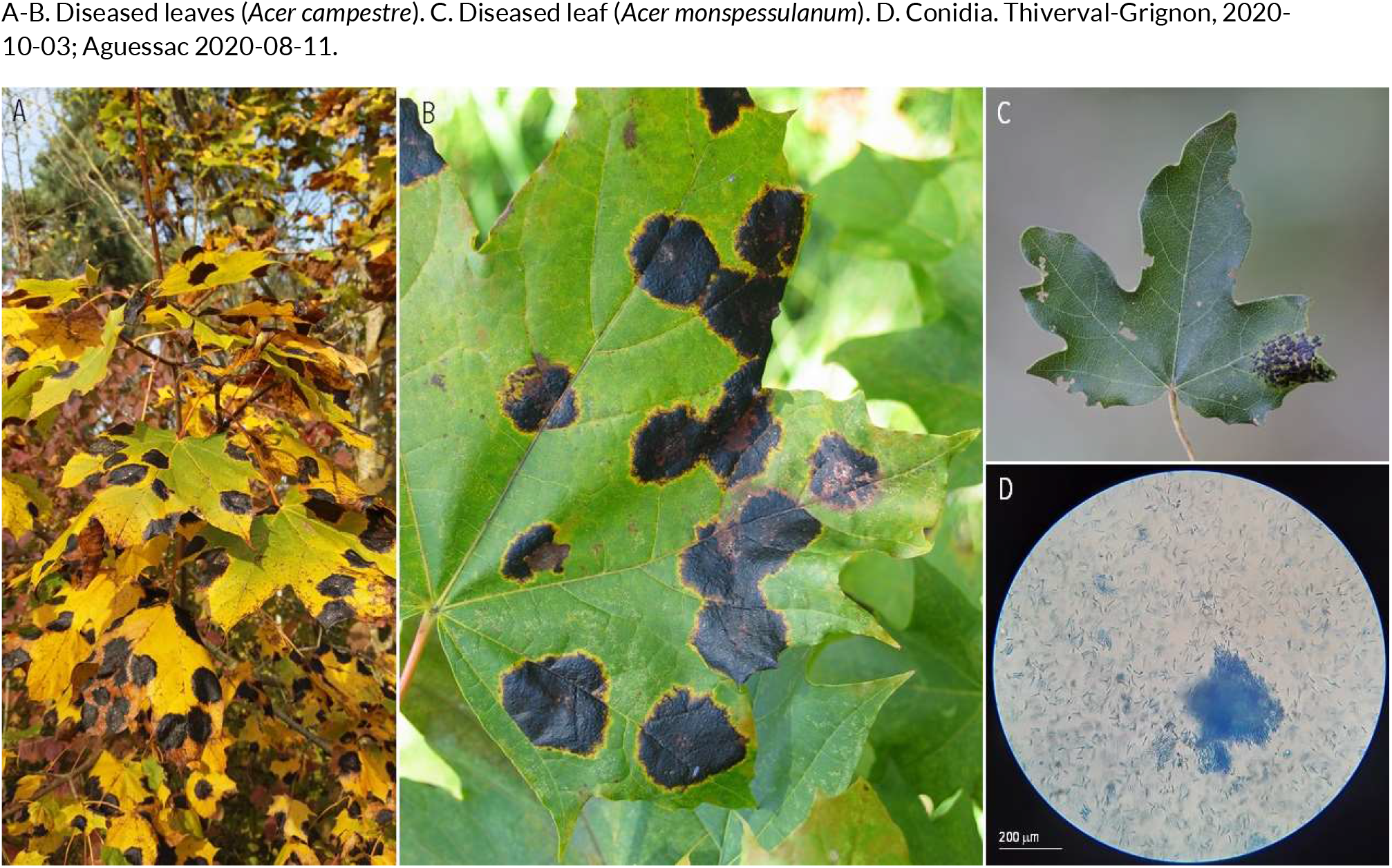
OTHER FUNGAL LEAF DISEASES | *Rhytisma acerinum* ex *Acer campestre* (field maple) and *Acer monspessulanum* (Montpellier maple)

**178.**
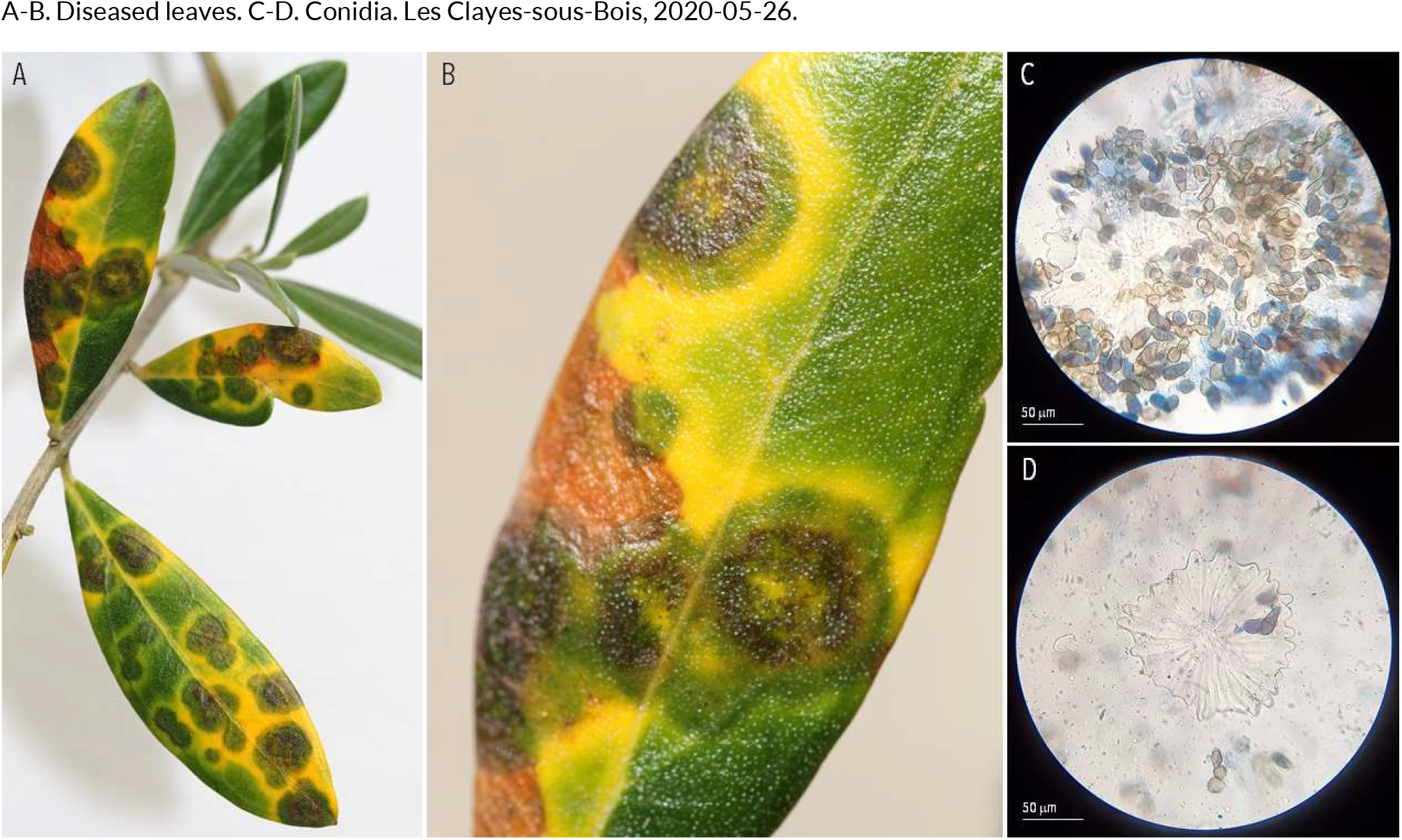
OTHER FUNGAL LEAF DISEASES | *Spilocaea oleaginea* ex *Olea europaea* (olive tree)

**179.**
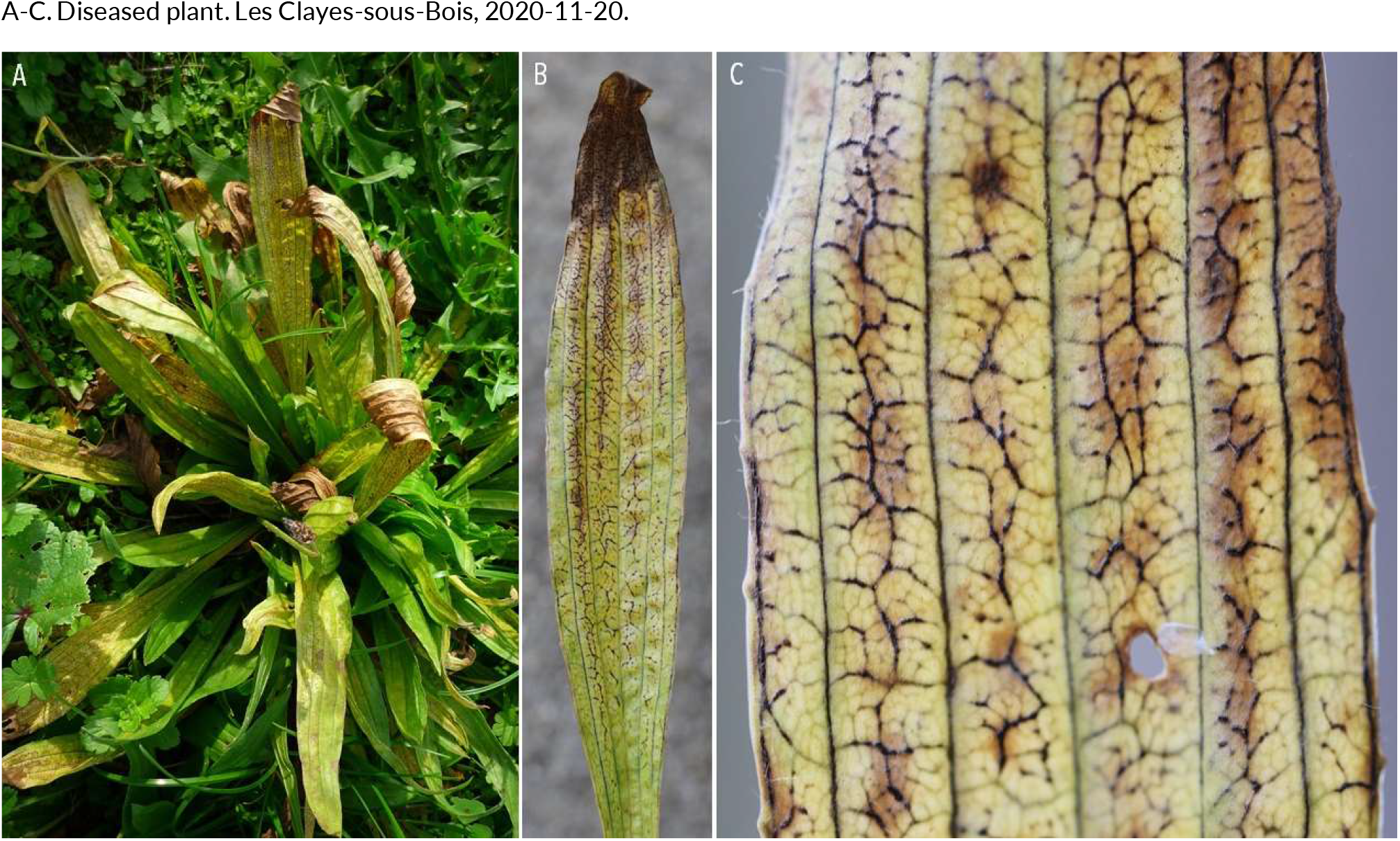
OTHER FUNGAL LEAF DISEASES | *Spilopodia nervisequa* ex *Plantago lanceolata* (ribwort plantain)

**180.**
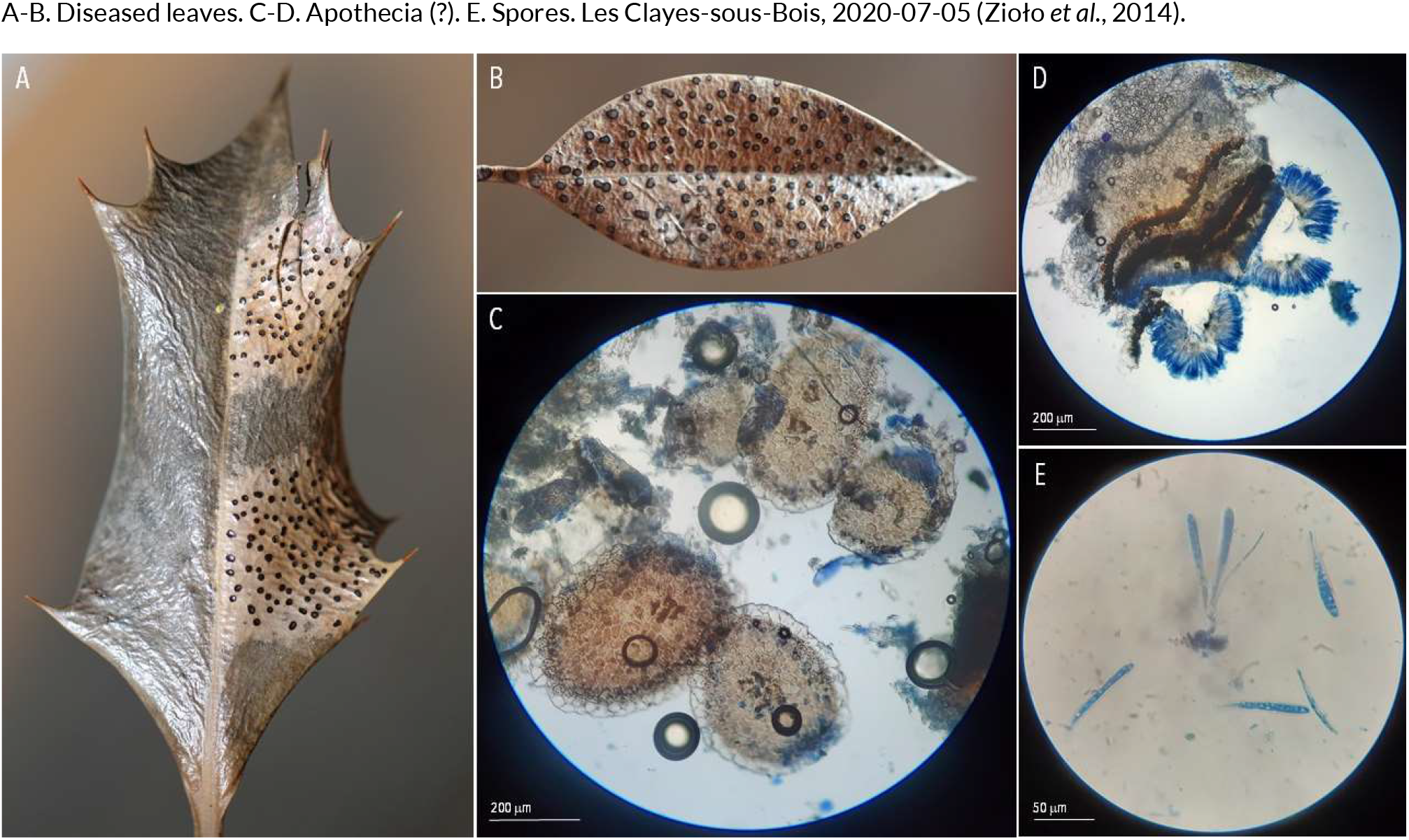
OTHER FUNGAL LEAF DISEASES | *Trochila ilicina* (?) *ex Ilex aquifolium* (English holly)

**181.**
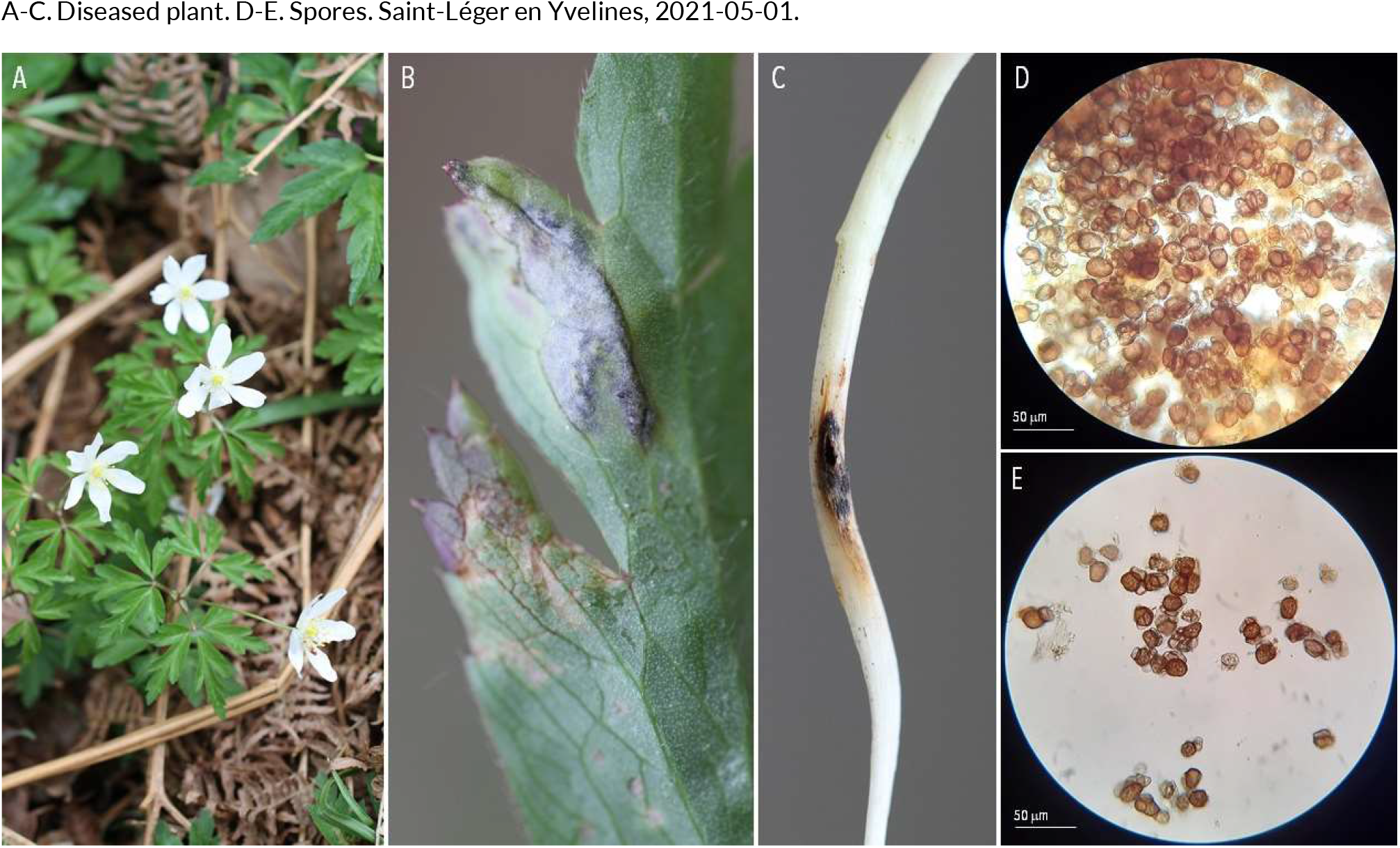
OTHER FUNGAL NON-LEAF DISEASES | *Urocystis anemones* ex *Anemone nemorosa* (wood anemone)

**182.**
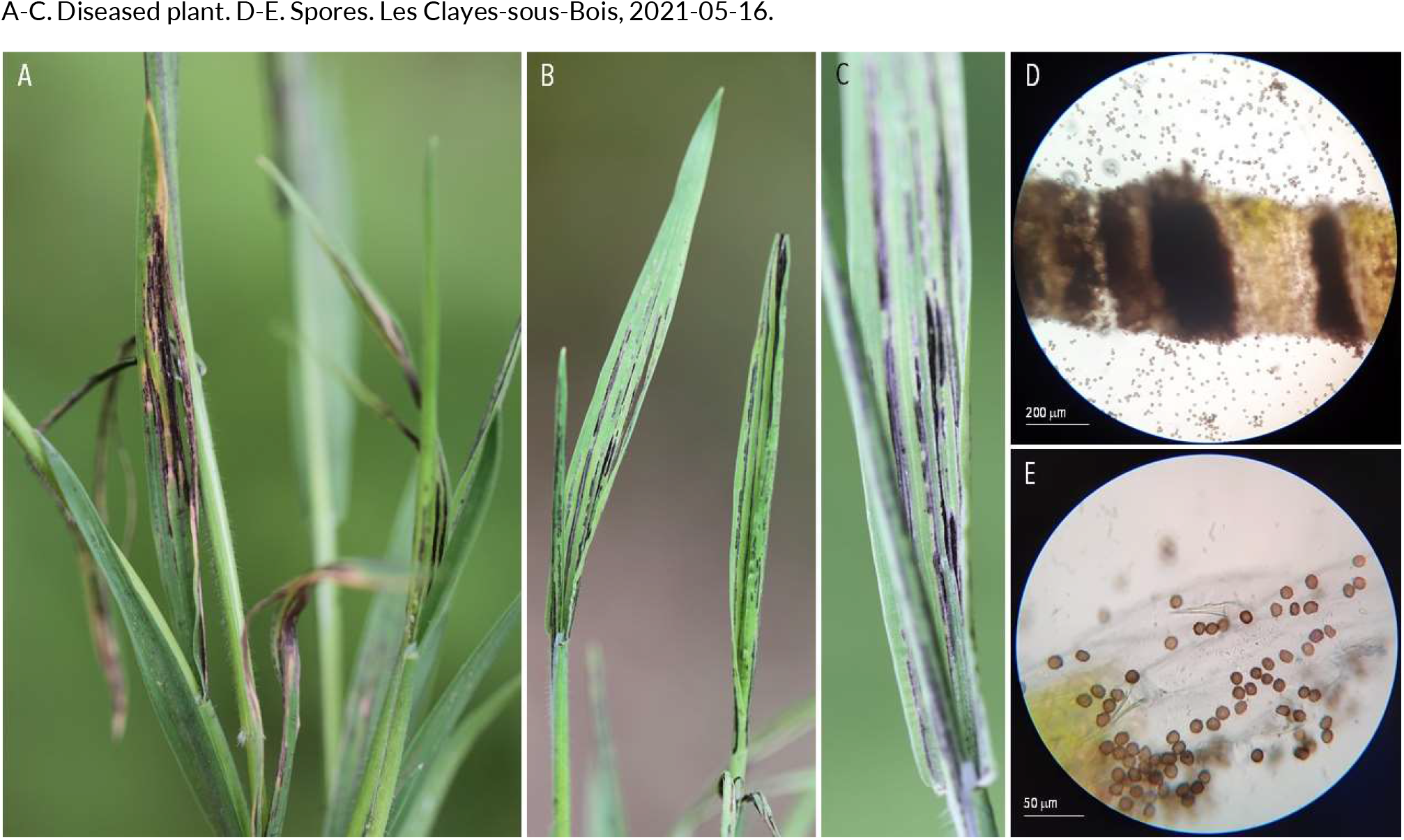
OTHER FUNGAL NON-LEAF DISEASES | *Ustilago salweyi* ex *Holcus lanatus* (common velvet grass)

**183.**
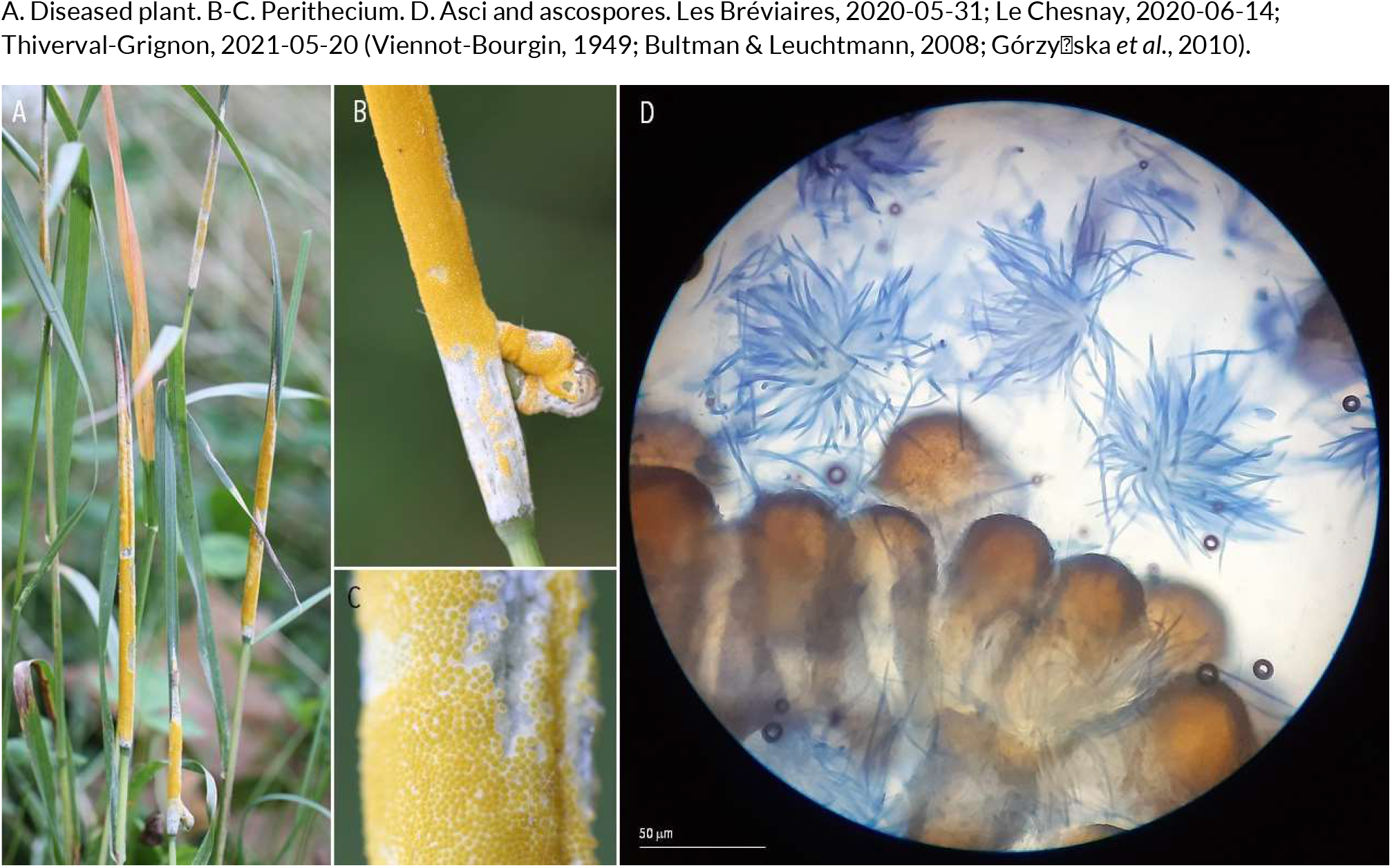
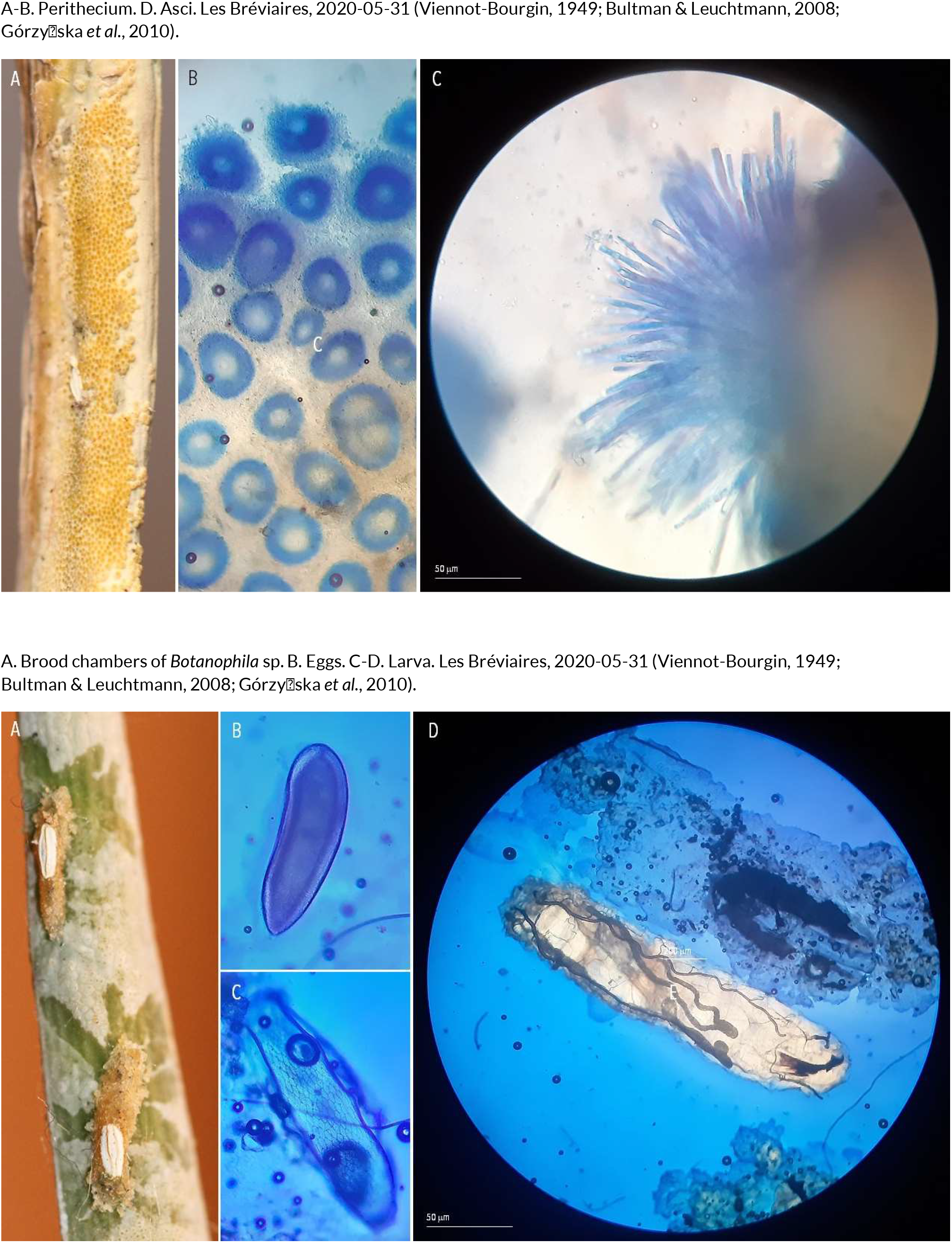
OTHER FUNGAL NON-LEAF DISEASES | *Epichloë typhina* ex *Dactylis glomerata* (cock’s-foot)

**184.**
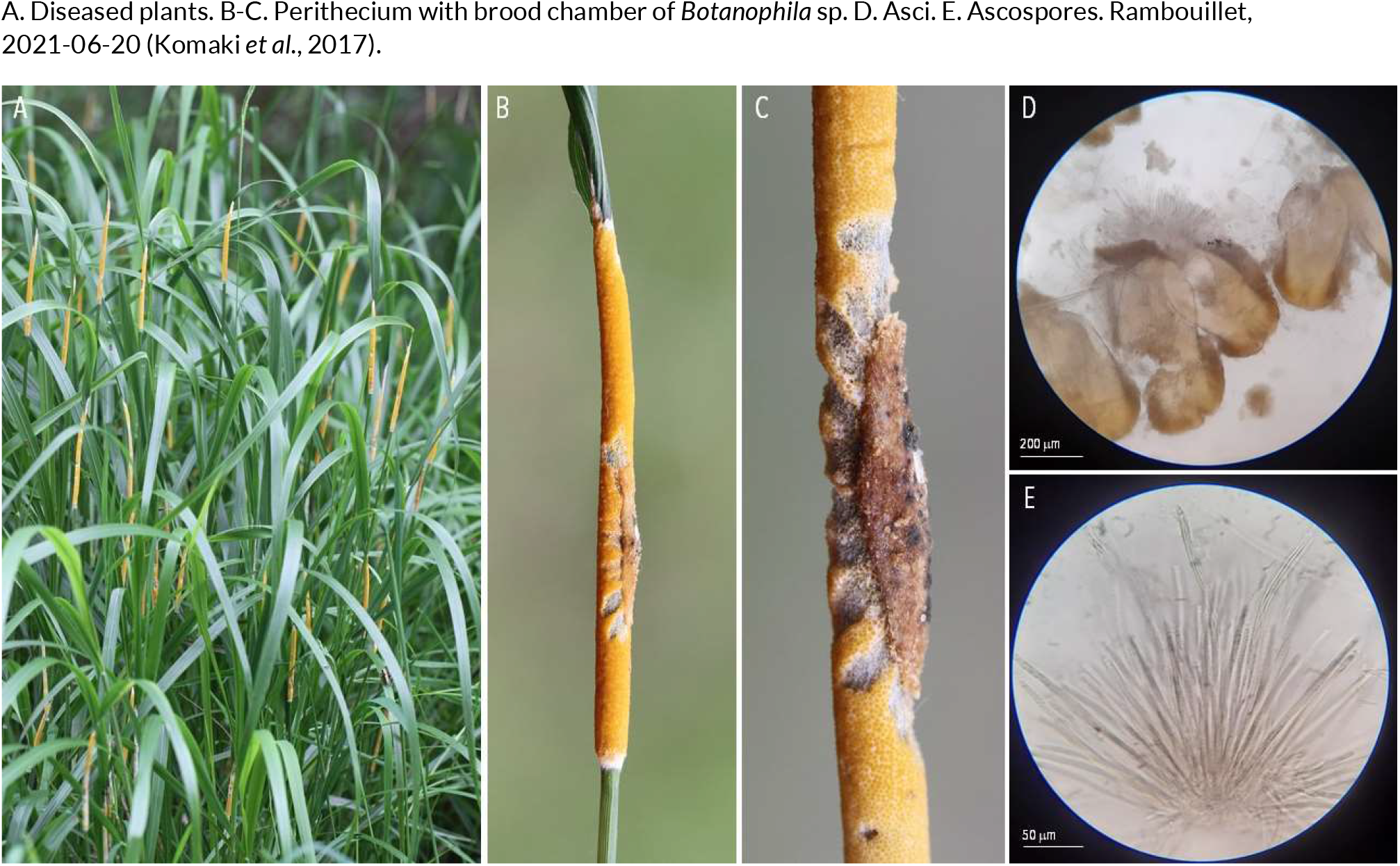
OTHER FUNGAL NON-LEAF DISEASES | *Epichloë sylvatica* ex *Brachypodium pinnatum* subsp. *rupestre* (torgrass)

**185.**
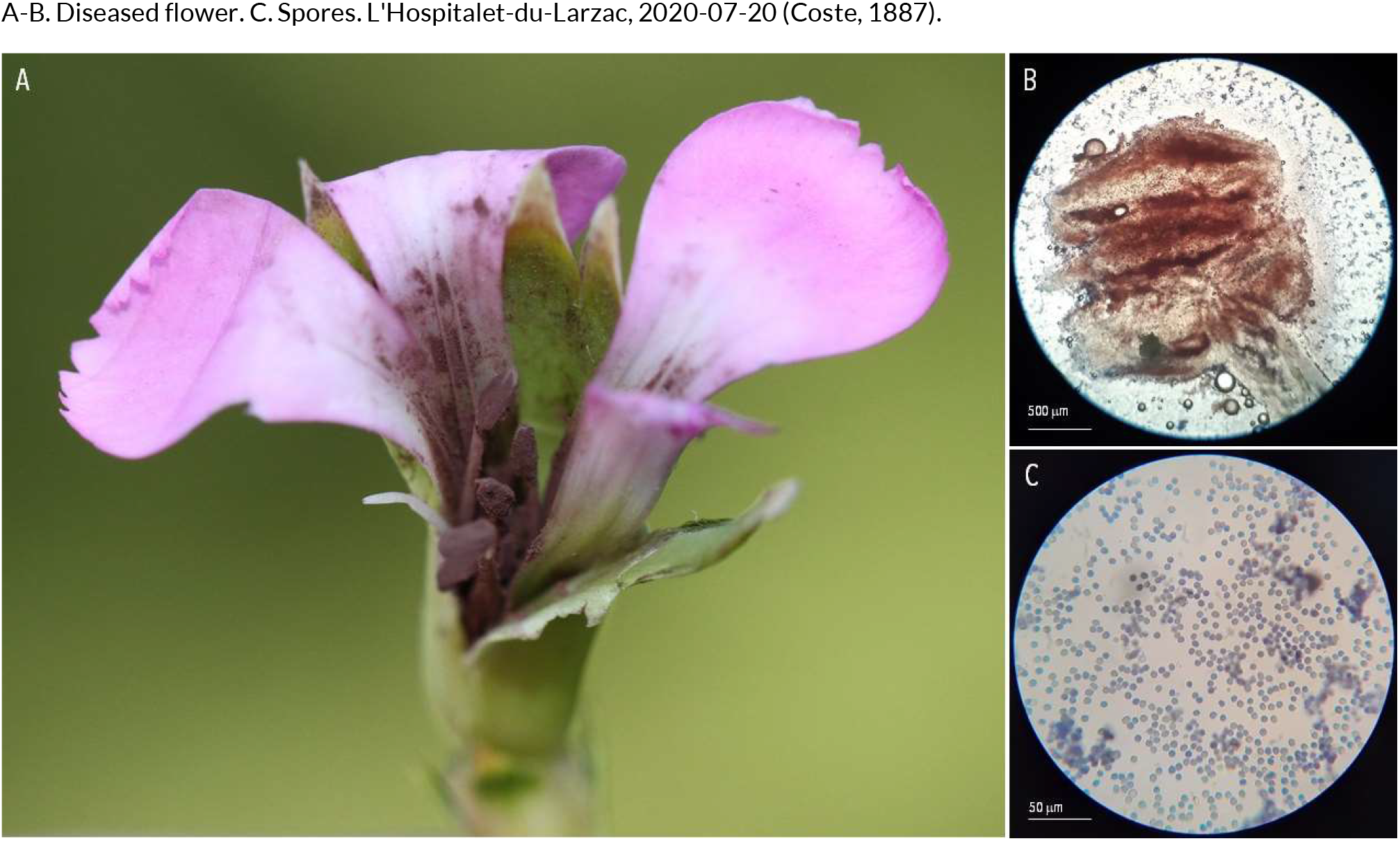
OTHER FUNGAL NON-LEAF DISEASES | *Microbotryum dianthorum* ex *Dianthus longicaulis*.

**186.**
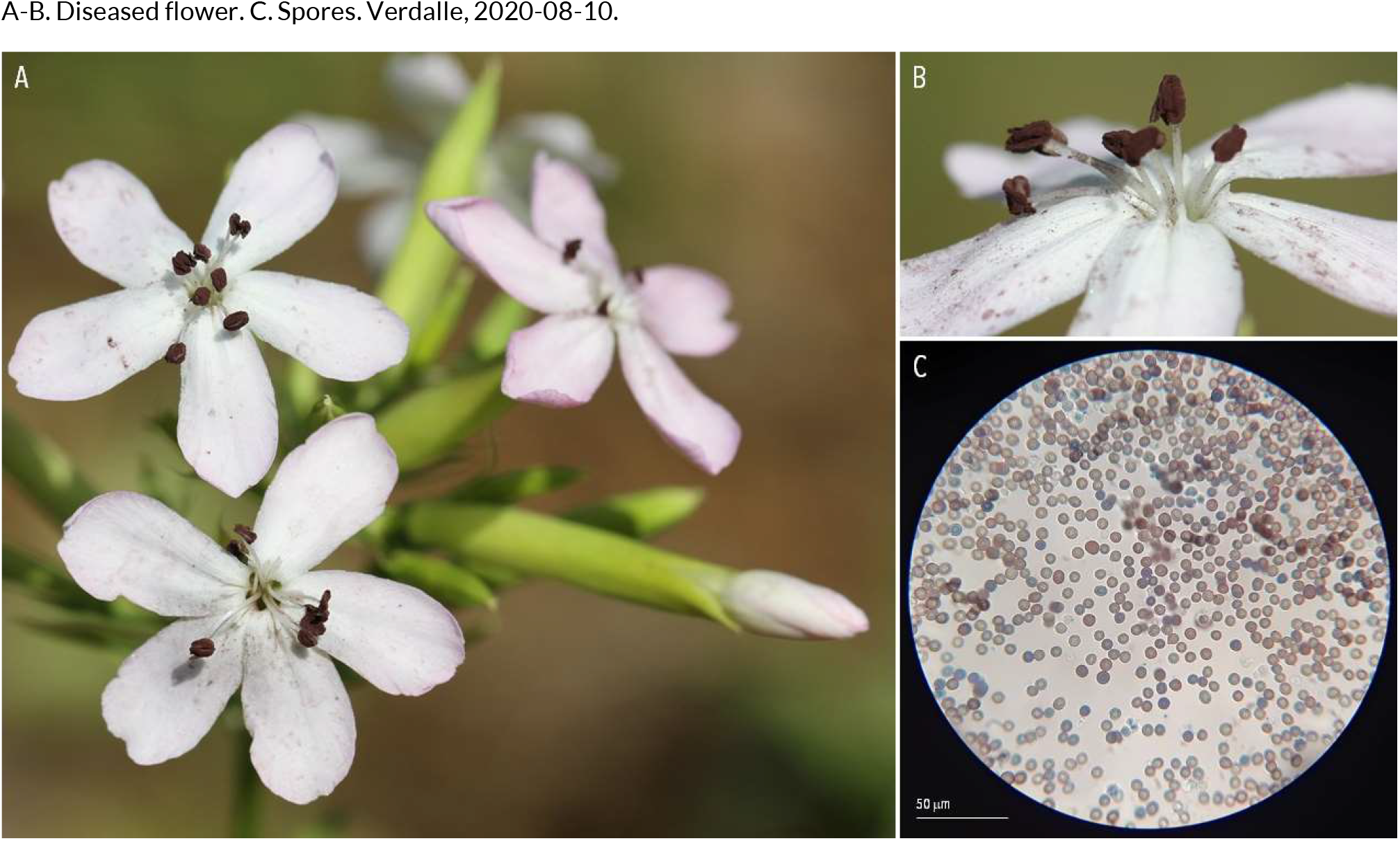
OTHER FUNGAL NON-LEAF DISEASES | *Microbotryum saponariae* ex *Saponaria officinalis* (soapwort)

**187.**
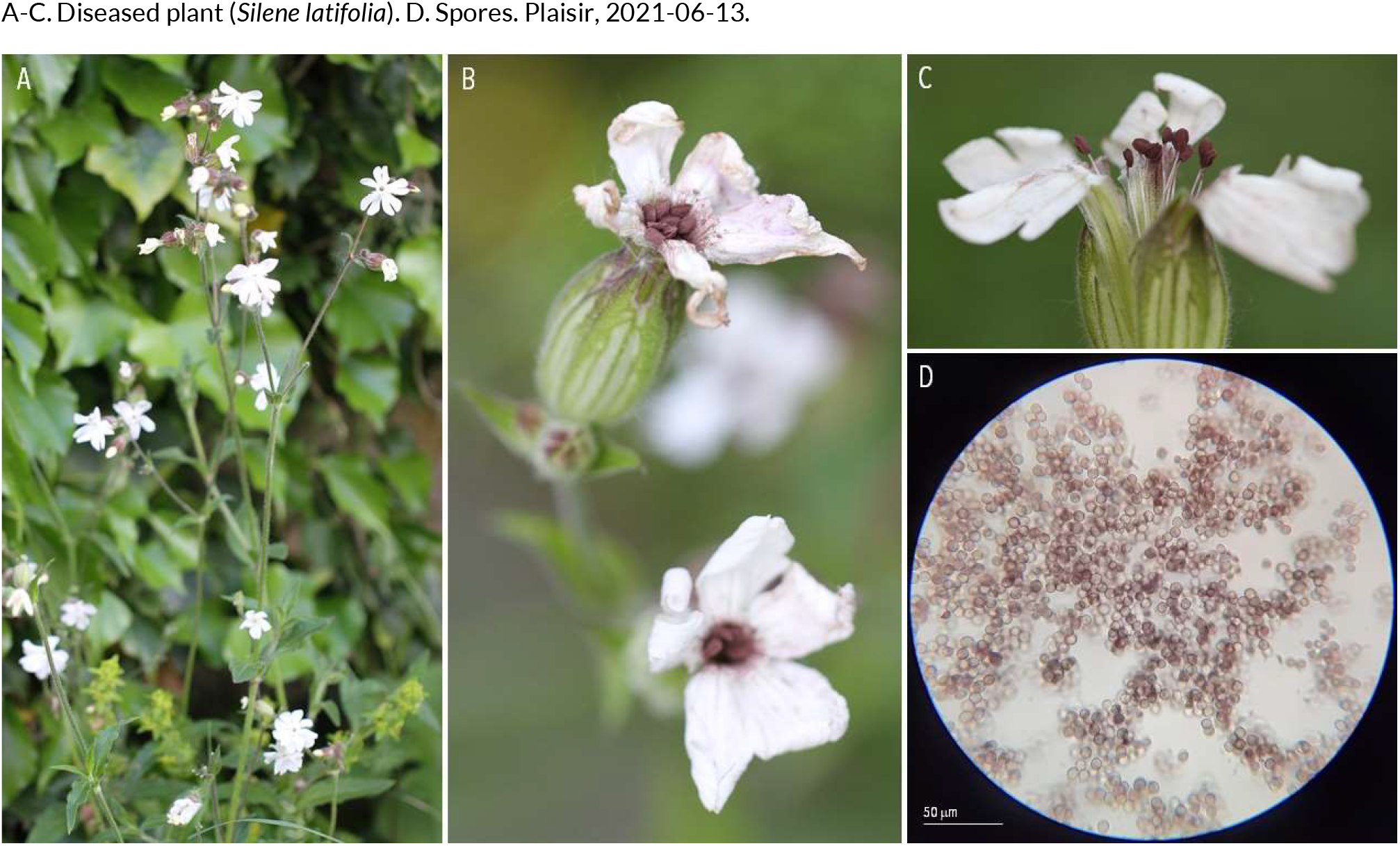
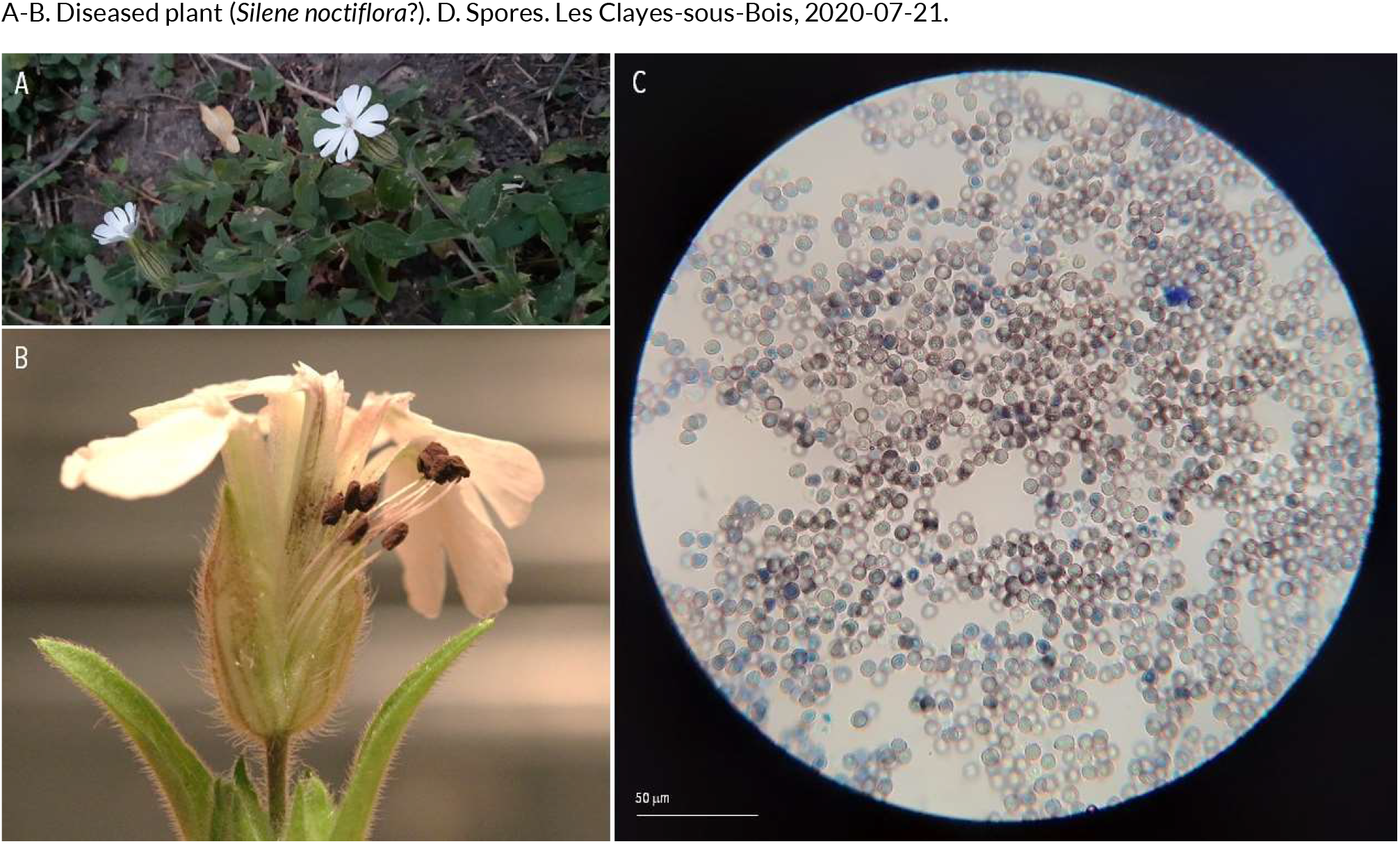
OTHER FUNGAL NON-LEAF DISEASES | *Microbotryum lychnidis-dioica* ex *Silene latifolia* (white campion) and *Silene noctiflora* (night-flowering catchfly)

**188.**
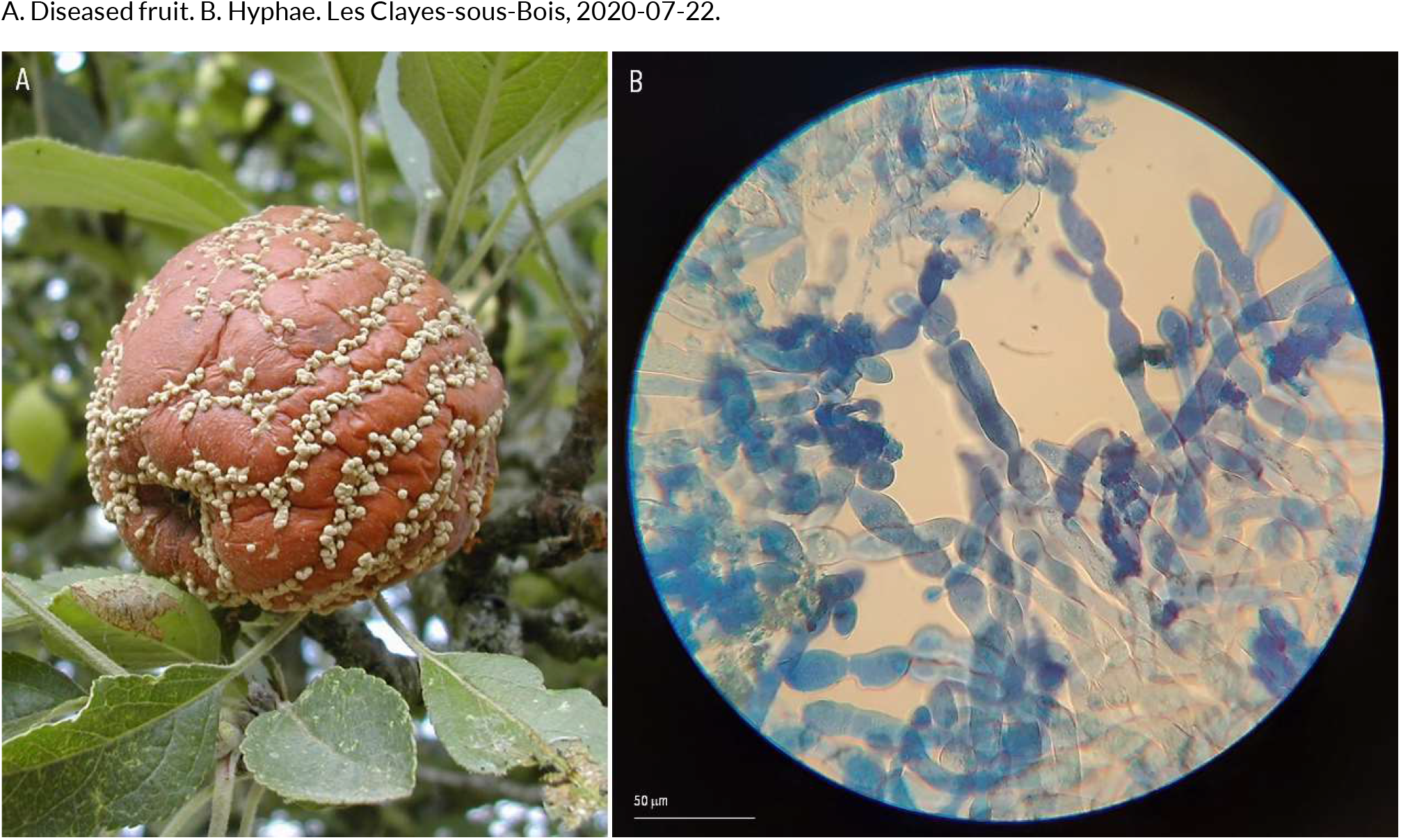
OTHER FUNGAL NON-LEAF DISEASES | *Monilia fructigena* ex *Malus domestica* (apple tree)

**189.**
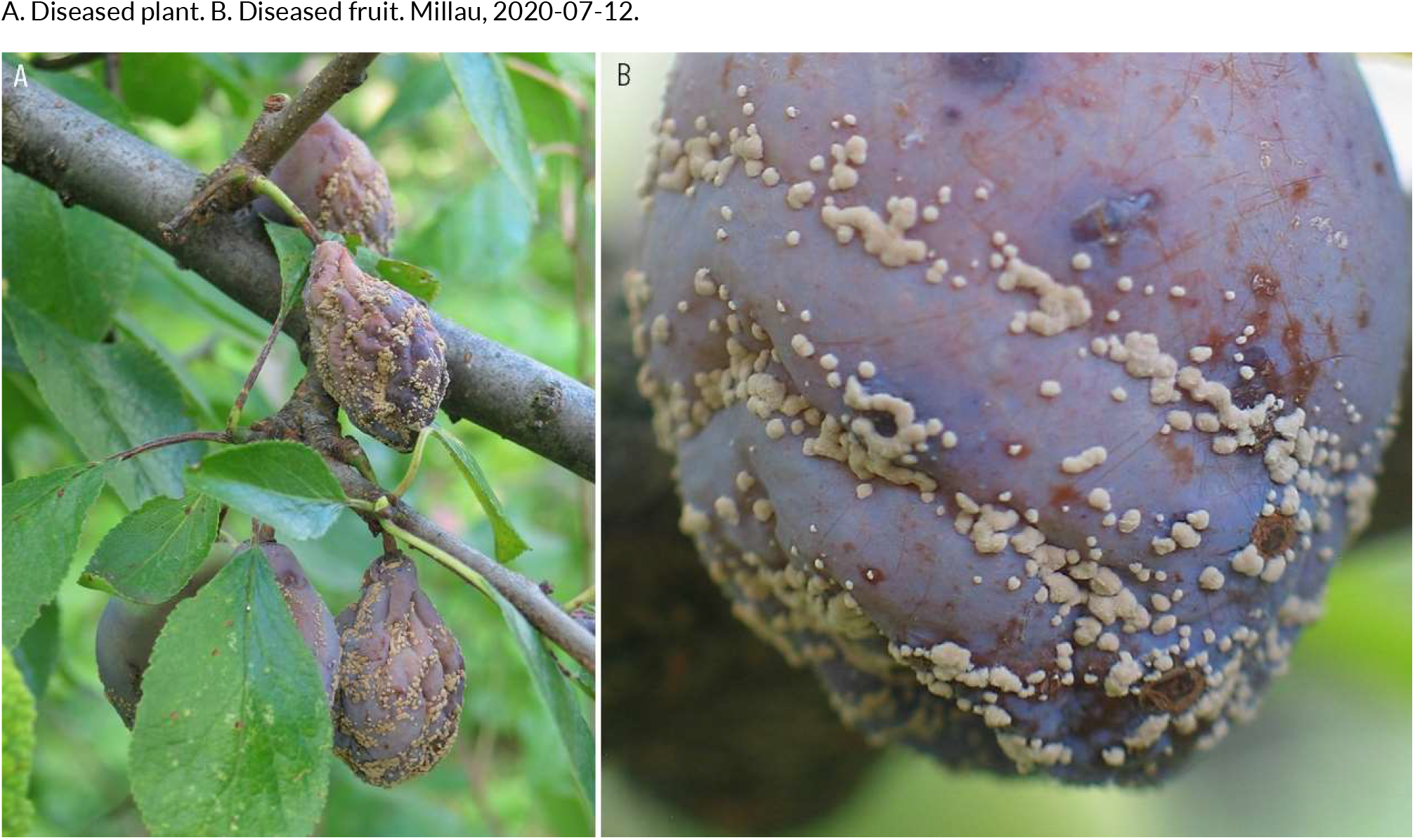
OTHER FUNGAL NON-LEAF DISEASES | *Monilia laxa* ex *Prunus domestica* (plum)

**190.**
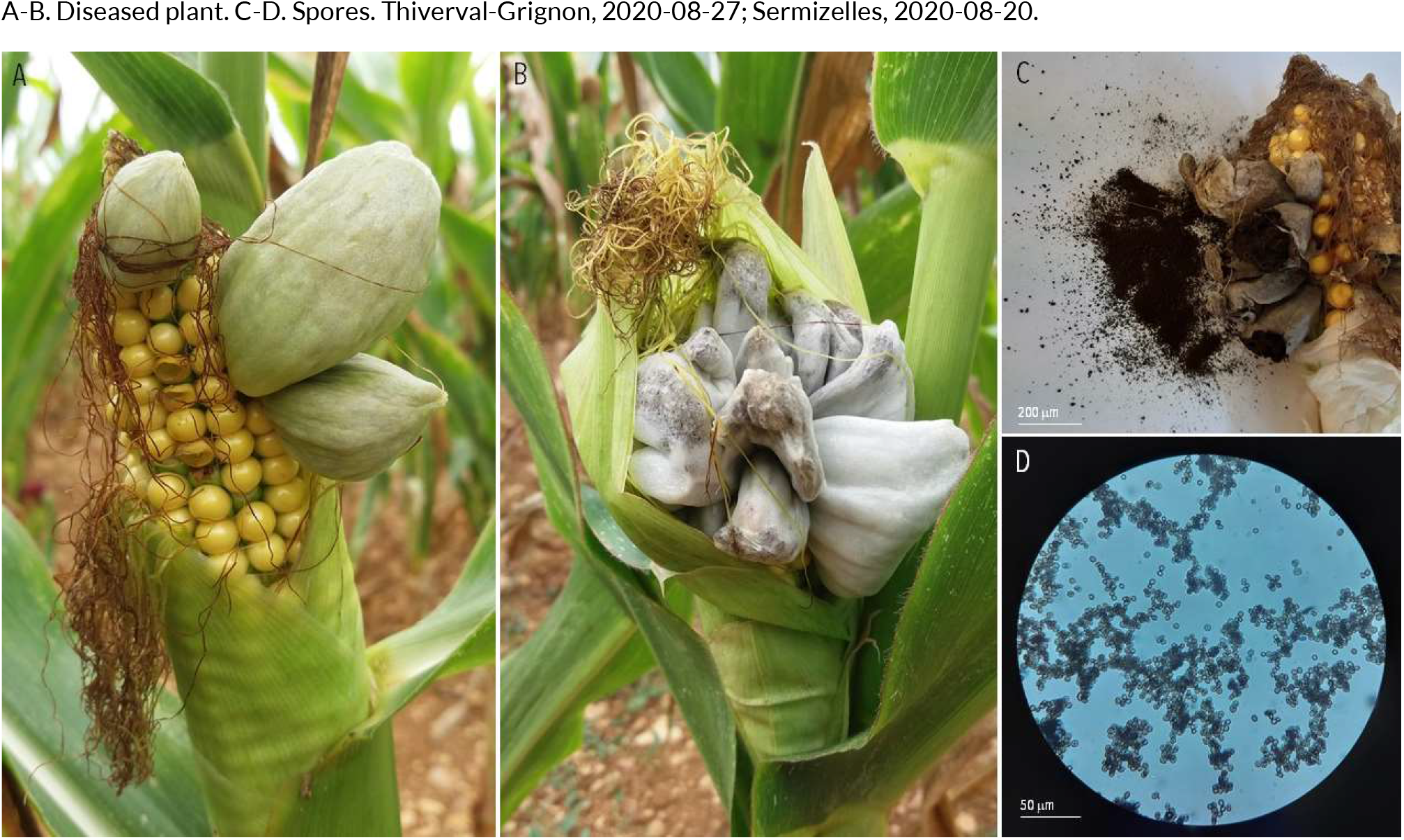
OTHER FUNGAL NON-LEAF DISEASES | *Ustilago maydis* ex *Zea mays* (maize)

**191.**
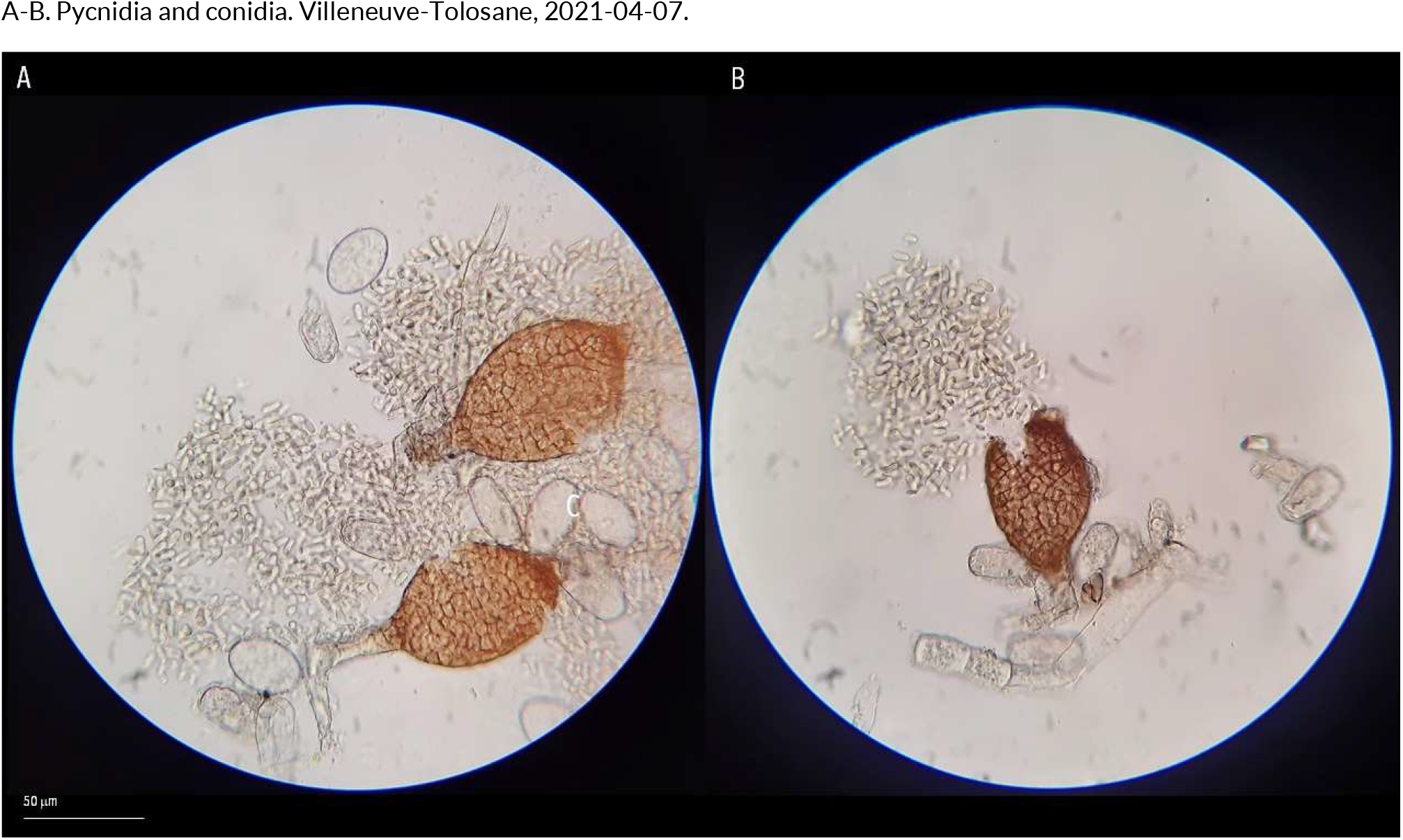
HYPERPARASITE | *Ampelomyces quisqualis* (hyperparasite of Golovinomyces cichoracearum) ex *Urospermum dalechampii* (smooth golden fleece)

**192.**
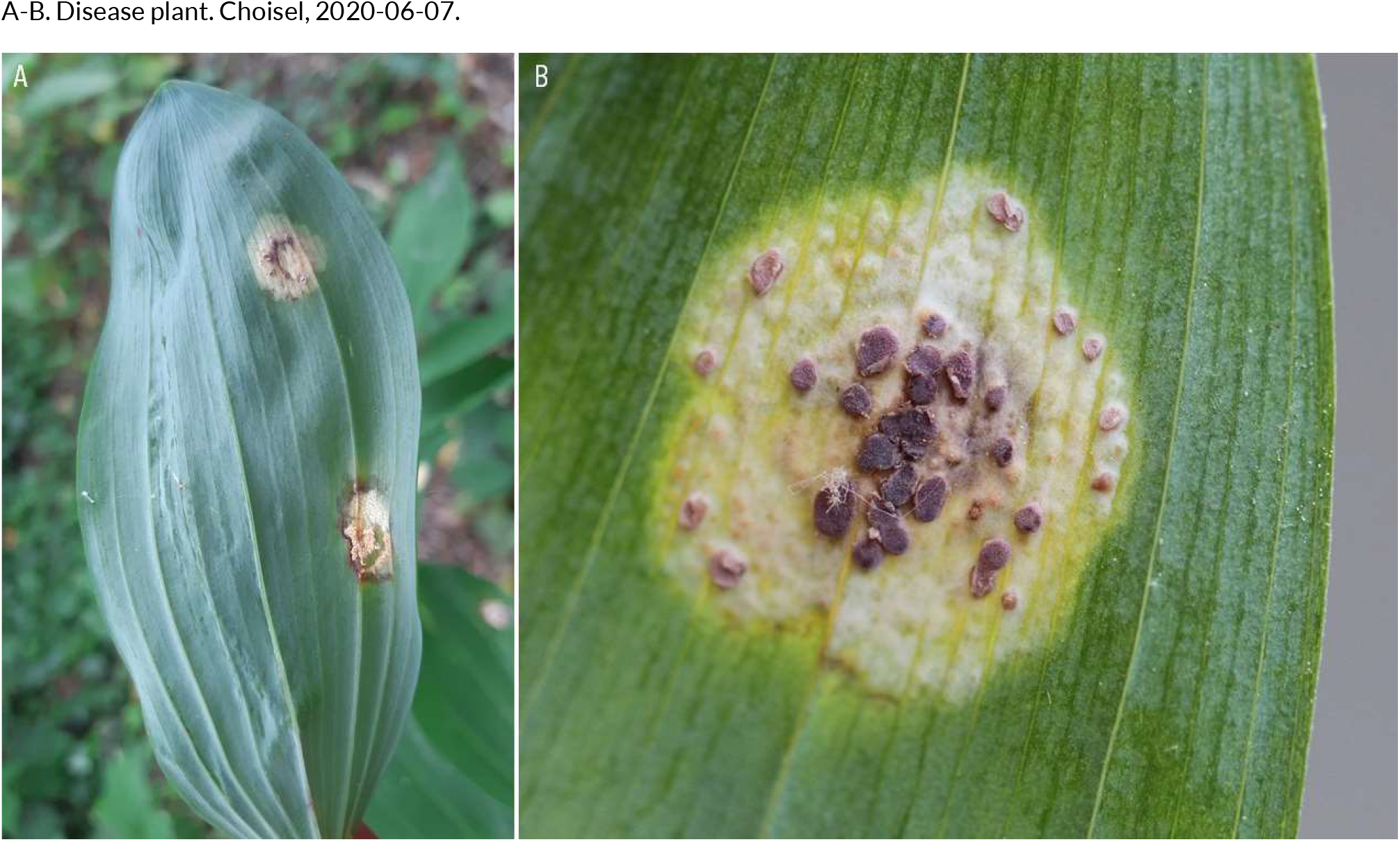
HYPERPARASITE | *Tuberculina persicina* (hyperparasite of Puccinia sessilis) ex *Polygonatum multiflorum* (Solomon’s seal)

**193.**
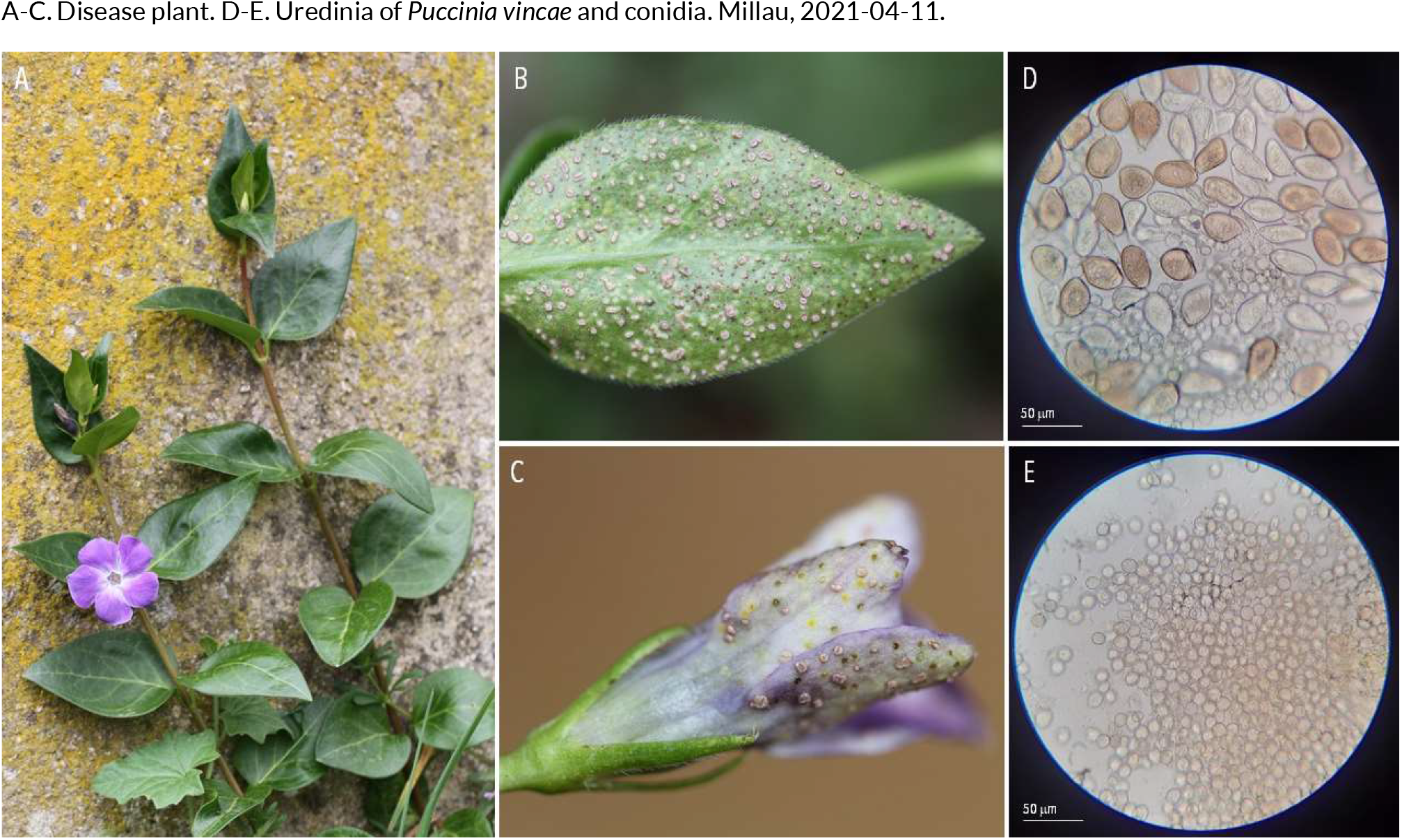
HYPERPARASITE | *Tuberculina sbrozzii* (hyperparasite of Puccinia vincae) ex *Vinca major* (bigleaf periwinkle)

**194.**
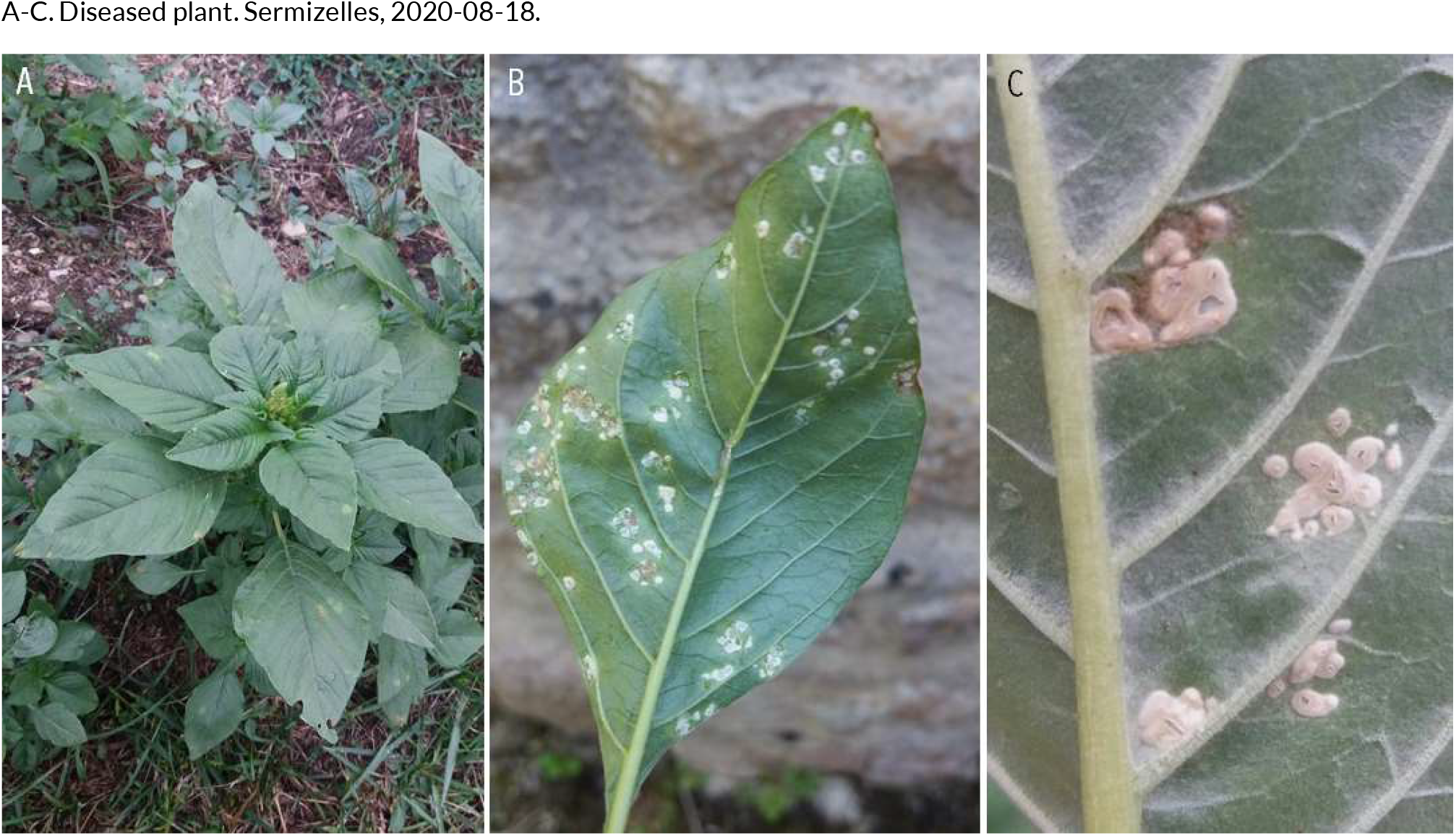
NON-FUNGAL DISEASES | *Albugo bliti* ex *Amaranthus retroflexus* (redroot pigweed)

**195.**
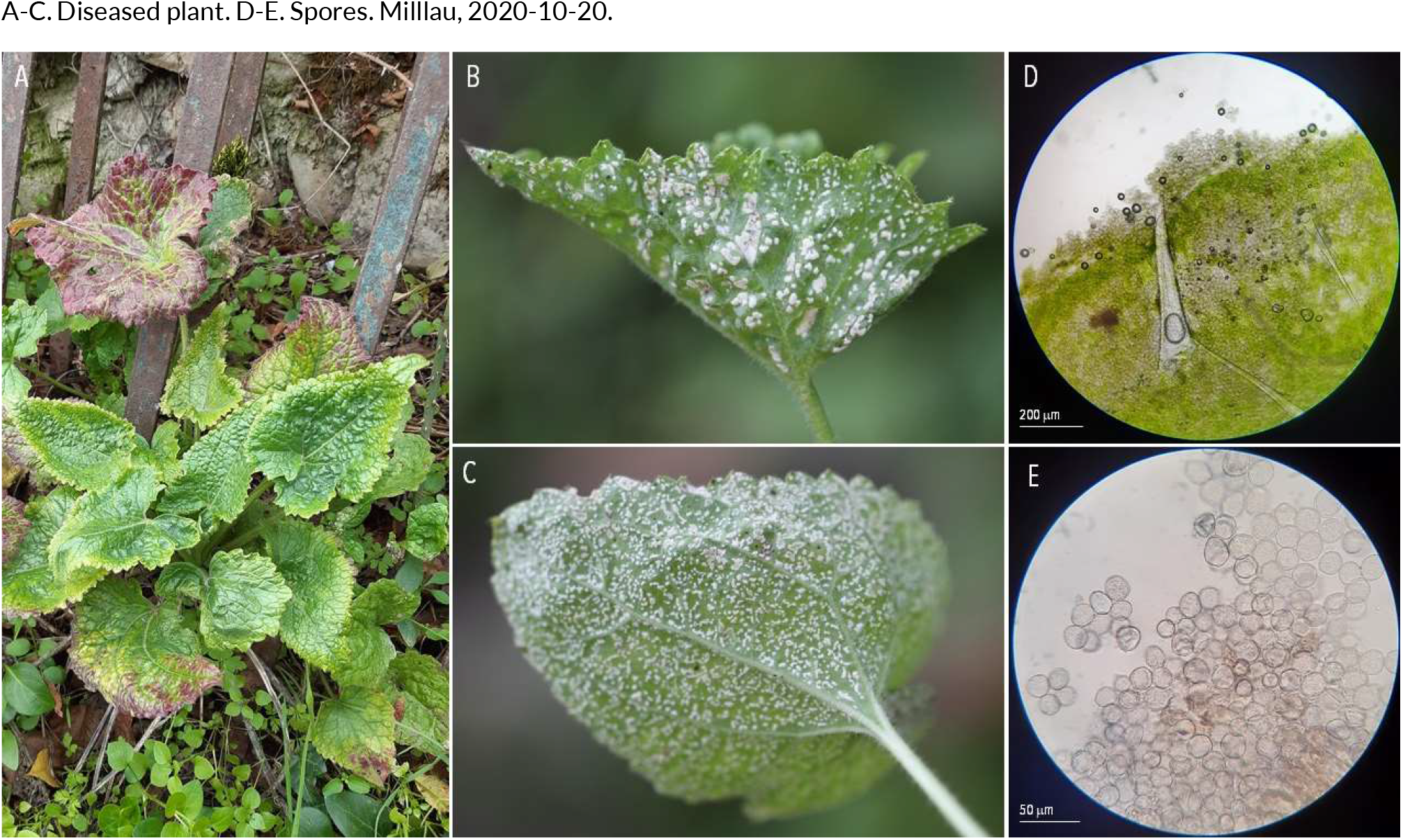
NON-FUNGAL DISEASES | *Albugo candida* ex *Lunaria annua* (honesty)

**196.**
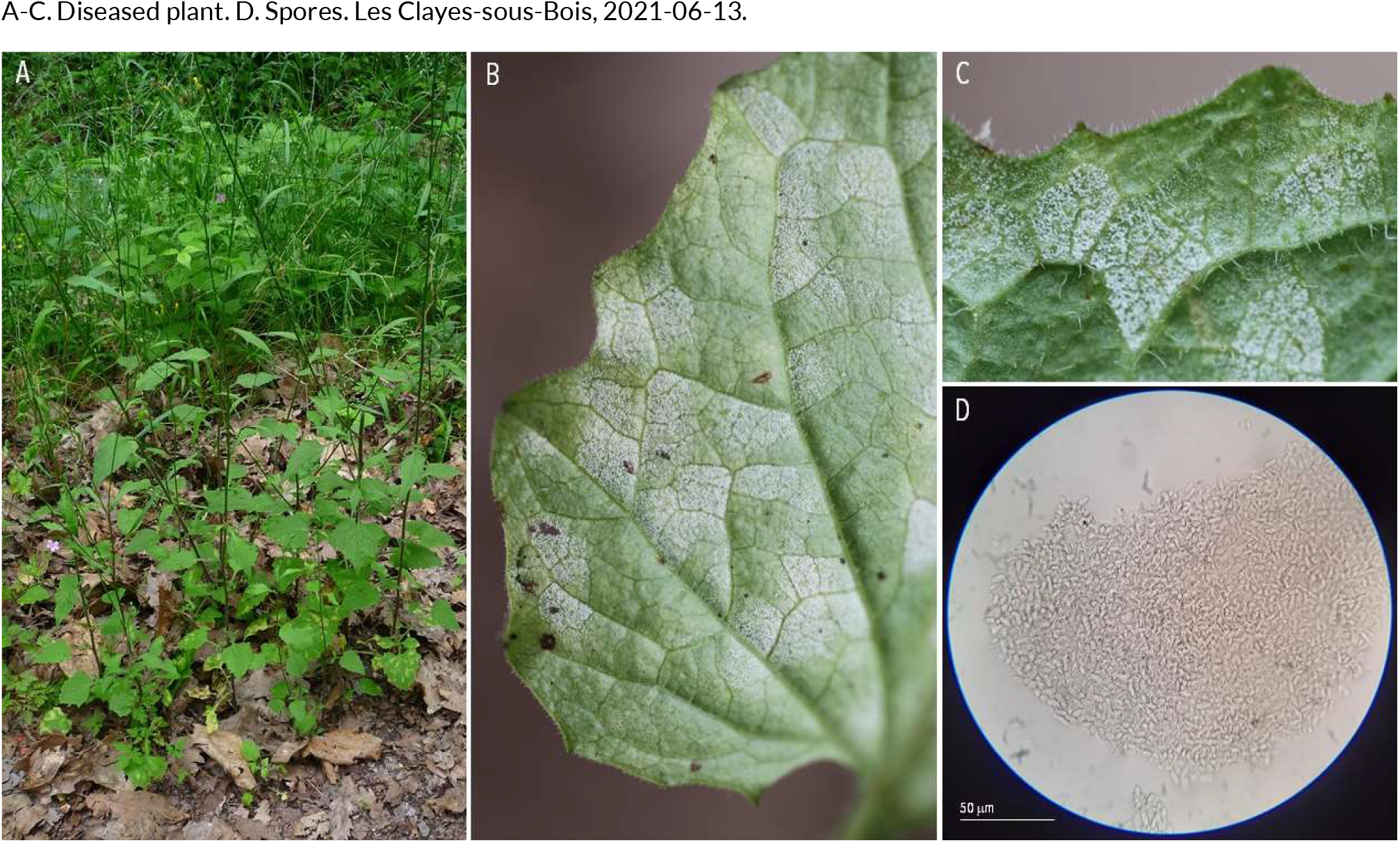
NON-FUNGAL DISEASES | *Bremia lapsanae* ex *Lapsana communis* (nipplewort)

## Appendix 2

Example of conversations on Twitter (single tweets, threads, and the reactions that they have induced).

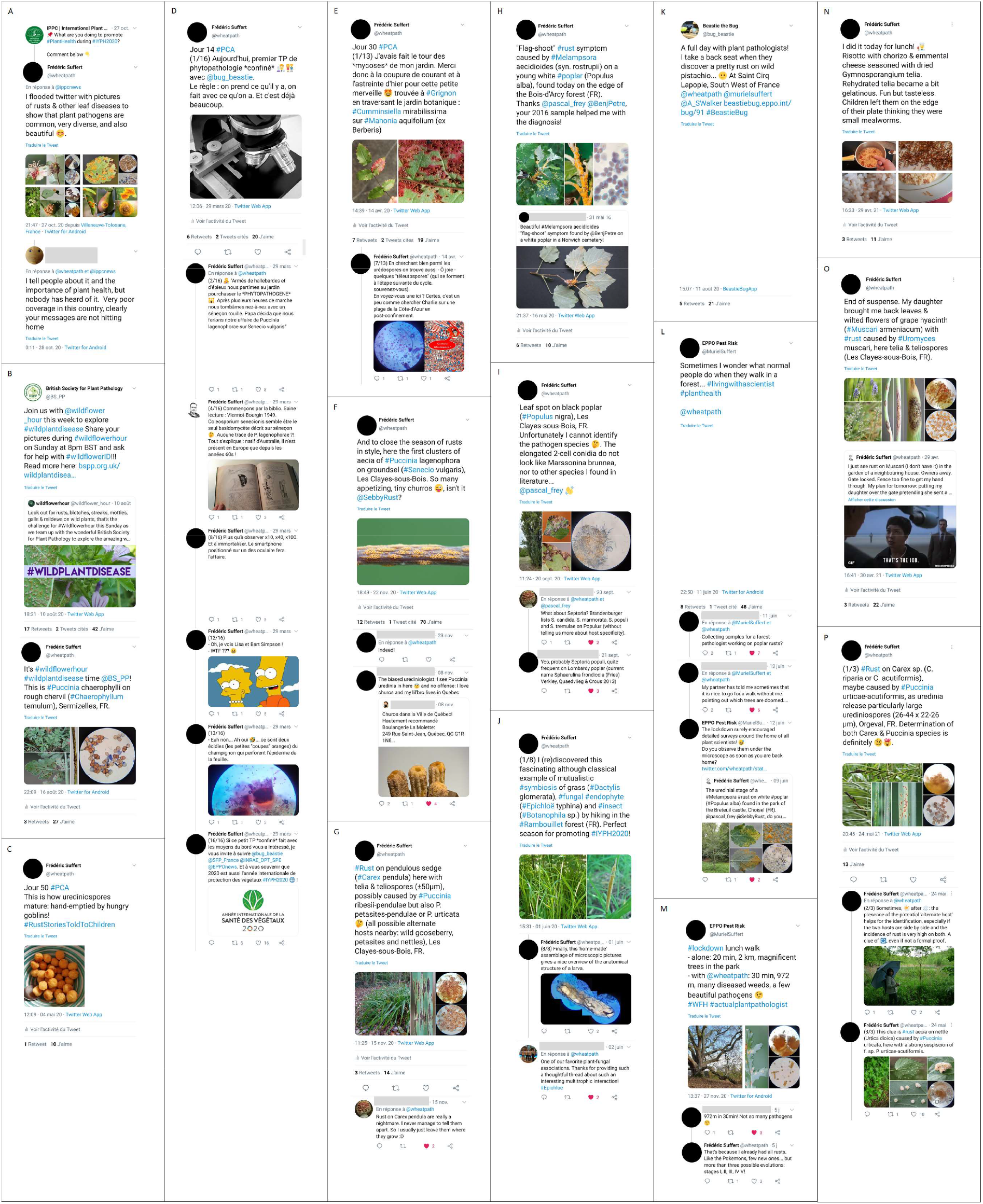

